# Isolation and characterization of *Pseudomonas chlororaphis* strain ST9 and its potential as a bioinoculant for agriculture

**DOI:** 10.1101/2020.12.23.424151

**Authors:** Iris Bertani, Elisa Zampieri, Cristina Bez, Andrea Volante, Vittorio Venturi, Stefano Monaco

## Abstract

The development of biotechnologies based on beneficial microorganisms for improving soil fertility and crop yields could help addressing many current agriculture challenges, such as food security, climate change, pests control, soil depletion while decreasing the use of chemical fertilizers and pesticides. Plant Growth Promoting (PGP) microbes can be used as probiotics in order to increase plant tolerance/resistance to abiotic/biotic stresses and in this context strains belonging to the *Pseudomonas chlororaphis* group have shown to have potential as PGP candidates. In this work a new *P. chlororaphis* isolate is reported and tested for (i) *in vitro* PGP features, (ii) whole genome sequence analysis, and (iii) its effects on root microbiome, plant growth and on the expression of different plant genes in greenhouse experiments. The potential use of this *P. chlororaphis* strain as a plant probiotic is discussed.

## Introduction

Agriculture of the 21st century has several challenges to face. Among them are the increase in population and the growing demand for food in the context of climate change, soil depletion, competition between different land uses and the need to reduce chemical fertilizers and pesticides. The use of beneficial microorganisms in crop production is an attractive solution as they can have beneficial effects on soil and plant by improving soil fertility, plant yields and reducing the use of agrochemicals (Hayat *et al*., 2010). The development of next-generation sequencing methodologies has led to numerous studies on plant microbiomes documenting that plants are colonized and live in association with a large number of microorganisms. Many of these, indicated as PGP (Plant Growth Promoting) microbes, play important roles in plant health and resistance to biotic and abiotic stresses (Berendsen *et al*., 2012; Bulgarelli *et al*., 2013; Backer *et al*., 2018; Orozco-Mosqueda *et al*., 2018; Khatoon *et al*., 2020). PGP microbes allow the reduction of fertilizers requirements, improving crop nutrient use efficiency (nitrogen, phosphate, etc.), and protecting the host plant by pathogens invasion through niche exclusion mechanisms and antibacterial/antifungal compound production (Bender *et al*., 2016; Pii *et al*., 2015). PGP microorganisms can also elicit transcriptional changes in hormone-, defence- and cell wall-related genes (Spaepen *et al*., 2014), increase root length (Hong *et al*., 1991), and activate auxin-response genes that reinforce plant growth (Ruzzi and Aroca, 2015). In the last decades the interest in developing PGP probiotic microorganisms has increased with the goal to reduce chemical fertilization (Mehnaz, 2016) and pesticide use, both in conventional and organic farming, with the aim of offering healthier food and improving sustainability of crop production (Hazra *et al*., 2018). One class of microorganisms that are being studied for many years for their application are Plant Growth Promoting Rhizobacteria (PGPR) (Oleńska *et al*., 2020).

Among the group of PGPR, strains belonging to the *Pseudomonas chlororaphis* species have been found in association with a wide range of plants, both mono- and dicotyledonous, and both wild and cultivated (Biessy *et al*., 2019). *Pseudomonas chlororaphis* is currently classified as four subspecies, namely *chlororaphis, aureofaciens, aurantiaca*, and *piscium* (Peix *et al*., 2007; Burr *et al*., 2010). Several strains of *P. chlororaphis* have shown good potential for application as plant probiotics (Zhao *et al*., 2013; Chen *et al*., 2015; Arrebola *et al*, 2019; Anderson *et al*., 2020) due to their rhizosphere colonization abilities and plant associated beneficial phenotypes such as chemotaxis and motility (Arrebola *et al*., 2020), biofilm formation (Calderon *et a*l., 2013, 2014), P solubilization (Ahemad, 2015), ACC deaminase (Glick, 2014), IAA production (Kang *et al*., 2006; Oh *et al*., 2013) and biocontrol. *P. chlororaphis* strains produce different antifungal compounds such as Prn (pyrrolnitrin), PCN (phenazine-1-carboxamide), PCA (phenazine-1-carboxylic acid), 2-OH-PHZ (2-hydroxyphenazine), HPR (2-hexyl-5-propyl-alkylresorcinol) and HCN (hydrogen cyanide). These molecules inhibit the growth of various phytopathogens belonging to the *Fusarium* group (Chin-A-Woeng *et al*., 1998; Hu *et al*., 2014) and to different species of *Colletotrichum, Phytophthora, Pythium, Sclerotinia* and *Rhizoctonia* (Liu *et al*., 2007), protecting plants such as maize (Tagele *et al*., 2019), tomato (Zhang *et al*., 2015). In addition, good product formulation protocols have been developed and commercialized products having *P. chlororaphis* strains have been produced (Nam *et al*., 2018).

In this work, we report the isolation and characterization of *P. chlororaphis* strain ST9. This strain was able to colonize and persist in rice roots for the entire rice vegetative cycle and studies are presented of its effect on the root microbiome, on plant growth and on the expression of several plant genes. The potential use of this *P. chlororaphis* strain as a plant probiotic for rice cultivation is discussed.

## Material and Methods

### Strain isolation, growth and identification

One gram of bulk soil from Padriciano, Trieste, Italy (45° 39’ 32” North, 13° 50’ 28” East) was taken and resuspended in 5 ml of PBS (phosphate buffer solution). Serial dilutions were performed and plated on 1/6 Tryptic Soy medium (BD, Maryland, 21152. USA), solidified with 1.5% agar. Growth was allowed for 72 hrs at 28°C and single colonies were isolated and purified through further streaking. Several isolates displaying varying morphologies, were selected and identified. Identification occurred through 16S rRNA sequencing: briefly, primers fD1 and rR2 were used for the amplification of the 16S rRNA gene and primers 518F and 800R (Supplementary Table (ST) ST1) were used for its sequencing (Eurofins, Ebersberg, Germany). A small bacterial collection was established and *P. chlororaphis* ST9 was one of the isolates stored at −80°C for further studies.

### *In vitro* phenotypic characterization

Presence of lipolytic and proteolytic activities was determined by streaking the bacterial isolates on 1/6 TSA (Tryptic Soy Agar) medium amended with 1% Glyceryl tributyrin (Smeltzer *et a*l., 1992) and 2% of powder milk (Huber *et al*., 2001), respectively. Exopolysaccharide (EPS) production was tested by streaking the bacterial isolates on Yeast Extract Mannitol medium (Zlosnik *et al*., 2008) and indole acetic acid (IAA) production was verified as described by Bric *et al*. (1991) using the Salkowski reagent. The 1-aminocyclopropane-1-carboxylic acid (ACC) deaminase activity was detected by means of M9 minimal medium with ACC as unique N source (Penrose *et al*., 2003), while the ability to solubilize P was verified using the NPRBB growth medium (Nautiyal, 1999). N-acyl homoserine lactone (AHL) production was assessed by T-streak technique, using the biosensor *Chromobacterium violaceum* CV026 after incubation for 1-2 days (Steindler and Venturi, 2007). Motility was checked on M8 medium plates with 0.3% (swimming) or 0.5% (swarming) agar (Kohler *et al*., 2000). Anti-bacterial and anti-fungal activities were tested by plating different plant pathogens (*Dickeya zeae, Pseudomonas fuscovaginae, Magnaporthe oryzae, Fusarium* sp., *Aspergillus* sp. and *Phytophora infestans*) adjacent to a 16 hrs old streak of ST9 on TSA medium: pathogens growth was checked after 72-96 hrs. Plant growth promotion traits were tested on 300 surface sterilized rice seeds. In detail, 16 hrs pre-germinated seeds were submerged for 2 hrs in a ST9 bacterial suspension (OD_600_ 0.5) or PBS, as control; after inoculum application seeds were rinsed with sterile water and allowed to germinate in the dark at 30°C. One hundred seeds were evaluated for emergence after 3 days, 100 seeds were grown for 10 days and then used to measure coleoptile length and the growth of the last 100 seeds was interrupted after 12 days for dry mass weight.

### Generation of a rifampicin resistant spontaneous mutants

The *P. chlororaphis* ST9 strain was grown in 1/6 TS medium for 16 hrs at 30°C with 100 rpm shaking. A 1/100 dilution of the overnight culture was put in 1/6 TS medium with 15 µg/ml rifampicin and the culture was grown again in the same growth conditions. The procedure was repeated, increasing concentration of Rifampicin (25, 50 and 100 µg/ml), and the final culture was plated on 1/6 TS. One colony was chosen and streaked on TSArif100; its identity, *P. chlororaphis* ST9, was confirmed again by 16S rRNA sequencing.

### Bacterial genomic DNA extraction and sequencing

Genomic DNA extraction for genome sequencing was performed with the pronase Sarkosyl lysis method (Better *et al*., 1983). 2 µg of DNA, quantified through Nanodrop (Thermo Scientific, Waltham, Massachusetts, USA) and checked by gel electrophoresis, were used for genome sequencing by the Exeter Sequencing Service (University of Exeter, Exeter, UK). The sequence was carried out using the Illumina technique with the Hiseq 2500 platform with the 125 base pair paired end system. Reads were assembled using SPAdes 3.9.03 (Bankevich *et al*., 2012). The assembled *P. chlororaphis* ST9 genome was uploaded in the RAST Annotation Server (Overbeek *et al*., 2014) and was automatically annotated using the RASTtk annotation scheme (Brettin *et al*., 2015). The genome sequence is presented as a unique contig and it is available on the RAST server (http://rast.nmpdr.org; “guest” as login and password) under Genome ID 286.2086.

### *P. chlororaphis* ST9 taxonomy

Multi-locus Sequence Analysis (MLSA) was performed using 5 housekeeping genes: *recA* (recombinase A), *gyrB* (DNA gyrase subunit B), *rpoD* (RNA polymerase sigma factor), *carA* (Carbamoyl-phosphate synthase small chain) and *atpD* (ATP synthase subunit beta), plus the 16S rRNA locus. Sequences of these loci were obtained from the ST9 genome sequence. For the phylogenetic analysis, we concatenated the gene sequences in the following order: 16S-*recA-gyrB-rpoD-carA-atpD* resulting in a single sequence. For the same genes, orthologue sequences from 16 *P. chlororaphis* species were obtained from the NCBI database and chained. The used sequences are reported in Supplementary Materials SM1. As outgroup, we used the orthologue concatenate from *Pseudomonas putida KT2440 and P. fluorescens Pf-01*. The phylogenetic analysis was performed using the NGPhylogeny.fr public platform (Lemoine *et al*., 2019).

### Plant inoculation

Seeds of *O. sativa* L. cv. Baldo were surface-sterilized with 50% sodium hypochlorite solution (commercial bleach) for 60 min, rinsed with sterile water and incubated in a wet dark environment for seven days at 30°C for germination. Two growth conditions (treatments) were considered: with and without the ST9 inoculation. The roots of 70 seedlings were soaked for 60 min in a ST9 RifR bacterial solution at 0.5 of OD_600_ (treated, “T” plants), while the control was performed soaking the roots of 70 seedlings in a sterile PBS solution (untreated, “UT” plants). The inoculated and the control seedlings were transferred to plastic tubes containing 0.4% water-agar and Hoagland solution for 72 hrs and then transplanted into pots in greenhouse.

### Greenhouse experiments

For each of the two treatments, fourteen plastic pots (23 cm x 21 cm) were used, filled with non-sterile paddy field soil (47.8% sand, 9.4% clay, 42.8% silt, pH 6.4, organic matter 1.45%) taken from the experimental rice field of CREA-CI in Vercelli (VC, Italy). Pots were placed in greenhouse under uncontrolled temperature, light and humidity parameters. In each pot, five seedlings, undergoing the same treatment, were sown. Plants were watered every day with tap water and kept in a greenhouse for 90 days, following the natural photoperiod. Four time points were considered: T1, 10 days post inoculation (dpi); T2, 28 dpi; T3, 40 dpi and T4, 90 dpi. At T1, T2 and T3 ten plants per treatment were sampled, washed and stored at −80°C for gene expression analysis. At T1, T2 and T4 ten plants per treatment were sampled and utilized for bacterial counting and microbiome analysis. A time line of the experiment is presented in SF1. The remaining 20 plants were used for physiological and morphological evaluations. In particular, Nitrogen Balance Index (NBI), indicator of the plant nitrogen status at the beginning of flowering stage (Tremblay *et al*., 2012) was calculated as ratio between chlorophyll (CHL) and flavonoid (FLA) concentration recorded by the DUALEX 4 Scientific (Dx4) chlorophyll meter (Force-A, Paris, France) (Goulas *et al*., 2004). Measurements were carried out on both adaxial and abaxial faces of the panicle leaf for each plant. The total height was also measured for each plant as well as the dry weight of shoots (obtained after 48 h at 65°C).

### Colonization counts

The roots of 10 rice plants for each treatment at each time point were cleaned from soil and washed thoroughly under tap water before processing them. For T1, complete roots were macerated, while for T2 and T4, 600 mg of root samples were macerated and resuspended in 2 ml of PBS. Serial dilutions, up to 10^−5^, were made and 100 μL of each dilution was plated in triplicate on TSA (Tryptic soy agar) and TSA with rifampicin (50 µg/L). Plates were incubated at 28°C for 48 hrs and the emerged colonies were counted and presented as function of plated volume, dilution factor, macerated volume and starting material weight in order to determine the number of total cultivable CFUs (Colony Forming Units) and of rifampicin resistant CFUs present in 1 g of roots.

### Bacterial genomic DNA extraction and 16S rRNA gene amplicon library preparation and sequencing

Five hundred mg of root samples from 10 ST9-treated and untreated plants at T2 and T4 time points were used to extract DNA and perform microbiome analysis. DNA was extracted using the PowerSoil Kit (Qiagen, Hilden, Germany) following supplier instructions. For the 16S amplicon libraries preparation, 12.5 ng of DNA, quantified using Nanodrop, were used for each sample to prepare the 16S rRNA amplicon libraries. Library preparation was done using the Illumina methodology following the 16S Metagenomic Sequencing Library Preparation protocol (https://emea.support.illumina.com/content/dam/illumina-support/documents/documentation/chemistry_documentation/16s/16s-metagenomic-library-prep-guide-15044223-b.pdf). Sequencing was performed at CBM scrl (Trieste, Italy) with MiSeq methodology.

### Microbiome sequence analysis

FASTQ files were demultiplexed using the QIIME 1.9.1 split_libraries_fastq.py script and then analysed using DADA2 v1.4.0 (Callahan *et al*., 2016) adapting the methods from the DADA2 Pipeline Tutorial (1.4) and including dereplication, singletons and chimera removing (sample inference was performed using the inferred error model, while chimeric sequences were removed using the removeBimeraDenovo function). The Greengenes (GG) database (McDonald *et al*., 2012), giving a final Operational Taxonomic Unit (OTU) table, was employed to assign bacterial taxonomy by means of the assign Taxonomy function with a 97% sequence similarity. The resulting OTUs were clustered at Genus taxonomic level obtaining the final profile and abundance of bacterial taxa in the different samples. Statistical analysis was performed using the vegan package version 2.5-4 (Oksanen *et al*., 2013) and phyloseq package (McMurdie and Holmes, 2013) in R version 3.5.2 (R CoreTeam, 2014). Relative abundances of OTUs between samples were calculated. To test the differential representation of microbial taxa in diverse samples the Deseq2 package (Love *et al*., 2014) was used.

### RNA extraction and cDNA conversion

The RNA from three biological replicates for each thesis at each time point was extracted using the RNeasy Plant Mini Kit (Qiagen, Hilden, Germany), according to the manufacturer’s instructions. After the RNA extraction, any amount of DNA was removed using DNase (RQ1 RNase-Free DNase, Promega, Madison, Wisconsin, USA) and measured using Qubit (Invitrogen, ThermoFisher Scientific, Massachusetts, USA). The absence of genomic DNA was verified through PCR on RNA with the primers for the reference gene (*OsACT* Lee *et al*., 2010). Total RNA was used for each sample to synthesize the cDNA, according to the SuperScript II Reverse Transcriptase (Invitrogen, ThermoFisher Scientific, Massachusetts, USA) procedure using random primers.

### Primer selection

The genes analysed in this study (Table 1) were selected based on their regulation and role during the interaction between PGPR and rice. In particular, in addition to genes coding for a metallothionein – like protein and a multiple stress-responsive zinc-finger protein (Brusamarello-Santos *et al*., 2012), genes involved in ethylene and auxin pathways were considered (Song *et al*., 2009; Brusamarello-Santos *et al*., 2012; Vargas *et al*., 2012). Before RT-qPCR, all primers (Table ST1) were tested *in silico* on PRIMERBLAST and in PCR reactions on genomic DNA extracted from Baldo rice. The DNA extraction was performed using the DNeasy Plant Mini Kit (Qiagen, Hilden, Germania) according to the manufacturer’s instructions.

**Table 1:**
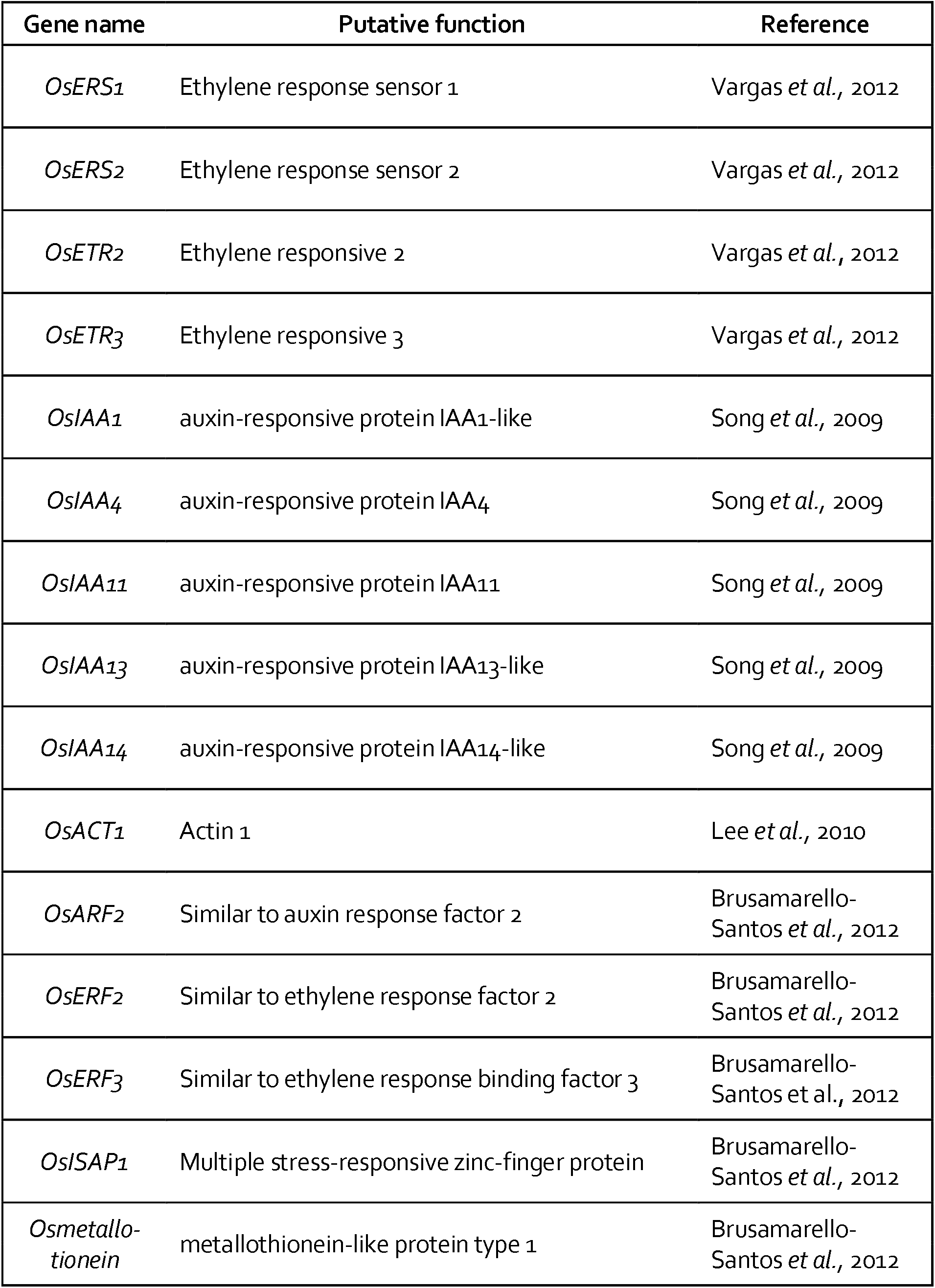
List of the genes considered for plant gene expression

### Gene expression analysis

Quantitative RT-PCR was carried out with 7500 Fast Real Time Systems (Applied Biosystem, ThermoFisher Scientific, Massachusetts, USA). Each PCR reaction was conducted on a total volume of 10 µl, containing 1 µl cDNA, 5 µl SYBR Green Reaction Mix and 0.3 µl of each primer (10 µM) using a 96-well plate. The used primers are listed in ST1. The following PCR programme, which includes the calculation of a melting curve, was used: 95°C for 10 min, 40 cycles of 95°C for 15 s, 60°C for 1 min, 95°C for 15 s, 60°C for 1 min, and 95°C for 15 s. All the reactions were performed for three biological and technical replicates. The baseline range and CT (cycle threshold) values were automatically calculated using the 7500 Fast Real Time Systems software. In order to compare data from different PCR runs or cDNA samples, the CT values of all the genes were normalized to the CT value of *OsACT1*, the reference gene. The candidate gene expression was normalized to that of the reference gene by subtracting the Ct value of the reference gene from the Ct value of the candidate gene efficiency correction, from the equation 2^−ΔΔCT;^ (Livak and Schmittgen, 2001), where ΔΔCT represents the ΔCT sample−ΔCT control.

### Data analysis

The statistical analysis of the phenotyping data was done using JMP7 (SAS Institute Inc., Cary, NC, 1989-2019). Concerning gene expression, statistical analyses were carried out using REST 2009, version 2.0.13 (Qiagen, Hilden, Germany) (Pfaffl *et al*., 2002), considering 0.05 as the *p*-value. Only significant expression values were considered and visualized as heat maps by a custom R (version 3.6.3) script (command “heatmap.2”). In order to reduce the data set dimension, a PCA (Principal Component Analysis) was carried out on ΔCT data for each biological replicate through R (version 3.6.3) (CRAN package “ggfortify”). The input files for R analysis were tabular file in .csv including ID# genes and ID# samples.

## Results

### *Identification and characterization of P. chlororaphis* ST9

It was of interest to isolate and characterize new *P. chlororaphis* strains from the soil in order to test their potential use as plant probiotics. Strain ST9 was isolated form bulk soil as described in the Materials and Methods section. The genome sequence of *P. chlororaphis* ST9 was determined and established that its chromosome was 6.7 Mb long with a GC content of 63%; according to the RASTtk annotation scheme it contains 6185 predicted protein-coding sequences (CDSs) and 83 RNAs. Phylogenetic analyses based on MSLA using 6 loci (16S rRNA, *recA, gyrB, rpoD, carA* and *atpD;* Hilario *et al*., 2004-) and 16 sequences of *P. chlororaphis* isolates revealed that strain ST9 belongs to the *Pseudomonas chlororaphis* subsp. *aurantiaca* group (Fig. SF2). As other *P. chlororaphis* strains, its genome possesses several loci encoding for anti-bacterial/anti-fungal compounds including operons for the biosynthesis of phenazine, of two R-tailocins bacteriocines and of pyrrolnitrin. Other loci potentially involved in microbial pathogen antagonism, cell-cell signal interferences, niche colonization, plant protection and plant-beneficial traits were found and reported in Table 3. In summary, *P. chlororaphis* ST9 contains many of the features that can potentially make this strain a plant probiotic and biocontrol agent.

In order to begin assessing possible PGP potential of *Pseudomonas chlororaphis* ST9, several *in vitro* phenotypes were determined. A summary of the tested phenotypes is summarized in Table 2. Among the performed tests, the ability of ST9 to inhibit the growth of a wide variety of pathogens, including rice pathogens such as *Dickeya zeae, Pseudomonas fuscovaginae* and *Magnaporthe oryzae* was observed. Other PGP related *in vitro* phenotypes such as lipolytic and proteolytic activities and motility were observed. In summary, this isolate possesses several phenotypic abilities many of which can be of importance in plant colonization and PGP.

**Table 2.**
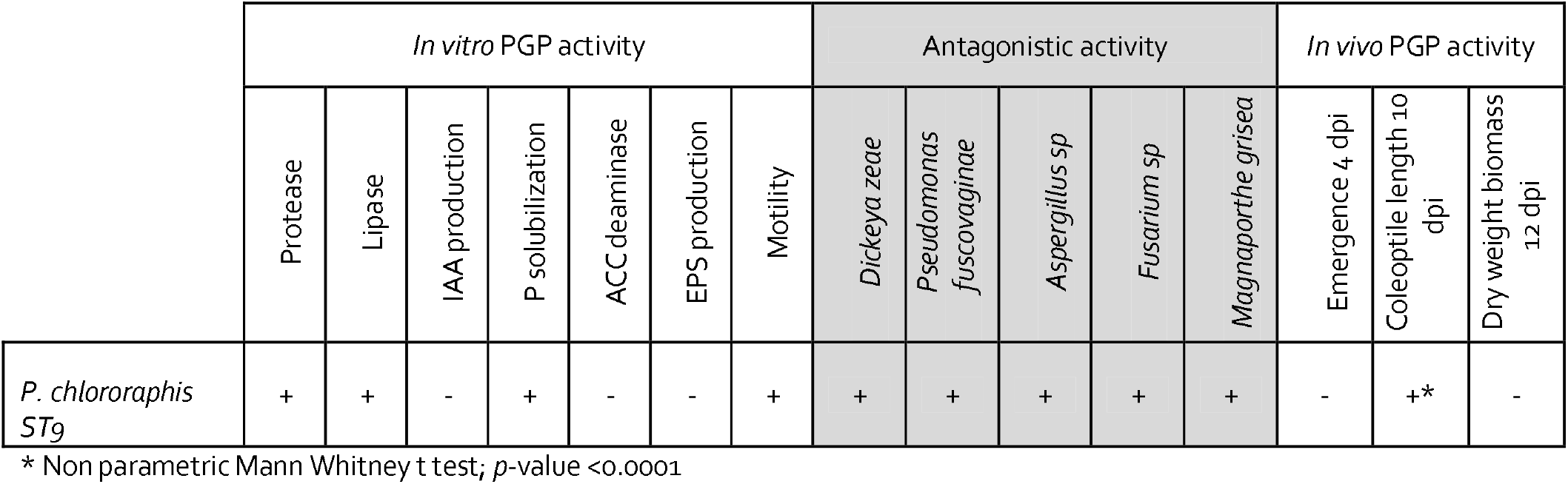
Phenotypic characterization of PGP features of *P. chlororaphis* ST9

**Table 3.**
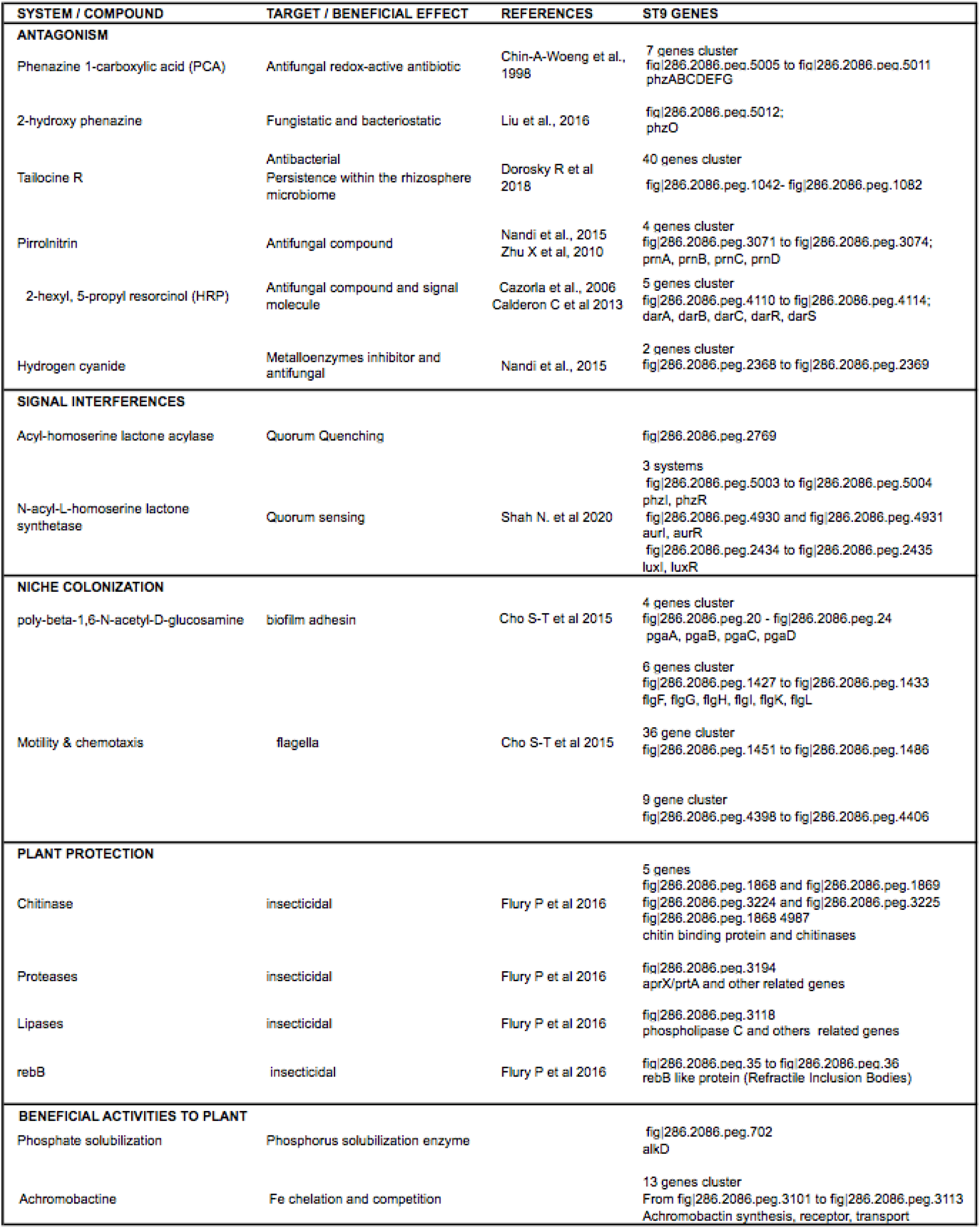
PGP and protection features in the genome of *P. chlororaphis*

### *P. chlororaphis* ST9 root colonization ability, persistence and effect on the microbiome in the rhizosphere

It was of interest to evaluate the ability of *P. chlororaphis* ST9 to colonize and persist in the rhizosphere of rice plants. Rice seeds were inoculated with *P. chlororaphis* ST9, grown in greenhouse and roots were then collected at T1 (10 dpi), T2 (28 dpi) and T4 (90 dpi) together with un-inoculated controls for ST9 strain CFU counts. Results are summarized in Figure 1 which show that the level of colonization after 10 days was significantly high (10^7^ cfu/g of root, -wet weight) then it decreased and stabilized around a value of 10^4^ cfu/g at 90 dpi. Colonies grown on TSA rifampicin plates were randomly chosen for identification through 16S rRNA amplification and sequencing to confirm their identity as *P. chlororaphis* ST9. Rifampicin resistant colonies present in the untreated plants resulted to be a mixture of different bacteria: some of them were part of an analogue experiment carried out in concomitance (data not shown). These results evidenced that under the conditions tested, *P. chlororaphis* ST9 was an efficient root colonizer and able to persist in the rhizosphere.

**Figure 1.**
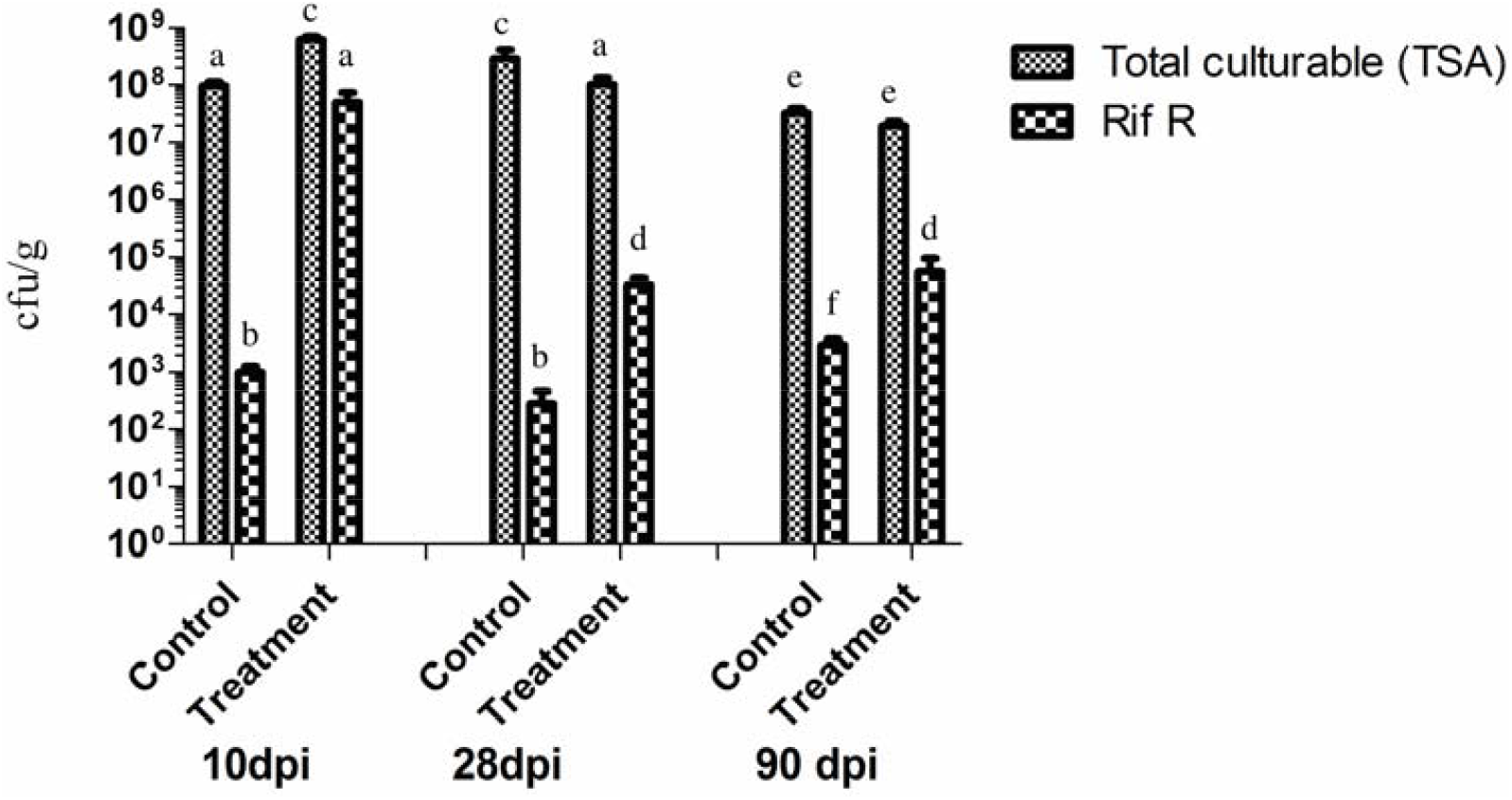
Rice root colonization by *P. chlororaphis* ST9 was evaluated at three time points: 10, 28 and 90 dpi (days post inoculation). The average of the cfu/g of fresh weight root of 10 plants for each treatment and time point is presented. Each sample was plated on TSA to count all cultivable bacteria, and on TSArif to count just the rifampicin resistant CFUs, mainly *P. chlororaphis* ST9. Statistical analysis: same letter means no significant difference (t-test, non-parametric, Mann Whitney, *p*-value < 0.05).

It was of interest to determine the possible effects of strain ST9 colonization on the rhizosphere microbiome. A 16S rDNA community profiling approach was performed, characterizing and comparing the microbial root community at 28 and 90 dpi in ST9 treated and untreated plants. The richness and diversity values of the bacterial communities after normalization are shown in Figure 2.

**Figure 2.**
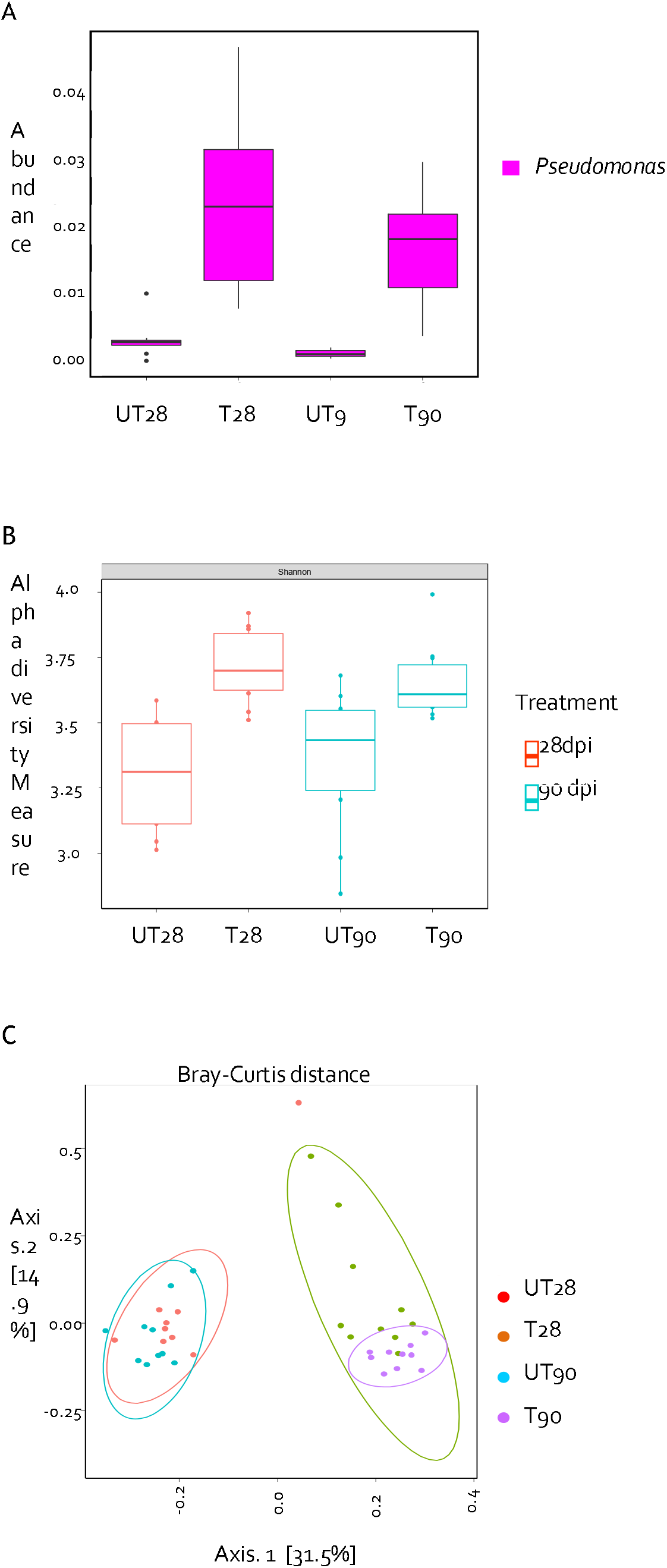
Microbiome analysis. (A) *Pseudomonas* genus abundance according to the plant treatment and time point; the Welch two Sample t-test was used to compare untreated *versus* treated samples at 28 dpi (*p*-value = 0.0006793) and at 90 dpi (*p*-value = 0.0003417). (B) Alpha Diversity (Shannon index) at community level in accordance to the treatment and the time point; the Wilcoxon rank sum test was used, *p*-value for untreated *versus* treated samples both at 28 and 90 dpi results < 0.01. (C) Principal component analysis of the samples in accordance to the treatment and the time point, similarity between the different communities was evaluated using the Bray-Curtis test; the bacterial communities of the untreated plants cluster together and differently from the bacterial communities colonizing the ST9 treated samples (*p*-value < 0.001). UT28: untreated samples at 28 dpi; T28: ST9 treated samples at 28 dpi; UT90: untreated samples at 90 dpi; T90: ST9 treated samples at 90 dpi.

The *Pseudomonas* genus, was significantly more abundant (*p*-value<0.001) in the *P. chlororaphis* ST9 inoculated plants as presented in Figure 2A. The presence of *P. chlororaphis* ST9 was confirmed by finding its 16S V3-V4 region sequence among the reads obtained by the NGS experiment. Significant differences (*p*-value = 8.66e-05) in Shannon alpha diversity (Figure 2B) were found between inoculated and un-inoculated samples both at 28 and 90 dpi. Beta diversity analysis based on Bray Curtis distance (Figure 2C) was performed to compare the microbial compositions of the two different tested conditions. The results obtained highlighted differences between the microbial populations in relation to the treatment. As result of the *P. chlororaphis* treatment, several genera resulted differently distributed as evidenced by the heatmap of Figure 3 and in SF3 and SF4. Among the genera more abundant in untreated samples were *Duganella, Clostridium, Aeromonas, Enterobacter and Vogesella*; the majority of them are also defined as animal/human pathogens and just few species, such as *Clostridium puniceum* (Lund *et al*., 1981) and *Enterobacter cloacae* (García-González *et al*., 2018), are correlated with plant diseases. On the other hand, genera such as *Janthinobacterium*, known for its antifungal features (Haack *et al*., 2016), *Flavobacterium*, often reported as a plant growth promoting rhizobacterium (Kolton *et al*., 2016), *Bacillus, Paenibacillus, Bradyrhizobium* and others known for their PGP potential (Sansinenea, 2019; Radhapriya *et al*., 2018; Greetatorn, *et al*., 2019; Ferreira *et al*., 2019; Berg, 2009; Souza *et al*., 2015) were significantly more abundant in the *P. chlororaphis* ST9 treated communities than in untreated ones.

**Figure 3.**
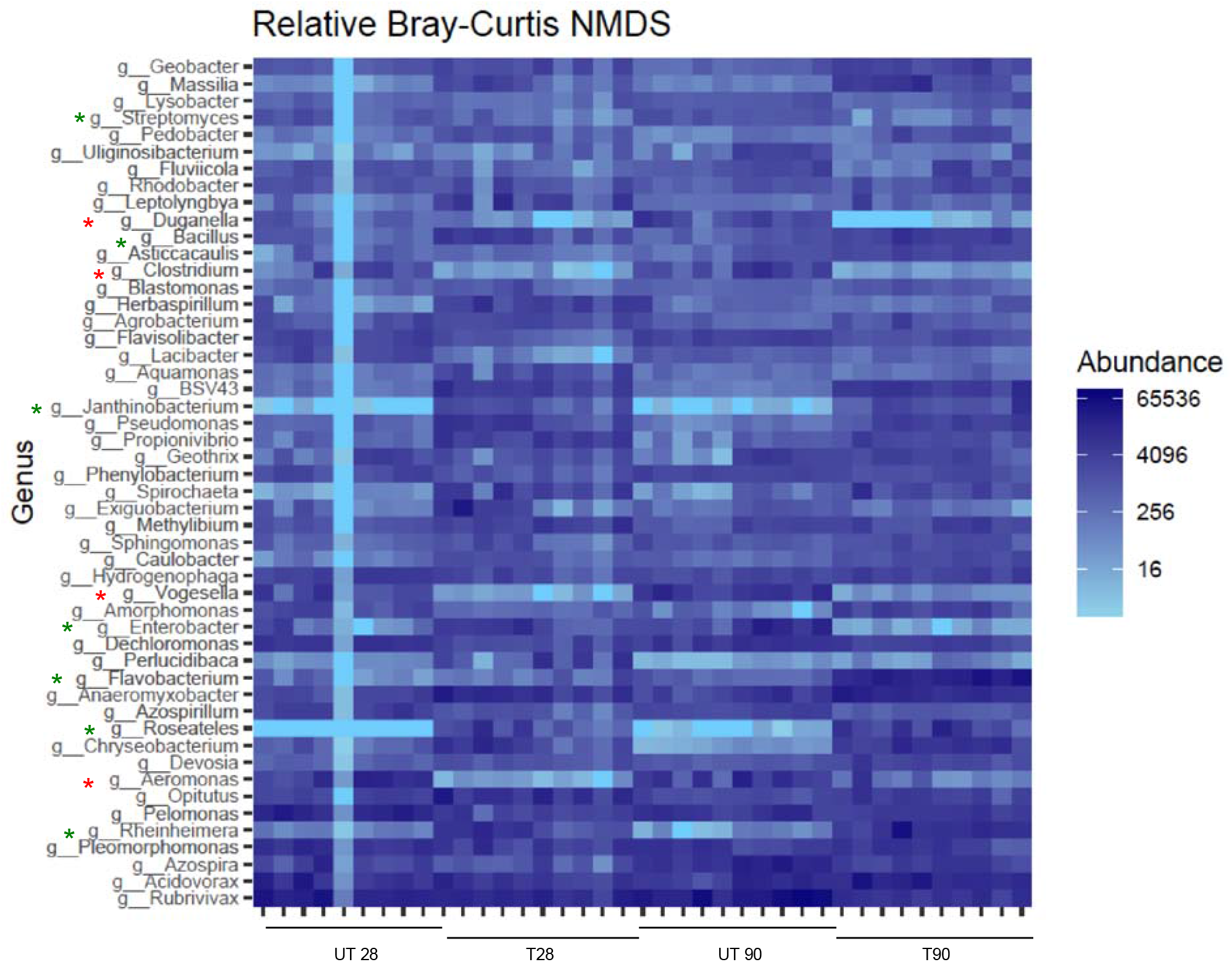
Heat map showing the relative abundance (% of sequencing reads) of the 50 predominant genera. Rows are bacterial genera. Columns are samples. Colors indicate taxa with a higher (blue) or lower (light blue) relative abundance in each sample. Genera showing a different distribution and abundance level between the samples are highlighted with *: green if enriched in the treated samples, red if enriched in the untreated ones. UT28: untreated samples at 28 dpi; T28: ST9 treated samples at 28 dpi; UT90: untreated samples at 90 dpi; T90: ST9 treated samples at 90 dpi.

### Plant gene expression and phenotypic analysis of *P. chlororaphis* inoculated plants

In order to determine the effects of *P. chlororaphis* ST9 on plant growth, several phenotypic parameters were assayed. Statistical analyses were carried out on chlorophyll and flavonoid content and on the nitrogen balance index (NBI) as physiological parameters. In addition, plant height and dry shoot biomass at 90 dpi (after flowering stage) was also established. Results did not show any statistical differences between control and ST9 inoculated plants [p(F)<0.05] (Figure 4), although a tendency on higher NBI and lower flavonoid contents was measured in inoculated plants compared with the control. It was concluded that under the conditions tested, no significant plant beneficial effect was observed upon seed inoculation of *P. chlororaphis* ST9.

**Figure 4:**
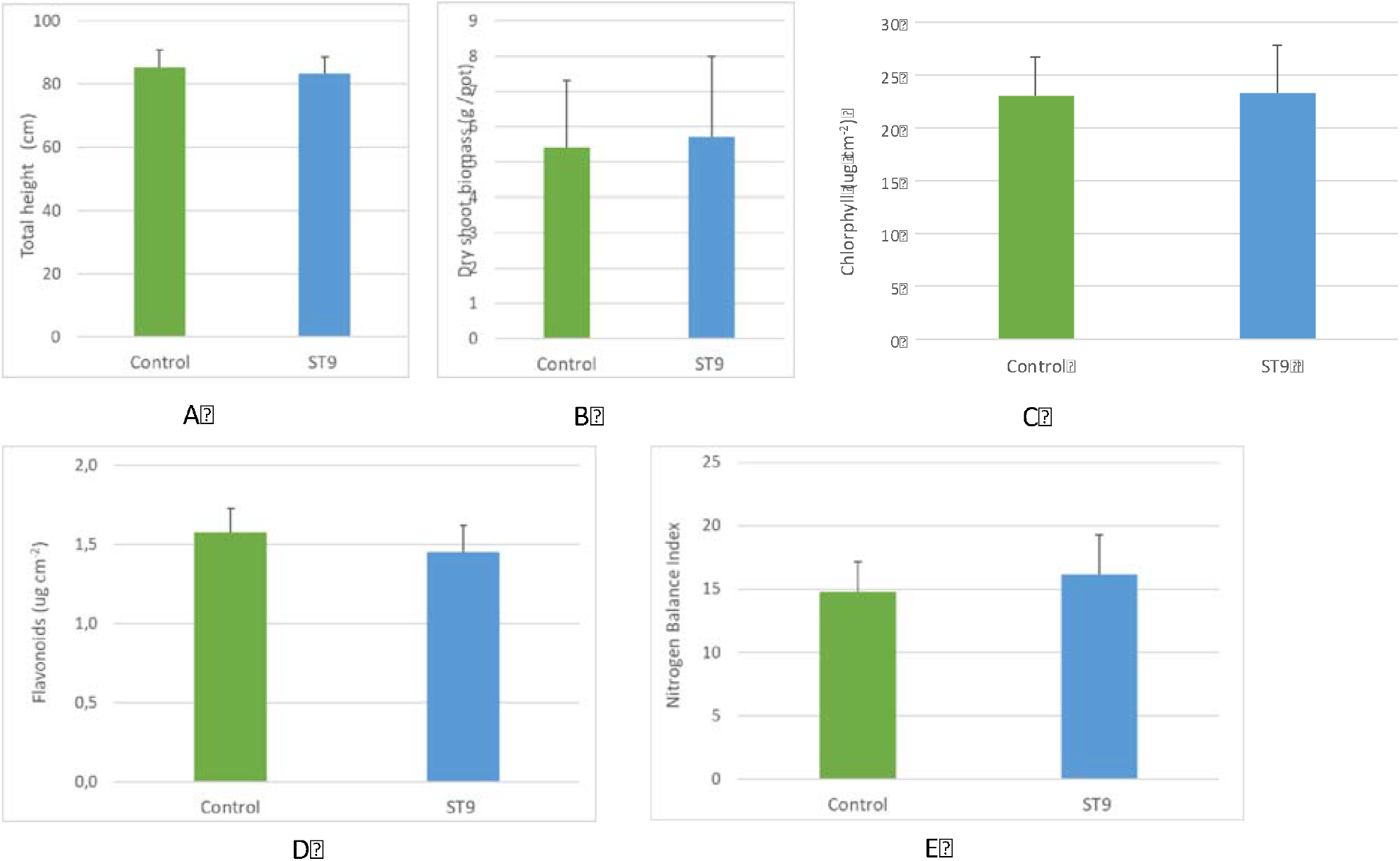
Total height (A), dry shoot biomass (B), chlorophyll (C) and flavonoid content (D), and NBI (E) in control un-inoculated and in ST9 inoculated plants.

In order to investigate the effect on PGPR beneficial plant-bacteria interactions, RT-qPCR was performed on 14 genes selected for their role in ethylene and auxin pathways (Table 1). In Supplementary Table 2 (ST2) the fold change was shown for genes that were significantly and not significantly differentially expressed, while in Figure 5 the heat map representation of the transcript levels coupled to a hierarchical clustering in ST9 inoculated plants is presented. At T2, significant differences in the expression of some loci was evidenced when compared to the other two time points. The most up- and down-regulated genes at T2 were *OsETR3* and *Osmetallotionein*, respectively. At T1, with the exception of *Osmetallotionein*, there were no significantly differentially regulated genes. At T3 the most up-regulated gene was *OsIAA14*, while *OsERF3* was the most down-regulated. The only genes that were never significantly differentially expressed were *OsIAA11* and *OsIAA4*. The PCA on ΔCT data of control (un-inoculated) and ST9 inoculated samples for each replicate and time point showed, without outlier observations, a clear distinction between control and inoculated samples at T2, where we have found the highest number of differently regulated genes (Figure 6). On the other hand, there was no separation between data at T1 and at T3, where we have found a reduced number of regulated genes. In summary, most loci displayed some level of variation in gene expression upon inoculation with strain ST9; one gene was regulated at T1, while 12 and five genes were regulated at T2 and T3, respectively.

**Figure 5:**
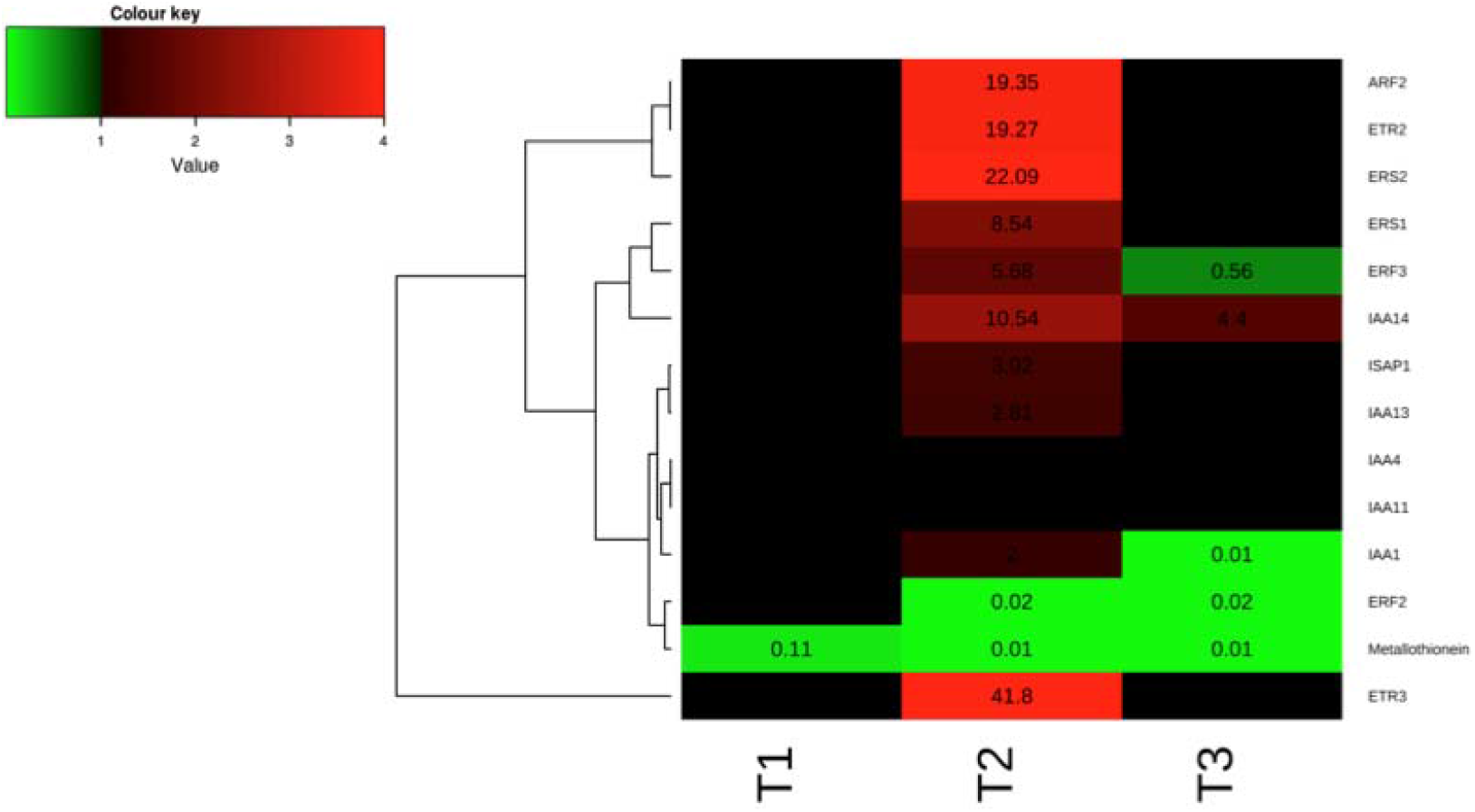
Heat map representation of the transcript levels coupled to a hierarchical clustering in ST9 inoculated plants. Each column represents a time point, while each row represents a gene. Expression levels are coloured green for low intensities and red for high intensities (see scale at the top left corner). The black cells represent genes not significantly different from those of the untreated samples.

**Figure 6:**
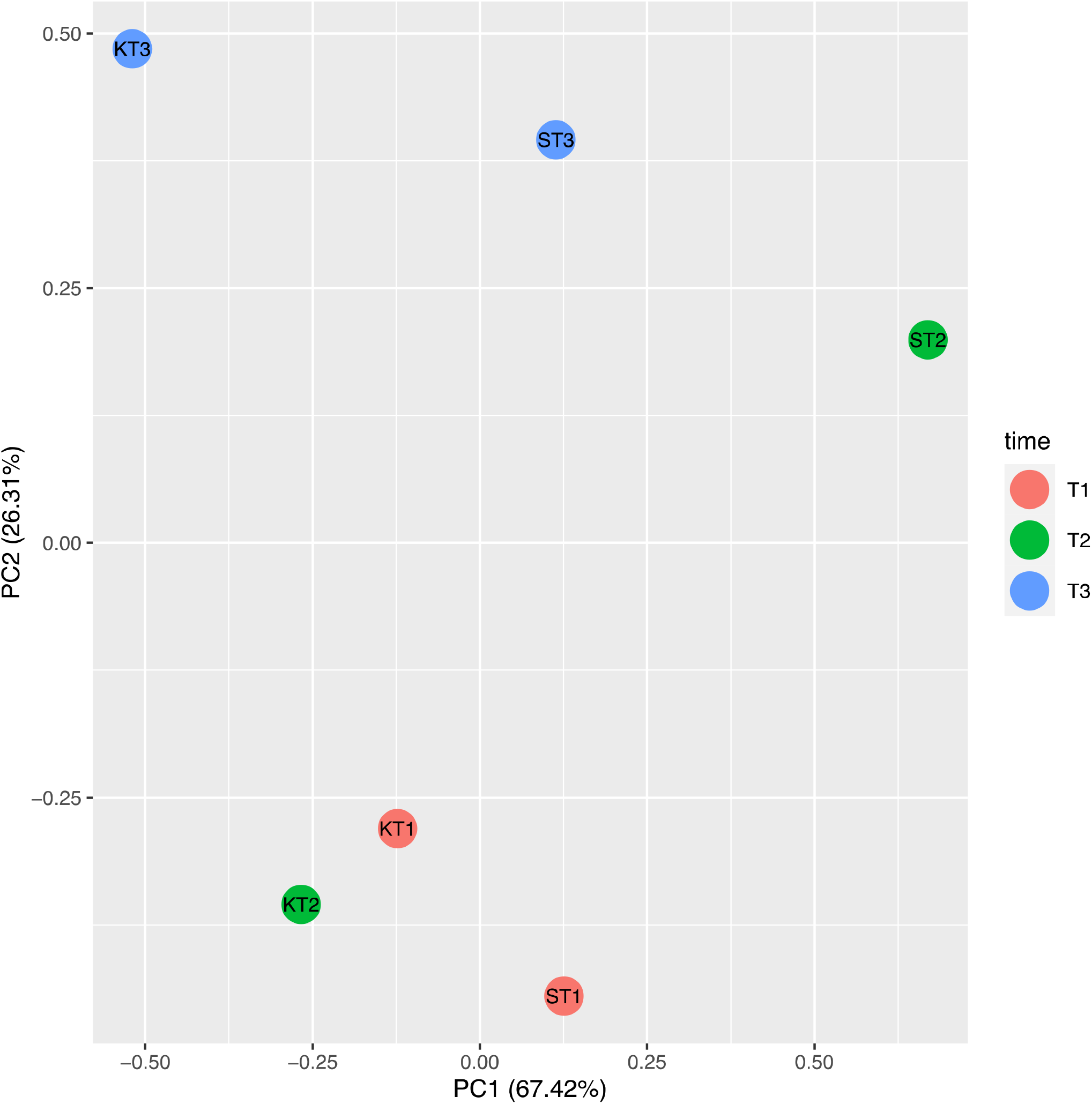
PCA carried out on ΔCT of each biological replicate of untreated control (K) and ST9 inoculated plants (S) at each time point (T1=10 dpi; T2=28 dpi; T3=40dpi). The cumulative variability percentage is given by the sum of 67.42% for PC1 and 26.31% for PC2, resulting in 93.73%. KT2 and ST2 are clearly separated in the space, differently by the other time points.

## Discussion

This study presents the identification of a *P. chlororaphis* strain and characterisation studies in relation to plant growth promotion (PGP). The work performed included root colonisation and persistence, effect on the root bacterial microbiome, effect on targeted plant gene expression and a greenhouse plant growth promotion test. Results showed that *P. chlororaphis* ST9 is a good rice root colonizer integrating well in the microbiome affecting plant gene expression however in our greenhouse experiment it did not result in a plant growth promotion effect.

*P. chlororaphis* was able to persist in the root compartment for 90 days without decreasing its abundance below a concentration of 10^4^ cfu/g of root. An initial good root colonization rate then slowly decreasing in the later stages of plant growth has also been observed for other PGPR strains. For example in Chaudhary *et al*. (2013) *Azotobacter* strain ST24 colonizes well the wheat rhizosphere decreasing during growth and reaching a density of 10^4^cfu/ml at 90 days after sowing. Similarly Solanky and Garg (2014) performed the inoculation of *Azotobacter* on *Brassica campestris* and again detected its persistence at 30 and 60 day post inoculation being approximately 2×10^4^ cfu per plant at 60 dpi. Mosquito *et al*. (2020) inoculated a *Kosakonia* sp. strain in rice in the field and it was persistent at 30 dpi, while it was undetectable at 60 and 90 dpi. The effect of ST9 inoculation on the total bacterial population evidenced an increase in biodiversity with an enrichment of certain bacterial genera (e. g. *Janthinobacterium, Flavobacterium, Bacillus, Paenibacillus, Bradyrhizobium*) which are known to establish beneficial association with plants (Berg, 2009; Souza *et al*., 2015; Radhapriya *et al*., 2018; Ferreira *et al*., 2019; Greetatorn, *et al*., 2019; Sansinenea, 2019).

No plant beneficial effect on the rice inoculated with strain ST9 was observed. The reason is not known regardless that the strain was present and persistent in the rhizosphere. It would be appropriate to challenge rice with biotic and abiotic stresses and determine whether the presence of strain ST9 would have resulted in a tolerance effect. On the bases of physiological parameters however ST9 inoculated plants displayed less stress symptoms than controls. A similar trend was also observed Andreozzi *et al*. (2019) where NBI was significantly lower in the uninoculated control with respect to the *Herbaspirillum huttiense* RCA24 + *Enterobacter cloacae* RCA25 inoculated Baldo rice plants.

Gene expression studies were performed with 14 rice loci related to ethylene and auxin pathways together with genes coding for a metallothionein–like protein and a multiple stress-responsive zinc-finger protein; a role for these genes during rice-PGPR interaction has been demonstrated (Song *et al*., 2009; Brusamarello-Santos *et al*., 2012; Vargas *et al*., 2012; Ambreetha *et al*., 2018). Genes *OsERS1, OsERS2, OsETR2, OsETR3*, homologs to *Arabidopsis thaliana* ethylene receptors, were transcribed at higher level in the *P. chlororaphis* ST9 inoculated plants. Similarly, Vargas *et al* (2012) also observed increase in expression of these loci when inoculated with *Azospirillum brasilense* and *Burkholderia kururiensis*. In their study rice varieties having more BNF (Biological Nitrogen Fixation) capacities showed higher bacterial colonization as well as up-regulation of ethylene receptors genes. In this study, these loci were significantly up-regulated at 28 dpi indicating that the Baldo rice variety displays good nitrogen fixing capacity as also reported by Andreozzi *et al*. (2019). The *OsERF2* and *OsERF3* genes, encoding for transcriptional factors related to ethylene, were also differently expressed in ST9 inoculated plants indicating that this strain could be involved in ethylene hormone levels. Indole-3-acetic acid (IAA) pathway modulation is important during an effective plant colonization, both by pathogenic and by nonpathogenic organisms (Brusamarello-Santos *et al*., 2012). In *P. chlororaphis* inoculated plants it was established that loci *OsIAA1, OsIAA13* and *OslAA14* were up-regulated at T2, *OsIAA1* was down-regulated at T3, while *OsIAA11* and *OsIAA4* were not differently transcribed in the three considered time points. Some of these trends are in accordance and others in contrast with previous observations (Brusamarello-Santos *et al*., 2012; Ambreetha *et al*., 2018); this could be due to the different timing and experimental applied conditions. Expression of genes associated to defense can be also affected by bacterial presence in the rhizosphere (Brusamarello-Santos *et al*., 2012). *OsISAP1*, encoding for a multiple stress-responsive zinc-finger protein, was up-regulated at T2, suggesting an activation of the plant defense mechanisms. The gene coding for a metallothionein, which is a metal binding protein involved in metal homeostasis, was always down-regulated at each time point in agreement with previous data (Brusamarello-Santos *et al*., 2012). The plant defense system could be primed by *P. chlororaphis* ST9 maintaining a low level of stress; future studies on the response to biotic stress of strain ST9 inoculated plant will determine whether the change of expression of these loci provides immunity to microbial pathogens.

Inoculated PGPR must interact or compete with other microorganisms in the rhizosphere microbiome and this can cause a short persistence of the inoculated bacteria possibly affecting its probiotic effects (Backer *et al*., 2018). The ability of *P. chlororaphis* ST9 to successfully colonize and persist in the rhizosphere is an important trait for its possible use as bioinoculant for agriculture. Furthermore, the plethora of antimicrobial activities encoded in its genome makes *P. chlororaphis* ST9 a good candidate for biotic stress tolerance tests upon its inoculation. Plant gene expression analysis demonstrated some positive effects of the presence of strain ST9 on the expression of plant hormonal pathways such as IAA and ethylene; further investigation should be carried out on rice defence genes in order to better clarify the type of interaction. *P. chlororaphis* strains are considered safe for the environment and human health (EPA, 2009) and their use in agriculture is has been permitted through the application of live microorganism formulations (Rosas 2017; Anderson *et al*., 2018) and via the production and purification of metabolites (Liu *et al*., 2016; Peng *et al*., 2018). Future experiments with *P. chlororaphis* ST9 will reveal its full potential as a bioinoculant for PGP, tolerance to abiotic and most importantly, biotic stresses.

