## Supplementary Materials for "Isolation and characterization of *Pseudomonas chlororaphis* strain ST9 and its potential as a bioinoculant for agriculture"

Supplementary Figures (SF)

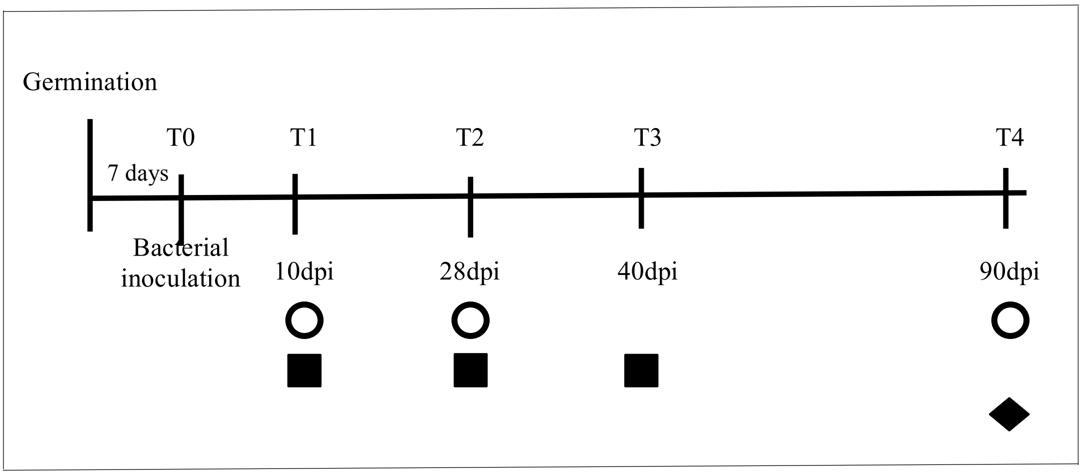


SF1. Time line of the sampling and use of the samples: O bacterial counting and microbiome analysis, ■ plant gene expression analysis, ◆ plant physiological and morphological evaluations.

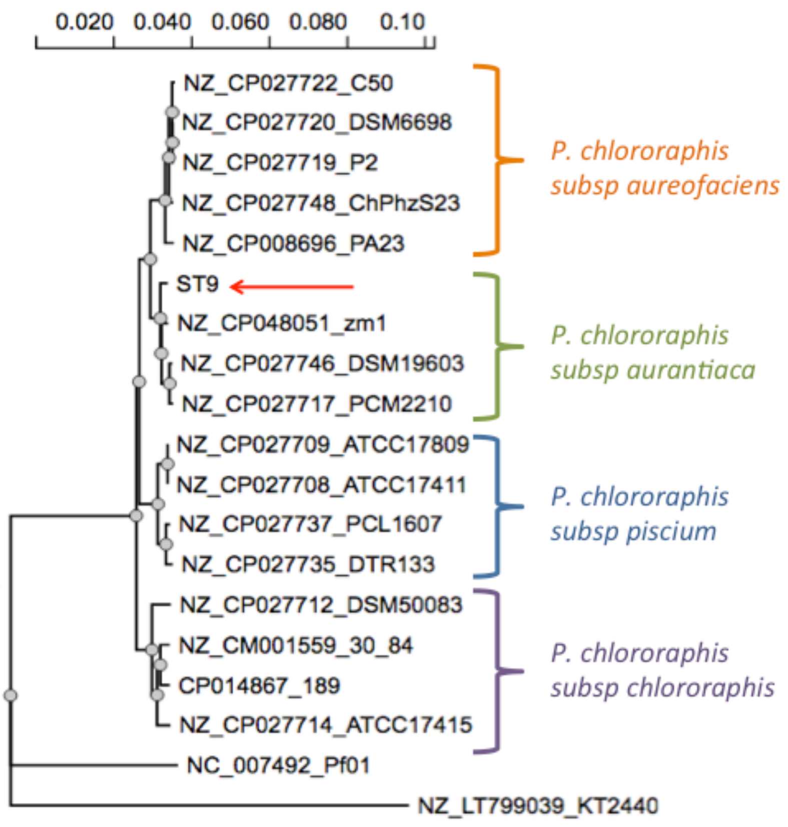


SF2. Phylogenetic tree reconstructed by the MSLA method based on six concatenated gene sequences (16S rRNA, *recA, gyrB*, *rpoD*, *carA*, *atpD* – 9371 nt-) of 17 *Pseudomonas chlororaphis* strains. The strains *P. putida* KT2440 and *P. fluorescens* Pf01 served as outgroups. The 4 subspecies of *P. chlororaphis* cluster separately and ST9 is part of the *aurantiaca* subspecie. The phylogenetic analysis was performed using the NGPhylogeny.fr public platform.

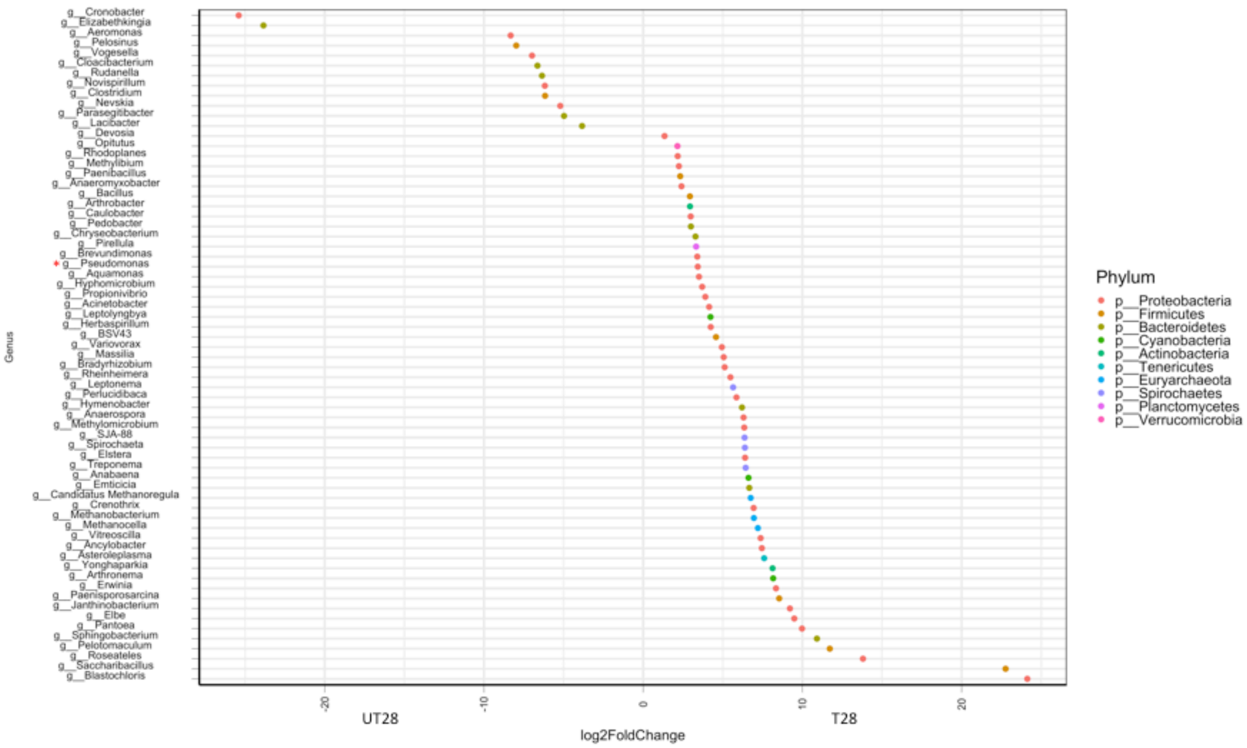


SF3. Differential representation of OTUs between ST9 inoculated and control samples at 28 days post inoculation.

Differential abundance of OTUs between the two groups of tested samples was assessed by fitting a local regression model with a negative binomial distribution to the data and testing for differential abundance with a likelihood ratio test as implemented in the R package DESeq2 (Love *et al.*, 2014) in conjunction with the phyloseq package. Taxa are represented as dots in the graph of fold change. A negative log2Foldchange indicates taxa more abundant in untreated samples, while a positive log2Foldchange indicates taxa more abundant in treated samples. Samples with a *p*-value less than 0.0001 and mean representation over all samples higher than 1 are shown. UT28: untreated samples at 28 dpi; T28: ST9 treated samples at 28 dpi.

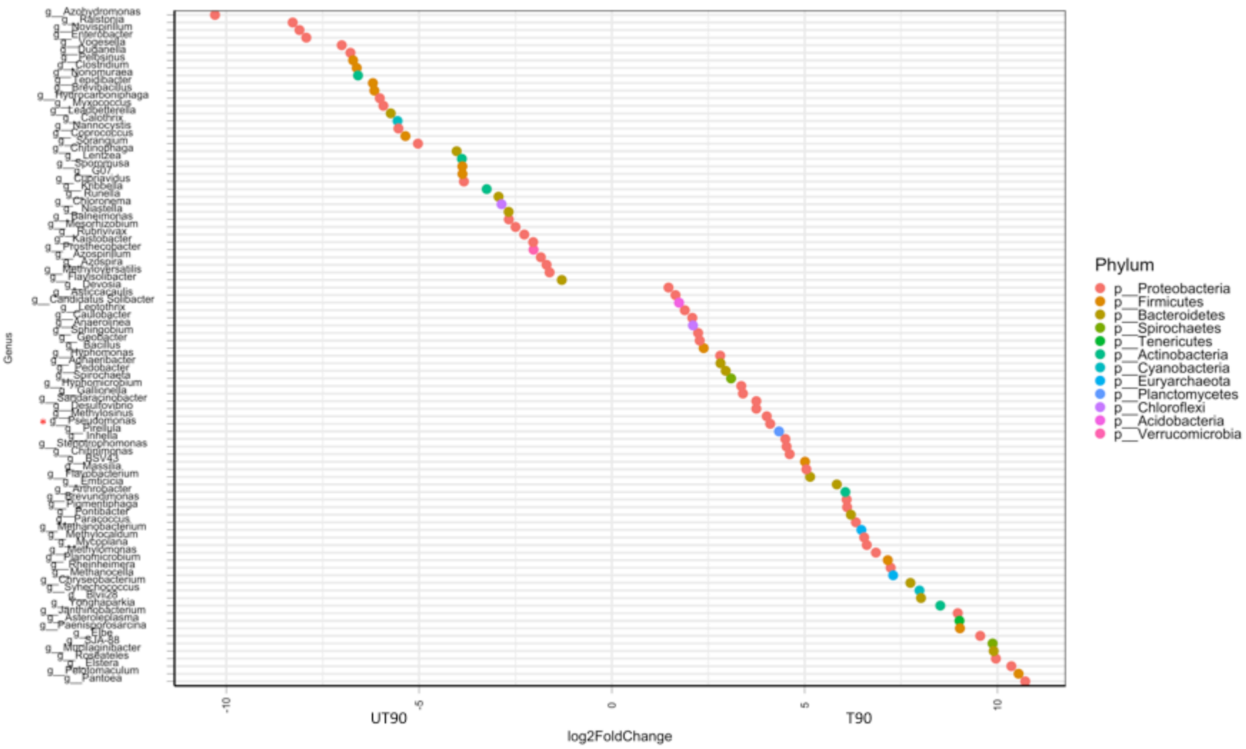


SF4. Differential representation of OTUs between ST9 inoculated and control samples at 90 days post inoculation. Differential abundance of OTUs between the two groups of tested samples was assessed by fitting a local regression model with a negative binomial distribution to the data and testing for differential abundance with a likelihood ratio test as implemented in the R package DESeq2 (Love *et al.,* 2014) in conjunction with the phyloseq package. Taxa are represented as dots in the graph of fold change. A negative log2Foldchange indicates taxa more abundant in untreated samples, while a positive log2Foldchange indicates taxa more abundant in treated samples. Samples with a *p*-value less than 0.0001 and mean representation over all samples higher than 1 are shown. UT90: untreated samples at 90 dpi; T90: ST9 treated samples at 90 dpi.

Supplementary Tables (ST)

Table ST1: List of the primers used in this study

| Primer name | Primer sequence (5'-3') | Gene name | Putative function | Reference |
|---|---|---|---|---|
| fD1<br>rP2 | AGAGTTTGATCCTGGCTCAG<br>ACGGCTACCTTGTTACGACTT | *16SrRNA* | 16S rRNA amplification | Weisburg *et al.* 1991 |
| 518F<br>800R | CCAGCAGCCGCGGTAATACG<br>TACCAGGGTATCTAATCC | *16SrRNA* | 16S rRNA sequencing | Lane 1991 |
| OsERS1f<br>OsERS1r | GAAAGGTCAGGCTTCTCTGAAATC<br>ATGCCGTCGATCAATTTACAGTAG | *OsERS1* | Ethylene response sensor 1 | Vargas *et al.*, 2012 |
| OsERS2f<br>OsERS2r | CCTCGGGTTCGCTACCAAT<br>GCATGGCGATGGCATCAT | *OsERS2* | Ethylene response sensor 2 | Vargas *et al.*, 2012 |
| OsETR2f<br>OsETR2r | CTTTAGCAGCACTGGGAGATGA<br>TGAGAACCATGAGGCTCTTTCA | *OsETR2* | Ethylene responsive 2 | Vargas *et al.*, 2012 |
| OsETR3f<br>OsETR3r | CGAGCTGGCGCGAATT<br>TTAGACAAACAGACCTCCAGCAAA | *OsETR3* | Ethylene responsive 3 | Vargas *et al.*, 2012 |
| OsIAA1f<br>OsIAA1r | ACCAAGAGCCGCTCAATGAG<br>ATCACACGTGGGCGAACATC | *OsIAA1* | auxin-responsive protein IAA1-like | Song *et al.*, 2009 |
| OsIAA4f<br>OsIAA4r | GCTCTTGCTGGATGGGTATGA<br>AGGTGATGGGCGTCTTGAAC | *OsIAA4* | auxin-responsive protein IAA4 | Song *et al.*, 2009 |
| OsIAA11f<br>OsIAA11r | AGTTGTCCATGGCGTTCCA<br>TGCTCTCCTTCAGCTGCTGAT | *OsIAA11* | auxin-responsive protein IAA11 | Song *et al.*, 2009 |
| OsIAA13f<br>OsIAA13r | CAAGGATGGTGACTGGATGCT<br>GATCCTCAAGCGTTTGCATGA | *OsIAA13* | auxin-responsive protein IAA13-like | Song *et al.*, 2009 |
| OsIAA14f<br>OsIAA14r | CCGTCGCCTATGAGGACAA<br>TTATCCGCAGCTTCTTGCAA | *OsIAA14* | auxin-responsive protein IAA14-like | Song *et al.*, 2009 |
| OsACT1f | GTATCCATGAGACTACATACAACT | *OsACT1* | Actin 1 | Lee *et al.*, 2010 |

| | | | | |
|---|---|---|---|---|
| OsACT1r | TACTCAGCCTTGGCAATCCACA | | | |
| ARF2-like f<br>ARF2-like r | GGCCTGAATCAAGTTGGAGATC<br>CTATCTGGCCGCGGAATAGTT | *OsARF2* | Similar to auxin response factor 2 | Brusamarello-Santos *et al.*, 2012 |
| ERF2-like f<br>ERF2-like r | CCGGCAAGGGTAGAGATGGT<br>TCAACATCGAAATCCCAAGAACT | *OsERF2* | Similar to ethylene response factor 2 | Brusamarello-Santos *et al.*, 2012 |
| ERF3-like f<br>ERF3-like r | TGCAGCAGCCTATGCAGATC<br>CGCGAGGACACTGCTTGAT | *OsERF3* | Similar to ethylene response binding factor 3 | Brusamarello-Santos *et al.*, 2012 |
| OsISAP1 f<br>OsISAP1 r | GCAATCCTCATCACACAGCAA<br>CCCTCTTGGTCTCAGGCTCTCT | *OsISAP1* | Multiple stress-responsive zinc-finger protein | Brusamarello-Santos *et al.*, 2012 |
| Metallo-thionein f<br>Metallo-thionein r | CAAACTGCTCCTGCGGAAAG<br>ACGACGGTGGCCTTGGT | *Osmetallo-tionein* | metallothionein-like protein type 1 | Brusamarello-Santos *et al.*, 2012 |

Table ST2: Fold change at each time point

| **Gene** | **Time** | | |
|---|---|---|---|
| | T1 | T2 | T3 |
| *OsERF2* | 0.22 | 0.02* | 0.02* |
| *OsMetallothionein* | 0.11* | 0.01* | 0.01* |
| *OsIAA1* | 0.22 | 0.002* | 0.01* |
| *OsIAA11* | 0.21 | 0.05 | 0.43 |
| *OsIAA13* | 2.54 | 2.81* | 0.3 |
| *OsIAA14* | 1.8 | 10.54* | 4.4* |
| *OsISAP1* | 1.16 | 3.02* | 0.62 |
| *OsETR3* | 1.72 | 41.8* | 1.88 |
| *OsARF2* | *0,69* | *19,35** | *1,16* |
| ERS1 | 0,3 | 8,54* | 2,04 |
| *OsERS2* | 0,87 | 22,09* | 2,46 |
| *OsERF3* | 0,81 | 5,68* | 0,56* |
| *OsIAA4* | 0,37 | 3,22 | 6,3 |
| *OsETR2* | 0,73 | 19,27* | 2,19 |

*: statistically significant different (P< 0.05)

Supplementary Materials (SM)

## SM1

Sequences used for the taxonomic analysis of *P. chlororaphys* ST9.

```
NCBI Reference Sequence: NZ_CP014867.1
Strain: 189
Chained genes: 16S rRNA, recA, gyrB, rpoD, carA, atpD
>NZ_CP014867.1_189
GAACTGAAGAGTTTGATCATGGCTCAGATTGAACGCTGGCGGCAGGCCTAACACATGCAAGTCGAGCGGTAGAGAGAAGC
TTGCTTCTCTTGAGAGCGGCGGACGGGTGAGTAATGCCTAGGAATCTGCCTGGTAGTGGGGGATAACGTCCGGAAACGGA
CGCTAATACCGCATACGTCCTACGGGAGAAAGCAGGGGACCTTCGGGCCTTGCGCTATCAGATGAGCCTAGGTCGGATTA
GCTAGTTGGTGAGGTAATGGCTCACCAAGGCGACGATCCGTAACTGGTCTGAGAGGATGATCAGTCACACTGGAACTGAG
ACACGGTCCAGACTCCTACGGGAGGCAGCAGTGGGGAATATTGGACAATGGGCGAAAGCCTGATCCAGCCATGCCGCGTG
TGTGAAGAAGGTCTTCGGATTGTAAAGCACTTTAAGTTGGGAGGAAGGGTACTTGCCTAATACGTGAGTATTTTGACGTT
ACCGACAGAATAAGCACCGGCTAACTCTGTGCCAGCAGCCGCGGTAATACAGAGGGTGCAAGCGTTAATCGGAATTACTG
GGCGTAAAGCGCGCGTAGGTGGTTCGTTAAGTTGGATGTGAAATCCCCGGGCTCAACCTGGGAACTGCATCCAAAACTGG
CGAGCTAGAGTATGGTAGAGGGTGGTGGAATTTCCTGTGTAGCGGTGAAATGCGTAGATATAGGAAGGAACACCAGTGGC
GAAGGCGACCACCTGGACTGATACTGACACTGAGGTGCGAAAGCGTGGGGAGCAAACAGGATTAGATACCCTGGTAGTCC
ACGCCGTAAACGATGTCAACTAGCCGTTGGGAGCCTTGAGCTCTTAGTGGCGCAGCTAACGCATTAAGTTGACCGCCTGG
GGAGTACGGCCGCAAGGTTAAAACTCAAATGAATTGACGGGGGCCCGCACAAGCGGTGGAGCATGTGGTTTAATTCGAAG
CAACGCGAAGAACCTTACCAGGCCTTGACATCCAATGAACTTTCCAGAGATGGATTGGTGCCTTCGGGAACATTGAGACA
GGTGCTGCATGGCTGTCGTCAGCTCGTGTCGTGAGATGTTGGGTTAAGTCCCGTAACGAGCGCAACCCTTGTCCTTAGTT
ACCAGCACGTAATGGTGGGCACTCTAAGGAGACTGCCGGTGACAAACCGGAGGAAGGTGGGGATGACGTCAAGTCATCAT
GGCCCTTACGGCCTGGGCTACACACGTGCTACAATGGTCGGTACAGAGGGTTGCCAAGCCGCGAGGTGGAGCTAATCCCA
CAAAACCGATCGTAGTCCGGATCGCAGTCTGCAACTCGACTGCGTGAAGTCGGAATCGCTAGTAATCGCGAATCAGAATG
TCGCGGTGAATACGTTCCCGGGCCTTGTACACACCGCCCGTCACACCATGGGAGTGGGTTGCACCAGAAGTAGCTAGTCT
AACCTTCGGGAGGACGGTTACCACGGTGTGATTCATGACTGGGGTGAAGTCGTAACAAGGTAGCCGTAGGGGAACCTGCG
GCTGGATCACCTCCTTAAATGGACGACAACAAGAAGAAAGCCTTGGCTGCGGCCCTGGGTCAGATCGAACGTCAATTCGG
CAAGGGTGCCGTAATGCGTATGGGCGATCACGACCGCCAGGCGATCCCGGCCATTTCCACTGGCTCTCTGGGTCTGGACA
TCGCACTCGGCATCGGCGGCCTGCCAAAGGGTCGTATTGTTGAAATCTACGGTCCGGAATCGTCCGGTAAAACCACCCTG
ACCCTGTCCGTGATTGCCCAGGCACAGAAGATGGGCGCCACTTGCGCCTTCGTCGACGCCGAGCATGCACTGGACCCGGA
ATACGCCGGCAAGCTGGGGGTCAACGTTGACGACCTGCTGGTTTCCCAGCCGGACACCGGTGAACAGGCACTGGAAATCA
CCGATATGCTGGTGCGCTCCAATGCCATCGACGTGATCGTGATCGACTCCGTGGCGGCACTGGTGCCCAAGGCCGAGATC
GAAGGCGAGATGGGCGACATGCACGTGGGCCTGCAGGCCCGCCTGATGTCCCAGGCGCTACGCAAGATCACCGGCAACAT
CAAGAACGCCAACTGCCTGGTGATCTTCATCAACCAGATCCGTATGAAAATCGGCGTGATGTTCGGCAGCCCGGAAACCA
CCACCGGTGGTAACGCGCTGAAGTTCTACGCTTCGGTTCGTCTGGATATCCGTCGTACTGGCGCGGTGAAGGAAGGTGAC
GAAGTCGTCGGTAGCGAAACCCGGGTCAAGATCGTCAAGAACAAGGTGGCTCCACCGTTCCGCCAGGCTGAATTCCAGAT
CCTGTACGGCAAGGGTATCTACCTGAACGGCGAGATCATCGATCTGGGCGTGCTGCATGGTTTCCTCGAGAAGTCCGGTG
CCTGGTACAGCTACCAGGGCAACAAGATCGGTCAGGGCAAGGCCAACTCGGCCAAGTTCCTGCAGGACAACCCGGAAATC
GGTAATGCCCTCGAGAAGCAGATTCGCGACAAGCTGCTGGCTCCGACCGCTGATGTCAAAGCTTCGCCGGTCAACGAGAC
CATCGATGACATGGCCGACGCGGATATCTGAATGAGCGAAGAAAACACGTACGACTCGAGCAGCATTAAAGTGCTGAAAG
GTTTGGATGCCGTACGCAAACGTCCCGGTATGTACATTGGTGACACCGACGATGGCAGCGGTCTGCACCATATGGTGTTC
GAGGTGGTCGATAACTCGATCGACGAAGCTCTGGCCGGCCATTGCGACGACATCAGCATCATCATCCACCCGGACGAATC
CATTACCGTGCGTGACAACGGTCGCGGCATCCCGGTAGACGTGCATAAAGAAGAAGGCGTTTCCGCAGCCGAGGTCATCA
TGACTGTGCTGCACGCCGGCGGTAAGTTCGACGATAACTCCTACAAAGTATCCGGCGGTCTGCACGGTGTGGGTGTGTCG
GTAGTGAACGCCCTGTCCGAAGAGCTGGTCCTGACCGTTCGCCGCAGTGGCAAGATCTGGGAACAGACCTACGTTCACGG
TGTGCCTCAGGCGCCTATGGCGATCGTCGGTGACAGCGAAACCACCGGTACCCAGATTCACTTCAAGGCTTCCAGCGAGA
CCTTCAAGAACATCCACTTCAGCTGGGACATCCTGGCCAAGCGGATTCGTGAACTGTCCTTCCTCAACTCCGGTGTCGGT
ATCGTTCTGAAGGACGAGCGCAGCGGCAAGGAAGAGCTGTTCAAGTACGAAGGCGGCTTGCGTGCGTTCGTTGAATACCT
GAACACCAACAAGACTGCGGTCAACCAGGTGTTCCACTTCAACGTGCAGCGTGAAGATGGCATCGGCGTGGAAATCGCCC
TGCAGTGGAACGACAGCTTCAACGAGAACCTGCAGTGCTTCACCAACAACATTCCGCAGCGCGACGGCGGCACCCACCTG
GTGGGCTTCCGTTCGGCGCTGACGCGTAACCTGAACAACTACATCGAGCAGGAAGGCCTGGCGAAGAAGCACAAGGTCGC
CACCACCGGTGACGATGCCCGCGAAGGCCTGACCGCGATCATTTCGGTCAAGGTGCCGGATCCGAAGTTCAGCTCCCAGA
CCAAAGACAAGCTGGTGTCTTCCGAAGTGAAGACCGCGGTCGAACAGGAAATGGGTAAGTACTTCTCCGACTTCCTGCTG
GAAAACCCGAACGAAGCCAAACTGGTGGTCGGCAAGATGCTCGACGCCGCCCGTGCCCGTGAAGCGGCGCGTAAAGCCCG
TGAGATGACCCGTCGTAAAGGCGCGCTGGATATCGCCGGCCTGCCGGGCAAACTGGCGGACTGCCAGGAAAAAGACCCTG
CCCTTTCCGAACTCTACCTGGTGGAAGGTGACTCTGCTGGCGGCTCCGCCAAGCAGGGACGCAACCGTAAGACCCAGGCG
ATCCTGCCGCTCAAGGGCAAGATCCTCAACGTCGAGAAAGCGCGCTTCGACAAGATGATTTCCTCGCAAGAGGTCGGCAC
CTTGATCACTGCACTCGGTTGCGGCATCGGCCGCGAAGAGTACAACATCGACAAGCTGCGTTATCACAACATCATCATCA
TGACCGACGCCGACGTCGACGGTTCGCACATCCGTACCCTACTCCTGACCTTCTTCTTCCGTCAGTTGCCGGAGCTGATC
```

GAGCGTGGCTACATCTACATCGCTCAACCGCCGCTGTACAAGGTCAAGAAAGGCAAGCAAGAGCAATACATCAAAGACGA
CGACGCCATGGAAGAGTACATGACGCAGTCGGCCCTGGAAGATGCGAGCCTGCACCTGAACGAAGAAGCACCGGGTATTT
CCGGCGAGGCGCTGGAGCGCCTGGTGAACGACTTCCGCATGGTCATGAAGACCCTCAAGCGTCTGTCGCGCCTCTACCCT
CAAGAGCTGACCGAGCACTTCATCTACCTGCCGGCCGTGAGCCTGGAGCAACTCTCCGATCACGCGGCGATGCAGGATTG
GCTGGCCCAATATGAAGTCCGCCTGCGCACCGTCGAGAAGTCCGGCCTGGTCTACAAGGCCAGCCTGCGTGAAGACCGTG
AACGTAATGTGTGGCTGCCAGAGGTCGAACTGATCTCCCACGGCCTGTCGAACTACGTCACCTTCAACCGCGACTTCTTC
GGCAGCAATGACTACAAGACCGTCGTCACCCTCGGCGCTCAACTGAGCTCCCTGCTGGACGAAGGTGCTTATATTCAGCG
CGGCGAGCGCAAGAAGGCGGTGACCGAGTTCAAGGAAGCCCTGGACTGGCTGATGACCGAAAGCACCAAGCGCCATACCA
TCCAGCGATACAAAGGTCTGGGTGAGATGAACCCGGATCAGCTGTGGGAAACCACCATGGACCCAAGCGTGCGCCGCATG
CTCAAGGTCACTATCGAAGACGCCATCGGCGCCGACCAGATCTTCAACACCCTGATGGGTGATGCGGTCGAGCCTCGTCG
TGACTTCATCGAAAGCAACGCCCTGGCGGTATCCAACCTGGACTTCTGAATGTCCGGAAAAGCGCAACAGCAGTCTCGCC
TCAAAGAGTTGATCAGCCGTGGTCGTGAGCAGGGTTACCTGACTTACGCGGAGGTCAACGACCACCTGCCGGAGGATATT
TCAGATCCGGAACAGGTGGAAGACATCATCCGCATGATCAACGACATGGGGATCAACGTATTCGAGAGTGCTCCGGATGC
GGATGCCCTTTTGTTGGCCGAAGCCGATACCGACGAAGCAGCAGCTGAAGAAGCCGCCGCAGCGTTGGCGGCTGTGGAAA
CCGACATTGGTCGCACTACCGACCCAGTGCGTATGTACATGCGCGAAATGGGTACGGTAGAGCTGCTCACACGTGAAGGC
GAAATCGAAATCGCCAAGCGTATCGAAGAGGGCATCCGTGAAGTGATGGGCGCGATCGCGCACTTCCCTGGCACGGTTGA
GCACATCCTCTCCGAATACACTCGCGTCACCACCGAAGGTGGTCGCCTGTCCGACGTCCTGAGCGGTTACATCGACCCGG
ACGACGGCATCGCGCCGCCTGCCGCCGAAGTACCACCGCCTGTCGATGCCAAGGCCGCGAAAGCGGACGACGACACCGAC
GACGATGACGCCGAAGCCAGTGACGACGAAGAAGAAGCCGAAAGCGGTCCGGATCCGGTCATCGCAGCCCAGCGCTTTGG
CGCCGTTGCCGACCAGATGGAAATCACTCGCAAGGCGCTGAAAAAGCACGGTCGTGAACACAAGCAAGCCCTGGCTGAAA
TGCTGGCCCTGGCTGAACTGTTCATGCCGATCAAACTGGTTCCGAAGCAATTCGAAGGCCTGGTTGAACGCGTTCGTAGC
GCCCTGGATCGCCTGCGTCAGCAAGAGCGCGCGATCATGCAACTCTGTGTTCGTGATGCTCGCATGCCACGCGCCGACTT
CCTGCGCCAGTTCCCTGGCAATGAAGTGGACGAAAGCTGGTCCGATGCACTGGCCAAAGGCAAGGCCAAGTACGCCGAAG
CCATCGGTCGCCTGCAACCGGACATCATCCGTTGCCAGCAGAAGCTGACCGCGCTCGAGACCGAGACCGGCCTGACGATC
GCCGAGATCAAGGACATCAACCGTCGCATGTCGATCGGCGAGGCCAAGGCCCGTCGCGCGAAGAAAGAGATGGTCGAAGC
CAACTTGCGCCTGGTGATCTCCATCGCCAAGAAGTACACCAACCGTGGCCTGCAGTTCCTGGACCTGATCCAGGAAGGCA
ACATCGGCTTGATGAAAGCGGTAGACAAGTTCGAATACCGTCGCGGCTACAAGTTCTCGACTTATGCCACCTGGTGGATC
CGTCAGGCGATCACTCGCTCGATCGCCGACCAGGCCCGCACCATCCGTATTCCGGTGCACATGATCGAGACGATCAACAA
GCTCAACCGTATTTCCCGGCAGATGCTGCAGGAAATGGGTCGCGAACCGACCCCGGAAGAGCTGGGCGAACGCATGGAAA
TGCCTGAGGACAAGATCCGCAAGGTATTGAAGATCGCTAAAGAGCCGATCTCCATGGAAACCCCGATCGGTGATGACGAA
GACTCCCATCTGGGCGACTTCATCGAAGACTCGACCATGCAGTCGCCAATCGATGTCGCCACCGTTGAGAGCCTCAAGGA
AGCGACTCGCGAAGTTCTCTCCGGCCTCACTGCCCGTGAAGCCAAGGTACTGCGCATGCGCTTCGGCATCGACATGAATA
CCGACCACACCCTCGAGGAAGTCGGTAAGCAGTTCGACGTTACCCGTGAGCGGATTCGCCAGATCGAAGCCAAGGCGCTG
CGCAAGCTGCGCCACCCGACGAGAAGCGAGCACCTGCGCTCCTTCCTCGACGAGTAATTGACTAAGCCAGCCATACTCGC
CCTTGCTGATGGCAGCATTTTTCGCGGCGAAGCCATTGGAGCCGACGGTCAAACCGTTGGTGAGGTGGTGTTTAACACCG
CAATGACCGGCTATCAGGAAATCCTTACCGATCCTTCCTACGCCCAACAGATCGTTACCCTGACTTACCCGCATATCGGC
AATACCGGCACCACGCCGGAAGACGCCGAGTCCGATCGTGTCTGGTCGGCCGGTCTGGTGATTCGCGACCTGCCACTGGT
TGCGAGCAACTGGCGTAACACCCTGTCCCTGTCCGATTACCTGAAAGCCAACAACGTTGTGGCGATCGCCGGTATCGACA
CCCGTCGCCTGACGCGCATCCTGCGCGAGAAAGGCGCGCAGAACGGCTGCATCATAGCCGGCGACAACATTTCCGACGAA
GCGGCGATTGCCGCAGCGCGCGGCTTCCCTGGCCTGAAAGGCATGGATTTGGCGAAGGTCGTCAGCACCAAGGAAAGCTA
CGAGTGGCGCTCCAGCGTCTGGAGCCTGAAGACCGACAGTCATCCGACTATCGAGGCTTCCGAGCTGCCGTACCACGTGG
TCGCCTACGACTACGGCGTCAAGCTGAACATCCTGCGCATGCTGGTCGAGCGCGGTTGCCGCGTGACCGTAGTGCCTGCG
CAAACTTCGGCCAGCGACGTCCTGGCACTCAAGCCTGACGGTGTGTTCCTGTCCAACGGCCCTGGCGACCCCGAGCCTTG
CGATTACGCCATCCAGGCGATCAAGGATGTGCTGGAAACCGAGATCCCGGTCTTCGGTATCTGCCTGGGCCACCAACTGC
TGGCACTGGCCTCCGGCGCCAAGACGGTGAAGATGGGCCACGGCCACCACGGTGCCAACCACCCGGTCCAGGACCTGGAC
AGCGGTGTAGTGATGATCACCAGCCAGAACCACGGTTTTGCGGTGGACGAAGCCACCCTGCCAGGCAACGTGCGGGCGAT
CCACAAATCGCTGTTCGACGGCACCCTGCAAGGCATCGAACGTACCGACAAGAGCGCATTCAGCTTCCAGGGCCACCCTG
AAGCGAGCCCGGGCCCGAACGATGTGGCGCCGCTGTTCGATCGTTTCATCAACGAGATGGCCAAGCGACGCTGAATGAGT
AGCGGACGTATCGTTCAAATCATCGGCGCCGTTATCGACGTGGAATTTCCACGCGACAGCGTACCGAGCATCTACGACGC
CTTGAAGGTTCAAGGCGCCGAAACCACTCTGGAAGTTCAGCAGCAGCTGGGCGACGGCGTGGTACGTACCATTGCGATGG
GCTCCACCGAGGGCTTGAAGCGCGGTCTGGACGTCAACAACACTGGCGCAGCCATCTCCGTACCGGTCGGTAAAGCGACC
CTGGGCCGGATCATGGACGTACTGGGCAACCCGATCGACGAAGCTGGTCCGATCGGCGAAGAAGAGCGTTGGGGCATTCA
CCGTCCTGCGCCGTCCTTCGCTGAACAAGCTGGCGGCAACGACCTCCTGGAAACCGGCATCAAGGTTATCGACCTGGTTT
GCCCGTTCGCCAAGGGCGGTAAAGTCGGTCTGTTTGGTGGTGCCGGTGTGGGCAAAACCGTAAACATGATGGAACTGATC
CGTAACATCGCCATCGAGCACAGCGGTTATTCCGTGTTCGCCGGTGTGGGTGAGCGTACTCGTGAGGGTAACGACTTCTA
CCACGAGATGAAGGATTCCAACGTTCTGGACAAAGTGGCACTGGTATACGGCCAGATGAACGAGCCGCCGGGAAACCGTC
TGCGCGTAGCTCTGACCGGCCTGACCATGGCCGAGAAGTTCCGTGACGAAGGTAACGACGTTCTGCTGTTCGTCGACAAC
ATCTATCGTTACACCCTGGCCGGTACCGAAGTATCCGCACTGCTGGGCCGTATGCCTTCCGCAGTAGGTTACCAGCCGAC
CCTGGCTGAAGAGATGGGCGTTCTGCAAGAACGTATCACTTCGACCAAGCAAGGCTCGATCACCTCGATCCAAGCGGTAT
ACGTACCTGCGGACGACTTGACCGACCCGTCGCCAGCGACCACCTTCGCCCACTTGGACGCCACCGTCGTACTGTCCCGT
GACATCGCTTCCCTGGGTATCTACCCAGCGGTAGATCCACTGGACTCGACTTCCCGTCAGCTGGACCCGAACGTGATCGG
CAACGAGCACTATGAAACCGCTCGCGGCGTTCAGTACGTGCTGCAGCGCTACAAAGAGCTGAAGGACATCATTGCGATCC
TGGGTATGGACGAACTGTCCGAAGCCGACAAGCAACTGGTATCCCGCGCTCGTAAGATCCAGCGCTTCCTGTCGCAGCCG
TTCTTCGTGGCTGAAGTCTTCACTGGTTCTCCAGGCAAATACGTTTCCCTGAAAGATACCATCGCTGGCTTCAAAGGCAT

CCTCAACGGTGACTACGACCACCTGCCAGAACAAGCGTTCTACATGGTCGGCGGCATCGAAGAAGCGATCGAGAAAGCCA
AGAAACTGTAA

NCBI Reference Sequence: NZ_ CP027714.1
Strain: ATCC17415
Chained genes: 16S rRNA, recA, gyrB, rpoD, carA, atpD
>NZ_CP027714_ATCC17415
GAACTGAAGAGTTTGATCATGGCTCAGATTGAACGCTGGCGGCAGGCCTAACACATGCAAGTCGAGCGGTAGAGAGAAGC
TTGCTTCTCTTGAGAGCGGCGGACGGGTGAGTAATGCCTAGGAATCTGCCTGGTAGTGGGGGATAACGTCCGGAAACGGA
CGCTAATACCGCATACGTCCTACGGGAGAAAGCAGGGGACCTTCGGGCCTTGCGCTATCAGATGAGCCTAGGTCGGATTA
GCTAGTTGGTGAGGTAATGGCTCACCAAGGCGACGATCCGTAACTGGTCTGAGAGGATGATCAGTCACACTGGAACTGAG
ACACGGTCCAGACTCCTACGGGAGGCAGCAGTGGGGAATATTGGACAATGGGCGAAAGCCTGATCCAGCCATGCCGCGTG
TGTGAAGAAGGTCTTCGGATTGTAAAGCACTTTAAGTTGGGAGGAAGGGTACTTACCTAATACGTGAGTATTTTGACGTT
ACCGACAGAATAAGCACCGGCTAACTCTGTGCCAGCAGCCGCGGTAATACAGAGGGTGCAAGCGTTAATCGGAATTACTG
GGCGTAAAGCGCGCGTAGGTGGTTCGTTAAGTTGGATGTGAAATCCCCGGGCTCAACCTGGGAACTGCATCCAAAACTGG
CGAGCTAGAGTATGGTAGAGGGTGGTGGAATTTCCTGTGTAGCGGTGAAATGCGTAGATATAGGAAGGAACACCAGTGGC
GAAGGCGACCACCTGGACTGATACTGACACTGAGGTGCGAAAGCGTGGGGAGCAAACAGGATTAGATACCCTGGTAGTCC
ACGCCGTAAACGATGTCAACTAGCCGTTGGGAGCCTTGAGCTCTTAGTGGCGCAGCTAACGCATTAAGTTGACCGCCTGG
GGAGTACGGCCGCAAGGTTAAAACTCAAATGAATTGACGGGGGCCCGCACAAGCGGTGGAGCATGTGGTTTAATTCGAAG
CAACGCGAAGAACCTTACCAGGCCTTGACATCCAATGAACTTTCCAGAGATGGATTGGTGCCTTCGGGAACATTGAGACA
GGTGCTGCATGGCTGTCGTCAGCTCGTGTCGTGAGATGTTGGGTTAAGTCCCGTAACGAGCGCAACCCTTGTCCTTAGTT
ACCAGCACATAATGGTGGGCACTCTAAGGAGACTGCCGGTGACAAACCGGAGGAAGGTGGGGATGACGTCAAGTCATCAT
GGCCCTTACGGCCTGGGCTACACACGTGCTACAATGGTCGGTACAGAGGGTTGCCAAGCCGCGAGGTGGAGCTAATCCCA
CAAAACCGATCGTAGTCCGGATCGCAGTCTGCAACTCGACTGCGTGAAGTCGGAATCGCTAGTAATCGCGAATCAGAATG
TCGCGGTGAATACGTTCCCGGGCCTTGTACACACCGCCCGTCACACCATGGGAGTGGGTTGCACCAGAAGTAGCTAGTCT
AACCTTCGGGAGGACGGTTACCACGGTGTGATTCATGACTGGGGTGAAGTCGTAACAAGGTAGCCGTAGGGGAACCTGCG
GCTGGATCACCTCCTTAAATGGACGACAACAAGAAGAAAGCCTTGGCTGCGGCCCTGGGTCAGATCGAACGTCAATTCGG
CAAGGGTGCCGTAATGCGTATGGGCGATCACGACCGCCAGGCGATCCCGGCCATTTCCACTGGCTCTCTGGGTCTGGACA
TCGCACTCGGCATCGGCGGCCTGCCAAAGGGCCGTATTGTTGAAATCTACGGTCCGGAATCGTCCGGTAAAACCACCCTG
ACCCTGTCCGTGATTGCCCAGGCACAGAAGATGGGCGCCACTTGCGCCTTCGTCGACGCCGAGCACGCACTGGACCCGGA
ATACGCCGGCAAGCTGGGGGTCAATGTTGACGACCTGCTGGTTTCCCAGCCGGACACCGGTGAACAGGCACTGGAAATCA
CCGACATGCTGGTGCGCTCCAATGCCATCGACGTGATCGTGATCGACTCCGTGGCGGCACTGGTGCCCAAGGCCGAGATC
GAAGGCGAGATGGGCGACATGCACGTGGGCCTGCAGGCCCGCCTGATGTCCCAGGCGCTGCGCAAGATCACCGGCAACAT
CAAGAACGCCAACTGCCTGGTGATCTTCATCAACCAGATCCGTATGAAAATCGGCGTGATGTTCGGCAGCCCGGAAACCA
CCACCGGTGGTAACGCGCTGAAGTTCTACGCTTCGGTTCGTCTGGATATCCGTCGTACTGGCGCGGTGAAGGAAGGTGAC
GAAGTCGTCGGTAGCGAAACCCGGGTCAAGATCGTCAAGAACAAGGTGGCTCCACCGTTCCGCCAGGCTGAATTCCAGAT
CCTGTACGGCAAGGGTATCTACCTGAACGGCGAGATCATCGATCTGGGCGTGCTGCACGGTTTCCTCGAGAAGTCCGGTG
CCTGGTACAGCTACCAGGGCAACAAGATCGGTCAGGGCAAGGCCAACTCGGCCAAGTTCCTGCAGGACAACCCGGAAATC
GGTAACGCCCTCGAGAAGCAGATTCGCGACAAGCTGCTGGCTCCGACCGCTGATGTCAAAGCTTCGCCGGTCAACGAGAC
CATCGATGACATGGCCGACGCGGATATCTGAATGAGCGAAGAAAACACGTACGACTCGAGCAGCATTAAAGTGCTGAAAG
GTTTGGATGCCGTACGCAAACGTCCCGGTATGTACATTGGTGACACCGACGATGGCAGCGGTCTGCACCATATGGTGTTC
GAGGTGGTCGATAACTCGATCGACGAAGCTCTGGCCGGCCATTGCGACGACATCAGCATCATCATCCACCCGGACGAATC
CATTACCGTGCGTGACAACGGTCGCGGCATCCCGGTAGACGTGCATAAAGAAGAAGGCGTTTCCGCAGCCGAGGTCATCA
TGACCGTACTGCACGCCGGCGGTAAGTTCGACGATAACTCCTACAAAGTATCCGGCGGTCTGCACGGTGTGGGTGTGTCG
GTAGTGAACGCCCTGTCCGAAGAACTGGTCCTGACCGTTCGCCGCAGTGGCAAGATCTGGGAACAGACCTACGTTCATGG
TGTGCCTCAGGCACCTATGGCGATCGTCGGTGACAGCGAAACCACTGGTACCCAGATTCACTTCAAGGCTTCCAGCGAGA
CCTTCAAGAACATCCACTTCAGCTGGGACATCCTGGCCAAGCGGATTCGTGAACTGTCCTTCCTCAACTCCGGTGTCGGT
ATCGTTCTGAAGGACGAGCGCAGCGGCAAGGAAGAGCTGTTCAAGTACGAAGGCGGCTTGCGTGCGTTCGTTGAATACCT
GAACACCAACAAGACTGCGGTCAACCAGGTGTTCCACTTCAATGTGCAGCGTGAAGACGGCATCGGCGTGGAAATCGCCC
TGCAGTGGAACGACAGCTTCAACGAGAACCTGCAGTGCTTTACCAACAACATTCCGCAGCGCGACGGCGGCACCCACCTG
GTGGGCTTCCGTTCGGCGCTTACGCGTAACCTGAACAACTACATCGAGCAGGAAGGCCTGGCGAAGAAGCACAAGGTCGC
CACCACCGGTGACGATGCCCGCGAAGGCCTGACCGCGATCATTTCGGTCAAGGTGCCGGATCCGAAGTTCAGCTCCCAGA
CCAAAGACAAGCTGGTTTCTTCCGAAGTGAAGACCGCAGTCGAACAGGAAATGGGCAAGTACTTCTCCGACTTCCTGCTG
GAAAACCCGAACGAAGCCAAGCTGGTGGTCGGCAAGATGCTCGACGCCGCCCGTGCCCGTGAAGCGGCGCGTAAAGCCCG
TGAGATGACCCGTCGTAAAGGCGCGCTGGATATCGCCGGCCTGCCGGGCAAACTGGCGGACTGCCAGGAAAAAGACCCTG
CCCTTTCCGAACTCTACCTGGTGGAAGGTGACTCTGCTGGCGGCTCCGCCAAGCAGGGACGCAACCGTAAGACCCAGGCG
ATCCTGCCGCTCAAGGGCAAGATCCTCAACGTCGAGAAAGCGCGCTTCGACAAGATGATTTCCTCGCAAGAGGTCGGCAC
CTTGATCACTGCACTCGGTTGCGGCATCGGTCGCGAAGAGTACAACATCGACAAGCTGCGTTATCACAACATCATCATCA
TGACCGACGCCGACGTCGACGGTTCGCACATCCGCACCCTGCTGCTGACCTTCTTCTTCCGTCAGTTGCCGGAGCTGATC
GAGCGTGGCTACATCTACATCGCTCAACCGCCGCTGTACAAGGTCAAGAAAGGCAAGCAAGAGCAATACATCAAAGACGA
CGACGCCATGGAAGAGTACATGACGCAGTCGGCCCTGGAAGATGCGAGCCTGCACCTGAACGAAGAAGCACCGGGTATTT
CCGGCGAGGCGCTGGAGCGCCTGGTGAACGACTTCCGCATGGTCATGAAGACCCTCAAGCGTCTGTCGCGCCTCTACCCT
CAAGAGCTGACCGAGCACTTCATCTACCTGCCGGCCGTGAGCCTGGAGCAACTCTCCGATCACGCGGCGATGCAGGATTG
GCTGGCCCAATATGAAGTCCGCCTGCGCACCGTCGAGAAGTCCGGCCTGGTCTACAAGGCCAGCCTGCGTGAAGACCGTG

AACGTAATGTGTGGCTGCCAGAGGTCGAACTGATCTCCCACGGTCTGTCGAACTACGTCACCTTCAACCGCGACTTCTTC
GGCAGCAATGACTACAAGACCGTCGTCACCCTCGGCGCTCAACTGAGCTCCCTGCTGGACGAAGGCGCTTATATTCAGCG
CGGCGAGCGCAAGAAGGCGGTGACCGAGTTCAAGGAAGCCCTGGACTGGCTGATGACCGAAAGCACCAAGCGCCACACCA
TCCAGCGATACAAAGGTCTGGGCGAGATGAACCCGGATCAGCTGTGGGAAACCACCATGGACCCAAGCGTGCGCCGCATG
CTCAAGGTCACCATCGAAGACGCCATCGGCGCCGACCAGATCTTCAACACCCTGATGGGTGATGCGGTCGAGCCTCGTCG
TGACTTCATCGAAAGCAACGCTCTGGCGGTATCCAACCTGGACTTCTGAATGTCCGGAAAAGCGCAACAGCAGTCTCGCC
TCAAAGAGTTGATCAGCCGTGGTCGTGAGCAGGGTTACCTGACTTACGCGGAGGTCAACGACCACCTGCCGGAGGATATT
TCAGATCCGGAACAGGTGGAAGACATCATCCGCATGATCAACGACATGGGGATCAACGTATTCGAGAGTGCTCCGGATGC
GGATGCCCTTTTGTTGGCCGAAGCCGATACCGACGAAGCAGCAGCTGAAGAAGCCGCCGCAGCGTTGGCGGCTGTGGAAA
CCGACATTGGGCGCACTACCGACCCAGTGCGTATGTACATGCGCGAAATGGGTACGGTAGAGCTGCTCACACGTGAAGGC
GAAATCGAAATCGCCAAGCGTATCGAAGAGGGTATCCGTGAAGTGATGGGCGCGATCGCGCACTTCCCTGGCACGGTTGA
GCACATCCTCTCCGAATACACTCGCGTCACCACCGAAGGTGGTCGCCTGTCCGACGTCCTGAGCGGTTACATCGACCCGG
ACGACGGCATCGCGCCGCCTGCCGCCGAAGTACCACCGCCTGTCGATGCCAAGGCCGCGAAAGCGGACGACGACACCGAC
GACGATGACGCCGAAGCCAGTGACGACGAAGAAGAAGCCGAAAGCGGTCCGGATCCGGTCATTGCAGCCCAGCGCTTTGG
CGCCGTTGCCGACCAGATGGAAATCACTCGCAAGGCGCTGAAAAAGCACGGTCGTGAACACAAGCAAGCCCTGGCTGAAA
TGCTGGCCCTGGCTGAACTGTTCATGCCGATCAAACTGGTTCCGAAGCAATTCGAAGGCCTGGTTGAACGCGTTCGTAGC
GCCCTGGATCGCCTGCGTCAGCAAGAGCGCGCGATCATGCAACTCTGTGTTCGTGATGCTCGCATGCCACGCGCCGACTT
CCTGCGCCAGTTCCCTGGCAATGAAGTGGACGAAAGCTGGTCCGACGCACTGGCCAAAGGCAAGGCCAAGTACGCCGAAG
CCATCGGTCGCCTGCAGCCGGACATCATCCGTTGCCAGCAGAAGCTGACCGCGCTCGAGACCGAGACCGGCCTGACGATC
GCCGAGATCAAGGACATCAACCGTCGCATGTCGATCGGCGAGGCCAAGGCCCGTCGCGCGAAGAAAGAGATGGTCGAAGC
CAACTTGCGTCTGGTGATCTCCATCGCCAAGAAGTACACCAACCGTGGCCTGCAGTTCCTCGACCTGATCCAGGAAGGCA
ACATCGGCTTGATGAAAGCGGTAGACAAGTTCGAATACCGTCGCGGCTACAAGTTCTCGACTTATGCCACCTGGTGGATC
CGTCAGGCGATCACTCGCTCGATCGCCGACCAGGCCCGCACCATCCGTATTCCGGTGCACATGATCGAGACGATCAACAA
GCTCAACCGTATTTCCCGGCAGATGCTGCAGGAAATGGGTCGCGAACCGACCCCGGAAGAGCTGGGCGAACGCATGGAAA
TGCCTGAGGACAAGATCCGCAAGGTATTGAAGATCGCTAAAGAGCCGATCTCCATGGAAACCCCGATCGGTGATGACGAA
GACTCCCATCTGGGCGACTTCATCGAAGACTCGACCATGCAGTCGCCAATCGATGTCGCCACCGTTGAGAGCCTCAAGGA
AGCGACTCGCGAAGTTCTCTCCGGCCTCACTGCCCGTGAAGCCAAGGTACTGCGCATGCGCTTCGGCATCGACATGAATA
CCGACCACACCCTCGAGGAAGTCGGTAAGCAGTTCGACGTTACCCGTGAGCGGATTCGCCAGATCGAAGCCAAGGCGCTG
CGCAAGCTACGCCATCCGACGAGAAGCGAGCACCTGCGCTCCTTCCTCGACGAGTAATTGACTAAGCCAGCCATACTCGC
CCTTGCTGATGGCAGCATTTTTCGCGGCGAAGCCATTGGAGCCGACGGTCAAACCGTTGGTGAGGTGGTGTTTAACACCG
CAATGACCGGCTATCAGGAAATCCTTACCGATCCTTCCTACGCCCAACAGATCGTTACCCTGACTTACCCGCATATCGGC
AATACCGGCACCACGCCGGAAGACGCCGAGTCCGATCGTGTCTGGTCGGCCGGTCTGGTGATTCGCGACCTGCCACTGGT
TGCGAGCAACTGGCGTAATACCCTGTCCCTGTCCGACTACCTGAAAGCCAACAACGTTGTGGCGATCGCCGGTATCGACA
CCCGTCGCCTGACGCGCATCCTGCGCGAGAAAGGCGCGCAGAACGGCTGCATCATGGCCGGCGACAATATCTCCGACGAA
GCGGCGATTGCCGCAGCGCGCGGCTTCCCTGGTCTGAAAGGCATGGATCTGGCGAAGGTCGTCAGCACCAAGAAAACCTA
CGAGTGGCGCTCCAGCGTCTGGAGCCTGAAGACCGACAGTCATCCGACTATCGAGGCTTCCGAGCTGCCGTACCACGTGG
TCGCCTACGACTACGGCGTCAAGCTGAACATCCTGCGCATGCTGGTCGAGCGTGGTTGCCGCGTGACCGTAGTGCCTGCG
CAAACTCCGGCCAGCGACGTCCTGGCACTCAAGCCTGACGGTGTGTTCCTGTCCAACGGCCCTGGCGACCCCGAGCCTTG
CGATTACGCCATCCAGGCGATTAAGGACGTGCTGGAAACCGAGATCCCGGTCTTCGGTATCTGCCTGGGCCACCAACTGC
TGGCATTGGCCTCCGGCGCCAAGACGGTGAAGATGGGCCACGGCCACCACGGTGCCAACCACCCGGTCCAGGACCTGGAC
AGCGGTGTAGTGATGATCACCAGCCAGAACCACGGTTTTGCGGTGGACGAAGCCACCCTGCCAGGCAACGTGCGGGCGAT
CCACAAATCGCTGTTCGACGGCACCCTGCAAGGCATCGAACGTACCGACAAGAGCGCATTCAGCTTCCAGGGCCACCCTG
AAGCGAGCCCGGGCCCGAACGATGTGGCGCCGCTGTTCGATCGTTTCATCAACGAGATGGCCAAGCGACGCTGAATGAGT
AGCGGACGTATCGTTCAAATCATCGGCGCCGTTATCGACGTGGAATTTCCACGCGACAGCGTACCGAGCATCTACGACGC
CTTGAAGGTTCAAGGCGCCGAAACCACTCTGGAAGTTCAGCAGCAGCTGGGCGACGGCGTGGTACGTACCATTGCGATGG
GCTCCACCGAGGGCTTGAAGCGCGGTCTGGACGTCAACAACACTGGCGCAGCCATCTCCGTACCGGTCGGTAAAGCGACC
CTGGGCCGGATCATGGACGTACTGGGCAACCCGATCGACGAAGCTGGCCCGATCGGCGAAGAAGAGCGTTGGGGCATTCA
CCGTCCTGCGCCGTCCTTCGCTGAACAAGCTGGCGGCAACGACCTCCTGGAAACCGGCATCAAGGTTATCGACCTGGTTT
GCCCGTTCGCCAAGGGCGGTAAAGTCGGTCTGTTCGGTGGTGCCGGTGTGGGCAAAACCGTAAACATGATGGAACTGATC
CGTAACATCGCCATCGAGCACAGCGGTTATTCCGTGTTCGCCGGTGTGGGTGAGCGTACTCGTGAGGGTAACGACTTCTA
CCACGAGATGAAGGATTCCAACGTTCTGGACAAAGTGGCACTGGTATACGGCCAGATGAACGAGCCGCCGGGAAACCGTC
TGCGCGTAGCTCTGACCGGCCTGACCATGGCCGAGAAGTTCCGTGACGAAGGTAACGACGTTCTGCTGTTCGTCGACAAC
ATCTATCGTTACACCCTGGCCGGTACCGAAGTATCCGCACTGCTGGGCCGTATGCCTTCCGCAGTAGGTTACCAGCCGAC
CCTGGCTGAAGAGATGGGCGTTCTGCAAGAACGTATCACTTCGACCAAGCAAGGCTCGATCACCTCGATCCAAGCGGTAT
ACGTACCTGCGGACGACTTGACCGACCCGTCGCCAGCGACCACCTTCGCCCACTTGGACGCCACCGTCGTACTGTCCCGT
GACATCGCTTCCCTGGGTATCTACCCAGCGGTAGATCCACTGGACTCGACTTCCCGTCAGCTGGACCCGAACGTGATCGG
CAACGAGCACTATGAAACCGCTCGCGGCGTTCAGTACGTGCTGCAGCGCTACAAAGAGCTGAAGGACATCATTGCGATCC
TGGGTATGGACGAACTGTCCGAAGCCGACAAGCAACTGGTATCCCGCGCTCGTAAGATCCAGCGCTTCCTGTCGCAGCCG
TTCTTCGTGGCTGAAGTCTTCACTGGTTCTCCAGGCAAATACGTTTCCCTGAAAGACACCATCGCTGGCTTCAAAGGCAT
CCTCAACGGTGACTACGACCACCTGCCAGAACAAGCGTTCTACATGGTCGGCGGCATCGAAGAAGCGATCGAGAAAGCCA
AGAAACTGTAA

NCBI Reference Sequence: NZ_CP027735.1
Strain: DTR133

Chained genes: 16S rRNA, recA, gyrB, rpoD, carA, atpD
>NZ_CP027735_DTR133
GAACTGAAGAGTTTGATCATGGCTCAGATTGAACGCTGGCGGCAGGCCTAACACATGCAAGTCGAGCGGTAGAGAGGTGC
TTGCACCTCTTGAGAGCGGCGGACGGGTGAGTAATGCCTAGGAATCTGCCTGGTAGTGGGGGATAACGTTCGGAAACGGA
CGCTAATACCGCATACGTCCTACGGGAGAAAGCAGGGGACCTTCGGGCCTTGCGCTATCAGATGAGCCTAGGTCGGATTA
GCTAGTTGGTGAGGTAATGGCTCACCAAGGCGACGATCCGTAACTGGTCTGAGAGGATGATCAGTCACACTGGAACTGAG
ACACGGTCCAGACTCCTACGGGAGGCAGCAGTGGGGAATATTGGACAATGGGCGAAAGCCTGATCCAGCCATGCCGCGTG
TGTGAAGAAGGTCTTCGGATTGTAAAGCACTTTAAGTTGGGAGGAAGGGTACTTACCTAATACGTGAGTATTTTGACGTT
ACCGACAGAATAAGCACCGGCTAACTCTGTGCCAGCAGCCGCGGTAATACAGAGGGTGCAAGCGTTAATCGGAATTACTG
GGCGTAAAGCGCGCGTAGGTGGTTCGTTAAGTTGAATGTGAAATCCCCGGGCTCAACCTGGGAACTGCATCCAAAACTGG
CGAGCTAGAGTATGGTAGAGGGTGGTGGAATTTCCTGTGTAGCGGTGAAATGCGTAGATATAGGAAGGAACACCAGTGGC
GAAGGCGACCACCTGGACTGATACTGACACTGAGGTGCGAAAGCGTGGGGAGCAAACAGGATTAGATACCCTGGTAGTCC
ACGCCGTAAACGATGTCAACTAGCCGTTGGGAGCCTTGAGCTCTTAGTGGCGCAGCTAACGCATTAAGTTGACCGCCTGG
GGAGTACGGCCGCAAGGTTAAAACTCAAATGAATTGACGGGGGCCCGCACAAGCGGTGGAGCATGTGGTTTAATTCGAAG
CAACGCGAAGAACCTTACCAGGCCTTGACATCCAATGAACTTTCCAGAGATGGATTGGTGCCTTCGGGAACATTGAGACA
GGTGCTGCATGGCTGTCGTCAGCTCGTGTCGTGAGATGTTGGGTTAAGTCCCGTAACGAGCGCAACCCTTGTCCTTAGTT
ACCAGCACGTAATGGTGGGCACTCTAAGGAGACTGCCGGTGACAAACCGGAGGAAGGTGGGGATGACGTCAAGTCATCAT
GGCCCTTACGGCCTGGGCTACACACGTGCTACAATGGTCGGTACAGAGGGTTGCCAAGCCGCGAGGTGGAGCTAATCCCA
TAAAACCGATCGTAGTCCGGATCGCAGTCTGCAACTCGACTGCGTGAAGTCGGAATCGCTAGTAATCGCGAATCAGAATG
TCGCGGTGAATACGTTCCCGGGCCTTGTACACACCGCCCGTCACACCATGGGAGTGGGTTGCACCAGAAGTAGCTAGTCT
AACCTTCGGGAGGACGGTTACCACGGTGTGATTCATGACTGGGGTGAAGTCGTAACAAGGTAGCCGTAGGGGAACCTGCG
GCTGGATCACCTCCTTAAATGGACGACAACAAGAAGAAAGCCTTGGCTGCGGCCCTGGGTCAGATCGAACGTCAATTCGG
CAAGGGTGCCGTAATGCGTATGGGCGATCACGATCGCCAGGCGATCCCGGCCATTTCCACTGGCTCTCTGGGTCTGGACA
TCGCACTCGGCATCGGCGGCCTGCCAAAAGGCCGTATTGTTGAAATCTACGGTCCGGAATCGTCCGGTAAAACCACCCTG
ACCCTGTCGGTGATTGCCCAGGCACAGAAGATGGGCGCCACCTGCGCCTTCGTCGACGCCGAGCACGCACTGGACCCGGA
GTACGCCGGCAAACTGGGGGTCAACGTTGACGACCTGCTGGTTTCCCAGCCGGACACCGGCGAACAGGCGCTGGAAATCA
CCGACATGCTGGTGCGCTCCAATGCCATCGACGTGATCGTGATCGACTCCGTGGCGGCGCTGGTACCCAAGGCCGAGATC
GAAGGCGAGATGGGCGACATGCACGTGGGCCTGCAGGCTCGCCTGATGTCCCAGGCGCTGCGCAAGATCACCGGTAACAT
CAAGAACGCCAACTGCCTGGTGATCTTCATCAACCAGATCCGTATGAAAATCGGCGTGATGTTCGGCAGCCCGGAAACCA
CCACCGGTGGTAACGCGCTGAAGTTCTACGCTTCGGTTCGTCTGGATATCCGTCGTACTGGCGCGGTGAAGGAAGGTGAC
GAAGTCGTCGGTAGCGAAACCCGGGTCAAGATCGTCAAGAACAAGGTGGCTCCACCCTTCCGTCAAGCTGAGTTCCAGAT
CCTGTACGGCAAGGGTATCTACCTGAACGGCGAGATCATCGATCTGGGCGTGCTGCACGGTTTCCTAGAGAAGTCCGGTG
CCTGGTACAGCTACCAGGGCAACAAGATCGGTCAGGGCAAGGCCAACTCGGCCAAGTTCCTGCAGGACAATCCGGAAATC
GGTAATGCCCTCGAGAAGCAGATTCGCGACAAGCTGCTGGCTCCGAGCGGAGATACCAAGGCTCTGCCCGTCAACGAGAC
CATCGATGACATGGCCGACGCGGATATCTGAATGAGCGAAGAAAACACGTACGACTCGAGCAGCATTAAAGTGCTGAAAG
GTTTGGATGCCGTACGCAAACGTCCCGGTATGTACATTGGTGACACCGACGATGGCAGCGGTCTGCACCATATGGTGTTC
GAGGTGGTCGACAACTCGATCGACGAAGCTCTGGCCGGCCACTGCGACGACATCAGCATCATCATCCACCCGGACGAATC
CATTACCGTGCGTGACAACGGTCGCGGCATCCCGGTAGACGTGCATAAAGAAGAAGGCGTTTCCGCAGCCGAGGTCATCA
TGACCGTGCTGCACGCCGGCGGTAAGTTCGACGACAACTCCTACAAGGTATCCGGCGGTCTGCACGGTGTGGGTGTGTCG
GTAGTGAATGCCCTGTCCGAAGAACTGGTGCTGACCGTTCGCCGCAGTGGCAAGATCTGGGAACAGACCTACGTTCACGG
TGTGCCTCAGGCGCCTATGGCGATCGTCGGTGACAGCGAGACCACTGGTACCCAGATTCACTTCAAGGCTTCCAGCGAGA
CCTTCAAGAACATCCATTTCAGCTGGGACATCCTGGCCAAGCGGATTCGTGAACTGTCCTTCCTCAACTCCGGTGTCGGT
ATCGTTCTGAAGGACGAGCGCAGCGGCAAGGAAGAACTGTTCAAATACGAAGGCGGCCTGCGCGCGTTCGTCGAATACCT
GAACACCAACAAGACTGCGGTCAACCAGGTGTTCCACTTCAACGTGCAGCGTGAAGACGGCATCGGCGTGGAAATCGCCC
TGCAGTGGAACGACAGCTTCAACGAAAACCTGCAGTGCTTCACCAACAACATTCCGCAGCGCGACGGCGGTACTCACCTG
GTGGGCTTCCGCTCGGCACTGACGCGTAACCTGAACAACTACATCGAGCAGGAAGGTCTGGCGAAGAAGCACAAGGTCGC
CACCACCGGTGACGATGCCCGCGAAGGCCTGACCGCGATCATTTCGGTCAAGGTGCCGGATCCGAAGTTCAGCTCCCAGA
CCAAAGACAAGCTGGTGTCTTCCGAAGTAAAGACCGCGGTCGAACAGGAAATGGGCAAGTACTTCTCCGACTTCCTGCTG
GAAAACCCGAACGAAGCCAAGCTGGTGGTCGGCAAGATGCTCGACGCCGCCCGCGCCCGTGAAGCGGCGCGTAAGGCCCG
TGAGATGACCCGCCGTAAAGGTGCGCTGGATATCGCCGGCCTGCCGGGCAAACTGGCGGACTGCCAGGAAAAGGACCCTG
CCCTTTCCGAACTCTACCTGGTGGAAGGTGACTCTGCTGGCGGCTCCGCCAAGCAGGGACGCAACCGCAAGACCCAGGCG
ATTCTGCCGCTCAAGGGCAAGATTCTTAACGTCGAGAAAGCGCGCTTCGACAAGATGATTTCCTCGCAAGAGGTCGGCAC
CTTGATCACTGCACTCGGCTGCGGCATCGGCCGCGAAGAGTACAACATCGACAAGCTGCGTTATCACAACATCATCATCA
TGACCGACGCCGACGTCGACGGTTCGCACATCCGCACCCTGCTGCTGACTTTCTTCTTCCGTCAGCTGCCGGAGCTGATC
GAGCGTGGCTACATCTACATCGCTCAACCGCCGCTGTACAAGGTCAAGAAAGGCAAGCAAGAGCAATACATCAAAGACGA
CGACGCCATGGAAGAGTACATGACGCAGTCGGCCCTGGAAGATGCGAGCCTGCACCTGAACGAAGAAGCACCGGGTATTT
CCGGCGAGGCGCTGGAGCGCCTGGTGAACGACTTCCGCATGGTCATGAAAACCCTCAAGCGTCTGTCGCGCCTGTACCCT
CAGGAGCTGACCGAACACTTCATCTACCTGCCAGCCGTGAGCCTGGAGCAACTCTCCGATCACGCAGCGATGCAGGATTG
GTTGGCCCAATATGAAGTCCGCCTGCGCACCGTCGAGAAGTCCGGCCTGGTCTACAAGGCCAGCCTGCGTGAAGACCGTG
AACGTAATGTCTGGCTGCCAGAGGTCGAACTGATCTCCCACGGCCTGTCGAACTACGTCACCTTCAACCGCGACTTCTTC
GGCAGCAATGACTACAAGACCGTCGTCACCCTCGGCGCTCAACTGAGCTCCCTGCTGGACGAAGGCGCTTATATTCAGCG
CGGCGAACGCAAGAAGGCGGTGACCGAGTTCAAGGAAGCCCTGGACTGGCTGATGACCGAAAGCACCAAGCGCCACACCA
TCCAGCGATACAAAGGTCTGGGCGAGATGAACCCGGACCAGCTGTGGGAAACCACCATGGACCCAAGCGTGCGCCGCATG
CTCAAGGTCACCATCGAAGACGCCATCGGCGCCGACCAGATCTTCAACACCCTGATGGGTGATGCGGTCGAGCCTCGTCG

CGACTTCATCGAAAGCAACGCCCTGGCGGTATCCAACCTGGACTTCTGAATGTCCGGAAAAGCGCAACAGCAGTCTCGCC
TCAAAGAGTTGATCAGCCGTGGTCGTGAGCAGGGTTACCTGACTTACGCGGAGGTCAACGACCACCTGCCGGAGGATATT
TCAGATCCGGAACAGGTGGAAGACATCATCCGCATGATCAACGACATGGGGATCAACGTATTCGAGAGTGCTCCGGATGC
GGATGCCCTTTTGTTGGCCGAAGCCGATACCGACGAAGCAGCAGCTGAAGAAGCCGCCGCAGCGTTGGCGGCTGTGGAAA
CCGACATTGGTCGCACTACCGACCCAGTGCGTATGTACATGCGCGAAATGGGCACGGTAGAGCTGCTCACACGTGAAGGC
GAAATCGAAATCGCCAAGCGTATCGAAGAGGGCATCCGTGAAGTGATGGGCGCGATCGCGCACTTCCCTGGCACGGTTGA
GCACATCCTCTCCGAATACACTCGCGTCACCACCGAAGGTGGCCGCCTGTCCGACGTCCTGAGCGGTTACATCGACCCGG
ATGACGGCATTGCGCCGCCTGCCGCCGAAGTACCACCGCCTGTCGATGCCAAGGCCGCGAAAGCGGACGACGACACCGAC
GACGATGACGCCGAAGCCAGTGACGACGAAGAAGAAGCCGAAAGCGGTCCGGATCCGGTCATCGCAGCCCAGCGCTTTGG
CGCCGTTGCCGACCAGATGGAAATCACCCGCAAGGCGCTGAAAAAGCACGGTCGTGAACACAAGCAAGCCCTGGCTGAAA
TGCTGGCCCTGGCTGAACTGTTCATGCCAATCAAACTGGTTCCGAAGCAATTCGAAGGCCTGGTTGAACGTGTTCGTAGC
GCCCTGGATCGCCTGCGTCAGCAAGAGCGCGCGATCATGCAGCTCTGTGTTCGTGATGCCCGCATGCCACGCGCCGACTT
CCTGCGCCAGTTCCCTGGCAATGAAGTGGACGAAAGCTGGTCCGACGCGCTGGCCAAAGGCAAGGCCAAGTACGCCGAAG
CCATCGGCCGCCTGCAGCCGGACATCATCCGTTGCCAGCAGAAGCTGACCGCGCTCGAGACCGAGACCGGCCTGACGATT
GCCGAGATCAAGGACATCAACCGTCGCATGTCGATCGGCGAGGCCAAGGCCCGTCGCGCGAAGAAAGAGATGGTCGAAGC
CAACTTGCGCCTGGTGATCTCCATCGCCAAGAAGTACACCAACCGTGGCCTGCAGTTCCTCGACCTGATCCAGGAAGGCA
ACATCGGTTTGATGAAAGCGGTAGACAAGTTCGAATACCGTCGCGGCTACAAGTTCTCGACTTATGCCACCTGGTGGATC
CGTCAGGCGATCACTCGCTCGATCGCCGACCAGGCCCGCACCATCCGTATTCCGGTGCACATGATCGAGACGATCAACAA
GCTCAACCGTATTTCCCGGCAGATGTTGCAGGAAATGGGTCGCGAACCGACTCCGGAAGAGCTGGGCGAACGCATGGAAA
TGCCTGAGGACAAGATCCGCAAGGTATTGAAGATCGCTAAAGAGCCGATCTCCATGGAAACCCCGATCGGTGATGACGAA
GACTCCCATCTGGGTGACTTCATCGAAGACTCGACCATGCAGTCGCCAATCGATGTCGCCACCGTTGAGAGCCTCAAGGA
AGCGACTCGCGAAGTCCTCTCCGGCCTCACTGCCCGTGAAGCCAAGGTACTGCGCATGCGCTTCGGCATCGACATGAATA
CCGATCACACCCTTGAGGAAGTCGGTAAGCAGTTCGACGTTACCCGTGAGCGGATTCGTCAGATCGAAGCCAAGGCGCTG
CGCAAGCTGCGCCACCCGACGAGAAGCGAGCATCTGCGCTCCTTCCTCGACGAGTAATTGACTAAGCCAGCCATACTCGC
CCTTGCTGATGGCAGCATTTTTCGCGGCGAAGCCATTGGAGCCGACGGTCAAACCGTTGGTGAGGTGGTGTTTAACACCG
CAATGACCGGCTATCAGGAAATCCTTACCGATCCTTCCTACGCCCAACAGATCGTTACCCTGACTTACCCGCACATCGGC
AACACTGGCACCACGCCGGAAGACGCCGAGTCCGATCGTGTCTGGTCGGCCGGTCTGGTGATTCGCGACCTGCCACTGGT
TGCGAGCAACTGGCGTAACACCCTGTCCCTGTCCGACTACCTGAAAGCCAACAATGTCGTGGCGATCGCCGGTATCGACA
CCCGTCGCCTGACGCGCATCCTGCGTGAAAAAGGCGCGCAGAACGGCTGCATCATGGCCGGCGACAATATCTCCGACGAA
GCAGCGATTGCCGCAGCGCGCGGCTTCCCTGGCCTGAAAGGCATGGATCTGGCGAAGGTCGTCAGCACCAAGGAAAGCTA
CGAGTGGCGCTCCAGCGTCTGGAGCCTGAAGACCGACAGTCATCCGACTATCGAAGCTTCCGAGCTGCCTTACCACGTGG
TTGCCTACGACTACGGCGTCAAGCTGAACATCCTGCGCATGTTGGTCGAGCGCGGCTGCCGCGTGACCGTAGTGCCTGCG
CAAACTCCGGCCAGCGACGTCCTGGCACTCAAGCCTGACGGTGTGTTCCTGTCCAACGGTCCTGGCGACCCCGAGCCTTG
TGATTACGCCATCCAGGCGATCAAGGACGTGCTGGAAACCGAGATCCCGGTCTTCGGTATCTGTCTGGGTCACCAATTGC
TGGCTCTGGCCTCCGGTGCCAAGACAGTGAAGATGGGCCACGGCCACCATGGCGCCAACCACCCGGTCCAGGACCTGGAC
AGCGGTGTAGTGATGATCACCAGCCAAAACCACGGTTTTGCGGTGGACGAAACTACCCTGCCAGGCAACGTGCGGGCGAT
CCACAAGTCGCTGTTCGATGGCACCCTGCAAGGCATCGAGCGTACCGACAAGAGCGCATTCAGCTTCCAGGGCCACCCTG
AAGCGAGCCCGGGCCCGAACGATGTGGCGCCGCTGTTCGATCGTTTCATCAACGAGATGGCCAAGCGACGCTGAATGAGT
AGCGGACGTATCGTTCAAATCATCGGCGCCGTTATCGACGTGGAATTTCCACGCGACAGCGTACCGAGCATCTACGACGC
CTTGAAGGTTCAAGGCGCCGAAACCACTCTGGAAGTTCAGCAGCAGCTGGGCGACGGCGTGGTACGTACCATTGCGATGG
GCTCCACCGAGGGCTTGAAGCGCGGTCTGGACGTCAACAACACTGGCGCAGCCATCTCCGTACCGGTCGGTAAAGCGACC
CTGGGCCGGATCATGGACGTACTGGGCAACCCGATCGACGAAGCTGGCCCGATCGGCGAAGAAGAGCGTTGGGGCATTCA
CCGTCCTGCGCCGACCTTCGCTGAACAAGCTGGCGGCAACGACCTCCTGGAAACCGGCATCAAGGTTATCGACCTGGTTT
GCCCGTTCGCCAAGGGCGGTAAAGTCGGTCTGTTCGGTGGTGCCGGTGTGGGCAAAACCGTAAACATGATGGAACTGATC
CGTAACATCGCCATCGAGCACAGCGGTTATTCCGTGTTCGCCGGTGTGGGTGAGCGTACTCGTGAGGGTAACGACTTCTA
CCACGAGATGAAGGATTCCAACGTTCTGGACAAAGTGGCACTGGTATACGGCCAGATGAACGAGCCGCCGGGAAACCGTC
TGCGCGTAGCTCTGACCGGCCTGACCATGGCCGAGAAGTTCCGTGACGAAGGTAACGACGTTCTGCTGTTCGTCGACAAC
ATCTATCGTTACACCCTGGCCGGTACCGAAGTATCCGCACTGCTGGGCCGTATGCCTTCGGCAGTAGGTTACCAGCCGAC
CCTGGCTGAAGAGATGGGCGTTCTGCAAGAACGTATCACTTCGACCAAGCAAGGCTCGATCACCTCGATCCAAGCGGTAT
ACGTACCTGCGGACGACTTGACCGACCCGTCGCCAGCGACCACCTTCGCCCACTTGGACGCCACCGTCGTACTGTCCCGT
GACATCGCTTCCCTGGGTATCTACCCAGCGGTAGACCCACTGGATTCGACTTCCCGTCAGCTGGACCCGAACGTGATCGG
CAACGAGCACTACGAAACCGCTCGCGGCGTTCAGTACGTGCTGCAGCGCTACAAAGAGCTGAAGGACATCATTGCGATCC
TGGGTATGGACGAACTGTCCGAAGCCGACAAGCAACTGGTATCCCGCGCTCGTAAGATCCAGCGCTTCCTGTCGCAGCCG
TTCTTCGTGGCTGAAGTCTTCACTGGTTCTCCAGGCAAATACGTTTCCCTGAAAGACACCATCGCTGGCTTCAAAGGCAT
CCTCAACGGTGACTACGACCACCTGCCAGAACAAGCGTTCTACATGGTCGGCGGCATCGAAGAAGCGATCGAGAAAGCCA
AGAAACTGTAA

NCBI Reference Sequence: NZ_CP027748.1
Strain: ChPhzS23
Chained genes: 16S rRNA, recA, gyrB, rpoD, carA, atpD
>NZ_CP027748_ChPhzS23
GAACTGAAGAGTTTGATCATGGCTCAGATTGAACGCTGGCGGCAGGCCTAACACATGCAAGTCGAGCGGTAGAGAGAAGC
TTGCTTCTCTTGAGAGCGGCGGACGGGTGAGTAATGCCTAGGAATCTGCCTGGTAGTGGGGGATAACGTTCGGAAACGGA
CGCTAATACCGCATACGTCCTACGGGAGAAAGCAGGGGACCTTCGGGCCTTGCGCTATCAGATGAGCCTAGGTCGGATTA

GCTAGTTGGTGAGGTAATGGCTCACCAAGGCGACGATCCGTAACTGGTCTGAGAGGATGATCAGTCACACTGGAACTGAG
ACACGGTCCAGACTCCTACGGGAGGCAGCAGTGGGGAATATTGGACAATGGGCGAAAGCCTGATCCAGCCATGCCGCGTG
TGTGAAGAAGGTCTTCGGATTGTAAAGCACTTTAAGTTGGGAGGAAGGGTACTTACCTAATACGTGAGTATTTTGACGTT
ACCGACAGAATAAGCACCGGCTAACTCTGTGCCAGCAGCCGCGGTAATACAGAGGGTGCAAGCGTTAATCGGAATTACTG
GGCGTAAAGCGCGCGTAGGTGGTTCGTTAAGTTGGATGTGAAATCCCCGGGCTCAACCTGGGAACTGCATCCAAAACTGG
CGAGCTAGAGTATGGTAGAGGGTGGTGGAATTTCCTGTGTAGCGGTGAAATGCGTAGATATAGGAAGGAACACCAGTGGC
GAAGGCGACCACCTGGACTGATACTGACACTGAGGTGCGAAAGCGTGGGGAGCAAACAGGATTAGATACCCTGGTAGTCC
ACGCCGTAAACGATGTCAACTAGCCGTTGGGAGCCTTGAGCTCTTAGTGGCGCAGCTAACGCATTAAGTTGACCGCCTGG
GGAGTACGGCCGCAAGGTTAAAACTCAAATGAATTGACGGGGGCCCGCACAAGCGGTGGAGCATGTGGTTTAATTCGAAG
CAACGCGAAGAACCTTACCAGGCCTTGACATCCAATGAACTTTCCAGAGATGGATTGGTGCCTTCGGGAACATTGAGACA
GGTGCTGCATGGCTGTCGTCAGCTCGTGTCGTGAGATGTTGGGTTAAGTCCCGTAACGAGCGCAACCCTTGTCCTTAGTT
ACCAGCACGTCATGGTGGGCACTCTAAGGAGACTGCCGGTGACAAACCGGAGGAAGGTGGGGATGACGTCAAGTCATCAT
GGCCCTTACGGCCTGGGCTACACACGTGCTACAATGGTCGGTACAGAGGGTTGCCAAGCCGCGAGGTGGAGCTAATCCCA
TAAAACCGATCGTAGTCCGGATCGCAGTCTGCAACTCGACTGCGTGAAGTCGGAATCGCTAGTAATCGCGAATCAGAATG
TCGCGGTGAATACGTTCCCGGGCCTTGTACACACCGCCCGTCACACCATGGGAGTGGGTTGCACCAGAAGTAGCTAGTCT
AACCTTCGGGAGGACGGTTACCACGGTGTGATTCATGACTGGGGTGAAGTCGTAACAAGGTAGCCGTAGGGGAACCTGCG
GCTGGATCACCTCCTTAATATGGACGACAACAAGAAGAAAGCCTTGGCTGCGGCCCTGGGTCAGATCGAACGTCAATTCG
GCAAGGGTGCCGTAATGCGTATGGGCGATCACGACCGCCAGGCGATCCCGGCCATTTCCACTGGCTCTCTGGGTCTGGAC
ATCGCACTCGGCATCGGCGGCCTGCCAAAAGGTCGTATTGTTGAAATCTACGGTCCGGAATCGTCCGGTAAAACCACCCT
GACCCTGTCCGTGATTGCCCAGGCACAGAAGATGGGCGCCACCTGCGCCTTCGTCGACGCCGAGCACGCACTGGACCCGG
AATACGCCGGCAAACTGGGGGTCAACGTTGACGACCTGCTGGTTTCCCAGCCGGACACCGGCGAACAGGCGCTGGAAATC
ACCGACATGCTGGTGCGCTCCAATGCCATCGACGTGATCGTGATCGACTCCGTGGCGGCACTGGTACCCAAGGCCGAGAT
CGAAGGCGAGATGGGCGACATGCACGTGGGCCTGCAGGCCCGCCTGATGTCCCAGGCGCTGCGCAAGATCACCGGTAACA
TCAAGAACGCCAACTGCCTGGTGATCTTCATCAACCAGATCCGTATGAAAATCGGCGTGATGTTCGGCAGCCCGGAAACC
ACCACCGGCGGTAACGCGCTGAAGTTCTACGCTTCGGTTCGTCTGGACATCCGTCGTACTGGCGCGGTGAAGGAAGGCGA
CGAAGTCGTCGGTAGCGAAACCCGGGTCAAGATCGTCAAGAACAAGGTGGCTCCACCGTTCCGCCAGGCTGAATTCCAGA
TCCTGTACGGCAAGGGTATCTACCTGAACGGCGAGATCATCGATCTGGGCGTGCTGCACGGTTTCCTCGAGAAGTCCGGT
GCCTGGTACAGCTACCAGGGCAACAAGATCGGTCAGGGCAAGGCCAACTCGGCCAAGTTCCTGCAGGACAATCCGGAAAT
CGGCAATGCCCTCGAGAAGCAGATTCGCGACAAGCTGCTGGCTCCAAGCGCTGATGTCAAAGCTTCGCCGGTCAACGAGA
CCATCGATGACATGGCTGACGCGGATATCTGAATGAGCGAAGAAAACACGTACGACTCGAGCAGCATTAAAGTGCTGAAA
GGTTTGGATGCCGTACGCAAACGTCCCGGTATGTACATTGGTGACACCGACGATGGCAGCGGTCTGCACCATATGGTGTT
CGAGGTGGTCGATAACTCGATCGACGAAGCTCTGGCCGGCCATTGCGACGACATCAGCATCATCATCCACCCGGACGAAT
CCATTACCGTGCGTGACAACGGTCGCGGCATCCCGGTAGACGTGCATAAAGAAGAAGGCGTTTCCGCGGCCGAGGTCATC
ATGACCGTACTGCACGCCGGCGGTAAGTTCGACGATAACTCCTACAAAGTATCCGGCGGTCTGCACGGTGTGGGTGTGTC
GGTAGTGAACGCCCTGTCCGAAGAACTGGTCCTGACCGTTCGCCGCAGCGGAAAGATCTGGGAACAGACCTACGTTCACG
GTGTGCCTCAGGCGCCTATGGCGATCGTCGGTGACAGCGAAACCACCGGTACCCAGATTCACTTCAAGGCGTCCAGCGAG
ACCTTCAAGAACATCCATTTCAGCTGGGACATCCTGGCCAAGCGGATTCGTGAACTGTCCTTCCTCAACTCCGGTGTCGG
TATCGTTCTGAAGGACGAACGCAGTGGCAAGGAAGAGCTGTTCAAGTACGAAGGCGGCCTGCGTGCGTTCGTTGAATACC
TGAACACCAACAAGACCGCGGTCAACCAGGTGTTCCACTTCAATGTGCAGCGTGAAGATGGCATCGGCGTGGAAATCGCC
CTGCAGTGGAACGACAGCTTCAACGAAAACCTGCAGTGCTTCACCAACAACATTCCGCAGCGCGATGGCGGCACCCACTT
GGTGGGCTTCCGTTCGGCACTGACGCGTAACCTGAACAACTACATCGAACAGGAAGGTCTGGCGAAGAAGCACAAGGTCG
CCACCACCGGTGACGATGCCCGCGAAGGCCTGACCGCGATCATTTCGGTCAAGGTGCCGGATCCGAAGTTCAGCTCCCAG
ACCAAAGACAAGCTGGTGTCTTCCGAAGTGAAGACCGCGGTTGAACAGGAAATGGGCAAGTACTTCTCCGACTTCCTGCT
GGAAAACCCGAACGAAGCCAAGCTGGTGGTCGGCAAGATGCTCGACGCCGCCCGTGCCCGTGAAGCGGCGCGTAAGGCTC
GTGAGATGACCCGCCGTAAAGGCGCGCTGGATATCGCCGGCCTGCCGGGCAAACTGGCGGACTGCCAGGAAAAAGACCCT
GCCCTTTCCGAACTCTACTTGGTGGAAGGTGACTCTGCTGGCGGCTCCGCCAAGCAGGGACGCAACCGTAAGACCCAGGC
GATTCTGCCGCTCAAGGGCAAGATCCTTAACGTCGAGAAAGCGCGCTTCGACAAGATGATTTCCTCGCAAGAGGTCGGCA
CCTTGATCACTGCACTCGGTTGCGGCATCGGCCGCGAAGAGTACAACATCGACAAGCTGCGTTATCACAACATCATCATC
ATGACCGACGCTGACGTCGACGGTTCGCACATCCGTACCCTGCTGCTGACCTTCTTCTTCCGTCAGCTGCCGGAGCTGAT
CGAGCGTGGCTACATCTACATCGCTCAACCGCCGCTGTACAAGGTCAAGAAAGGCAAGCAAGAGCAATACATCAAAGACG
ACGACGCCATGGAAGAGTACATGACGCAGTCGGCCCTGGAAGATGCGAGCCTGCACCTGAACGAAGAAGCACCGGGTATT
TCCGGCGAGGCGCTGGAGCGCCTGGTGAACGACTTCCGCATGGTCATGAAAACCCTCAAGCGTCTGTCGCGCCTGTACCC
TCAGGAGCTGACCGAGCACTTCATCTACCTGCCGGCCGTGAGCCTGGAGCAACTCTCCGATCACGCGGCCATGCAGGATT
GGCTGGCCCAATATGAAGTCCGCCTGCGCACCGTCGAGAAGTCCGGCCTGGTCTACAAGGCCAGCCTGCGTGAAGACCGT
GAACGTAATGTCTGGCTGCCAGAGGTCGAACTGATCTCCCACGGCCTGTCGAACTACGTCACCTTCAACCGCGACTTCTT
CGGTAGCAATGACTACAAGACCGTCGTTACCCTCGGCGCTCAACTGAGCTCCCTGCTGGACGAAGGCGCTTATATTCAGC
GTGGCGAACGCAAGAAGGCGGTGACCGAGTTCAAGGAAGCCCTGGACTGGCTGATGACCGAAAGCACCAAGCGCCACACC
ATCCAGCGATACAAAGGTCTGGGCGAGATGAACCCGGATCAGCTGTGGGAAACCACCATGGACCCAAGCGTGCGCCGTAT
GCTCAAGGTCACGATTGAAGATGCCATCGGCGCCGACCAGATCTTCAACACCCTGATGGGGGATGCGGTCGAGCCTCGTC
GCGACTTCATCGAAAGCAACGCCCTGGCGGTATCCAATCTGGACTTCTGAATGTCCGGAAAAGCGCAACAGCAGTCTCGC
CTCAAAGAGTTGATCAGCCGTGGTCGTGAGCAGGGTTACCTGACTTACGCGGAGGTCAACGACCACCTGCCGGAGGATAT
TTCAGATCCGGAACAGGTGGAAGACATCATCCGCATGATCAACGACATGGGGATCAACGTATTCGAGAGTGCTCCGGATG
CGGATGCCCTTTTGTTGGCCGAAGCCGATACCGACGAAGCAGCAGCTGAAGAAGCCGCCGCAGCGTTGGCGGCCGTGGAA
ACCGACATTGGTCGCACTACCGACCCCGTGCGTATGTACATGCGCGAAATGGGAACGGTAGAGCTGCTCACACGTGAAGG

CGAAATCGAAATCGCCAAGCGTATCGAAGAGGGCATCCGTGAAGTGATGGGCGCGATCGCGCACTTCCCTGGCACGGTTG
AGCACATCCTCTCCGAATACACTCGCGTCACCACCGAAGGTGGCCGCCTGTCCGACGTCCTGAGCGGTTACATCGACCCG
GACGACGGCATCGCGCCGCCTGCCGCCGAAGTACCACCGCCTGTCGATGCCAAGGCCGCAAAAGCGGACGACGACACCGA
CGACGATGACGCCGAAGCCAGTGACGACGAAGAAGAAGCCGAAAGCGGTCCGGATCCGGTCATCGCAGCCCAGCGCTTTG
GCGCCGTTGCCGACCAGATGGAAATCACCCGCAAGGCGCTGAAGAAGCACGGTCGCGAACACAAGCAAGCCCTGGCTGAA
ATGCTGGCCCTGGCTGAACTGTTCATGCCGATCAAACTGGTTCCGAAGCAATTCGAAGGCCTGGTTGAACGTGTTCGTAG
CGCCCTGGATCGCCTGCGTCAGCAAGAGCGCGCGATCATGCAGCTCTGTGTTCGTGATGCCCGCATGCCACGCGCCGACT
TCCTGCGCCAGTTCCCTGGCAATGAAGTGGACGAAAGCTGGTCCGACGCGCTGGCCAAAGGCAAGGCCAAGTACGCCGAA
GCCATCGGCCGCCTGCAGCCGGACATTATCCGTTGCCAGCAGAAGCTGACCGCGCTTGAGACCGAGACCGGCCTGACGAT
CGCCGAGATCAAGGACATCAACCGTCGCATGTCGATCGGCGAGGCCAAGGCCCGTCGCGCGAAGAAAGAGATGGTCGAAG
CCAACCTGCGTCTGGTGATCTCCATCGCCAAGAAGTACACCAACCGTGGCTTGCAATTCCTCGACCTGATCCAGGAAGGC
AACATCGGTTTGATGAAAGCGGTAGACAAGTTCGAATACCGTCGCGGCTACAAATTCTCGACTTATGCCACCTGGTGGAT
CCGTCAGGCGATCACTCGCTCGATCGCCGACCAGGCCCGCACCATCCGTATTCCGGTGCACATGATCGAGACGATCAACA
AGCTCAACCGTATTTCCCGGCAGATGTTGCAGGAAATGGGTCGCGAACCGACTCCGGAAGAGCTGGGCGAACGCATGGAA
ATGCCTGAGGACAAGATCCGCAAGGTATTGAAGATCGCTAAAGAGCCGATCTCCATGGAAACCCCGATCGGTGATGACGA
AGACTCCCATCTGGGTGACTTCATCGAAGACTCGACCATGCAGTCGCCAATCGATGTCGCCACCGTCGAGAGTCTTAAAG
AAGCGACTCGCGAAGTACTCTCCGGCCTCACTGCCCGTGAAGCCAAGGTACTGCGCATGCGCTTCGGCATCGACATGAAT
ACCGACCACACCCTCGAGGAAGTCGGTAAGCAGTTTGACGTTACCCGTGAGCGGATTCGTCAGATCGAAGCCAAGGCGCT
GCGCAAGCTGCGCCACCCGACGCGAAGCGAGCACCTGCGCTCCTTCCTCGACGAGTAATTGACTAAGCCAGCCATACTCG
CCCTTGCTGATGGCAGCATTTTTCGCGGCGAAGCCATTGGAGCCGACGGTCAAACCGTTGGTGAGGTGGTGTTTAACACC
GCAATGACCGGCTATCAGGAAATCCTTACCGATCCTTCCTACGCCCAACAGATCGTTACCCTGACTTACCCGCATATCGG
CAATACCGGCACCACGCCGGAAGACGCCGAGTCCGATCGTGTCTGGTCGGCCGGTCTGGTGATTCGCGACCTGCCTCTGG
TTGCGAGCAACTGGCGTAACACCCTGTCCCTGTCCGACTACCTGAAAGCCAACAATGTCGTGGCAATCGCCGGTATCGAC
ACCCGTCGCCTGACGCGCATCCTGCGCGAGAAAGGTGCGCAGAACGGCTGCATCATGGCCGGCGACAATATCTCCGACGA
AGCGGCGATTGCCGCTGCACGCGGCTTCCCGGGCCTGAAAGGCATGGATCTGGCGAAGGTCGTCAGCACCAAGGAAAGCT
ACGAGTGGCGCTCCAGTGTCTGGAACCTGAAGACCGACAGTCATCCGACCATCGAAGCTTCCGAGCTGCCTTACCACGTG
GTTGCCTACGACTACGGCGTCAAGCTGAACATCCTGCGCATGCTGGTCGAACGCGGTTGCCGCGTGACCGTGGTGCCTGC
GCAAACCCCGGCCAGCGAAGCGCTGGCGCTCAAGCCTGACGGTGTGTTCCTGTCCAACGGCCCTGGCGACCCCGAGCCTT
GCGATTACGCCATCCAGGCGATCAAGGACGTGCTGGAGACCGAGATTCCGGTCTTCGGTATCTGTCTGGGCCACCAACTG
CTGGCACTGGCCGCCGGCGCCAAGACAGTGAAGATGGGCCACGGCCACCACGGCGCCAACCACCCGGTCCAGGACCTGGA
CAGCGGTGTGGTGATGATCACCAGCCAGAACCACGGTTTTGCGGTGGACGAAGCCACCCTGCCGGGCAACGTGCGGGCGA
TCCACAAGTCGCTGTTCGACGGCACCCTGCAAGGCATCGAGCTGATCGACAAGAGCGCATTCAGCTTCCAGGGCCACCCT
GAAGCGAGCCCGGGCCCGAACGATGTGGCGCCGCTGTTCGATCGTTTCATCAACGAGATGGCCAAGCGACGCTGAATGAG
TAGCGGACGTATCGTTCAAATCATCGGCGCCGTTATCGACGTGGAATTTCCACGCGACAGCGTACCGAGCATCTACGACG
CCTTGAAGGTTCAAGGCGCCGAAACCACTCTGGAAGTTCAGCAGCAGCTGGGCGACGGCGTGGTACGTACCATTGCGATG
GGCTCCACCGAGGGCTTGAAGCGCGGTCTGGACGTCAACAACACTGGCGCAGCCATCTCCGTACCGGTCGGTAAAGCGAC
CCTGGGCCGGATCATGGACGTACTGGGCAACCCGATCGACGAAGCTGGCCCGATCGGCGAAGAAGAGCGTTGGGGCATTC
ACCGTCCTGCGCCGACCTTCGCTGAACAAGCTGGCGGCAACGACCTGCTGGAAACCGGCATCAAGGTTATCGACCTGGTT
TGCCCGTTCGCCAAGGGCGGTAAAGTCGGTCTGTTCGGTGGTGCCGGTGTGGGCAAAACCGTAAACATGATGGAACTGAT
CCGTAACATCGCCATCGAGCACAGCGGTTATTCCGTGTTCGCCGGTGTGGGTGAGCGTACTCGTGAGGGTAACGACTTCT
ACCACGAGATGAAGGACTCCAACGTTCTGGACAAAGTGGCACTGGTATACGGCCAGATGAACGAGCCGCCGGGAAACCGT
CTGCGCGTAGCTCTGACCGGCCTGACCATGGCCGAGAAGTTCCGTGACGAAGGTAACGACGTTCTGCTGTTCGTCGACAA
CATCTATCGTTACACCCTGGCCGGTACCGAAGTATCCGCACTGCTGGGCCGTATGCCTTCGGCAGTAGGTTACCAGCCGA
CCCTGGCTGAAGAAATGGGCGTTCTGCAAGAACGTATCACTTCGACCAAGCAAGGCTCGATCACCTCGATCCAAGCGGTA
TACGTACCTGCGGACGACTTGACCGACCCGTCGCCAGCGACCACCTTCGCCCACTTGGACGCCACCGTCGTTCTGTCCCG
TGACATCGCTTCCCTGGGTATCTACCCAGCGGTAGACCCACTGGACTCGACTTCCCGTCAGCTGGACCCGAACGTGATCG
GCACCGAGCACTACGAAACCGCTCGTGGCGTTCAGTACGTGCTGCAGCGCTACAAAGAGCTGAAGGACATCATTGCGATC
CTGGGTATGGACGAACTGTCCGAAGCCGACAAGCAACTGGTATCCCGCGCTCGTAAGATCCAGCGCTTCCTGTCGCAGCC
GTTCTTCGTGGCTGAAGTCTTCACTGGTTCTCCAGGCAAATACGTTTCCCTGAAAGACACCATCGCTGGCTTCAAAGGCA
TCCTCAACGGTGACTACGACCATCTGCCAGAACAAGCGTTCTACATGGTTGGTGGCATCGAAGAAGCGATCGAGAAAGCC
AAGAAACTGTAA

NCBI Reference Sequence: NZ_CP027720.1
Strain: DSM6698
Chained genes: 16S rRNA, recA, gyrB, rpoD, carA, atpD
>NZ_CP027720_DSM6698
GAACTGAAGAGTTTGATCATGGCTCAGATTGAACGCTGGCGGCAGGCCTAACACATGCAAGTCGAGCGGTAGAGAGAAGC
TTGCTTCTCTTGAGAGCGGCGGACGGGTGAGTAATGCCTAGGAATCTGCCTGGTAGTGGGGGATAACGTTCGGAAACGGA
CGCTAATACCGCATACGTCCTACGGGAGAAAGCAGGGGACCTTCGGGCCTTGCGCTATCAGATGAGCCTAGGTCGGATTA
GCTAGTTGGTGAGGTAATGGCTCACCAAGGCGACGATCCGTAACTGGTCTGAGAGGATGATCAGTCACACTGGAACTGAG
ACACGGTCCAGACTCCTACGGGAGGCAGCAGTGGGGAATATTGGACAATGGGCGAAAGCCTGATCCAGCCATGCCGCGTG
TGTGAAGAAGGTCTTCGGATTGTAAAGCACTTTAAGTTGGGAGGAAGGGTACTTACCTAATACGTGAGTATTTTGACGTT
ACCGACAGAATAAGCACCGGCTAACTCTGTGCCAGCAGCCGCGGTAATACAGAGGGTGCAAGCGTTAATCGGAATTACTG
GGCGTAAAGCGCGCGTAGGTGGTTCGTTAAGTTGGATGTGAAATCCCCGGGCTCAACCTGGGAACTGCATCCAAAACTGG

CGAGCTAGAGTATGGTAGAGGGTGGTGGAATTTCCTGTGTAGCGGTGAAATGCGTAGATATAGGAAGGAACACCAGTGGC
GAAGGCGACCACCTGGACTGATACTGACACTGAGGTGCGAAAGCGTGGGGAGCAAACAGGATTAGATACCCTGGTAGTCC
ACGCCGTAAACGATGTCAACTAGCCGTTGGGAGCCTTGAGCTCTTAGTGGCGCAGCTAACGCATTAAGTTGACCGCCTGG
GGAGTACGGCCGCAAGGTTAAAACTCAAATGAATTGACGGGGGCCCGCACAAGCGGTGGAGCATGTGGTTTAATTCGAAG
CAACGCGAAGAACCTTACCAGGCCTTGACATCCAATGAACTTTCCAGAGATGGATTGGTGCCTTCGGGAACATTGAGACA
GGTGCTGCATGGCTGTCGTCAGCTCGTGTCGTGAGATGTTGGGTTAAGTCCCGTAACGAGCGCAACCCTTGTCCTTAGTT
ACCAGCACGTCATGGTGGGCACTCTAAGGAGACTGCCGGTGACAAACCGGAGGAAGGTGGGGATGACGTCAAGTCATCAT
GGCCCTTACGGCCTGGGCTACACACGTGCTACAATGGTCGGTACAGAGGGTTGCCAAGCCGCGAGGTGGAGCTAATCCCA
TAAAACCGATCGTAGTCCGGATCGCAGTCTGCAACTCGACTGCGTGAAGTCGGAATCGCTAGTAATCGCGAATCAGAATG
TCGCGGTGAATACGTTCCCGGGCCTTGTACACACCGCCCGTCACACCATGGGAGTGGGTTGCACCAGAAGTAGCTAGTCT
AACCTTCGGGAGGACGGTTACCACGGTGTGATTCATGACTGGGGTGAAGTCGTAACAAGGTAGCCGTAGGGGAACCTGCG
GCTGGATCACCTCCTTAAATGGACGACAACAAGAAGAAAGCCTTGGCTGCGGCCCTGGGTCAGATCGAACGTCAATTCGG
CAAGGGTGCCGTAATGCGTATGGGCGATCACGACCGCCAGGCGATCCCGGCCATTTCCACTGGCTCTCTGGGTCTGGACA
TCGCACTCGGCATCGGCGGCCTGCCAAAAGGTCGTATTGTTGAAATCTACGGTCCGGAATCGTCCGGTAAAACCACCCTG
ACCCTGTCCGTGATTGCCCAGGCACAGAAGATGGGCGCCACCTGCGCCTTCGTCGACGCCGAGCACGCACTGGACCCGGA
ATACGCCGGCAAACTGGGGGTCAACGTTGACGACCTGCTGGTTTCCCAGCCGGACACCGGCGAACAGGCGCTGGAAATCA
CCGACATGCTGGTGCGCTCCAATGCCATCGACGTGATCGTGATCGACTCCGTGGCGGCACTGGTACCCAAGGCCGAGATC
GAAGGCGAGATGGGCGACATGCACGTGGGCCTGCAGGCCCGCCTGATGTCCCAGGCACTGCGCAAGATCACCGGTAACAT
CAAGAACGCCAACTGCCTGGTGATCTTCATCAACCAGATCCGTATGAAAATCGGCGTGATGTTCGGCAGCCCGGAAACCA
CCACCGGCGGTAACGCGCTGAAGTTCTACGCTTCGGTTCGTCTGGACATCCGTCGTACTGGCGCGGTGAAGGAAGGCGAC
GAAGTCGTCGGTAGCGAAACCCGGGTCAAGATCGTCAAGAACAAGGTGGCTCCACCGTTCCGCCAGGCTGAATTCCAGAT
CCTGTACGGCAAGGGTATCTACCTGAACGGCGAGATCATCGATCTGGGCGTGCTGCACGGTTTCCTCGAGAAGTCCGGTG
CCTGGTACAGCTACCAGGGCAACAAGATCGGTCAGGGCAAGGCCAACTCGGCCAAGTTCCTGCAGGACAATCCGGAAATC
GGCAATGCCCTCGAGAAGCAGATTCGCGACAAGCTGCTGGCTCCAACCGCTGATGTCAAAGCTTCGCCGGTCAACGAGAC
CATCGATGACATGGCTGACGCGGATATCTGAATGAGCGAAGAAAACACGTACGACTCGAGCAGCATTAAAGTGCTGAAAG
GTTTGGATGCCGTACGCAAACGTCCCGGTATGTACATTGGTGACACCGACGATGGCAGCGGTCTGCACCATATGGTGTTC
GAGGTGGTCGATAACTCGATCGACGAAGCTCTGGCCGGCCATTGCGACGACATCAGCATCATCATCCACCCGGACGAATC
CATTACCGTGCGTGACAACGGTCGCGGCATCCCGGTAGACGTGCATAAAGAAGAAGGCGTTTCCGCGGCCGAGGTCATCA
TGACCGTACTGCACGCCGGCGGTAAGTTCGACGATAACTCCTACAAAGTATCCGGCGGTCTGCACGGTGTGGGTGTGTCG
GTAGTGAACGCCCTGTCCGAAGAACTGGTCCTGACCGTTCGCCGCAGCGGAAAGATCTGGGAACAGACCTACGTTCACGG
TGTGCCTCAGGCGCCTATGGCGATCGTCGGTGACAGCGAAACCACCGGTACCCAGATTCACTTCAAGGCGTCCAGCGAGA
CCTTCAAGAACATCCATTTCAGCTGGGACATCCTGGCCAAGCGGATTCGTGAACTGTCCTTCCTCAACTCCGGTGTCGGT
ATCGTTCTGAAGGACGAACGCAGTGGCAAGGAAGAGCTGTTCAAGTACGAAGGCGGCCTGCGTGCGTTCGTTGAATACCT
GAACACCAACAAGACCGCGGTCAACCAGGTGTTCCACTTCAATGTGCAGCGTGAAGATGGCATCGGCGTGGAAATCGCCC
TGCAGTGGAACGACAGCTTCAACGAAAACCTGCAGTGCTTCACCAACAACATTCCGCAGCGCGATGGCGGCACCCACTTG
GTGGGCTTCCGTTCGGCACTGACGCGTAACCTGAACAACTACATCGAACAGGAAGGTCTGGCGAAGAAGCACAAGGTCGC
CACCACCGGTGACGATGCCCGCGAAGGCCTGACCGCGATCATTTCGGTCAAGGTGCCGGATCCGAAGTTCAGCTCCCAGA
CCAAAGACAAGCTGGTGTCTTCCGAAGTGAAGACCGCGGTTGAACAGGAAATGGGCAAGTACTTCTCCGACTTCCTGCTG
GAAAACCCGAACGAAGCCAAGCTGGTGGTCGGCAAGATGCTCGACGCCGCCCGTGCCCGTGAAGCGGCGCGTAAGGCTCG
TGAGATGACCCGCCGTAAAGGCGCGCTGGATATCGCCGGCCTGCCGGGCAAACTGGCGGACTGCCAGGAAAAAGACCCTG
CCCTTTCCGAACTCTACCTGGTGGAAGGTGACTCTGCTGGCGGCTCCGCCAAGCAGGGACGCAACCGTAAGACCCAGGCG
ATTCTGCCGCTCAAGGGCAAGATCCTTAACGTCGAGAAAGCGCGCTTCGACAAGATGATTTCCTCGCAAGAGGTCGGCAC
CTTGATCACTGCACTCGGTTGCGGCATCGGCCGCGAAGAGTACAACATCGACAAGCTGCGTTATCACAACATCATCATCA
TGACCGACGCTGACGTCGACGGTTCGCACATCCGTACCCTGCTGCTGACCTTCTTCTTCCGTCAGCTGCCGGAGCTGATC
GAGCGTGGCTACATCTACATCGCTCAACCGCCGCTGTACAAGGTCAAGAAAGGCAAGCAAGAGCAATACATCAAAGACGA
CGACGCCATGGAAGAGTACATGACGCAGTCGGCCCTGGAAGATGCGAGCCTGCACCTGAACGAAGAAGCACCGGGTATTT
CCGGCGAGGCGCTGGAGCGCCTGGTGAACGACTTCCGCATGGTCATGAAAACCCTCAAGCGTCTGTCGCGCCTGTACCCT
CAGGAGCTGACCGAGCACTTCATCTACCTGCCGGCCGTGAGCCTGGAGCAACTCTCCGATCACGCGGCCATGCAGGATTG
GCTGGCCCAATATGAAGTCCGCCTGCGCACCGTCGAGAAGTCCGGCCTGGTCTACAAGGCCAGCCTGCGTGAAGACCGTG
AACGTAATGTCTGGCTGCCAGAGGTCGAACTGATCTCCCACGGCCTGTCGAACTACGTCACCTTCAACCGCGACTTCTTC
GGTAGCAATGACTACAAGACCGTCGTTACCCTCGGCGCTCAACTGAGCTCCCTGCTGGACGAAGGCGCTTATATTCAGCG
TGGCGAACGCAAGAAGGCGGTGACCGAGTTCAAGGAAGCCCTGGACTGGCTGATGACCGAAAGCACCAAGCGCCACACCA
TCCAGCGATACAAAGGTCTGGGCGAGATGAACCCGGATCAGCTGTGGGAAACCACCATGGACCCAAGCGTGCGCCGTATG
CTCAAGGTCACGATTGAAGATGCCATCGGCGCCGACCAGATCTTCAACACCCTGATGGGGGATGCGGTCGAGCCTCGTCG
CGACTTCATCGAAAGCAACGCCCTGGCGGTATCCAATCTGGACTTCTGAATGTCCGGAAAAGCGCAACAGCAGTCTCGCC
TCAAAGAGTTGATCAGCCGTGGTCGTGAGCAGGGTTACCTGACTTACGCGGAGGTCAACGACCACCTGCCGGAGGATATT
TCAGATCCGGAACAGGTGGAAGACATCATCCGCATGATCAACGACATGGGGATCAACGTATTCGAGAGTGCTCCGGATGC
GGATGCCCTTTTGTTGGCCGAAGCCGATACCGACGAAGCAGCAGCTGAAGAAGCCGCCGCAGCGTTGGCGGCTGTGGAAA
CCGACATTGGTCGCACTACCGACCCCGTGCGTATGTACATGCGCGAAATGGGAACGGTAGAGCTGCTCACACGTGAAGGC
GAAATCGAAATCGCCAAGCGTATCGAAGAGGGCATCCGTGAAGTGATGGGCGCGATCGCGCACTTCCCTGGCACGGTTGA
GCACATCCTCTCCGAATACACTCGCGTCACCACCGAAGGTGGCCGCCTGTCCGACGTCCTGAGCGGTTACATCGACCCGG
ACGACGGCATTGCGCCGCCTGCCGCCGAAGTACCACCGCCTGTCGATGCCAAGGCTGCAAAAGCGGACGACGACACCGAC
GACGATGACGCCGAAGCCAGTGACGACGAAGAAGAAGCCGAAAGCGGTCCGGATCCGGTCATCGCAGCCCAGCGCTTTGG
CGCCGTTGCCGACCAGATGGAAATCACCCGCAAGGCGCTGAAGAAGCACGGTCGCGAACACAAGCAAGCCCTGGCCGAAA

TGCTGGCCCTGGCTGAACTGTTCATGCCGATCAAACTGGTTCCGAAGCAATTCGAAGGCCTGGTTGAACGTGTTCGTAGC
GCCCTGGATCGCCTGCGTCAGCAAGAGCGCGCGATCATGCAGCTCTGTGTTCGTGATGCCCGCATGCCACGCGCCGACTT
CCTGCGCCAGTTCCCTGGCAATGAAGTGGACGAAAGCTGGTCCGACGCGCTGGCCAAAGGCAAGGCCAAGTACGCCGAAG
CCATCGGCCGCCTGCAGCCGGACATCATCCGTTGCCAGCAGAAGCTGACCGCGCTCGAGACCGAGACCGGCCTGACGATC
GCCGAGATCAAGGACATCAACCGTCGCATGTCGATCGGCGAGGCCAAGGCCCGTCGCGCGAAGAAAGAGATGGTCGAAGC
CAACCTGCGTCTGGTGATCTCCATCGCCAAGAAGTACACCAACCGTGGCTTGCAATTCCTCGACCTGATCCAGGAAGGCA
ACATCGGTTTGATGAAAGCGGTAGACAAGTTCGAATACCGTCGCGGCTACAAATTCTCGACTTATGCCACCTGGTGGATC
CGTCAGGCGATCACTCGCTCGATCGCCGACCAGGCCCGCACCATCCGTATTCCGGTGCACATGATCGAGACGATCAACAA
GCTCAACCGTATTTCCCGGCAGATGTTGCAGGAAATGGGTCGCGAACCGACTCCGGAAGAGCTGGGCGAACGCATGGAAA
TGCCTGAGGACAAGATCCGCAAGGTATTGAAGATCGCTAAAGAGCCGATCTCCATGGAAACCCCGATCGGTGATGACGAA
GACTCCCATCTGGGTGACTTCATCGAAGACTCGACCATGCAGTCGCCAATCGATGTCGCCACCGTTGAGAGCCTTAAAGA
AGCGACTCGCGAAGTACTCTCCGGCCTCACTGCCCGTGAAGCCAAGGTACTGCGCATGCGCTTCGGCATCGACATGAATA
CCGACCACACCCTCGAGGAAGTCGGTAAGCAGTTCGACGTTACCCGTGAGCGGATTCGTCAGATCGAAGCCAAGGCGCTG
CGCAAGCTGCGCCACCCGACGCGAAGCGAGCACCTGCGCTCCTTCCTCGACGAGTAATTGACTAAGCCAGCCATACTCGC
CCTTGCTGATGGCAGCATTTTTCGCGGCGAAGCCATTGGAGCCGACGGTCAAACCGTTGGTGAGGTGGTGTTTAACACCG
CAATGACCGGCTATCAGGAAATCCTTACCGATCCTTCCTACGCCCAACAGATCGTTACCCTGACTTACCCGCATATCGGC
AATACCGGCACCACGCCGGAAGACGCCGAGTCCGATCGTGTCTGGTCGGCCGGTCTGGTGATTCGCGACCTGCCTCTGGT
TGCGAGCAACTGGCGTAACACCCTGTCCCTGTCCGACTACCTGAAAGCCAACAATGTCGTGGCGATCGCCGGTATCGACA
CCCGTCGCCTGACGCGCATCCTGCGCGAGAAAGGTGCGCAGAACGGCTGCATCATGGCCGGCGACAATATCTCCGACGAA
GCGGCGATTGCCGCTGCACGCGGCTTCCCGGGCCTGAAAGGCATGGATCTGGCGAAGGTCGTCAGTACCAAGGAAAGCTA
CGAGTGGCGCTCCAGTGTCTGGAACCTGAAGACCGACAGTCATCCGACCATCGAAGCTTCCGAGCTGCCTTACCACGTGG
TTGCCTACGACTACGGCGTCAAGCTGAACATCCTGCGCATGCTGGTCGAACGCGGTTGCCGCGTGACCGTGGTGCCTGCG
CAAACCCCGGCCAGCGAAGCTCTGGCGCTCAAGCCTGACGGTGTGTTCCTGTCCAACGGCCCTGGCGACCCCGAGCCTTG
CGATTACGCCATCCAGGCGATCAAGGACGTGCTGGAGACCGAGATTCCGGTCTTCGGTATCTGTCTGGGCCACCAACTGC
TGGCACTGGCCGCCGGCGCCAAGACAGTGAAGATGGGCCACGGCCACCACGGCGCCAACCACCCGGTCCAGGACCTGGAC
AGCGGTGTGGTGATGATCACCAGCCAGAACCACGGTTTTGCGGTGGACGAAGCCACCCTGCCGGGCAACGTGCGGGCGAT
CCACAAGTCGCTGTTCGACGGCACCCTGCAAGGCATCGAGCTGACCGACAAGAGCGCATTCAGCTTCCAGGGCCACCCTG
AAGCGAGCCCGGGCCCGAACGATGTGGCGCCGCTGTTCGATCGTTTCATCAACGAGATGGCCAAGCGACGCTGAATGAGT
AGCGGACGTATCGTTCAAATCATCGGCGCCGTTATCGACGTGGAATTTCCACGCGACAGCGTACCGAGCATCTACGACGC
CTTGAAGGTTCAAGGCGCCGAAACCACTCTGGAAGTTCAGCAGCAGCTGGGCGACGGCGTGGTACGTACCATTGCGATGG
GCTCCACCGAGGGCTTGAAGCGCGGTCTGGACGTCAACAACACTGGCGCAGCCATCTCCGTACCGGTCGGTAAAGCGACC
CTGGGCCGGATCATGGACGTACTGGGCAACCCGATCGACGAAGCTGGCCCGATCGGTGAAGAAGAGCGTTGGGGCATTCA
CCGTCCTGCGCCGACCTTCGCTGAACAAGCTGGCGGCAACGACCTGCTGGAAACCGGCATCAAGGTTATCGACCTGGTTT
GCCCGTTCGCCAAGGGCGGTAAAGTCGGTCTGTTCGGTGGTGCCGGTGTGGGCAAAACCGTAAACATGATGGAACTGATC
CGTAACATCGCCATCGAGCACAGCGGTTATTCCGTGTTCGCCGGTGTGGGTGAGCGTACTCGTGAGGGTAACGACTTCTA
CCACGAGATGAAGGATTCCAACGTTCTGGACAAAGTGGCACTGGTATACGGCCAGATGAACGAGCCGCCGGGAAACCGTC
TGCGCGTAGCTCTGACCGGCCTGACCATGGCCGAGAAGTTCCGTGACGAAGGTAACGACGTTCTGCTGTTCGTCGACAAC
ATCTATCGTTACACCCTGGCCGGTACCGAAGTATCCGCACTGCTGGGCCGTATGCCTTCGGCAGTAGGTTACCAGCCGAC
CCTGGCTGAAGAGATGGGCGTTCTGCAAGAACGTATCACTTCGACCAAGCAAGGCTCGATCACCTCGATCCAAGCGGTAT
ACGTACCTGCGGACGACTTGACCGACCCGTCGCCAGCGACCACCTTCGCCCACTTGGACGCCACCGTCGTTCTGTCCCGT
GACATCGCTTCCCTGGGTATCTACCCAGCGGTAGACCCACTGGACTCGACTTCCCGTCAGCTGGACCCGAACGTGATCGG
CACCGAGCACTACGAAACCGCTCGTGGCGTTCAGTACGTGCTGCAGCGCTACAAAGAGCTGAAGGACATCATTGCGATCC
TGGGTATGGACGAACTGTCCGAAGCCGACAAGCAACTGGTATCCCGCGCTCGTAAGATCCAGCGCTTCCTGTCGCAGCCG
TTCTTCGTGGCTGAAGTCTTCACTGGTTCTCCAGGCAAATACGTTTCCCTGAAAGACACCATCGCTGGCTTCAAAGGCAT
CCTCAACGGTGACTACGACCATCTGCCAGAACAAGCGTTCTACATGGTTGGTGGCATCGAAGAAGCGATCGAGAAAGCCA
AGAAACTGTAA

NCBI Reference Sequence: NZ_CP027737.1
Strain: PCL1607
Chained genes: 16S rRNA, recA, gyrB, rpoD, carA, atpD
>NZ_CP027737_PCL1607
GAACTGAAGAGTTTGATCATGGCTCAGATTGAACGCTGGCGGCAGGCCTAACACATGCAAGTCGAGCGGTAGAGAGGTGC
TTGCACCTCTTGAGAGCGGCGGACGGGTGAGTAATGCCTAGGAATCTGCCTGGTAGTGGGGGATAACGTTCGGAAACGGA
CGCTAATACCGCATACGTCCTACGGGAGAAAGCAGGGGACCTTCGGGCCTTGCGCTATCAGATGAGCCTAGGTCGGATTA
GCTAGTTGGTGAGGTAATGGCTCACCAAGGCGACGATCCGTAACTGGTCTGAGAGGATGATCAGTCACACTGGAACTGAG
ACACGGTCCAGACTCCTACGGGAGGCAGCAGTGGGGAATATTGGACAATGGGCGAAAGCCTGATCCAGCCATGCCGCGTG
TGTGAAGAAGGTCTTCGGATTGTAAAGCACTTTAAGTTGGGAGGAAGGGTACTTACCTAATACGTGAGTATTTTGACGTT
ACCGACAGAATAAGCACCGGCTAACTCTGTGCCAGCAGCCGCGGTAATACAGAGGGTGCAAGCGTTAATCGGAATTACTG
GGCGTAAAGCGCGCGTAGGTGGTTCGTTAAGTTGAATGTGAAATCCCCGGGCTCAACCTGGGAACTGCATCCAAAACTGG
CGAGCTAGAGTATGGTAGAGGGTGGTGGAATTTCCTGTGTAGCGGTGAAATGCGTAGATATAGGAAGGAACACCAGTGGC
GAAGGCGACCACCTGGACTGATACTGACACTGAGGTGCGAAAGCGTGGGGAGCAAACAGGATTAGATACCCTGGTAGTCC
ACGCCGTAAACGATGTCAACTAGCCGTTGGGAGCCTTGAGCTCTTAGTGGCGCAGCTAACGCATTAAGTTGACCGCCTGG
GGAGTACGGCCGCAAGGTTAAAACTCAAATGAATTGACGGGGGCCCGCACAAGCGGTGGAGCATGTGGTTTAATTCGAAG
CAACGCGAAGAACCTTACCAGGCCTTGACATCCAATGAACTTTCCAGAGATGGATTGGTGCCTTCGGGAACATTGAGACA

GGTGCTGCATGGCTGTCGTCAGCTCGTGTCGTGAGATGTTGGGTTAAGTCCCGTAACGAGCGCAACCCTTGTCCTTAGTT
ACCAGCACGTAATGGTGGGCACTCTAAGGAGACTGCCGGTGACAAACCGGAGGAAGGTGGGGATGACGTCAAGTCATCAT
GGCCCTTACGGCCTGGGCTACACACGTGCTACAATGGTCGGTACAGAGGGTTGCCAAGCCGCGAGGTGGAGCTAATCCCA
TAAAACCGATCGTAGTCCGGATCGCAGTCTGCAACTCGACTGCGTGAAGTCGGAATCGCTAGTAATCGCGAATCAGAATG
TCGCGGTGAATACGTTCCCGGGCCTTGTACACACCGCCCGTCACACCATGGGAGTGGGTTGCACCAGAAGTAGCTAGTCT
AACCTTCGGGAGGACGGTTACCACGGTGTGATTCATGACTGGGGTGAAGTCGTAACAAGGTAGCCGTAGGGGAACCTGCG
GCTGGATCACCTCCTTAAATGGACGACAACAAGAAGAAAGCCTTGGCTGCGGCCCTGGGTCAGATCGAACGTCAATTCGG
CAAGGGTGCCGTAATGCGTATGGGCGATCACGATCGCCAGGCGATCCCGGCCATTTCCACTGGCTCTCTGGGTCTGGACA
TCGCACTCGGCATCGGCGGCCTGCCAAAAGGCCGTATTGTTGAAATCTACGGTCCGGAATCGTCCGGTAAAACCACCCTG
ACCCTGTCGGTGATTGCCCAGGCACAGAAGATGGGCGCCACCTGCGCCTTCGTCGACGCCGAGCACGCACTGGACCCGGA
GTACGCCGGCAAACTGGGGGTCAACGTTGACGACCTGCTGGTTTCCCAGCCGGACACCGGCGAACAGGCGCTGGAAATCA
CCGACATGCTGGTGCGCTCCAATGCCATCGACGTGATCGTGATCGACTCCGTGGCGGCGCTGGTACCCAAGGCCGAGATC
GAAGGCGAGATGGGCGACATGCACGTGGGCCTGCAGGCTCGCCTGATGTCCCAGGCGCTGCGCAAGATCACCGGTAACAT
CAAGAACGCCAACTGCCTGGTGATCTTCATCAACCAGATCCGTATGAAAATCGGCGTGATGTTCGGCAGCCCGGAAACCA
CCACCGGTGGTAACGCGCTGAAGTTCTACGCTTCGGTTCGTCTGGATATCCGTCGTACTGGCGCGGTGAAGGAAGGTGAC
GAAGTCGTCGGTAGCGAAACCCGGGTCAAGATCGTCAAGAACAAGGTGGCTCCACCCTTCCGTCAAGCTGAGTTCCAGAT
CCTGTACGGCAAGGGTATCTACCTGAACGGCGAGATCATCGATCTGGGCGTGCTGCACGGTTTCCTAGAGAAGTCCGGTG
CCTGGTACAGCTACCAGGGCAACAAGATCGGTCAGGGCAAGGCCAACTCGGCCAAGTTCCTGCAGGACAATCCGGAAATC
GGTAATGCCCTCGAGAAGCAGATTCGCGACAAGCTGCTGGCTCCGAGCGGAGATACCAAGGCTCTGCCCGTCAACGAGAC
CATCGATGACATGGCCGACGCGGATATCTGAATGAGCGAAGAAAACACGTACGACTCGAGCAGCATTAAAGTGCTGAAAG
GTTTGGATGCCGTACGCAAACGTCCCGGTATGTACATTGGTGACACCGACGATGGCAGCGGTCTGCACCATATGGTGTTC
GAGGTGGTCGACAACTCGATCGACGAAGCTCTGGCCGGCCACTGCGACGACATCAGCATCATCATCCACCCGGACGAATC
CATTACCGTGCGTGACAACGGTCGCGGCATCCCGGTAGACGTGCATAAAGAAGAAGGCGTTTCCGCAGCCGAGGTCATCA
TGACCGTGCTGCACGCCGGCGGTAAGTTCGACGACAACTCCTACAAGGTATCCGGCGGTCTGCACGGTGTGGGTGTGTCG
GTAGTGAATGCCCTGTCCGAAGAACTGGTGCTGACCGTTCGCCGCAGTGGCAAGATCTGGGAACAGACCTACGTTCACGG
TGTGCCTCAGGCGCCTATGGCGATCGTCGGTGACAGCGAGACCACTGGTACCCAGATTCACTTCAAGGCTTCCAGCGAGA
CCTTCAAGAACATCCATTTCAGCTGGGACATCCTGGCCAAGCGGATTCGTGAACTGTCCTTCCTCAACTCCGGTGTCGGT
ATCGTTCTGAAGGACGAGCGCAGCGGCAAGGAAGAACTGTTCAAATACGAAGGCGGCCTGCGCGCGTTCGTCGAATACCT
GAACACCAACAAGACTGCGGTCAACCAGGTGTTCCACTTCAACGTGCAGCGTGAAGACGGCATCGGCGTGGAAATCGCCC
TGCAGTGGAACGACAGCTTCAACGAAAACCTGCAGTGCTTCACCAACAACATTCCGCAGCGCGACGGCGGTACTCACCTG
GTGGGCTTCCGCTCGGCACTGACGCGTAACCTGAACAACTACATCGAGCAGGAAGGTCTGGCGAAGAAGCACAAGGTCGC
CACCACCGGTGACGATGCCCGCGAAGGCCTGACCGCGATCATTTCGGTCAAGGTGCCGGATCCGAAGTTCAGCTCCCAGA
CCAAAGACAAGCTGGTGTCTTCCGAAGTAAAGACCGCGGTCGAACAGGAAATGGGCAAGTACTTCTCCGACTTCCTGCTG
GAAAACCCGAACGAAGCCAAGCTGGTGGTCGGCAAGATGCTCGACGCCGCCCGCGCCCGTGAAGCGGCGCGTAAGGCCCG
TGAGATGACCCGCCGTAAAGGTGCGCTGGATATCGCCGGCCTGCCGGGCAAACTGGCGGACTGCCAGGAAAAGGACCCTG
CCCTTTCCGAACTCTACCTGGTGGAAGGTGACTCTGCTGGCGGCTCCGCCAAGCAGGGACGCAACCGCAAGACCCAGGCG
ATTCTGCCGCTCAAGGGCAAGATTCTTAACGTCGAGAAAGCGCGCTTCGACAAGATGATTTCCTCGCAAGAGGTCGGCAC
CTTGATCACTGCACTCGGCTGCGGCATCGGCCGCGAAGAGTACAACATCGACAAGCTGCGTTATCACAACATCATCATCA
TGACCGACGCCGACGTCGACGGTTCGCACATCCGCACCCTGCTGCTGACTTTCTTCTTCCGTCAGCTGCCGGAGCTGATC
GAGCGTGGCTACATCTACATCGCTCAACCGCCGCTGTACAAGGTCAAGAAAGGCAAGCAAGAGCAATACATCAAAGACGA
CGACGCCATGGAAGAGTACATGACGCAGTCGGCCCTGGAAGATGCGAGCCTGCACCTGAACGAAGAAGCACCGGGTATTT
CCGGCGAGGCGCTGGAGCGCCTGGTGAACGACTTCCGCATGGTCATGAAAACCCTCAAGCGTCTGTCGCGCCTGTACCCT
CAGGAGCTGACCGAACACTTCATCTACCTGCCAGCCGTGAGCCTGGAGCAACTCTCCGATCACGCAGCGATGCAGGATTG
GTTGGCCCAATATGAAGTCCGCCTGCGCACCGTCGAGAAGTCCGGCCTGGTCTACAAGGCCAGCCTGCGTGAAGACCGTG
AACGTAATGTCTGGCTGCCAGAGGTCGAACTGATCTCCCACGGCCTGTCGAACTACGTCACCTTCAACCGCGACTTCTTC
GGCAGCAATGACTACAAGACCGTCGTCACCCTCGGCGCTCAACTGAGCTCCCTGCTGGACGAAGGCGCTTATATTCAGCG
CGGCGAACGCAAGAAGGCGGTGACCGAGTTCAAGGAAGCCCTGGACTGGCTGATGACCGAAAGCACCAAGCGCCACACCA
TCCAGCGATACAAAGGTCTGGGCGAGATGAACCCGGACCAGCTGTGGGAAACCACCATGGACCCAAGCGTGCGCCGCATG
CTCAAGGTCACCATCGAAGACGCCATCGGCGCCGACCAGATCTTCAACACCCTGATGGGTGATGCGGTCGAGCCTCGTCG
CGACTTCATCGAAAGCAACGCCCTGGCGGTATCCAACCTGGACTTCTGAATGTCCGGAAAAGCGCAACAGCAGTCTCGCC
TCAAAGAGTTGATCAGCCGTGGTCGTGAGCAGGGTTACCTGACTTACGCGGAGGTCAACGACCACCTGCCGGAGGATATT
TCAGATCCGGAACAGGTGGAAGACATCATCCGCATGATCAACGACATGGGGATCAACGTATTCGAGAGTGCTCCGGATGC
GGATGCCCTTTTGTTGGCCGAAGCCGATACCGACGAAGCAGCAGCTGAAGAAGCCGCCGCAGCGTTGGCGGCTGTGGAAA
CCGACATTGGTCGCACTACCGACCCAGTGCGTATGTACATGCGCGAAATGGGCACGGTAGAGCTGCTCACACGTGAAGGC
GAAATCGAAATCGCCAAGCGTATCGAAGAGGGCATCCGTGAAGTGATGGGCGCGATCGCGCACTTCCCTGGCACGGTTGA
GCACATCCTCTCCGAATACACTCGCGTCACCACCGAAGGTGGCCGCCTGTCCGACGTCCTGAGCGGTTACATCGACCCGG
ATGACGGCATTGCGCCGCCTGCCGCCGAAGTACCACCGCCTGTCGATGCCAAGGCCGCGAAAGCGGACGACGACACCGAC
GACGATGACGCCGAAGCCAGTGACGACGAAGAAGAAGCCGAAAGCGGTCCGGATCCGGTCATCGCAGCCCAGCGCTTTGG
CGCCGTTGCCGACCAGATGGAAATCACCCGCAAGGCGCTGAAAAAGCACGGTCGTGAACACAAGCAAGCCCTGGCTGAAA
TGCTGGCCCTGGCTGAACTGTTCATGCCAATCAAACTGGTTCCGAAGCAATTCGAAGGCCTGGTTGAACGTGTTCGTAGC
GCCCTGGATCGCCTGCGTCAGCAAGAGCGCGCGATCATGCAGCTCTGTGTTCGTGATGCCCGCATGCCACGCGCCGACTT
CCTGCGCCAGTTCCCTGGCAATGAAGTGGACGAAAGCTGGTCCGACGCGCTGGCCAAAGGCAAGGCCAAGTACGCCGAAG
CCATCGGCCGCCTGCAGCCGGACATCATCCGTTGCCAGCAGAAGCTGACCGCGCTCGAGACCGAGACCGGCCTGACGATT
GCCGAGATCAAGGACATCAACCGTCGCATGTCGATCGGCGAGGCCAAGGCCCGTCGCGCGAAGAAAGAGATGGTCGAAGC

CAACTTGCGCCTGGTGATCTCCATCGCCAAGAAGTACACCAACCGTGGCCTGCAGTTCCTCGACCTGATCCAGGAAGGCA
ACATCGGTTTGATGAAAGCGGTAGACAAGTTCGAATACCGTCGCGGCTACAAGTTCTCGACTTATGCCACCTGGTGGATC
CGTCAGGCGATCACTCGCTCGATCGCCGACCAGGCCCGCACCATCCGTATTCCGGTGCACATGATCGAGACGATCAACAA
GCTCAACCGTATTTCCCGGCAGATGTTGCAGGAAATGGGTCGCGAACCGACTCCGGAAGAGCTGGGCGAACGCATGGAAA
TGCCTGAGGACAAGATCCGCAAGGTATTGAAGATCGCTAAAGAGCCGATCTCCATGGAAACCCCGATCGGTGATGACGAA
GACTCCCATCTGGGTGACTTCATCGAAGACTCGACCATGCAGTCGCCAATCGATGTCGCCACCGTTGAGAGCCTCAAGGA
AGCGACTCGCGAAGTCCTCTCCGGCCTCACTGCCCGTGAAGCCAAGGTACTGCGCATGCGCTTCGGCATCGACATGAATA
CCGATCACACCCTTGAGGAAGTCGGTAAGCAGTTCGACGTTACCCGTGAGCGGATTCGTCAGATCGAAGCCAAGGCGCTG
CGCAAGCTGCGCCACCCGACGAGAAGCGAGCATCTGCGCTCCTTCCTCGACGAGTAATTGACTAAGCCAGCCATACTCGC
CCTTGCTGATGGCAGCATTTTTCGCGGCGAAGCCATTGGAGCCGACGGTCAAACCGTTGGTGAGGTGGTGTTTAACACCG
CAATGACCGGCTATCAGGAAATCCTTACCGATCCTTCCTACGCCCAACAGATCGTTACCCTGACTTACCCGCACATCGGC
AACACTGGCACCACGCCGGAAGACGCCGAGTCCGATCGTGTCTGGTCGGCCGGTCTGGTGATTCGCGACCTGCCACTGGT
TGCGAGCAACTGGCGTAACACCCTGTCCCTGTCCGACTACCTGAAAGCCAACAATGTCGTGGCGATCGCCGGTATCGACA
CCCGTCGCCTGACGCGCATCCTGCGTGAAAAAGGCGCGCAGAACGGCTGCATCATGGCCGGCGACAATATCTCCGACGAA
GCAGCGATTGCCGCAGCGCGCGGCTTCCCTGGCCTGAAAGGCATGGATCTGGCGAAGGTCGTCAGCACCAAGGAAAGCTA
CGAGTGGCGCTCCAGCGTCTGGAGCCTGAAGACCGACAGTCATCCGACTATCGAGGCTTCCGAGCTGCCTTACCACGTGG
TTGCCTACGACTACGGCGTCAAGCTGAACATCCTGCGCATGCTGGTCGAGCGCGGTTGCCGCGTGACCGTAGTGCCTGCG
CAAACCCCGGCCAGCGACGTCCTGGCACTCAAGCCTGACGGTGTGTTCCTGTCCAACGGTCCTGGTGACCCCGAGCCTTG
CGATTACGCCATCCAGGCGATCAAGGACGTGCTGGAAACCGAGATTCCGGTCTTCGGTATCTGCCTGGGCCACCAACTGC
TGGCACTGGCCTCCGGCGCCAAGACGGTGAAGATGGGCCACGGTCACCACGGTGCCAACCACCCGGTCCAGGACCTGGAC
AGCGGTGTAGTGATGATCACTAGCCAGAACCACGGTTTTGCGGTGGACGAAGCCACCCTGCCGGGCAACGTGCGGGCGAT
CCACAAGTCGCTGTTCGACGGCACCCTGCAAGGCATCGAGCGTACCGACAAGAGTGCATTCAGCTTCCAGGGCCACCCTG
AAGCGAGCCCGGGCCCGAACGATGTGGCGCCGCTGTTCGATCGTTTCATCAACGAGATGGTCAAGCGACGCTGAATGAGT
AGCGGACGTATCGTTCAAATCATCGGCGCCGTTATCGACGTGGAATTTCCACGCGACAGCGTACCGAGCATCTACGACGC
CTTGAAGGTTCAAGGCGCCGAAACCACTCTGGAAGTTCAGCAGCAGCTGGGCGACGGCGTGGTACGTACCATTGCGATGG
GCTCCACCGAGGGCTTGAAGCGCGGTCTGGACGTCAACAACACTGGCGCAGCCATCTCCGTACCGGTCGGTAAAGCGACC
CTGGGCCGGATCATGGACGTACTGGGCAACCCGATCGACGAAGCTGGCCCGATCGGCGAAGAAGAGCGTTGGGGCATTCA
CCGTCCTGCGCCGACCTTCGCTGAACAAGCTGGCGGCAACGACCTCCTGGAAACCGGCATCAAGGTTATCGACCTGGTTT
GCCCGTTCGCCAAGGGCGGTAAAGTCGGTCTGTTCGGTGGTGCCGGTGTGGGCAAAACCGTAAACATGATGGAACTGATC
CGTAACATCGCCATCGAGCACAGCGGTTATTCCGTGTTCGCCGGTGTGGGTGAGCGTACTCGTGAGGGTAACGACTTCTA
CCACGAGATGAAGGATTCCAACGTTCTGGACAAAGTGGCACTGGTATACGGCCAGATGAACGAGCCGCCGGGAAACCGTC
TGCGCGTAGCTCTGACCGGCCTGACCATGGCCGAGAAGTTCCGTGACGAAGGTAACGACGTTCTGCTGTTCGTCGACAAC
ATCTATCGTTACACCCTGGCCGGTACCGAAGTATCCGCACTGCTGGGCCGTATGCCTTCGGCAGTAGGTTACCAGCCGAC
CCTGGCTGAAGAGATGGGCGTTCTGCAAGAACGTATCACTTCGACCAAGCAAGGCTCGATCACCTCGATCCAAGCGGTAT
ACGTACCTGCGGACGACTTGACCGACCCGTCGCCAGCGACCACCTTCGCCCACTTGGACGCCACCGTCGTACTGTCCCGT
GACATCGCTTCCCTGGGTATCTACCCAGCGGTAGACCCACTGGATTCGACTTCCCGTCAGCTGGACCCGAACGTGATCGG
CAACGAGCACTACGAAACTGCTCGCGGCGTTCAGTACGTGCTGCAGCGCTACAAAGAGCTGAAGGACATCATTGCGATCC
TGGGTATGGACGAACTGTCCGAAGCCGACAAGCAACTGGTATCCCGCGCTCGTAAGATCCAGCGCTTCCTGTCGCAGCCG
TTCTTCGTGGCTGAAGTCTTCACTGGTTCTCCAGGCAAATACGTTTCCCTGAAAGACACCATCGCTGGCTTCAAAGGCAT
CCTCAACGGTGACTACGACCACCTGCCAGAACAAGCGTTCTACATGGTCGGCGGCATCGAAGAAGCGATCGAGAAAGCCA
AGAAACTGTAA

RAST server Genome ID: 286.2086
Strain: ST9
Chained genes: 16S rRNA, recA, gyrB, rpoD, carA, atpD
>ID_286.2086_ST9
GAACTGAAGAGTTTGATCATGGCTCAGATTGAACGCTGGCGGCAGGCCTAACACATGCAAGTCGAGCGGTAGAGAGAAGC
TTGCTTCTCTTGAGAGCGGCGGACGGGTGAGTAATGCCTAGGAATCTGCCTGGTAGTGGGGGATAACGTCCGGAAACGGA
CGCTAATACCGCATACGTCCTACGGGAGAAAGCAGGGGACCTTCGGGCCTTGCGCTATCAGATGAGCCTAGGTCGGATTA
GCTAGTTGGTGAGGTAATGGCTCACCAAGGCGACGATCCGTAACTGGTCTGAGAGGATGATCAGTCACACTGGAACTGAG
ACACGGTCCAGACTCCTACGGGAGGCAGCAGTGGGGAATATTGGACAATGGGCGAAAGCCTGATCCAGCCATGCCGCGTG
TGTGAAGAAGGTCTTCGGATTGTAAAGCACTTTAAGTTGGGAGGAAGGGTACTTACCTAATACGTGAGTATTTTGACGTT
ACCGACAGAATAAGCACCGGCTAACTCTGTGCCAGCAGCCGCGGTAATACAGAGGGTGCAAGCGTTAATCGGAATTACTG
GGCGTAAAGCGCGCGTAGGTGGTTCGTTAAGTTGGATGTGAAATCCCCGGGCTCAACCTGGGAACTGCATCCAAAACTGG
CGAGCTAGAGTATGGTAGAGGGTGGTGGAATTTCCTGTGTAGCGGTGAAATGCGTAGATATAGGAAGGAACACCAGTGGC
GAAGGCGACCACCTGGACTGATACTGACACTGAGGTGCGAAAGCGTGGGGAGCAAACAGGATTAGATACCCTGGTAGTCC
ACGCCGTAAACGATGTCAACTAGCCGTTGGGAGCCTTGAGCTCTTAGTGGCGCAGCTAACGCATTAAGTTGACCGCCTGG
GGAGTACGGCCGCAAGGTTAAAACTCAAATGAATTGACGGGGGCCCGCACAAGCGGTGGAGCATGTGGTTTAATTCGAAG
CAACGCGAAGAACCTTACCAGGCCTTGACATCCAATGAACTTTCCAGAGATGGATTGGTGCCTTCGGGAACATTGAGACA
GGTGCTGCATGGCTGTCGTCAGCTCGTGTCGTGAGATGTTGGGTTAAGTCCCGTAACGAGCGCAACCCTTGTCCTTAGTT
ACCAGCACGTTATGGTGGGCACTCTAAGGAGACTGCCGGTGACAAACCGGAGGAAGGTGGGGATGACGTCAAGTCATCAT
GGCCCTTACGGCCTGGGCTACACACGTGCTACAATGGTCGGTACAGAGGGTTGCCAAGCCGCGAGGTGGAGCTAATCCCA
TAAAACCGATCGTAGTCCGGATCGCAGTCTGCAACTCGACTGCGTGAAGTCGGAATCGCTAGTAATCGCGAATCAGAATG
TCGCGGTGAATACGTTCCCGGGCCTTGTACACACCGCCCGTCACACCATGGGAGTGGGTTGCACCAGAAGTAGCTAGTCT

AACCTTCGGGAGGACGGTTACCACGGTGTGATTCATGACTGGGGTGAAGTCGTAACAAGGTAGCCGTAGGGGAACCTGCG
GCTGGATCACCTCCTTAAATGGACGACAACAAGAAGAAAGCCTTGGCTGCGGCCCTGGGTCAGATCGAACGTCAATTCGG
CAAGGGTGCCGTAATGCGTATGGGCGATCACGACCGCCAGGCGATCCCGGCCATTTCCACTGGCTCTCTGGGTCTGGACA
TCGCACTCGGCATCGGCGGCCTGCCAAAAGGCCGTATTGTTGAAATCTACGGTCCGGAATCGTCCGGTAAAACCACCCTG
ACCCTGTCGGTGATTGCCCAGGCACAGAAGATGGGCGCCACCTGCGCCTTCGTCGACGCCGAGCACGCACTGGACCCGGA
ATACGCCGGCAAACTGGGGGTCAACGTTGACGACCTGCTGGTTTCCCAGCCGGACACCGGCGAACAGGCGCTGGAAATCA
CCGACATGCTGGTGCGCTCCAATGCCATCGACGTGATCGTGATCGACTCCGTGGCGGCACTGGTACCCAAGGCCGAGATC
GAAGGCGAGATGGGCGACATGCACGTGGGCCTGCAGGCCCGCCTGATGTCCCAGGCGCTGCGCAAGATCACCGGTAACAT
CAAGAACGCCAACTGCCTGGTGATCTTCATCAACCAGATCCGTATGAAAATCGGCGTGATGTTCGGCAGCCCGGAAACCA
CCACCGGTGGTAACGCGCTGAAGTTCTACGCTTCGGTTCGTCTGGACATCCGTCGTACTGGCGCGGTGAAGGAAGGCGAC
GAAGTCGTCGGTAGCGAAACCCGGGTCAAGATCGTCAAGAACAAGGTGGCTCCACCGTTCCGTCAGGCTGAATTCCAGAT
CCTGTACGGCAAGGGTATCTACCTGAACGGCGAGATCATCGATCTGGGCGTGCTGCACGGTTTCCTCGAGAAGTCCGGTG
CCTGGTACAGCTACCAGGGCAACAAGATCGGTCAGGGCAAGGCCAACTCGGCCAAGTTCCTGCAGGACAATCCGGAAATC
GGCAATGCCCTCGAGAAGCAGATTCGCGACAAGCTGCTGGCTCCAACCGCTGATGTCAAAGCTTCGCCGGTCAACGAGAC
CATCGATGACATGGCTGACGCGGATATCTGAATGAGCGAAGAAAACACGTACGACTCGAGCAGCATTAAAGTGCTGAAAG
GTTTGGATGCCGTACGCAAACGTCCCGGTATGTACATTGGTGACACCGACGATGGCAGCGGTCTGCACCATATGGTGTTC
GAGGTGGTCGATAACTCGATCGACGAAGCTCTGGCCGGCCACTGCGACGACATCAGCATCATCATCCACCCAGACGAATC
CATTACCGTGCGTGACAACGGTCGCGGCATCCCGGTAGACGTGCATAAAGAAGAAGGCGTTTCCGCGGCCGAGGTCATCA
TGACTGTGCTGCACGCCGGCGGTAAGTTCGACGACAACTCCTACAAAGTATCCGGCGGTCTGCACGGTGTGGGTGTGTCG
GTAGTGAACGCCCTGTCCGAAGAACTGGTCCTGACCGTTCGCCGCAGTGGCAAGATCTGGGAACAGACCTACGTTCACGG
TGTGCCTCAGGCGCCTATGGCGATCGTCGGTGACAGTGAAACCACCGGTACCCAGATTCACTTCAAGGCTTCCAGCGAGA
CCTTCAAGAACATCCATTTCAGCTGGGACATCCTGGCCAAGCGGATTCGTGAACTGTCCTTCCTCAACTCCGGTGTCGGT
ATCGTTCTGAAGGACGAGCGCAGCGGCAAGGAAGAACTGTTCAAGTACGAAGGCGGTCTGCGTGCGTTCGTTGAATACCT
GAACACCAACAAGACCGCGGTCAACCAGGTGTTCCACTTCAATGTGCAGCGTGAAGATGGCATCGGCGTGGAAATCGCCC
TGCAGTGGAACGACAGCTTCAACGAAAACCTGCAGTGCTTCACCAACAACATTCCGCAGCGCGACGGCGGCACCCACTTG
GTGGGCTTCCGTTCGGCACTGACACGTAACCTGAACAACTACATCGAACAGGAAGGTCTGGCGAAGAAGCACAAGGTCGC
CACCACCGGTGACGATGCCCGCGAAGGCCTGACCGCGATCATTTCGGTCAAGGTGCCGGATCCGAAGTTCAGCTCCCAGA
CCAAAGACAAGCTGGTGTCTTCCGAAGTGAAGACCGCGGTCGAACAGGAAATGGGCAAGTACTTCTCCGACTTCCTGCTG
GAAAACCCGAACGAAGCCAAGCTGGTGGTCGGCAAGATGCTCGACGCCGCCCGTGCCCGTGAAGCGGCGCGTAAGGCTCG
CGAGATGACCCGCCGTAAAGGTGCGCTGGATATCGCCGGCCTGCCGGGCAAACTGGCGGACTGCCAGGAAAAAGACCCTG
CCCTTTCCGAACTCTACCTGGTGGAAGGTGACTCTGCTGGCGGCTCCGCCAAGCAGGGACGCAACCGTAAGACCCAGGCG
ATTCTGCCGCTCAAGGGCAAGATCCTTAACGTCGAGAAAGCGCGTTTCGACAAGATGATTTCCTCGCAAGAGGTCGGCAC
CTTGATCACTGCACTCGGTTGCGGCATCGGCCGCGAAGAGTACAACATCGACAAGCTGCGTTATCACAACATCATCATCA
TGACCGACGCCGACGTCGACGGTTCGCACATCCGTACCCTGCTGCTGACCTTCTTCTTCCGTCAGCTGCCGGAGCTGATC
GAGCGTGGCTACATCTACATCGCTCAACCGCCGCTGTACAAGGTCAAGAAAGGCAAGCAAGAGCAATACATCAAAGACGA
CGACGCCATGGAAGAGTACATGACGCAGTCGGCCCTGGAAGATGCGAGCCTGCACCTGAACGAAGACGCACCGGGTATTT
CCGGCGAGGCGCTGGAGCGCCTGGTGAACGACTTCCGCATGGTCATGAAAACCCTCAAGCGTCTGTCGCGCCTGTACCCT
CAGGAGCTGACCGAGCACTTCATCTACCTGCCGGCCGTGAGCCTGGAGCAGCTCTCCGATCACGCGGCGATGCAGGATTG
GCTGGCCCAATATGAAGTCCGCCTGCGCACCGTCGAGAAGTCCGGCCTGGTCTACAAGGCCAGCCTGCGTGAAGACCGTG
AACGTAATGTCTGGCTGCCAGAGGTCGAACTGATCTCCCACGGCCTGTCGAACTACGTCACCTTCAACCGCGACTTCTTC
GGCAGCAATGACTACAAGACCGTCGTCACCCTCGGTGCTCAACTGAGCTCCCTGCTGGACGAAGGCGCTTATATTCAGCG
TGGCGAACGCAAGAAGGCAGTGACCGAGTTCAAGGAAGCCCTGGACTGGCTGATGACCGAAAGTACCAAGCGCCACACCA
TCCAGCGATACAAAGGTCTGGGCGAGATGAACCCGGATCAGCTGTGGGAAACCACTATGGACCCAAGCGTGCGCCGTATG
CTCAAGGTCACCATCGAAGACGCCATCGGCGCCGACCAGATCTTCAACACCCTGATGGGTGATGCGGTCGAGCCTCGTCG
CGACTTCATCGAAAGCAACGCCCTGGCGGTATCCAACCTGGACTTCTGAATGTCCGGAAAAGCGCAACAGCAGTCTCGCC
TCAAAGAGTTGATCAGCCGTGGTCGTGAGCAGGGTTACCTGACTTACGCGGAGGTCAACGACCACCTGCCGGAGGATATT
TCAGATCCGGAACAGGTGGAAGACATCATCCGCATGATCAACGACATGGGGATCAACGTATTCGAGAGTGCTCCGGATGC
GGATGCCCTTTTGTTGGCCGAAGCCGATACCGACGAAGCAGCAGCTGAAGAAGCCGCCGCAGCGTTGGCGGCTGTGGAAA
CCGACATTGGTCGCACTACCGACCCCGTGCGTATGTACATGCGCGAAATGGGTACGGTAGAGCTGCTCACACGTGAAGGC
GAAATCGAAATCGCCAAGCGTATCGAAGAGGGCATCCGTGAAGTGATGGGCGCGATCGCGCACTTCCCTGGCACGGTTGA
GCACATCCTCTCCGAATACACTCGCGTCACCACCGAAGGTGGCCGCCTGTCCGACGTCCTGAGCGGTTACATCGACCCGG
ACGACGGTATTGCGCCGCCTGCCGCCGAAGTACCACCGCCTGTCGATGCCAAGGCCGCAAAAGCGGACGACGACACCGAC
GACGATGACGCCGAAGCCAGTGACGACGAAGAAGAAGCCGAAAGCGGTCCGGATCCGGTCATCGCAGCCCAGCGCTTTGG
CGCCGTTGCCGACCAGATGGAAATTACCCGCAAGGCGCTGAAAAAGCACGGTCGCGAACACAAGCAAGCCCTGGCTGAAA
TGCTGGCCCTGGCTGAGCTGTTCATGCCGATCAAACTGGTTCCGAAGCAATTCGAAGGCCTGGTTGAACGTGTTCGTAGC
GCCCTGGATCGCCTGCGTCAGCAAGAGCGCGCGATCATGCAGCTCTGTGTTCGTGATGCCCGCATGCCACGCGCCGACTT
CCTGCGCCAGTTCCCTGGCAATGAAGTGGACGAAAGCTGGTCCGACGCACTGGCCAAAGGCAAGGCCAAGTACGCCGAAT
CCATTGGCCGCCTGCAGCCGGATATCATCCGTTGCCAGCAGAAGCTGACCGCGCTCGAGACCGAGACCGGCCTGACGATC
GCCGAGATCAAGGACATCAACCGTCGCATGTCGATCGGCGAGGCCAAGGCCCGTCGCGCGAAGAAAGAGATGGTCGAAGC
CAACTTGCGTCTGGTGATCTCCATCGCCAAGAAGTACACCAACCGTGGCCTGCAATTCCTCGACCTGATCCAGGAAGGCA
ACATCGGTTTGATGAAAGCGGTAGACAAGTTCGAATACCGTCGCGGCTACAAGTTCTCGACTTATGCCACCTGGTGGATC
CGTCAGGCGATCACTCGCTCGATCGCCGACCAGGCCCGCACCATCCGTATTCCGGTGCACATGATCGAGACGATCAACAA
GCTCAACCGTATTTCCCGGCAGATGTTGCAGGAAATGGGTCGCGAACCGACTCCGGAAGAGCTGGGCGAACGCATGGAAA
TGCCTGAAGACAAGATCCGCAAGGTATTGAAGATCGCTAAAGAGCCGATCTCCATGGAAACCCCGATCGGTGATGACGAA

GACTCCCATCTGGGTGACTTCATCGAAGACTCGACCATGCAGTCGCCAATCGATGTCGCCACCGTTGAGAGCCTTAAAGA
AGCGACTCGCGAAGTACTCTCCGGCCTCACTGCCCGTGAAGCCAAGGTACTGCGCATGCGCTTCGGCATCGACATGAATA
CCGACCACACCCTCGAGGAAGTCGGTAAGCAGTTCGACGTTACCCGTGAGCGGATTCGTCAGATCGAAGCCAAGGCGCTG
CGCAAGCTGCGCCACCCGACGCGAAGCGAGCACCTGCGCTCCTTCCTCGACGAGTAATTGACTAAGCCAGCCATACTCGC
CCTTGCTGATGGCAGCATTTTTCGCGGCGAAGCCATTGGAGCCGACGGTCAGACCGTTGGTGAGGTGGTGTTTAACACCG
CAATGACCGGCTATCAGGAAATCCTTACCGATCCTTCCTACGCCCAACAGATCGTTACCCTGACTTACCCGCACATCGGC
AATACCGGCACCACGCCGGAAGACGCCGAGTCCGATCGTGTCTGGTCGGCCGGTCTGGTGATTCGCGACCTGCCACTGGT
TGCGAGCAACTGGCGTAACACCCTGTCCCTGTCCGACTACCTGAAAGCCAACAACGTTGTGGCGATCGCCGGTATCGACA
CCCGTCGCCTGACGCGCATCCTGCGCGAGAAAGGCGCGCAGAACGGCTGCATCATGGCCGGCGACAATATCTCCGACGAA
GCGGCGATTGCCGCTGCGCGCGGCTTCCCTGGCCTGAAAGGCATGGATCTGGCGAAGGTCGTCAGCACCAAGGAAAGCTA
CGAATGGCGTTCCAGCGTCTGGAGCCTGAAGACCGACAGTCATCCGACCATCGAGGCTTCCGAGCTGCCTTACCACGTGG
TTGCCTACGACTACGGCGTCAAGCTGAACATCCTGCGCATGCTGGTCGAGCGCGGTTGCCGCGTGACCGTGGTACCTGCG
CAAACCCCGGCCAGCGACGTCCTGGCGCTCAAGCCTGACGGTGTGTTTCTGTCCAACGGTCCTGGCGACCCCGAGCCTTG
CGATTACGCGATCCAGGCGATCAAGGACGTGCTGGAGACCGAGATCCCGGTCTTCGGTATCTGCCTGGGCCACCAACTGC
TGGCGCTGGCCGCCGGCGCCAAGACAGTGAAGATGGGCCACGGCCACCACGGTGCCAACCACCCGGTCCAGGACCTGGAC
AGCGGTGTAGTGATGATCACCAGCCAGAACCACGGTTTTGCGGTGGACGAAACCACCCTGCCGGGCAACGTGCGGGCGAT
CCACAAGTCGCTGTTCGACGGCACCCTGCAAGGCATCGAGTTGACCGACAAGAGCGCATTCAGCTTCCAGGGGCACCCTG
AAGCGAGCCCGGGCCCGAACGATGTGGCGCCGCTGTTCGATCGTTTCATCAACGAGATGGCCAAGCGACGCTGAATGAGT
AGCGGACGTATCGTTCAAATCATCGGCGCCGTTATCGACGTGGAATTTCCACGCGACAGCGTACCGAGCATCTACGACGC
CTTGAAGGTTCAAGGCGCCGAAACCACTCTGGAAGTTCAGCAGCAGCTGGGCGACGGCGTGGTACGTACCATTGCGATGG
GCTCCACCGAGGGCTTGAAGCGCGGTCTGGACGTCAACAACACTGGCGCAGCCATCTCCGTACCGGTCGGTAAAGCGACC
CTGGGCCGGATCATGGACGTACTGGGCAACCCGATCGACGAAGCTGGCCCGATCGGCGAAGAAGAGCGTTGGGGCATTCA
CCGTCCTGCGCCGACCTTCGCTGAACAAGCTGGCGGCAACGACCTGCTGGAAACCGGCATCAAGGTTATCGACCTGGTTT
GCCCGTTCGCCAAGGGCGGTAAAGTCGGTCTGTTCGGTGGTGCCGGTGTGGGCAAAACCGTAAACATGATGGAACTGATC
CGTAACATCGCCATCGAGCACAGCGGTTATTCCGTGTTCGCCGGTGTGGGTGAGCGTACTCGTGAGGGTAACGACTTCTA
CCACGAGATGAAGGATTCCAACGTTCTGGACAAAGTGGCACTGGTATACGGCCAGATGAACGAGCCGCCGGGAAACCGTC
TGCGCGTAGCTCTGACCGGCCTGACCATGGCCGAGAAGTTCCGTGACGAAGGTAACGACGTTCTGCTGTTCGTCGACAAC
ATCTATCGTTACACCCTGGCCGGTACCGAAGTATCCGCACTGCTGGGCCGTATGCCTTCGGCAGTAGGTTACCAGCCGAC
CCTGGCTGAAGAGATGGGCGTTCTGCAAGAACGTATCACTTCGACCAAGCAAGGCTCGATCACCTCGATCCAAGCGGTAT
ACGTGCCTGCGGACGACTTGACCGACCCGTCGCCAGCGACCACCTTCGCCCACTTGGACGCCACCGTCGTTCTGTCCCGT
GACATCGCTTCCCTGGGTATCTACCCAGCGGTAGACCCACTGGACTCGACTTCCCGTCAGCTGGACCCGAACGTGATCGG
CAACGAGCACTACGAAACCGCTCGCGGCGTTCAGTACGTGCTGCAGCGCTACAAAGAGCTGAAGGACATCATTGCGATCC
TGGGTATGGACGAACTGTCCGAAGCCGACAAGCAACTGGTATCCCGCGCTCGTAAGATCCAGCGCTTCCTGTCGCAGCCG
TTCTTCGTGGCTGAAGTCTTCACTGGTTCTCCAGGCAAATACGTTTCCCTGAAAGACACCATCGCTGGCTTCAAAGGCAT
CCTCAACGGTGACTACGACCATCTGCCAGAACAAGCGTTCTACATGGTTGGTGGCATCGAAGAAGCGATCGAGAAAGCCA
AGAAACTGTAA

NCBI Reference Sequence: NZ_CP027708.1
Strain: ATCC17411
Chained genes: 16S rRNA, recA, gyrB, rpoD, carA, atpD
>NZ_CP027708_ATCC17411
GAACTGAAGAGTTTGATCATGGCTCAGATTGAACGCTGGCGGCAGGCCTAACACATGCAAGTCGAGCGGTAGAGAGGTGC
TTGCACCTCTTGAGAGCGGCGGACGGGTGAGTAATGCCTAGGAATCTGCCTGGTAGTGGGGGATAACGTTCGGAAACGGA
CGCTAATACCGCATACGTCCTACGGGAGAAAGCAGGGGACCTTCGGGCCTTGCGCTATCAGATGAGCCTAGGTCGGATTA
GCTAGTTGGTGAGGTAATGGCTCACCAAGGCGACGATCCGTAACTGGTCTGAGAGGATGATCAGTCACACTGGAACTGAG
ACACGGTCCAGACTCCTACGGGAGGCAGCAGTGGGGAATATTGGACAATGGGCGAAAGCCTGATCCAGCCATGCCGCGTG
TGTGAAGAAGGTCTTCGGATTGTAAAGCACTTTAAGTTGGGAGGAAGGGTACTTACCTAATACGTGAGTATTTTGACGTT
ACCGACAGAATAAGCACCGGCTAACTCTGTGCCAGCAGCCGCGGTAATACAGAGGGTGCAAGCGTTAATCGGAATTACTG
GGCGTAAAGCGCGCGTAGGTGGTTCGTTAAGTTGAATGTGAAATCCCCGGGCTCAACCTGGGAACTGCATCCAAAACTGG
CGAGCTAGAGTATGGTAGAGGGTGGTGGAATTTCCTGTGTAGCGGTGAAATGCGTAGATATAGGAAGGAACACCAGTGGC
GAAGGCGACCACCTGGACTGATACTGACACTGAGGTGCGAAAGCGTGGGGAGCAAACAGGATTAGATACCCTGGTAGTCC
ACGCCGTAAACGATGTCAACTAGCCGTTGGGAGCCTTGAGCTCTTAGTGGCGCAGCTAACGCATTAAGTTGACCGCCTGG
GGAGTACGGCCGCAAGGTTAAAACTCAAATGAATTGACGGGGGCCCGCACAAGCGGTGGAGCATGTGGTTTAATTCGAAG
CAACGCGAAGAACCTTACCAGGCCTTGACATCCAATGAACTTTCCAGAGATGGATTGGTGCCTTCGGGAACATTGAGACA
GGTGCTGCATGGCTGTCGTCAGCTCGTGTCGTGAGATGTTGGGTTAAGTCCCGTAACGAGCGCAACCCTTGTCCTTAGTT
ACCAGCACGTAATGGTGGGCACTCTAAGGAGACTGCCGGTGACAAACCGGAGGAAGGTGGGGATGACGTCAAGTCATCAT
GGCCCTTACGGCCTGGGCTACACACGTGCTACAATGGTCGGTACAGAGGGTTGCCAAGCCGCGAGGTGGAGCTAATCCCA
TAAAACCGATCGTAGTCCGGATCGCAGTCTGCAACTCGACTGCGTGAAGTCGGAATCGCTAGTAATCGCGAATCAGAATG
TCGCGGTGAATACGTTCCCGGGCCTTGTACACACCGCCCGTCACACCATGGGAGTGGGTTGCACCAGAAGTAGCTAGTCT
AACCTTCGGGAGGACGGTTACCACGGTGTGATTCATGACTGGGGTGAAGTCGTAACAAGGTAGCCGTAGGGGAACCTGCG
GCTGGATCACCTCCTTAAATGGACGACAACAAGAAGAAAGCCTTGGCTGCGGCCCTGGGTCAGATCGAACGTCAATTCGG
CAAGGGTGCCGTAATGCGTATGGGCGATCACGATCGCCAGGCGATCCCGGCCATTTCCACTGGCTCTCTGGGTCTGGACA
TCGCACTCGGCATCGGCGGCCTGCCAAAAGGCCGTATTGTTGAAATCTACGGTCCGGAATCGTCCGGTAAAACCACCCTG
ACCCTGTCGGTGATCGCCCAGGCACAGAAGATGGGCGCCACCTGCGCCTTCGTCGACGCCGAGCACGCACTGGACCCGGA

ATACGCCGGCAAACTGGGGGTCAACGTTGACGACCTGCTGGTTTCCCAGCCGGACACCGGCGAACAGGCGCTGGAAATCA
CCGACATGCTGGTGCGCTCCAATGCCATCGACGTGATCGTGATCGACTCCGTGGCGGCGCTGGTACCCAAGGCCGAGATC
GAAGGCGAGATGGGCGACATGCACGTGGGCCTGCAGGCTCGCCTGATGTCCCAGGCGCTGCGCAAGATCACCGGTAACAT
CAAGAACGCCAACTGCCTGGTGATCTTCATCAACCAGATCCGTATGAAAATCGGCGTGATGTTCGGCAGCCCGGAAACCA
CCACCGGTGGTAACGCGCTGAAGTTCTACGCTTCGGTTCGTCTGGACATCCGTCGTACTGGCGCGGTGAAGGAAGGTGAC
GAAGTCGTCGGTAGCGAAACCCGGGTCAAGATCGTCAAGAACAAGGTGGCTCCACCTTTCCGTCAAGCTGAGTTCCAGAT
CCTGTACGGCAAGGGTATCTACCTGAACGGCGAGATCATCGATCTGGGCGTGCTGCACGGTTTCCTCGAGAAGTCCGGTG
CCTGGTACAGCTACCAGGGCAACAAGATCGGTCAGGGCAAGGCCAACTCGGCCAAGTTCCTGCAGGACAATCCGGAAATC
GGTAATGCCCTCGAGAAGCAGATTCGCGACAAGCTGCTGGCTCCGAGCGGAGATACCAAGGCTCTGCCCGTCAACGAGAC
CATCGATGACATGGCCGACGCGGATATCTGAATGAGCGAAGAAAACACGTACGACTCGAGCAGCATTAAAGTGCTGAAAG
GTTTGGATGCCGTACGCAAACGTCCCGGTATGTACATTGGTGACACCGACGATGGCAGCGGTCTGCACCATATGGTGTTC
GAGGTGGTCGACAACTCGATCGACGAAGCTCTGGCCGGCCACTGCGACGACATCAGCATCATCATCCACCCGGACGAATC
CATTACCGTGCGTGACAACGGTCGCGGCATCCCGGTAGACGTGCATAAAGAAGAAGGCGTTTCCGCAGCCGAGGTCATCA
TGACCGTGCTGCACGCCGGCGGTAAGTTCGACGACAACTCCTACAAGGTATCCGGCGGTCTGCACGGTGTGGGTGTGTCG
GTAGTGAACGCCCTGTCCGAAGAACTGGTGCTGACCGTTCGCCGCAGTGGCAAGATCTGGGAACAGACCTACGTTCACGG
TGTGCCTCAGGCGCCTATGGCAATTGTCGGTGACAGCGAGACCACCGGTACCCAGATTCACTTCAAGGCTTCCAGCGAGA
CCTTCAAGAACATCCATTTCAGCTGGGACATCCTGGCCAAGCGGATTCGTGAACTGTCCTTCCTCAACTCCGGTGTCGGT
ATCGTTCTGAAGGACGAGCGCAGCGGCAAGGAAGAGCTGTTCAAGTACGAAGGCGGCCTGCGCGCATTCGTTGAATACCT
GAACACCAACAAGACTGCGGTCAACCAGGTGTTCCACTTCAACGTGCAGCGTGAAGACGGCATCGGCGTGGAAATCGCCC
TGCAGTGGAACGACAGCTTCAACGAAAACCTGCAGTGCTTCACCAACAACATTCCGCAGCGCGACGGCGGCACCCACCTG
GTGGGCTTCCGCTCGGCACTGACGCGTAACCTGAACAACTACATCGAGCAGGAAGGTCTGGCGAAGAAGCACAAGGTGGC
CACCACCGGTGACGATGCCCGCGAAGGCCTGACCGCGATCATTTCGGTCAAGGTGCCGGATCCGAAGTTCAGCTCCCAGA
CCAAAGATAAGCTGGTGTCTTCCGAAGTGAAGACCGCAGTCGAACAGGAAATGGGCAAGTACTTCTCCGACTTCCTGCTG
GAAAACCCGAACGAAGCCAAGCTGGTGGTCGGCAAGATGCTCGACGCCGCCCGTGCCCGTGAAGCGGCGCGTAAGGCTCG
TGAGATGACCCGCCGTAAAGGTGCGCTGGATATCGCCGGCCTGCCGGGCAAACTGGCGGACTGCCAGGAAAAGGACCCTG
CCCTTTCCGAACTCTACCTGGTGGAAGGTGACTCTGCTGGCGGCTCCGCCAAGCAGGGACGCAACCGCAAGACCCAGGCG
ATTCTGCCGCTCAAGGGCAAGATTCTTAACGTCGAGAAAGCGCGCTTCGACAAGATGATTTCCTCGCAAGAGGTCGGCAC
CTTGATCACTGCACTCGGCTGCGGCATCGGCCGCGAAGAGTACAACATCGACAAGCTGCGTTATCACAACATCATCATCA
TGACCGACGCCGACGTCGACGGTTCGCACATCCGCACCCTGCTGCTGACTTTCTTCTTCCGTCAGCTGCCGGAGCTGATC
GAGCGTGGCTACATCTACATCGCTCAACCGCCGCTGTACAAGGTCAAGAAAGGCAAGCAAGAGCAATACATCAAAGACGA
CGACGCCATGGAAGAGTACATGACGCAGTCGGCCCTGGAAGATGCGAGCCTGCACCTGAACGAAGAAGCACCGGGTATTT
CCGGCGAGGCGCTGGAGCGCCTGGTGAACGACTTCCGCATGGTCATGAAAACCCTCAAGCGCCTGTCGCGCCTGTACCCT
CAGGAGCTGACCGAACACTTCATCTACCTGCCAGCCGTGAGCCTGGAGCAACTCTCCGATCACGCAGCGATGCAGGATTG
GTTGGCCCAATATGAAGTCCGCCTGCGCACCGTCGAGAAGTCCGGCCTGGTCTACAAGGCCAGCCTGCGTGAAGACCGTG
AACGTAATGTCTGGCTGCCAGAGGTCGAACTGATCTCCCACGGCCTGTCGAACTACGTCACCTTCAACCGCGACTTCTTC
GGCAGCAATGACTACAAGACCGTCGTCACCCTCGGCGCTCAACTGAGCTCCCTGCTGGACGAAGGCGCTTATATTCAGCG
TGGCGAACGCAAGAAGGCGGTGACCGAGTTCAAGGAAGCCCTGGACTGGCTGATGACCGAAAGCACCAAGCGCCACACCA
TCCAGCGATACAAAGGTCTGGGCGAGATGAACCCGGATCAGCTGTGGGAAACCACCATGGACCCAAGCGTGCGCCGCATG
CTCAAGGTCACCATCGAAGACGCCATCGGCGCCGACCAGATCTTCAACACCCTGATGGGTGATGCGGTCGAGCCTCGTCG
CGACTTCATCGAAAGCAACGCCCTGGCGGTATCCAACCTGGACTTCTGAATGTCCGGAAAAGCGCAACAGCAGTCTCGCC
TCAAAGAGTTGATCAGCCGTGGTCGTGAGCAGGGTTACCTGACTTACGCGGAGGTCAACGACCACCTGCCGGAGGATATT
TCAGATCCGGAACAGGTGGAAGACATCATCCGCATGATCAACGACATGGGGATCAACGTATTCGAGAGTGCTCCGGATGC
GGATGCCCTTTTGTTGGCCGAAGCCGATACCGACGAAGCAGCAGCTGAAGAAGCCGCCGCAGCGTTGGCGGCTGTGGAAA
CCGACATTGGTCGCACTACCGACCCAGTGCGTATGTACATGCGCGAAATGGGCACGGTAGAGCTGCTCACACGTGAAGGC
GAAATCGAAATCGCCAAGCGTATCGAAGAGGGCATCCGTGAAGTGATGGGCGCGATCGCGCACTTCCCTGGCACGGTTGA
GCACATCCTCTCCGAATACACTCGCGTCACCACCGAAGGTGGCCGCCTGTCCGACGTCCTGAGCGGTTACATCGACCCGG
ATGACGGCATTGCGCCGCCTGCCGCCGAAGTACCACCGCCTGTCGATGCCAAGGCCGCGAAAGCGGACGACGACACCGAC
GACGATGACGCCGAAGCCAGTGACGACGAAGAAGAAGCCGAAAGCGGTCCGGATCCGGTCATCGCAGCCCAGCGCTTTGG
CGCCGTTGCCGACCAGATGGAAATCACCCGCAAGGCGCTGAAAAAGCACGGTCGCGAACACAAGCAAGCCCTGGCTGAAA
TGCTGGCCCTGGCTGAACTGTTCATGCCGATCAAACTGGTTCCGAAGCAATTCGAAGGCCTGGTTGAACGTGTTCGTAGC
GCCCTGGATCGCCTGCGTCAGCAAGAGCGCGCGATCATGCAGCTCTGTGTTCGTGATGCCCGCATGCCACGCGCCGACTT
CCTGCGCCAGTTCCCTGGCAATGAAGTGGACGAAAGCTGGTCCGACGCGCTGGCCAAAGGCAAGGCCAAGTACGCCGAAG
CCATCGGCCGCCTGCAGCCGGACATCATCCGTTGCCAGCAGAAGCTGACCGCGCTCGAGACCGAGACCGGCCTGACGATT
GCCGAGATCAAGGACATCAACCGTCGCATGTCGATCGGCGAGGCCAAGGCCCGTCGCGCGAAGAAAGAGATGGTCGAAGC
CAACTTGCGCCTGGTGATCTCCATCGCCAAGAAGTACACCAACCGTGGCCTGCAGTTCCTCGACCTGATCCAGGAAGGCA
ACATCGGTTTGATGAAAGCGGTAGACAAGTTCGAATACCGTCGCGGCTACAAGTTCTCGACTTATGCCACCTGGTGGATC
CGTCAGGCGATCACTCGCTCGATCGCCGACCAGGCCCGCACCATCCGTATTCCGGTGCACATGATCGAGACGATCAACAA
GCTCAACCGTATTTCCCGGCAGATGTTGCAGGAAATGGGTCGCGAACCGACTCCGGAAGAGCTGGGCGAACGCATGGAAA
TGCCTGAGGACAAGATCCGCAAGGTATTGAAGATCGCTAAAGAGCCGATCTCCATGGAAACCCCGATCGGTGATGACGAA
GACTCCCATCTGGGTGACTTCATCGAAGACTCGACCATGCAGTCGCCAATCGATGTCGCCACCGTTGAGAGCCTCAAGGA
AGCGACTCGCGAAGTCCTCTCCGGCCTCACTGCCCGTGAAGCCAAGGTACTGCGCATGCGCTTCGGCATCGACATGAATA
CCGACCACACCCTTGAGGAAGTCGGTAAGCAGTTCGACGTTACCCGTGAGCGGATTCGTCAGATCGAAGCCAAGGCGCTG
CGCAAGCTGCGCCACCCGACGAGAAGCGAGCATCTGCGCTCCTTCCTCGACGAGTAATTGACTAAGCCAGCCATACTCGC
CCTTGCTGATGGCAGCATTTTTCGCGGCGAAGCCATTGGAGCCGACGGTCAGACCGTTGGTGAGGTGGTGTTTAACACCG

CAATGACCGGCTATCAGGAAATCCTTACCGATCCTTCCTACGCCCAACAGATCGTTACCCTGACTTACCCGCACATCGGC
AACACTGGCACCACGCCGGAAGACGCCGAGTCCGATCGTGTCTGGTCGGCCGGTCTGGTGATTCGCGACCTGCCACTGGT
TGCGAGCAACTGGCGTAACACCCTGTCCCTGTCCGATTACCTGAAAGCCAACAATGTCGTGGCGATCGCCGGTATCGACA
CCCGTCGCCTGACGCGCATCCTGCGCGAGAAAGGCGCACAGAACGGCTGCATCATGGCCGGCGACAACATCTCCGACGAA
GCGGCGATTGCTGCTGCACGCGGCTTCCCTGGCCTGAAAGGCATGGATCTGGCGAAGGTCGTCAGCACCAAGGAAAGCTA
CGAGTGGCGCTCCAGCGTCTGGAGCCTGAAGACCGACAGTCATCCGACTATCGAGGCTTCCGAGCTGCCTTACCACGTGG
TTGCCTACGACTACGGCGTCAAGCTGAACATCCTGCGCATGCTGGTCGAGCGCGGTTGCCGCGTGACCGTAGTGCCTGCG
CAAACCCCGGCCAGCGACGTCCTGGCACTCAAGCCTGACGGTGTGTTCCTGTCCAACGGTCCTGGCGACCCCGAGCCTTG
CGATTACGCCATCCAGGCGATCAAGGACGTGCTGGAAACCGAGATTCCGGTCTTCGGTATCTGCCTGGGCCACCAACTGC
TGGCACTGGCCTCCGGCGCCAAGACGGTGAAAATGGGCCACGGCCACCACGGCGCCAACCACCCGGTCCAGGACCTGGAC
AGCGGTGTGGTGATGATCACCAGCCAGAACCACGGTTTTGCGGTGGACGAAACCACCCTGCCGGGCAACGTGCGGGCGAT
CCACAAGTCGCTGTTCGATGGCACCCTGCAAGGCATCGAGCGTACCGACAAGAGCGCATTCAGCTTCCAGGGCCACCCTG
AAGCGAGCCCGGGCCCGAACGATGTGGCGCCGCTGTTCGATCGTTTCATCAACGAGATGGCCAAGCGACGCTGAATGAGT
AGCGGACGTATCGTTCAAATCATCGGCGCCGTTATCGACGTGGAATTTCCACGCGACAGCGTACCGAGCATCTACGACGC
CTTGAAGGTTCAAGGCGCCGAAACCACTCTGGAAGTTCAGCAGCAGCTGGGCGACGGCGTGGTACGTACCATTGCGATGG
GCTCCACCGAGGGCTTGAAGCGCGGTCTGGACGTCAACAACACTGGCGCCGCCATCTCCGTACCGGTCGGTAAAGCGACC
CTGGGCCGGATCATGGACGTGCTGGGCAACCCGATCGACGAAGCTGGCCCGATCGGCGAAGAAGAGCGTTGGGGCATTCA
CCGTCCTGCGCCGACCTTCGCTGAACAAGCTGGCGGCAACGACCTCCTGGAAACCGGCATCAAGGTTATCGACCTGGTTT
GCCCGTTCGCCAAGGGCGGTAAAGTCGGTCTGTTCGGTGGTGCCGGTGTGGGCAAAACCGTAAACATGATGGAACTGATC
CGTAACATCGCCATCGAGCACAGCGGTTATTCCGTGTTCGCTGGTGTGGGTGAGCGTACTCGTGAGGGTAACGACTTCTA
CCACGAGATGAAGGATTCCAACGTTCTGGACAAAGTGGCACTGGTATACGGCCAGATGAACGAGCCGCCGGGAAACCGTC
TGCGCGTAGCTCTGACCGGCCTGACCATGGCCGAGAAGTTCCGTGACGAAGGTAACGACGTTCTGCTGTTCGTCGACAAC
ATCTATCGTTACACCCTGGCCGGTACCGAAGTATCCGCACTGCTGGGCCGTATGCCTTCGGCAGTAGGTTACCAGCCGAC
CCTGGCTGAAGAGATGGGCGTTCTGCAAGAACGTATCACTTCGACCAAGCAAGGCTCGATCACCTCGATCCAAGCGGTAT
ACGTACCTGCGGACGACTTGACCGACCCGTCGCCAGCGACCACCTTCGCCCACTTGGACGCCACCGTCGTACTGTCCCGT
GACATCGCTTCCCTGGGTATCTACCCAGCGGTAGACCCACTGGATTCGACTTCCCGTCAGCTGGACCCGAACGTGATCGG
CAACGAGCACTACGAAACCGCTCGCGGCGTTCAGTACGTGCTGCAGCGCTACAAAGAGCTGAAGGACATCATTGCGATCC
TGGGTATGGACGAACTGTCCGAAGCCGACAAGCAACTGGTATCCCGCGCTCGTAAGATCCAGCGCTTCCTGTCGCAGCCG
TTCTTCGTGGCTGAAGTCTTCACTGGTTCTCCAGGCAAATACGTTTCCCTGAAAGACACCATCGCTGGCTTCAAAGGCAT
CCTCAACGGTGACTACGACCACCTGCCAGAACAAGCGTTCTACATGGTCGGCGGCATCGAAGAAGCGATCGAGAAAGCCA
AGAAACTGTAA

NCBI Reference Sequence: NZ_CP027722.1
Strain: C50
Chained genes: 16S rRNA, recA, gyrB, rpoD, carA, atpD
>NZ_CP027722_C50
GAACTGAAGAGTTTGATCATGGCTCAGATTGAACGCTGGCGGCAGGCCTAACACATGCAAGTCGAGCGGTAGAGAGAAGC
TTGCTTCTCTTGAGAGCGGCGGACGGGTGAGTAATGCCTAGGAATCTGCCTGGTAGTGGGGGATAACGTTCGGAAACGGA
CGCTAATACCGCATACGTCCTACGGGAGAAAGCAGGGGACCTTCGGGTCTTGCGCTATCAGATGAGCCTAGGTCGGATTA
GCTAGTTGGTGAGGTAATGGCTCACCAAGGCGACGATCCGTAACTGGTCTGAGAGGATGATCAGTCACACTGGAACTGAG
ACACGGTCCAGACTCCTACGGGAGGCAGCAGTGGGGAATATTGGACAATGGGCGAAAGCCTGATCCAGCCATGCCGCGTG
TGTGAAGAAGGTCTTCGGATTGTAAAGCACTTTAAGTTGGGAGGAAGGGTACTTACCTAATACGTGAGTATTTTGACGTT
ACCGACAGAATAAGCACCGGCTAACTCTGTGCCAGCAGCCGCGGTAATACAGAGGGTGCAAGCGTTAATCGGAATTACTG
GGCGTAAAGCGCGCGTAGGTGGTTCGTTAAGTTGGATGTGAAATCCCCGGGCTCAACCTGGGAACTGCATCCAAAACTGG
CGAGCTAGAGTATGGTAGAGGGTGGTGGAATTTCCTGTGTAGCGGTGAAATGCGTAGATATAGGAAGGAACACCAGTGGC
GAAGGCGACCACCTGGACTGATACTGACACTGAGGTGCGAAAGCGTGGGGAGCAAACAGGATTAGATACCCTGGTAGTCC
ACGCCGTAAACGATGTCAACTAGCCGTTGGGAGCCTTGAGCTCTTAGTGGCGCAGCTAACGCATTAAGTTGACCGCCTGG
GGAGTACGGCCGCAAGGTTAAAACTCAAATGAATTGACGGGGGCCCGCACAAGCGGTGGAGCATGTGGTTTAATTCGAAG
CAACGCGAAGAACCTTACCAGGCCTTGACATCCAATGAACTTTCCAGAGATGGATTGGTGCCTTCGGGAACATTGAGACA
GGTGCTGCATGGCTGTCGTCAGCTCGTGTCGTGAGATGTTGGGTTAAGTCCCGTAACGAGCGCAACCCTTGTCCTTAGTT
ACCAGCACGTCATGGTGGGCACTCTAAGGAGACTGCCGGTGACAAACCGGAGGAAGGTGGGGATGACGTCAAGTCATCAT
GGCCCTTACGGCCTGGGCTACACACGTGCTACAATGGTCGGTACAGAGGGTTGCCAAGCCGCGAGGTGGAGCTAATCCCA
TAAAACCGATCGTAGTCCGGATCGCAGTCTGCAACTCGACTGCGTGAAGTCGGAATCGCTAGTAATCGCGAATCAGAATG
TCGCGGTGAATACGTTCCCGGGCCTTGTACACACCGCCCGTCACACCATGGGAGTGGGTTGCACCAGAAGTAGCTAGTCT
AACCTTCGGGAGGACGGTTACCACGGTGTGATTCATGACTGGGGTGAAGTCGTAACAAGGTAGCCGTAGGGGAACCTGCG
GCTGGATCACCTCCTTAAATGGACGACAACAAGAAGAAAGCCTTGGCTGCGGCCCTGGGTCAGATCGAACGTCAATTCGG
CAAGGGTGCCGTAATGCGTATGGGCGATCACGATCGCCAGGCGATCCCGGCCATTTCCACTGGCTCTCTGGGTCTGGACA
TCGCGCTCGGCATCGGCGGCCTGCCAAAAGGCCGTATTGTTGAAATCTACGGTCCGGAATCGTCCGGTAAAACCACCCTG
ACCCTGTCCGTGATTGCCCAGGCACAGAAGATGGGCGCCACCTGCGCCTTCGTCGACGCCGAGCACGCACTGGACCCGGA
ATACGCCGGCAAACTGGGGGTCAACGTTGACGACCTGCTGGTTTCCCAGCCGGACACCGGCGAACAGGCGCTGGAAATCA
CCGACATGCTGGTGCGCTCCAATGCCATCGACGTGATCGTGATCGACTCCGTGGCGGCACTGGTACCCAAGGCCGAGATC
GAAGGCGAGATGGGCGACATGCACGTGGGCCTGCAGGCCCGCCTGATGTCCCAGGCGCTGCGCAAGATCACCGGTAACAT
CAAGAACGCCAACTGTCTGGTGATCTTCATCAACCAGATCCGTATGAAAATCGGCGTGATGTTCGGCAGCCCGGAAACCA
CCACCGGCGGTAACGCGCTGAAGTTCTACGCCTCGGTTCGTCTGGACATCCGTCGTACTGGCGCGGTGAAGGAAGGCGAC

GAAGTCGTCGGTAGCGAAACCCGGGTCAAGATCGTCAAGAACAAGGTGGCTCCACCGTTCCGTCAGGCTGAATTCCAGAT
CCTGTACGGCAAGGGTATCTACCTGAACGGCGAGATCATCGATCTGGGCGTGCTGCACGGTTTCCTCGAGAAGTCCGGTG
CCTGGTACAGCTACCAGGGCAACAAGATCGGTCAGGGCAAGGCCAACTCGGCCAAGTTCCTGCAGGACAATCCGGAAATC
GGCAATGCCCTCGAGAAGCAGATTCGCGACAAGCTGCTGGCTCCAACCGCTGATGTCAAAGCTTCGCCGGTCAACGAGAC
CATCGATGACATGGCTGACGCGGATATCTGAATGAGCGAAGAAAACACGTACGACTCGAGCAGCATTAAAGTGCTGAAAG
GTTTGGATGCCGTACGCAAACGTCCCGGTATGTACATTGGTGACACCGACGATGGCAGCGGTCTGCACCATATGGTGTTC
GAGGTGGTCGATAACTCGATCGACGAAGCTCTGGCCGGCCATTGCGACGACATCAGCATCATCATCCACCCGGACGAATC
CATTACCGTGCGTGACAACGGTCGCGGCATCCCGGTAGACGTGCATAAAGAAGAAGGCGTTTCCGCGGCCGAGGTCATCA
TGACCGTACTGCACGCCGGCGGTAAGTTCGACGATAACTCCTACAAAGTATCCGGCGGTCTGCACGGTGTGGGTGTGTCG
GTAGTGAACGCCCTGTCCGAAGAACTGGTCCTGACCGTTCGCCGCAGCGGAAAGATCTGGGAACAGACCTACGTTCACGG
TGTGCCTCAGGCGCCTATGGCGATCGTCGGTGACAGCGAAACCACCGGTACCCAGATTCACTTCAAGGCGTCCAGCGAGA
CCTTCAAGAACATCCATTTCAGCTGGGACATCCTGGCCAAGCGGATTCGTGAACTGTCCTTCCTCAACTCCGGTGTCGGT
ATCGTTCTGAAGGACGAACGCAGTGGCAAGGAAGAGCTGTTCAAGTACGAAGGCGGCCTGCGTGCGTTCGTTGAATACCT
GAACACCAACAAGACCGCGGTCAACCAGGTGTTCCACTTCAATGTGCAGCGTGAAGATGGCATCGGCGTGGAAATCGCCC
TGCAGTGGAACGACAGCTTCAACGAAAACCTGCAGTGCTTCACCAACAACATTCCGCAGCGCGATGGCGGCACCCACTTG
GTGGGCTTCCGTTCGGCACTGACGCGTAACCTGAACAACTACATCGAACAGGAAGGTCTGGCGAAGAAGCACAAGGTCGC
CACCACCGGTGACGATGCCCGCGAAGGCCTGACCGCGATCATTTCGGTCAAGGTGCCGGATCCGAAGTTCAGCTCCCAGA
CCAAAGACAAGCTGGTGTCTTCCGAAGTGAAGACCGCGGTTGAACAGGAAATGGGCAAGTACTTCTCCGACTTCCTGCTG
GAAAACCCGAACGAAGCCAAGCTGGTGGTCGGCAAGATGCTCGACGCCGCCCGTGCCCGTGAAGCGGCGCGTAAGGCTCG
TGAGATGACCCGCCGTAAAGGCGCGCTGGATATCGCCGGCCTGCCGGGCAAACTGGCGGACTGCCAGGAAAAAGACCCTG
CCCTTTCCGAACTCTACCTGGTGGAAGGTGACTCTGCTGGCGGCTCCGCCAAGCAGGGACGCAACCGTAAGACCCAGGCG
ATTCTGCCGCTCAAGGGCAAGATCCTTAACGTCGAGAAAGCGCGCTTCGACAAGATGATTTCCTCGCAAGAGGTCGGCAC
CTTGATCACTGCACTCGGTTGCGGCATCGGCCGCGAAGAGTACAACATCGACAAGCTGCGTTATCACAACATCATCATCA
TGACCGACGCTGACGTCGACGGTTCGCACATCCGTACCCTGCTGCTGACCTTCTTCTTCCGTCAGCTGCCGGAGCTGATC
GAGCGTGGCTACATCTACATCGCTCAACCGCCGCTGTACAAGGTCAAGAAAGGCAAGCAAGAGCAATACATCAAAGACGA
CGACGCCATGGAAGAGTACATGACGCAGTCGGCCCTGGAAGATGCGAGCCTGCACCTGAACGAAGAAGCACCGGGTATTT
CCGGCGAGGCGCTGGAGCGCCTGGTGAACGACTTCCGCATGGTCATGAAAACCCTCAAGCGTCTGTCGCGCCTGTACCCT
CAGGAGCTGACCGAGCACTTCATCTACCTGCCGGCCGTGAGCCTGGAGCAACTCTCCGATCACGCGGCCATGCAGGATTG
GCTGGCCCAATATGAAGTCCGCCTGCGCACCGTCGAGAAGTCCGGCCTGGTCTACAAGGCCAGCCTGCGTGAAGACCGTG
AACGTAATGTCTGGCTGCCAGAGGTCGAACTGATCTCCCACGGCCTGTCGAACTACGTCACCTTCAACCGCGACTTCTTC
GGTAGCAATGACTACAAGACCGTCGTTACCCTCGGCGCTCAACTGAGCTCCCTGCTGGACGAAGGCGCTTATATTCAGCG
TGGCGAACGCAAGAAGGCGGTGACCGAGTTCAAGGAAGCCCTGGACTGGCTGATGACCGAAAGCACCAAGCGCCACACCA
TCCAGCGATACAAAGGTCTGGGCGAGATGAACCCGGATCAGCTGTGGGAAACCACCATGGACCCAAGCGTGCGCCGTATG
CTCAAGGTCACGATTGAAGATGCCATCGGCGCCGACCAGATCTTCAACACCCTGATGGGGGATGCGGTCGAGCCTCGTCG
CGACTTCATCGAAAGCAACGCCCTGGCGGTATCCAATCTGGACTTCTGAATGTCCGGAAAAGCGCAACAGCAGTCTCGCC
TCAAAGAGTTGATCAGCCGTGGTCGTGAGCAGGGTTACCTGACTTACGCGGAGGTCAACGACCACCTGCCGGAGGATATT
TCAGATCCGGAACAGGTGGAAGACATCATCCGCATGATCAACGACATGGGGATCAACGTATTCGAGAGTGCTCCGGATGC
GGATGCCCTTTTGTTGGCCGAAGCCGATACCGACGAAGCAGCAGCTGAAGAAGCCGCCGCAGCGTTGGCGGCTGTGGAAA
CCGACATTGGTCGCACTACCGACCCCGTGCGTATGTACATGCGCGAAATGGGAACGGTAGAGCTGCTCACACGTGAAGGC
GAAATCGAAATCGCCAAGCGTATCGAAGAGGGCATCCGTGAAGTGATGGGCGCGATCGCGCACTTCCCTGGCACGGTTGA
GCACATCCTCTCCGAATACACTCGCGTCACCACCGAAGGTGGCCGCCTGTCCGACGTCCTGAGCGGTTACATCGACCCGG
ACGACGGCATTGCGCCGCCTGCCGCCGAAGTACCACCGCCTGTCGATGCCAAGGCTGCAAAAGCGGACGACGACACCGAC
GACGATGACGCCGAAGCCAGTGACGACGAAGAAGAAGCCGAAAGCGGTCCGGATCCGGTCATCGCAGCCCAGCGCTTTGG
CGCCGTTGCCGACCAGATGGAAATCACCCGCAAGGCGCTGAAGAAGCACGGTCGCGAACACAAGCAAGCCCTGGCCGAAA
TGCTGGCCCTGGCTGAACTGTTCATGCCGATCAAACTGGTTCCGAAGCAATTCGAAGGCCTGGTTGAACGTGTTCGTAGC
GCCCTGGATCGCCTGCGTCAGCAAGAGCGCGCGATCATGCAGCTCTGTGTTCGTGATGCCCGCATGCCACGCGCCGACTT
CCTGCGCCAGTTCCCTGGCAATGAAGTGGACGAAAGCTGGTCCGACGCGCTGGCCAAAGGCAAGGCCAAGTACGCCGAAG
CCATCGGCCGCCTGCAGCCGGACATCATCCGTTGCCAGCAGAAGCTGACCGCGCTCGAGACCGAGACCGGCCTGACGATC
GCCGAGATCAAGGACATCAACCGTCGCATGTCGATCGGCGAGGCCAAGGCCCGTCGCGCGAAGAAAGAGATGGTCGAAGC
CAACCTGCGTCTGGTGATCTCCATCGCCAAGAAGTACACCAACCGTGGCTTGCAATTCCTCGACCTGATCCAGGAAGGCA
ACATCGGTTTGATGAAAGCGGTAGACAAGTTCGAATACCGTCGCGGCTACAAATTCTCGACTTATGCCACCTGGTGGATC
CGTCAGGCGATCACTCGCTCGATCGCCGACCAGGCCCGCACCATCCGTATTCCGGTGCACATGATCGAGACGATCAACAA
GCTCAACCGTATTTCCCGGCAGATGTTGCAGGAAATGGGTCGCGAACCGACTCCGGAAGAGCTGGGCGAACGCATGGAAA
TGCCTGAGGACAAGATCCGCAAGGTATTGAAGATCGCTAAAGAGCCGATCTCCATGGAAACCCCGATCGGTGATGACGAA
GACTCCCATCTGGGTGACTTCATCGAAGACTCGACCATGCAGTCGCCAATCGATGTCGCCACCGTTGAGAGCCTTAAAGA
AGCGACTCGCGAAGTACTCTCCGGCCTCACTGCCCGTGAAGCCAAGGTACTGCGCATGCGCTTCGGCATCGACATGAATA
CCGACCACACCCTCGAGGAAGTCGGTAAGCAGTTCGACGTTACCCGTGAGCGGATTCGTCAGATCGAAGCCAAGGCGCTG
CGCAAGCTGCGCCACCCGACGCGAAGCGAGCACCTGCGCTCCTTCCTCGACGAGTAATTGACTAAGCCAGCCATACTCGC
CCTTGCTGATGGCAGCATTTTTCGCGGCGAAGCCATTGGAGCCGACGGTCAAACCGTTGGTGAGGTGGTGTTTAACACCG
CAATGACCGGCTATCAGGAAATCCTTACCGATCCTTCCTACGCCCAACAGATCGTTACCCTGACTTACCCGCATATCGGC
AATACCGGCACCACGCCGGAAGACGCCGAGTCCGATCGTGTCTGGTCGGCCGGTCTGGTGATTCGCGACCTGCCTCTGGT
TGCGAGCAACTGGCGTAACACCCTGTCCCTGTCCGACTACCTGAAAGCCAACAATGTCGTGGCGATCGCCGGTATCGACA
CCCGTCGCCTGACGCGCATCCTGCGCGAGAAAGGTGCGCAGAACGGCTGCATCATGGCCGGCGACAATATCTCCGACGAA
GCGGCGATTGCCGCTGCACGCGGCTTCCCGGGCCTGAAAGGCATGGATCTGGCGAAGGTCGTCAGTACCAAGGAAAGCTA

CGAGTGGCGCTCCAGTGTCTGGAACCTGAAGACCGACAGTCATCCGACCATCGAAGCTTCCGAGCTGCCTTACCACGTGG
TTGCCTACGACTACGGCGTCAAGCTGAACATCCTGCGCATGCTGGTCGAACGCGGTTGCCGCGTGACCGTGGTGCCTGCG
CAAACCCCGGCCAGCGAAGCTCTGGCGCTCAAGCCTGACGGTGTGTTCCTGTCCAACGGCCCTGGCGACCCCGAGCCTTG
CGATTACGCCATCCAGGCGATCAAGGACGTGCTGGAGACCGAGATTCCGGTCTTCGGTATCTGTCTGGGCCACCAACTGC
TGGCACTGGCCGCCGGCGCCAAGACAGTGAAGATGGGCCACGGCCACCACGGCGCCAACCACCCGGTCCAGGACCTGGAC
AGCGGTGTGGTGATGATCACCAGCCAGAACCACGGTTTTGCGGTGGACGAAGCCACCCTGCCGGGCAACGTGCGGGCGAT
CCACAAGTCGCTGTTCGACGGCACCCTGCAAGGCATCGAGCTGACCGACAAGAGCGCATTCAGCTTCCAGGGCCACCCTG
AAGCGAGCCCGGGCCCGAACGATGTGGCGCCGCTGTTCGATCGTTTCATCAACGAGATGGCCAAGCGACGCTGAATGAGT
AGCGGACGTATCGTTCAAATCATCGGCGCCGTTATCGACGTGGAATTTCCACGCGACAGCGTACCGAGCATCTACGACGC
CTTGAAGGTTCAAGGCGCCGAAACCACTCTGGAAGTTCAGCAGCAGCTGGGCGACGGCGTGGTACGTACCATTGCGATGG
GCTCCACCGAGGGCTTGAAGCGCGGTCTGGACGTCAACAACACTGGCGCAGCCATCTCCGTACCGGTCGGTAAAGCGACC
CTGGGCCGGATCATGGACGTACTGGGCAACCCGATCGACGAAGCTGGCCCGATCGGTGAAGAAGAGCGTTGGGGCATTCA
CCGTCCTGCGCCGACCTTCGCTGAACAAGCTGGCGGCAACGACCTGCTGGAAACCGGCATCAAGGTTATCGACCTGGTTT
GCCCGTTCGCCAAGGGCGGTAAAGTCGGTCTGTTCGGTGGTGCCGGTGTGGGCAAAACCGTAAACATGATGGAACTGATC
CGTAACATCGCCATCGAGCACAGCGGTTATTCCGTGTTCGCCGGTGTGGGTGAGCGTACTCGTGAGGGTAACGACTTCTA
CCACGAGATGAAGGATTCCAACGTTCTGGACAAAGTGGCACTGGTATACGGCCAGATGAACGAGCCGCCGGGAAACCGTC
TGCGCGTAGCTCTGACCGGCCTGACCATGGCCGAGAAGTTCCGTGACGAAGGTAACGACGTTCTGCTGTTCGTCGACAAC
ATCTATCGTTACACCCTGGCCGGTACCGAAGTATCCGCACTGCTGGGCCGTATGCCTTCGGCAGTAGGTTACCAGCCGAC
CCTGGCTGAAGAGATGGGCGTTCTGCAAGAACGTATCACTTCGACCAAGCAAGGCTCGATCACCTCGATCCAAGCGGTAT
ACGTACCTGCGGACGACTTGACCGACCCGTCGCCAGCGACCACCTTCGCCCACTTGGACGCCACCGTCGTTCTGTCCCGT
GACATCGCTTCCCTGGGTATCTACCCAGCGGTAGACCCACTGGACTCGACTTCCCGTCAGCTGGACCCGAACGTGATCGG
CACCGAGCACTACGAAACCGCTCGTGGCGTTCAGTACGTGCTGCAGCGCTACAAAGAGCTGAAGGACATCATTGCGATCC
TGGGTATGGACGAACTGTCCGAAGCCGACAAGCAACTGGTATCCCGCGCTCGTAAGATCCAGCGCTTCCTGTCGCAGCCG
TTCTTCGTGGCTGAAGTCTTCACTGGTTCTCCAGGCAAATACGTTTCCCTGAAAGACACCATCGCTGGCTTCAAAGGCAT
CCTCAACGGTGACTACGACCATCTGCCAGAACAAGCGTTCTACATGGTTGGTGGCATCGAAGAAGCGATCGAGAAAGCCA
AGAAACTGTAA

NCBI Reference Sequence: NZ_CM001559.1
Strain: 30-84
Chained genes: 16S rRNA, recA, gyrB, rpoD, carA, atpD
>NZ_CM001559_30_84
GAACTGAAGAGTTTGATCATGGCTCAGATTGAACGCTGGCGGCAGGCCTAACACATGCAAGTCGAGCGGTAGAGAGAAGC
TTGCTTCTCTTGAGAGCGGCGGACGGGTGAGTAATGCCTAGGAATCTGCCTGGTAGTGGGGGATAACGTCCGGAAACGGA
CGCTAATACCGCATACGTCCTACGGGAGAAAGCAGGGGACCTTCGGGCCTTGCGCTATCAGATGAGCCTAGGTCGGATTA
GCTAGTTGGTGAGGTAATGGCTCACCAAGGCGACGATCCGTAACTGGTCTGAGAGGATGATCAGTCACACTGGAACTGAG
ACACGGTCCAGACTCCTACGGGAGGCAGCAGTGGGGAATATTGGACAATGGGCGAAAGCCTGATCCAGCCATGCCGCGTG
TGTGAAGAAGGTCTTCGGATTGTAAAGCACTTTAAGTTGGGAGGAAGGGTACTTACCTAATACGTGAGTATTTTGACGTT
ACCGACAGAATAAGCACCGGCTAACTCTGTGCCAGCAGCCGCGGTAATACAGAGGGTGCAAGCGTTAATCGGAATTACTG
GGCGTAAAGCGCGCGTAGGTGGTTCGTTAAGTTGGATGTGAAATCCCCGGGCTCAACCTGGGAACTGCATCCAAAACTGG
CGAGCTAGAGTATGGTAGAGGGTGGTGGAATTTCCTGTGTAGCGGTGAAATGCGTAGATATAGGAAGGAACACCAGTGGC
GAAGGCGACCACCTGGACTGATACTGACACTGAGGTGCGAAAGCGTGGGGAGCAAACAGGATTAGATACCCTGGTAGTCC
ACGCCGTAAACGATGTCAACTAGCCGTTGGGAGCCTTGAGCTCTTAGTGGCGCAGCTAACGCATTAAGTTGACCGCCTGG
GGAGTACGGCCGCAAGGTTAAAACTCAAATGAATTGACGGGGGCCCGCACAAGCGGTGGAGCATGTGGTTTAATTCGAAG
CAACGCGAAGAACCTTACCAGGCCTTGACATCCAATGAACTTTCCAGAGATGGATTGGTGCCTTCGGGAACATTGAGACA
GGTGCTGCATGGCTGTCGTCAGCTCGTGTCGTGAGATGTTGGGTTAAGTCCCGTAACGAGCGCAACCCTTGTCCTTAGTT
ACCAGCACGTAATGGTGGGCACTCTAAGGAGACTGCCGGTGACAAACCGGAGGAAGGTGGGGATGACGTCAAGTCATCAT
GGCCCTTACGGCCTGGGCTACACACGTGCTACAATGGTCGGTACAGAGGGTTGCCAAGCCGCGAGGTGGAGCTAATCCCA
CAAAACCGATCGTAGTCCGGATCGCAGTCTGCAACTCGACTGCGTGAAGTCGGAATCGCTAGTAATCGCGAATCAGAATG
TCGCGGTGAATACGTTCCCGGGCCTTGTACACACCGCCCGTCACACCATGGGAGTGGGTTGCACCAGAAGTAGCTAGTCT
AACCTTCGGGAGGACGGTTACCACGGTGTGATTCATGACTGGGGTGAAGTCGTAACAAAGTAGCCGTAGGGGAACCTGCG
GCTGGATCACCTCCTTAAATGGACGACAACAAGAAGAAAGCCTTGGCTGCGGCCCTGGGTCAGATCGAACGTCAATTCGG
CAAGGGTGCCGTAATGCGTATGGGCGATCACGACCGCCAGGCGATCCCGGCCATTTCCACTGGCTCTCTGGGTCTGGACA
TCGCACTCGGCATCGGCGGCCTGCCAAAGGGCCGTATTGTTGAAATCTACGGTCCGGAATCGTCCGGTAAAACCACCCTG
ACCCTGTCCGTGATTGCCCAGGCACAGAAGATGGGCGCCACCTGCGCCTTCGTCGACGCCGAGCACGCACTGGACCCGGA
ATACGCCGGCAAGCTGGGGGTCAACGTTGACGACCTGCTGGTTTCCCAGCCGGACACCGGTGAACAGGCACTGGAAATCA
CCGATATGCTGGTGCGCTCCAATGCCATTGACGTGATCGTGATCGACTCCGTGGCGGCACTGGTGCCCAAGGCCGAGATC
GAAGGCGAGATGGGCGACATGCACGTGGGCCTGCAGGCCCGCCTGATGTCCCAGGCGCTGCGCAAGATCACCGGCAACAT
CAAGAACGCCAACTGCCTGGTGATCTTCATCAACCAGATCCGTATGAAAATCGGCGTGATGTTCGGCAGCCCGGAAACCA
CCACCGGTGGTAACGCGCTGAAGTTCTACGCTTCGGTTCGTCTGGATATCCGTCGTACTGGCGCGGTGAAGGAAGGTGAC
GAAGTCGTCGGTAGCGAAACCCGGGTCAAGATCGTCAAGAACAAGGTGGCTCCACCGTTCCGCCAGGCTGAATTCCAGAT
CCTGTACGGCAAGGGTATCTACCTGAACGGCGAGATCATCGATCTGGGCGTGCTGCACGGTTTCCTCGAGAAGTCCGGTG
CCTGGTACAGCTACCAGGGCAACAAGATCGGTCAGGGCAAGGCCAACTCGGCCAAGTTCCTGCAGGACAACCCGGAAATC
GGTAACGCCCTCGAGAAGCAGATTCGCGACAAGCTGCTGGCTCCGACTGCTGATGTCAAAGCTTCGCCGGTCAACGAGAC
CATCGATGACATGGCCGACGCGGATATCTGAATGAGCGAAGAAAACACGTACGACTCGAGCAGCATTAAAGTGCTGAAAG

GTTTGGATGCCGTACGCAAACGTCCCGGTATGTACATTGGTGACACCGACGATGGCAGCGGTCTGCACCATATGGTGTTC
GAGGTGGTCGATAACTCGATCGACGAAGCTCTGGCCGGCCATTGTGACGACATCAGCATCATCATCCACCCGGACGAATC
CATTACCGTGCGTGACAACGGTCGCGGCATCCCGGTAGACGTGCATAAAGAAGAAGGCGTTTCCGCAGCCGAGGTCATCA
TGACTGTGCTGCACGCCGGCGGTAAGTTCGACGATAACTCCTACAAAGTATCCGGCGGTCTGCACGGTGTGGGTGTGTCG
GTAGTAAACGCCCTGTCCGAAGAGCTGGTCTTGACCGTTCGCCGCAGTGGCAAGATCTGGGAACAGACCTACGTTCACGG
TGTGCCTCAGGCGCCTATGGCGATCGTCGGTGACAGCGAAACCACCGGTACCCAGATTCACTTCAAGGCTTCCAGCGAGA
CCTTCAAGAACATCCACTTCAGCTGGGACATCCTGGCCAAGCGGATTCGTGAACTGTCCTTCCTCAACTCCGGTGTCGGT
ATCGTTCTGAAGGACGAGCGCAGCGGCAAGGAAGAGCTGTTCAAGTACGAAGGCGGCTTGCGTGCGTTCGTTGAATACCT
GAACACCAACAAGACTGCGGTCAACCAGGTGTTCCACTTCAACGTGCAGCGTGAAGATGGCATCGGCGTGGAAATCGCCC
TGCAGTGGAACGACAGCTTCAACGAGAACCTGCAGTGCTTCACCAACAACATTCCGCAGCGCGACGGCGGCACCCACCTG
GTGGGCTTCCGTTCGGCGCTGACGCGTAACCTGAACAACTACATCGAGCAGGAAGGCCTGGCGAAGAAGCACAAGGTCGC
CACCACCGGTGACGATGCCCGCGAAGGCCTGACCGCGATCATTTCGGTCAAGGTGCCGGATCCGAAGTTCAGCTCCCAGA
CCAAAGACAAGCTGGTGTCTTCCGAAGTGAAGACCGCGGTCGAACAGGAAATGGGTAAGTACTTCTCCGACTTCCTGCTG
GAAAACCCGAACGAAGCCAAACTGGTGGTCGGCAAGATGCTCGACGCCGCCCGTGCCCGTGAAGCGGCGCGTAAAGCCCG
TGAGATGACCCGTCGTAAAGGCGCGCTGGATATCGCCGGCCTGCCGGGCAAACTGGCGGACTGCCAGGAAAAAGACCCTG
CCCTTTCCGAACTCTACCTGGTGGAAGGTGACTCTGCTGGCGGCTCCGCCAAGCAGGGACGCAACCGTAAGACCCAGGCG
ATCCTGCCGCTCAAGGGCAAGATCCTCAACGTCGAGAAAGCGCGCTTCGACAAGATGATTTCCTCGCAAGAGGTCGGCAC
CTTGATCACTGCACTCGGTTGCGGCATCGGCCGCGAAGAGTACAACATCGACAAGCTGCGTTATCACAACATCATCATCA
TGACCGACGCCGACGTCGACGGTTCGCACATCCGTACCCTGCTGCTGACCTTCTTCTTCCGTCAGTTGCCGGAGCTGATC
GAGCGTGGCTACATCTACATCGCTCAACCGCCGCTGTACAAGGTCAAGAAAGGCAAGCAAGAGCAATACATCAAAGACGA
CGACGCCATGGAAGAGTACATGACGCAGTCGGCCCTGGAGGATGCGAGCCTGCACCTGAACGAAGAAGCACCGGGTATTT
CCGGCGAGGCGCTGGAGCGTCTGGTGAACGACTTCCGCATGGTCATGAAGACCCTCAAGCGTCTGTCGCGCCTGTACCCT
CAAGAGCTGACCGAGCACTTCATCTACCTGCCGGCCGTGAGCCTGGAGCAACTCTCCGATCACGCGGCCATGCAGGATTG
GCTGGCCCAATATGAAGTCCGCCTGCGCACCGTCGAGAAGTCCGGCCTGGTCTACAAGGCCAGCCTGCGTGAAGACCGTG
AACGTAATGTGTGGCTGCCAGAGGTCGAACTGATCTCCCACGGCCTGTCGAACTACGTCACCTTCAACCGCGACTTCTTC
GGCAGCAATGACTACAAGACCGTCGTCACCCTCGGTGCTCAACTGAGCTCCCTGCTGGACGAAGGCGCTTATATTCAGCG
CGGCGAGCGCAAGAAGGCGGTGACCGAGTTCAAGGAAGCCCTGGACTGGCTGATGACCGAAAGCACCAAGCGCCACACCA
TCCAGCGATACAAAGGTCTGGGCGAGATGAACCCGGATCAGTTGTGGGAAACCACCATGGACCCAAGCGTGCGCCGCATG
CTCAAGGTCACCATCGAAGACGCCATCGGCGCCGACCAGATCTTCAACACCCTGATGGGTGATGCGGTCGAGCCTCGTCG
TGACTTCATCGAAAGCAACGCCCTGGCGGTATCCAACCTGGACTTCTGAATGTCCGGAAAAGCGCAACAGCAGTCTCGCC
TCAAAGAGTTGATCAGCCGTGGTCGTGAGCAGGGTTACCTGACTTACGCGGAGGTCAACGACCACCTGCCGGAGGATATT
TCAGATCCGGAACAGGTGGAAGACATCATCCGCATGATCAACGACATGGGGATCAACGTATTCGAGAGTGCTCCGGATGC
GGATGCCCTTTTGTTGGCCGAAGCCGATACCGACGAAGCAGCAGCTGAAGAAGCCGCCGCAGCGTTGGCGGCTGTGGAAA
CCGACATTGGTCGCACTACCGACCCAGTGCGTATGTACATGCGCGAAATGGGTACGGTAGAGCTGCTCACACGTGAAGGC
GAAATCGAAATCGCCAAGCGTATCGAAGAGGGCATCCGTGAAGTGATGGGCGCGATCGCGCACTTCCCTGGCACGGTTGA
GCACATCCTCTCCGAATACACTCGCGTCACCACCGAAGGTGGTCGCCTGTCCGACGTCCTGAGCGGTTACATCGACCCGG
ACGACGGCATCGCGCCGCCTGCCGCCGAAGTACCACCGCCTGTCGATGCCAAGGCCGCGAAAGCGGACGACGACACCGAC
GACGATGACGCCGAAGCCAGTGACGACGAAGAAGAAGCCGAAAGCGGTCCGGATCCGGTCATCGCAGCCCAGCGCTTTGG
CGCCGTTGCCGACCAGATGGAAATCACCCGCAAGGCGCTGAAAAAGCACGGTCGTGAACACAAGCAAGCCCTGGCTGAAA
TGCTGGCCCTGGCTGAACTGTTCATGCCGATCAAACTGGTTCCGAAGCAATTCGAAGGCCTGGTTGAACGCGTTCGTAGC
GCCCTGGATCGCCTGCGTCAGCAAGAGCGCGCGATCATGCAACTCTGTGTTCGTGATGCTCGCATGCCACGCGCCGACTT
CCTGCGCCAGTTCCCTGGCAATGAAGTGGACGAAAGCTGGTCCGACGCACTGGCCAAAGGCAAGGCCAAGTACGCCGAAG
CCATCGGTCGCCTGCAACCGGACATCATCCGTTGCCAGCAGAAGCTGACCGCGCTCGAGACCGAGACCGGCCTGACGATC
GCCGAGATCAAGGACATCAACCGTCGCATGTCGATCGGCGAAGCCAAGGCCCGTCGCGCGAAGAAAGAGATGGTCGAAGC
CAACTTGCGTCTGGTGATCTCCATCGCCAAGAAGTACACCAACCGTGGCCTGCAGTTCCTCGACCTGATCCAGGAAGGCA
ACATCGGTTTGATGAAAGCGGTAGACAAGTTCGAATACCGTCGCGGCTACAAGTTCTCGACTTATGCGACCTGGTGGATC
CGTCAGGCGATCACTCGCTCGATCGCCGACCAGGCCCGTACCATCCGTATTCCGGTGCACATGATCGAGACGATCAACAA
GCTCAACCGTATTTCCCGGCAGATGCTGCAGGAAATGGGTCGCGAACCGACCCCGGAAGAGCTGGGCGAACGCATGGAAA
TGCCTGAGGACAAGATCCGCAAGGTATTGAAGATCGCTAAAGAGCCGATCTCCATGGAAACCCCGATCGGTGATGACGAA
GACTCCCATCTGGGCGACTTCATCGAAGACTCGACCATGCAGTCGCCAATCGATGTCGCCACCGTTGAGAGCCTCAAGGA
AGCGACTCGCGAAGTACTCTCCGGCCTCACTGCCCGTGAAGCCAAGGTACTGCGCATGCGCTTCGGCATCGACATGAATA
CCGACCACACCCTCGAGGAAGTCGGTAAGCAGTTCGACGTTACCCGTGAGCGGATTCGCCAGATCGAAGCCAAGGCGCTG
CGCAAGCTGCGCCACCCGACGAGAAGCGAGCACCTGCGCTCCTTCCTCGACGAGTAATTGACTAAGCCAGCCATACTCGC
CCTTGCTGATGGCAGCATTTTTCGCGGCGAAGCCATTGGAGCCGACGGTCAAACCGTTGGTGAGGTGGTGTTTAACACCG
CAATGACCGGCTATCAGGAAATCCTTACCGATCCTTCCTACGCCCAACAGATCGTTACCCTGACTTACCCGCATATCGGC
AATACCGGCACCACGCCGGAAGACGCCGAGTCCGATCGTGTCTGGTCGGCCGGTCTGGTGATTCGCGACCTGCCACTGTT
TGCGAGCAACTGGCGTAACACCCTGTCCCTGTCCGATTACCTGAAAGCCAACAACGTTGTGGCGATCGCCGGTATCGACA
CCCGTCGCCTGACGCGCATCCTGCGCGAGAAAGGCGCGCAGAACGGCTGCATCATGGCCGGCGACAACATTTCCGACGAA
GCGGCGATTGCCGCAGCGCGCGGCTTCCCTGGCCTGAAAGGCATGGATCTGGCGAAGGTCGTCAGCACCAAGGAAAGCTA
CGAGTGGCGCTCCAGCGTCTGGAGCCTGAAGACCGACAGTCATCCGACCATCGAGGCTTCCGAGCTGCCGTACCACGTGG
TCGCCTACGATTACGGCGTCAAGCTGAACATCCTGCGCATGCTGGTCGAGCGCGGTTGCCGCGTGACCGTAGTGCCTGCG
CAAACTCCGGCCAGCGACGTCCTGGCACTCAAGCCTGACGGTGTGTTCCTGTCCAACGGCCCTGGCGACCCCGAGCCTTG
CGATTACGCCATCCAGGCGATCAAGGATGTGCTGGAAACCGAGATCCCGGTCTTCGGTATCTGCCTGGGCCACCAACTGC
TGGCACTGGCCTCTGGCGCCAAGACGGTGAAGATGGGCCACGGCCACCACGGTGCCAACCACCCGGTCCAGGACCTGGAC

AGCGGTGTAGTGATGATCACCAGCCAGAACCACGGTTTTGCGGTGGACGAAGCCACCCTGCCAGGCAACGTGCGGGCGAT
CCACAAATCGCTGTTCGACGGCACCCTGCAAGGCATCGAACGTACCGACAAGAGCGCATTCAGCTTCCAGGGCCACCCTG
AAGCGAGCCCGGGCCCGAACGATGTGGCGCCGCTGTTCGATCGTTTCATCAACGAGATGGCCAAGCGACGCTGAATGAGT
AGCGGACGTATCGTTCAAATCATCGGCGCCGTTATCGACGTGGAATTTCCACGCGACAGCGTACCGAGCATCTACGACGC
CTTGAAGGTTCAAGGCGCCGAAACCACTCTGGAAGTTCAGCAGCAGCTGGGCGACGGCGTGGTACGTACCATTGCGATGG
GCTCCACCGAGGGCTTGAAGCGCGGTCTGGACGTCAACAACACTGGCGCAGCCATCTCCGTACCGGTCGGTAAAGCGACC
CTGGGCCGGATCATGGACGTACTGGGCAACCCGATCGACGAAGCTGGTCCGATCGGCGAAGAAGAGCGTTGGGGCATTCA
CCGTCCTGCGCCGACCTTCGCTGAACAAGCTGGCGGCAACGACCTGCTGGAAACCGGCATCAAGGTTATCGACCTGGTTT
GCCCGTTCGCCAAGGGCGGTAAAGTCGGTCTGTTCGGTGGTGCCGGTGTGGGCAAAACCGTAAACATGATGGAACTGATC
CGTAACATCGCCATCGAGCACAGCGGTTATTCCGTGTTCGCCGGTGTGGGTGAGCGTACTCGTGAGGGTAACGACTTCTA
CCACGAGATGAAGGATTCCAACGTTCTGGACAAAGTGGCACTGGTATACGGCCAGATGAACGAGCCGCCGGGAAACCGTC
TGCGCGTAGCTCTGACCGGCCTGACCATGGCCGAGAAGTTCCGTGACGAAGGTAACGACGTTCTGCTGTTCGTCGACAAC
ATCTATCGTTACACCCTGGCCGGTACCGAAGTATCCGCACTGCTGGGCCGTATGCCTTCCGCAGTAGGTTACCAGCCGAC
CCTGGCTGAAGAGATGGGCGTTCTGCAAGAACGTATCACTTCGACCAAGCAAGGCTCGATCACCTCGATCCAAGCGGTAT
ACGTACCTGCGGACGACTTGACCGACCCGTCGCCAGCGACCACCTTCGCCCACTTGGACGCCACCGTCGTACTGTCCCGT
GACATCGCTTCCCTGGGTATCTACCCAGCGGTAGATCCACTGGACTCGACTTCCCGTCAGCTGGACCCGAACGTGATCGG
CAACGAGCACTATGAAACCGCTCGCGGCGTTCAGTACGTGCTGCAGCGCTACAAAGAGCTGAAGGACATCATTGCGATCC
TGGGTATGGACGAACTGTCCGAAGCCGACAAGCAACTGGTATCCCGCGCTCGTAAGATCCAGCGCTTCCTGTCGCAGCCG
TTCTTCGTGGCTGAAGTCTTCACTGGTTCTCCAGGCAAATACGTTTCCCTGAAAGACACCATCGCTGGCTTCAAAGGCAT
CCTCAACGGTGACTACGACCACCTGCCAGAACAAGCGTTCTACATGGTCGGCGGCATCGAAGAAGCGATCGAGAAAGCCA
AGAAACTGTAA

NCBI Reference Sequence: NZ_CP027712.1
Strain: DSM50083
Chained genes: 16S rRNA, recA, gyrB, rpoD, carA, atpD
>NZ_CP027712_DSM50083
GAACTGAAGAGTTTGATCATGGCTCAGATTGAACGCTGGCGGCAGGCCTAACACATGCAAGTCGAGCGGTAGAGAGGTGC
TTGCACCTCTTGAGAGCGGCGGACGGGTGAGTAATGCCTAGGAATCTGCCTGGTAGTGGGGGATAACGTTCGGAAACGGA
CGCTAATACCGCATACGTCCTACGGGAGAAAGCAGGGGACCTTCGGGCCTTGCGCTATCAGATGAGCCTAGGTCGGATTA
GCTAGTTGGTGAGGTAATGGCTCACCAAGGCGACGATCCGTAACTGGTCTGAGAGGATGATCAGTCACACTGGAACTGAG
ACACGGTCCAGACTCCTACGGGAGGCAGCAGTGGGGAATATTGGACAATGGGCGAAAGCCTGATCCAGCCATGCCGCGTG
TGTGAAGAAGGTCTTCGGATTGTAAAGCACTTTAAGTTGGGAGGAAGGGTACTTACCTAATACGTGAGTATTTTGACGTT
ACCGACAGAATAAGCACCGGCTAACTCTGTGCCAGCAGCCGCGGTAATACAGAGGGTGCAAGCGTTAATCGGAATTACTG
GGCGTAAAGCGCGCGTAGGTGGTTCGTTAAGTTGGATGTGAAATCCCCGGGCTCAACCTGGGAACTGCATCCAAAACTGG
CGAGCTAGAGTATGGTAGAGGGTGGTGGAATTTCCTGTGTAGCGGTGAAATGCGTAGATATAGGAAGGAACACCAGTGGC
GAAGGCGACCACCTGGACTGATACTGACACTGAGGTGCGAAAGCGTGGGGAGCAAACAGGATTAGATACCCTGGTAGTCC
ACGCCGTAAACGATGTCAACTAGCCGTTGGGAGCCTTGAGCTCTTAGTGGCGCAGCTAACGCATTAAGTTGACCGCCTGG
GGAGTACGGCCGCAAGGTTAAAACTCAAATGAATTGACGGGGGCCCGCACAAGCGGTGGAGCATGTGGTTTAATTCGAAG
CAACGCGAAGAACCTTACCAGGCCTTGACATCCAATGAACTTTCCAGAGATGGATTGGTGCCTTCGGGAACATTGAGACA
GGTGCTGCATGGCTGTCGTCAGCTCGTGTCGTGAGATGTTGGGTTAAGTCCCGTAACGAGCGCAACCCTTGTCCTTAGTT
ACCAGCACGTAATGGTGGGCACTCTAAGGAGACTGCCGGTGACAAACCGGAGGAAGGTGGGGATGACGTCAAGTCATCAT
GGCCCTTACGGCCTGGGCTACACACGTGCTACAATGGTCGGTACAGAGGGTTGCCAAGCCGCGAGGTGGAGCTAATCCCA
CAAAACCGATCGTAGTCCGGATCGCAGTCTGCAACTCGACTGCGTGAAGTCGGAATCGCTAGTAATCGCGAATCAGAATG
TCGCGGTGAATACGTTCCCGGGCCTTGTACACACCGCCCGTCACACCATGGGAGTGGGTTGCACCAGAAGTAGCTAGTCT
AACCTTCGGGAGGACGGTTACCACGGTGTGATTCATGACTGGGGTGAAGTCGTAACAAGGTAGCCGTAGGGGAACCTGCG
GCTGGATCACCTCCTTAAATGGACGACAACAAGAAGAAAGCCTTGGCTGCGGCCCTGGGTCAGATCGAACGTCAATTCGG
CAAGGGTGCCGTAATGCGTATGGGCGATCACGACCGCCAGGCGATCCCGGCCATTTCCACTGGCTCTCTGGGTCTGGACA
TCGCACTCGGCATCGGCGGCCTGCCAAAGGGCCGTATTGTCGAAATCTACGGTCCGGAATCGTCCGGTAAAACCACCCTG
ACCTTGTCCGTGATTGCCCAGGCACAGAAGATGGGCGCCACTTGCGCCTTCGTCGACGCCGAGCACGCACTGGACCCGGA
ATACGCCGGCAAGCTGGGGGTCAACGTTGACGACCTGCTGGTTTCCCAGCCGGACACCGGTGAACAGGCACTGGAAATCA
CCGACATGTTGGTGCGCTCCAATGCCATCGACGTGATCGTGATCGACTCCGTGGCAGCACTGGTGCCCAAGGCCGAGATC
GAAGGCGAGATGGGCGACATGCACGTGGGCCTGCAGGCCCGCCTGATGTCCCAGGCGCTGCGCAAGATCACCGGCAACAT
CAAGAACGCCAACTGCCTGGTGATCTTCATCAACCAGATCCGTATGAAAATCGGCGTGATGTTCGGCAGCCCGGAAACCA
CCACCGGTGGTAACGCGCTGAAGTTCTACGCTTCGGTTCGCCTGGATATCCGTCGTACTGGCGCGGTGAAGGAAGGTGAC
GAAGTCGTCGGTAGCGAAACCCGGGTCAAGATCGTCAAGAACAAGGTGGCTCCACCGTTCCGCCAGGCTGAATTCCAGAT
CCTGTACGGCAAGGGTATCTACCTGAACGGCGAGATCATCGATCTGGGCGTGCTGCACGGTTTCCTCGAGAAGTCTGGTG
CCTGGTACAGCTACCAGGGCAACAAGATCGGTCAGGGCAAGGCCAACTCGGCCAAGTTCCTGCAGGACAACCCGGAAATC
GGTAATGCCCTCGAGAAGCAGATTCGCGACAAGCTGCTGGCTCCGACCGCTGATGTCAAAGCTTCGCCGGTCAACGAGAC
CATCGATGACATGGCCGACGCGGATATCTGAATGAGCGAAGAAAACACGTACGACTCGAGCAGCATTAAAGTGCTGAAAG
GTTTGGATGCCGTACGCAAACGTCCCGGTATGTACATTGGTGACACCGACGATGGCAGCGGTCTGCACCATATGGTGTTC
GAGGTGGTCGATAACTCGATCGACGAAGCTCTGGCCGGCCATTGCGACGACATCAGCATCATCATCCACCCGGACGAATC
CATTACCGTGCGCGACAACGGTCGCGGCATCCCGGTAGACGTGCATAAAGAAGAAGGCGTTTCCGCAGCCGAGGTCATCA
TGACCGTGCTGCACGCCGGCGGTAAGTTCGACGATAACTCCTACAAAGTATCCGGCGGTCTGCACGGTGTGGGTGTGTCG
GTAGTGAACGCCCTGTCCGAAGAACTGGTCTTGACCGTTCGCCGCAGTGGCAAGATCTGGGAACAGACCTACGTTCACGG

TGTGCCTCAGGCGCCTATGGCTATCGTCGGTGACAGCGAAACCACCGGTACCCAGATTCACTTCAAGGCTTCCAGCGAAA
CCTTCAAGAACATCCACTTCAGCTGGGACATCCTGGCCAAGCGGATTCGTGAACTGTCCTTCCTCAACTCCGGTGTTGGT
ATCGTTCTGAAGGACGAGCGCAGCGGCAAGGAAGAACTGTTCAAGTACGAAGGCGGCTTGCGTGCGTTCGTTGAATACCT
GAATACCAACAAGACTGCGGTCAACCAGGTGTTCCACTTCAATGTGCAGCGTGAAGACGGCATTGGCGTGGAAATCGCAC
TGCAGTGGAACGACAGCTTCAACGAAAACCTGCAGTGCTTCACCAACAACATTCCGCAGCGCGACGGCGGCACCCACCTG
GTGGGCTTCCGTTCGGCACTGACGCGTAACCTGAACAACTACATCGAACAGGAAGGCCTGGCGAAGAAGCACAAGGTCGC
CACCACCGGTGACGATGCCCGCGAAGGCCTGACCGCGATCATTTCGGTCAAGGTGCCGGATCCGAAGTTCAGCTCCCAGA
CCAAAGACAAGCTGGTGTCTTCCGAAGTGAAGACCGCGGTCGAACAGGAAATGGGCAAGTACTTCTCCGACTTCCTGCTG
GAAAACCCGAACGAAGCCAAGCTGGTGGTCGGCAAGATGCTCGACGCCGCCCGTGCCCGTGAAGCGGCGCGTAAAGCCCG
TGAGATGACCCGTCGTAAAGGCGCGCTGGATATCGCCGGCCTGCCGGGCAAACTGGCGGACTGCCAGGAAAAAGACCCTG
CTCTTTCCGAACTCTACCTGGTGGAAGGTGACTCTGCTGGCGGCTCCGCCAAGCAGGGACGCAACCGTAAGACCCAGGCG
ATCCTGCCGCTCAAGGGCAAGATCCTCAACGTCGAGAAAGCGCGCTTCGACAAGATGATTTCCTCGCAAGAGGTCGGCAC
CTTGATCACTGCACTCGGTTGCGGCATCGGCCGCGAAGAGTACAACATCGACAAGCTGCGTTATCACAACATCATCATCA
TGACCGACGCCGACGTCGACGGTTCGCACATCCGTACCCTCCTGCTGACCTTCTTTTTCCGTCAGTTGCCGGAGCTGATC
GAGCGTGGCTACATCTACATCGCTCAACCGCCGCTGTACAAGGTCAAGAAAGGCAAGCAAGAGCAATACATCAAAGACGA
CGACGCCATGGAAGAGTACATGACGCAGTCGGCCCTGGAAGATGCGAGCCTGCACCTGAACGAAGAAGCACCGGGTATTT
CCGGCGAGGCGCTGGAGCGCCTGGTGAACGACTTCCGCATGGTCATGAAGACCCTCAAGCGTCTGTCGCGCCTGTACCCT
CAAGAGCTGACCGAGCACTTCATCTACCTGCCGGCCGTGAGCCTGGAGCAACTCTCCGATCACGCGGCGATGCAGGATTG
GCTGGCCCAGTATGAAGTCCGCCTGCGCACCGTCGAGAAGTCCGGCCTGGTCTACAAGGCCAGTCTGCGTGAAGACCGTG
AACGTAATGTCTGGCTGCCAGAGGTCGAACTGATCTCCCACGGCCTGTCGAACTACGTCACCTTCAACCGCGATTTCTTC
GGCAGTAATGACTACAAGACCGTCGTCACCCTCGGGGCTCAACTGAGCTCCCTGCTGGACGAAGGCGCTTATATTCAGCG
CGGCGAGCGCAAGAAGGCAGTGACCGAGTTCAAGGAAGCCCTGGACTGGCTGATGACCGAAAGCACCAAACGCCACACCA
TCCAGCGATATAAAGGTCTGGGTGAGATGAACCCGGACCAGCTGTGGGAAACCACCATGGACCCAAGCGTGCGCCGCATG
CTCAAGGTCACCATCGAAGACGCCATCGGCGCCGACCAGATCTTCAACACCCTGATGGGTGATGCGGTCGAGCCTCGTCG
CGACTTCATCGAAAGCAACGCCCTGGCGGTATCCAACCTGGACTTCTGAATGTCCGGAAAAGCGCAACAGCAGTCTCGCC
TCAAAGAGTTGATCAGCCGTGGTCGTGAGCAGGGTTACCTGACTTACGCGGAGGTCAACGACCACCTGCCGGAGGATATT
TCAGATCCGGAACAGGTGGAAGACATCATCCGCATGATCAACGACATGGGGATCAACGTATTCGAGAGTGCTCCGGATGC
GGATGCCCTTTTGTTGGCCGAAGCCGATACCGACGAAGCAGCAGCTGAAGAAGCCGCCGCAGCGTTGGCGGCAGTGGAAA
CCGACATTGGTCGCACTACCGACCCAGTGCGTATGTACATGCGCGAAATGGGTACGGTAGAGCTGCTCACACGTGAAGGC
GAAATCGAAATCGCCAAGCGTATCGAAGAGGGCATCCGTGAAGTGATGGGCGCGATCGCGCACTTCCCTGGCACGGTTGA
GCACATCCTCTCCGAATACACTCGCGTCACCACCGAAGGTGGTCGCCTGTCCGACGTCCTGAGCGGTTACATCGACCCGG
ACGACGGCATCGCGCCGCCTGCCGCCGAAGTACCACCGCCTGTCGATGCCAAGGCCGCGAAAGCGGACGACGACACCGAC
GACGATGACGCCGAAGCCAGTGACGACGAAGAAGAAGCCGAAAGCGGTCCGGATCCGGTCATCGCAGCCCAGCGCTTTGG
CGCCGTTGCCGACCAGATGGAAATCACCCGCAAGGCGCTGAAAAAGCACGGTCGTGAACACAAGCAAGCCCTGGCTGAAA
TGCTGGCCCTGGCTGAACTGTTCATGCCGATCAAACTGGTTCCGAAGCAATTCGAAGGCCTGGTTGAACGCGTTCGTAGC
GCCCTGGATCGCCTGCGTCAGCAAGAGCGCGCGATCATGCAGCTCTGTGTTCGTGATGCTCGCATGCCACGCGCCGACTT
CCTGCGCCAGTTCCCTGGCAATGAAGTGGACGAAAGCTGGTCCGACGCACTGGCCAAAGGCAAGGCCAAGTACGCCGAAG
CCATCGGTCGCCTGCAGCCGGACATCATTCGTTGCCAGCAGAAGCTGACCGCGCTCGAGACCGAGACCGGCCTGACGATC
GCCGAGATCAAGGACATCAACCGTCGCATGTCGATCGGCGAGGCCAAGGCCCGTCGCGCGAAGAAAGAGATGGTCGAAGC
CAACTTGCGTCTGGTGATCTCCATCGCCAAGAAGTACACCAACCGTGGCCTGCAGTTCCTCGACCTGATCCAGGAAGGCA
ACATCGGCTTGATGAAAGCGGTAGACAAGTTCGAATACCGTCGCGGCTACAAGTTCTCGACTTATGCCACCTGGTGGATC
CGTCAGGCGATCACTCGCTCGATCGCCGACCAGGCCCGCACCATCCGTATTCCGGTGCACATGATCGAGACGATCAACAA
GCTCAACCGTATTTCCCGGCAGATGCTGCAGGAAATGGGCCGCGAACCGACCCCGGAAGAGCTGGGCGAACGCATGGAAA
TGCCTGAGGACAAGATCCGCAAGGTATTGAAGATCGCTAAAGAGCCGATCTCCATGGAAACCCCGATCGGTGATGACGAA
GACTCCCATCTGGGCGACTTCATCGAAGACTCGACCATGCAGTCGCCAATCGATGTCGCTACCGTTGAGAGCCTCAAGGA
AGCGACTCGCGAAGTACTCTCCGGCCTCACTGCCCGTGAAGCCAAGGTACTGCGCATGCGCTTCGGTATCGACATGAATA
CCGACCATACCCTCGAGGAAGTCGGTAAGCAGTTCGACGTTACCCGTGAGCGGATTCGCCAGATCGAAGCCAAGGCGCTG
CGCAAGCTGCGCCACCCGACGAGAAGCGAGCACCTGCGCTCCTTCCTCGACGAGTAATTGACTAAGCCAGCCATACTCGC
CCTTGCTGATGGCAGCATTTTTCGCGGCGAAGCCATTGGAGCCGACGGTCAAACCGTTGGTGAGGTGGTGTTTAACACCG
CAATGACCGGCTATCAGGAAATCCTTACCGATCCTTCCTACGCCCAACAGATCGTTACCCTGACTTACCCGCATATCGGC
AATACCGGCACCACGCCGGAAGACGCCGAGTCCGATCGTGTCTGGTCGGCCGGTCTGGTGATTCGCGACCTGCCACTGGT
TGCGAGCAACTGGCGTAACACCCTGTCCCTGTCCGATTACCTGAAAGCCAACAATGTTGTGGCGATCGCCGGTATCGACA
CCCGTCGCCTGACGCGCATCCTGCGCGAGAAAGGCGCGCAGAACGGCTGCATCATGGCCGGTGACAATATCTCCGACGAA
GCGGCGATTGCCGCAGCGCGCGGCTTCCCTGGCCTGAAAGGCATGGATCTGGCGAAGGTCGTCAGCACCAAGGAAAGCTA
CGAGTGGCGCTCCAGCGTCTGGAGCCTGAAGACCGACAGTCATCCGACCATCGAGGCTTCCGAGCTGCCGTACCACGTGG
TTGCCTACGACTACGGCGTCAAGCTGAACATCCTGCGCATGCTGGTCGAGCGCGGTTGCCGCGTGACCGTAGTGCCTGCG
CAAACTCCGGCCAGCGACGTCCTGGCACTCAAGCCTGATGGCGTGTTCCTGTCCAACGGCCCTGGCGACCCCGAGCCTTG
CGATTACGCCATCCAGGCGATCAAGGACGTGCTGGAAACCGAGATCCCGGTCTTCGGTATCTGCCTGGGCCACCAACTGC
TGGCACTGGCCTCCGGCGCCAAGACGGTGAAGATGGGCCACGGCCACCACGGTGCCAACCACCCGGTCCAGGACCTGGAC
AGCGGTGTAGTGATGATCACCAGCCAGAACCACGGTTTTGCGGTGGACGAAGCCACCCTGCCAGGCAACGTGCGGGCGAT
CCACAAATCGCTGTTCGACGGTACCCTGCAAGGCATCGAACGTACCGACAAGAGCGCATTCAGCTTCCAGGGCCACCCTG
AAGCGAGCCCGGGCCCGAACGATGTGGCGCCGCTGTTCGATCGTTTCATCAACGAGATGGCCAAGCGACGCTGAATGAGT
AGCGGACGTATCGTTCAAATCATCGGCGCCGTTATCGACGTGGAATTTCCACGCGACAGCGTACCGAGCATCTACGACGC
CTTGAAGGTTCAAGGCGCCGAAACCACTCTGGAAGTTCAGCAGCAGCTGGGCGACGGCGTGGTACGTACCATTGCGATGG

GCTCCACCGAGGGCTTGAAGCGCGGTCTGGACGTCAACAACACTGGCGCAGCCATCTCCGTACCGGTCGGTAAAGCGACC
CTGGGCCGGATCATGGACGTACTGGGCAACCCGATCGACGAAGCTGGCCCGATCGGCGAAGAAGAGCGTTGGGGCATTCA
CCGTCCTGCGCCGTCCTTCGCTGAACAAGCTGGTGGCAACGACCTCCTGGAAACCGGCATCAAGGTTATCGACCTGGTTT
GCCCGTTCGCCAAGGGCGGTAAAGTCGGTCTGTTCGGTGGTGCCGGTGTGGGCAAGACCGTAAACATGATGGAACTGATC
CGTAACATCGCCATCGAGCACAGCGGTTATTCCGTGTTCGCCGGTGTGGGTGAGCGTACTCGTGAGGGTAACGACTTCTA
CCACGAGATGAAGGATTCCAACGTTCTGGACAAAGTGGCACTGGTATACGGCCAGATGAACGAGCCGCCGGGAAACCGTC
TGCGCGTAGCTCTGACCGGCCTGACCATGGCCGAGAAGTTCCGTGACGAAGGTAACGACGTTCTGCTGTTCGTCGACAAC
ATCTATCGTTACACCCTGGCCGGTACCGAAGTATCCGCACTGCTGGGCCGTATGCCTTCCGCAGTAGGTTACCAGCCGAC
CCTGGCTGAAGAGATGGGCGTTCTGCAAGAACGTATCACTTCGACCAAGCAAGGCTCGATCACCTCGATCCAAGCGGTAT
ACGTACCTGCGGACGACTTGACCGACCCGTCGCCAGCGACCACCTTCGCCCACTTGGACGCCACAGTCGTACTGTCCCGT
GACATCGCTTCCCTGGGTATCTACCCAGCGGTAGATCCACTGGACTCGACTTCCCGTCAGCTGGACCCGAACGTGATCGG
CAACGAGCACTACGAAACCGCTCGCGGCGTTCAGTACGTGCTGCAGCGCTACAAAGAGCTGAAGGACATCATTGCGATCC
TGGGTATGGACGAACTGTCCGAAGCCGACAAGCAACTGGTATCCCGCGCTCGTAAGATCCAGCGCTTCCTGTCGCAGCCG
TTCTTCGTGGCTGAAGTCTTCACTGGTTCTCCAGGCAAATACGTTTCCCTGAAAGACACCATCGCTGGCTTCAAAGGCAT
CCTCAACGGTGACTACGACCACCTGCCAGAACAAGCGTTCTACATGGTCGGCGGCATCGAAGAAGCGATCGAGAAAGCCA
AGAAACTGTAA

NCBI Reference Sequence: NZ_CP027717.1
Strain: PCM2210
Chained genes: 16S rRNA, recA, gyrB, rpoD, carA, atpD
>NZ_CP027717_PCM2210
GAACTGAAGAGTTTGATCATGGCTCAGATTGAACGCTGGCGGCAGGCCTAACACATGCAAGTCGAGCGGTAGAGAGAAGC
TTGCTTCTCTTGAGAGCGGCGGACGGGTGAGTAATGCCTAGGAATCTGCCTGGTAGTGGGGGATAACGTCCGGAAACGGA
CGCTAATACCGCATACGTCCTACGGGAGAAAGCAGGGGACCTTCGGGCCTTGCGCTATCAGATGAGCCTAGGTCGGATTA
GCTAGTTGGTGAGGTAATGGCTCACCAAGGCGACGATCCGTAACTGGTCTGAGAGGATGATCAGTCACACTGGAACTGAG
ACACGGTCCAGACTCCTACGGGAGGCAGCAGTGGGGAATATTGGACAATGGGCGAAAGCCTGATCCAGCCATGCCGCGTG
TGTGAAGAAGGTCTTCGGATTGTAAAGCACTTTAAGTTGGGAGGAAGGGTACTTACCTAATACGTGAGTATTTTGACGTT
ACCGACAGAATAAGCACCGGCTAACTCTGTGCCAGCAGCCGCGGTAATACAGAGGGTGCAAGCGTTAATCGGAATTACTG
GGCGTAAAGCGCGCGTAGGTGGTTCGTTAAGTTGGATGTGAAATCCCCGGGCTCAACCTGGGAACTGCATCCAAAACTGG
CGAGCTAGAGTATGGTAGAGGGTGGTGGAATTTCCTGTGTAGCGGTGAAATGCGTAGATATAGGAAGGAACACCAGTGGC
GAAGGCGACCACCTGGACTGATACTGACACTGAGGTGCGAAAGCGTGGGGAGCAAACAGGATTAGATACCCTGGTAGTCC
ACGCCGTAAACGATGTCAACTAGCCGTTGGGAGCCTTGAGCTCTTAGTGGCGCAGCTAACGCATTAAGTTGACCGCCTGG
GGAGTACGGCCGCAAGGTTAAAACTCAAATGAATTGACGGGGGCCCGCACAAGCGGTGGAGCATGTGGTTTAATTCGAAG
CAACGCGAAGAACCTTACCAGGCCTTGACATCCAATGAACTTTCCAGAGATGGATTGGTGCCTTCGGGAACATTGAGACA
GGTGCTGCATGGCTGTCGTCAGCTCGTGTCGTGAGATGTTGGGTTAAGTCCCGTAACGAGCGCAACCCTTGTCCTTAGTT
ACCAGCACGTTATGGTGGGCACTCTAAGGAGACTGCCGGTGACAAACCGGAGGAAGGTGGGGATGACGTCAAGTCATCAT
GGCCCTTACGGCCTGGGCTACACACGTGCTACAATGGTCGGTACAGAGGGTTGCCAAGCCGCGAGGTGGAGCTAATCCCA
TAAAACCGATCGTAGTCCGGATCGCAGTCTGCAACTCGACTGCGTGAAGTCGGAATCGCTAGTAATCGCGAATCAGAATG
TCGCGGTGAATACGTTCCCGGGCCTTGTACACACCGCCCGTCACACCATGGGAGTGGGTTGCACCAGAAGTAGCTAGTCT
AACCTTCGGGAGGACGGTTACCACGGTGTGATTCATGACTGGGGTGAAGTCGTAACAAGGTAGCCGTAGGGGAACCTGCG
GCTGGATCACCTCCTTAAATGGACGACAACAAGAAGAAAGCCTTGGCTGCGGCCCTGGGTCAGATCGAACGTCAATTCGG
CAAGGGTGCCGTAATGCGTATGGGCGATCACGACCGCCAGGCGATCCCGGCCATTTCCACTGGCTCTCTGGGCCTGGACA
TCGCGCTCGGCATCGGCGGCCTGCCAAAAGGCCGTATTGTTGAAATCTACGGTCCGGAATCGTCCGGTAAAACCACCCTG
ACCCTGTCGGTGATTGCCCAGGCACAGAAGATGGGCGCCACCTGCGCCTTCGTCGACGCCGAGCACGCACTGGACCCGGA
ATACGCCGGCAAACTGGGGGTCAACGTTGACGACCTGCTGGTTTCCCAGCCGGACACCGGCGAACAGGCGCTGGAAATCA
CCGACATGCTGGTGCGTTCCAATGCCATCGACGTGATCGTGATCGACTCCGTGGCGGCACTGGTACCCAAGGCCGAGATC
GAAGGCGAGATGGGCGACATGCACGTGGGCCTGCAGGCCCGCCTGATGTCCCAGGCGCTGCGCAAGATCACCGGTAACAT
CAAGAACGCCAACTGCCTGGTGATCTTCATCAACCAGATCCGTATGAAAATCGGCGTGATGTTCGGCAGCCCGGAAACCA
CCACCGGCGGTAACGCGCTGAAGTTCTACGCCTCGGTTCGTCTGGACATCCGTCGTACTGGCGCGGTGAAGGAAGGCGAC
GAAGTCGTCGGTAGCGAAACCCGGGTCAAGATCGTCAAGAACAAGGTGGCTCCACCGTTCCGTCAGGCTGAATTCCAGAT
CCTGTACGGCAAGGGTATCTACCTGAACGGCGAGATCATCGATCTGGGCGTGCTGCACGGTTTCCTCGAGAAGTCCGGTG
CCTGGTACAGCTACCAGGGCAACAAGATCGGTCAGGGCAAGGCCAACTCGGCCAAGTTCCTGCAGGACAATCCGGAAATC
GGCAATGCCCTCGAGAAGCAGATTCGCGACAAGCTGCTGGCTCCAACCGCTGATGTCAAAGCTTCGCCGGTCAACGAGAC
CATCGATGACATGGCTGACGCGGATATCTGAATGAGCGAAGAAAACACGTACGACTCGAGCAGCATTAAAGTGCTGAAAG
GTTTGGATGCCGTACGCAAACGTCCCGGTATGTACATTGGTGACACCGACGATGGCAGCGGTCTGCACCATATGGTGTTC
GAGGTGGTCGATAACTCGATCGACGAAGCTCTGGCCGGCCACTGCGACGACATCAGCATCATCATCCACCCGGACGAATC
CATTACCGTGCGTGACAACGGTCGCGGCATCCCGGTAGACGTGCATAAAGAAGAAGGCGTTTCCGCGGCCGAGGTCATCA
TGACTGTGCTGCACGCCGGCGGTAAGTTCGACGACAACTCCTACAAAGTATCCGGCGGTCTGCACGGTGTGGGTGTGTCG
GTAGTGAACGCCCTGTCCGAAGAACTGGTCCTGACCGTTCGCCGCAGTGGCAAGATCTGGGAACAGACCTACGTTCACGG
TGTGCCTCAGGCGCCTATGGCGATCGTCGGTGACAGTGAAACCACCGGTACCCAGATTCACTTCAAGGCTTCCAGCGAGA
CCTTCAAGAACATCCATTTCAGCTGGGACATCCTGGCCAAGCGGATTCGTGAACTGTCCTTCCTCAACTCCGGTGTCGGT
ATCGTTCTGAAGGACGAGCGCAGCGGCAAGGAAGAACTGTTCAAGTACGAAGGCGGTCTGCGTGCGTTCGTTGAATACCT
GAACACCAACAAGACCGCGGTCAACCAGGTGTTCCACTTCAATGTGCAGCGTGAAGATGGCATCGGCGTGGAAATCGCCC
TGCAGTGGAACGACAGCTTCAACGAAAACCTGCAGTGCTTCACCAACAACATTCCGCAGCGCGACGGCGGCACCCACCTG

GTGGGCTTCCGTTCGGCACTGACGCGTAACCTGAACAACTACATCGAACAGGAAGGTCTGGCGAAGAAGCACAAGGTCGC
CACCACCGGTGACGATGCCCGCGAAGGCCTGACCGCAATCATTTCGGTCAAGGTGCCGGATCCGAAGTTCAGCTCCCAGA
CCAAAGACAAGCTGGTGTCTTCCGAAGTGAAGACCGCGGTCGAACAGGAAATGGGCAAGTACTTCTCCGACTTCCTGCTG
GAAAACCCGAACGAAGCCAAGCTGGTGGTCGGCAAGATGCTCGACGCCGCCCGTGCCCGTGAAGCGGCGCGTAAGGCTCG
CGAGATGACCCGCCGTAAAGGTGCGCTGGATATCGCCGGCCTGCCGGGCAAACTGGCGGACTGCCAGGAAAAAGACCCTG
CCCTTTCCGAACTCTACCTGGTGGAAGGTGACTCTGCTGGCGGCTCCGCCAAGCAGGGACGCAACCGTAAGACCCAGGCG
ATTCTGCCGCTCAAGGGCAAGATCCTTAACGTCGAGAAAGCGCGTTTCGACAAGATGATTTCCTCGCAAGAGGTCGGCAC
CTTGATCACTGCACTCGGTTGCGGCATCGGCCGCGAAGAGTACAACATCGACAAGCTGCGTTATCACAACATCATCATCA
TGACCGACGCCGACGTCGACGGTTCGCACATCCGTACCCTGCTGCTGACCTTCTTCTTCCGTCAGCTGCCGGAGCTGATC
GAGCGTGGCTACATCTACATCGCTCAACCGCCGCTGTACAAGGTCAAGAAAGGCAAGCAAGAGCAATACATCAAAGACGA
CGACGCCATGGAAGAGTACATGACGCAGTCGGCCCTGGAAGATGCGAGCCTGCACCTGAACGAAGACACACCGGGTATTT
CCGGCGAGGCGCTGGAGCGCCTGGTGAACGACTTCCGCATGGTCATGAAAACCCTCAAGCGTCTGTCGCGCCTGTACCCT
CAGGAGCTGACCGAGCACTTCATCTACCTGCCGGCCGTGAGCCTGGAGCAGCTCTCCGATCACGCGGCCATGCAGGATTG
GCTGGCCCAATATGAAGTCCGCCTGCGCACCGTCGAGAAGTCCGGCCTGGTCTACAAGGCCAGCCTGCGTGAAGACCGTG
AACGTAATGTCTGGCTGCCAGAGGTCGAACTGATCTCCCACGGCCTGTCGAACTACGTCACCTTCAACCGCGACTTCTTC
GGCAGCAATGACTACAAGACCGTCGTCACACTCGGCGCTCAACTGAGCTCCCTGCTGGACGAAGGCGCTTATATTCAGCG
TGGCGAACGCAAGAAGGCAGTGACCGAGTTCAAGGAAGCCCTGGACTGGCTGATGACCGAAAGCACCAAGCGCCACACCA
TCCAGCGATACAAAGGTCTGGGCGAGATGAACCCGGATCAGCTGTGGGAAACCACCATGGACCCAAGCGTGCGCCGTATG
CTCAAGGTCACCATCGAAGACGCCATCGGCGCCGACCAGATCTTCAACACCCTGATGGGTGATGCGGTCGAGCCTCGTCG
CGACTTCATCGAAAGCAACGCCCTGGCGGTATCCAACTTGGACTTCTGAATGTCCGGAAAAGCGCAACAGCAGTCTCGCC
TCAAAGAGTTGATCAGCCGTGGTCGTGAGCAGGGTTACCTGACTTACGCGGAGGTCAACGACCACCTGCCGGAGGATATT
TCAGATCCGGAACAGGTGGAAGACATCATCCGCATGATCAACGACATGGGGATCAACGTATTCGAGAGTGCTCCGGATGC
GGATGCCCTTTTGTTGGCCGAAGCCGATACCGACGAAGCAGCAGCTGAAGAAGCCGCCGCAGCGTTGGCGGCTGTGGAAA
CCGACATTGGTCGCACTACCGACCCCGTGCGTATGTACATGCGCGAAATGGGTACGGTAGAGCTGCTCACACGTGAAGGC
GAAATCGAAATCGCCAAGCGTATCGAAGAGGGCATCCGTGAAGTGATGGGCGCGATCGCGCACTTCCCTGGCACGGTTGA
GCACATCCTCTCCGAATACACTCGCGTCACCACCGAAGGTGGCCGCCTGTCCGACGTCCTGAGCGGTTACATCGACCCGG
ACGACGGTATTGCGCCGCCTGCCGCCGAAGTACCACCGCCTGTCGATGCCAAGGCCGCAAAAGCGGACGACGACACCGAC
GACGATGACGCCGAAGCCAGTGACGACGAAGAAGAAGCCGAAAGCGGTCCGGATCCGGTCATCGCAGCCCAGCGCTTTGG
CGCCGTTGCCGACCAGATGGAAATTACCCGCAAGGCGCTGAAAAAGCACGGTCGCGAACACAAGCAAGCCCTGGCTGAAA
TGCTGGCCCTGGCTGAGCTGTTCATGCCGATCAAACTGGTTCCGAAGCAATTCGAAGGCCTGGTTGAACGTGTTCGTAGC
GCCCTGGATCGCCTGCGTCAGCAAGAGCGCGCGATCATGCAGCTCTGTGTTCGTGATGCCCGCATGCCACGCGCCGACTT
CCTGCGCCAGTTCCCTGGCAATGAAGTGGACGAAAGCTGGTCCGACGCATTGGCCAAAGGCAAGGCCAAGTACGCCGAAG
CCATCGGCCGCCTGCAGCCGGATATCATCCGTTGCCAGCAGAAGCTGACAGCGCTCGAGACCGAGACTGGCCTGACGATC
GCCGAGATCAAGGACATCAACCGTCGCATGTCGATCGGCGAGGCCAAGGCCCGTCGCGCGAAGAAAGAGATGGTCGAAGC
CAACTTGCGTCTGGTGATCTCCATCGCCAAGAAGTACACCAACCGTGGCCTGCAATTCCTCGACCTGATCCAGGAAGGCA
ACATCGGTTTGATGAAAGCGGTAGACAAGTTCGAATACCGTCGCGGCTACAAATTCTCGACTTATGCCACCTGGTGGATC
CGTCAGGCGATCACTCGTTCGATCGCCGACCAGGCCCGCACCATCCGTATTCCGGTGCACATGATCGAGACGATCAACAA
GCTCAACCGTATTTCCCGGCAGATGTTGCAGGAAATGGGTCGCGAACCGACTCCGGAAGAGCTGGGCGAACGCATGGAAA
TGCCTGAGGACAAGATCCGCAAGGTATTGAAGATCGCTAAAGAGCCGATCTCCATGGAAACCCCGATCGGTGATGACGAA
GACTCCCATCTGGGTGACTTCATCGAAGACTCGACCATGCAGTCGCCAATCGATGTCGCCACCGTTGAGAGCCTTAAAGA
AGCGACTCGCGAAGTACTCTCCGGCCTCACTGCCCGTGAAGCCAAGGTACTGCGCATGCGCTTCGGCATCGACATGAATA
CCGACCACACCCTCGAGGAAGTCGGTAAGCAGTTCGACGTTACCCGTGAGCGGATTCGTCAGATCGAAGCCAAGGCGCTG
CGCAAGCTGCGCCACCCGACGCGAAGCGAGCATCTGCGCTCCTTCCTCGACGAGTAATTGACTAAGCCAGCCATACTCGC
CCTTGCTGATGGCAGCATTTTTCGCGGCGAAGCCATTGGAGCCGACGGTCAAACCGTTGGTGAGGTGGTGTTTAACACCG
CAATGACCGGCTATCAGGAAATCCTTACCGATCCTTCCTACGCCCAACAGATCGTTACCCTGACTTACCCGCACATCGGC
AATACCGGCACCACGCCGGAAGACGCCGAGTCCGATCGTGTCTGGTCGGCCGGTCTGGTGATTCGCGACCTGCCACTGGT
TGCGAGCAACTGGCGTAACACCTTGTCCCTGTCCGACTACCTGAAAGCCAACAATGTTGTGGCGATCGCCGGTATCGACA
CCCGTCGTCTGACGCGCATCCTGCGCGAGAAAGGCGCGCAGAACGGCTGCATCATGGCCGGCGACAATATCTCCGACGAA
GCGGCGATTGCCGCTGCGCGCGGCTTCCCGGGCCTGAAAGGCATGGATCTGGCGAAGGTCGTCAGCACCAAGGAAAGCTA
CGAGTGGCGCTCCAGCGTCTGGAGCCTGAAGACCGACAGTCACCCGACCATCGAGGCTTCCGAGCTGCCTTACCACGTGG
TTGCCTACGACTACGGCGTCAAGCTGAACATCCTGCGCATGCTGGTCGAGCGCGGTTGCCGCGTGACCGTGGTACCTGCG
CAAACCCCGGCCAGCGACGTCCTGGCGCTCAAGCCTGACGGTGTGTTCCTGTCCAACGGTCCTGGCGACCCCGAGCCTTG
CGATTACGCGATCCAGGCGATCAAGGACGTGCTGGAGACCGAGATACCGGTCTTCGGGATCTGCCTGGGCCACCAACTGC
TGGCGCTGGCCGCCGGCGCCAAGACAGTGAAGATGGGCCACGGCCACCACGGTGCCAACCACCCGGTCCAGGACCTGGAC
AGCGGTGTAGTGATGATCACCAGTCAGAACCACGGTTTTGCGGTGGACGAAACCACCCTGCCGGGCAACGTGCGGGCGAT
CCACAAGTCGTTGTTCGACGGCACCCTGCAAGGCATCGAGCTGACCGACAAGAGCGCATTCAGCTTCCAGGGCCACCCTG
AAGCGAGCCCGGGCCCGAACGATGTGGCGCCGCTGTTCGATCGTTTCATCAACGAGATGGCCAAGCGACGCTGAATGAGT
AGCGGACGTATCGTTCAAATCATCGGCGCCGTTATCGACGTGGAATTTCCACGCGACAGCGTACCGAGCATCTACGACGC
CTTGAAGGTTCAAGGCGCCGAAACCACTCTGGAAGTTCAGCAGCAGCTGGGCGACGGCGTGGTACGTACCATTGCGATGG
GCTCCACCGAGGGCTTGAAGCGCGGTCTGGACGTCAACAACACTGGCGCAGCCATCTCCGTACCGGTCGGTAAAGCGACC
CTGGGCCGGATCATGGACGTACTGGGCAACCCGATCGACGAAGCTGGCCCGATCGGCGAAGAAGAGCGTTGGGGCATTCA
CCGTCCTGCGCCGACCTTCGCTGAACAAGCTGGTGGCAACGACCTGCTGGAAACCGGCATCAAGGTTATCGACCTGGTTT
GCCCGTTCGCCAAGGGCGGTAAAGTTGGTCTGTTCGGTGGTGCCGGTGTGGGCAAAACCGTAAACATGATGGAACTGATC
CGTAACATCGCCATCGAGCACAGCGGTTATTCCGTGTTCGCCGGTGTGGGCGAGCGTACTCGTGAGGGTAACGACTTCTA

CCACGAGATGAAGGATTCCAACGTTCTGGACAAAGTGGCACTGGTATACGGCCAGATGAACGAGCCGCCGGGAAACCGTC
TGCGCGTAGCTCTGACCGGCCTGACCATGGCCGAGAAGTTCCGTGACGAAGGTAACGACGTTCTGCTGTTCGTCGACAAC
ATCTATCGTTACACCCTGGCCGGTACCGAAGTATCCGCACTGCTGGGCCGTATGCCTTCGGCAGTAGGTTACCAGCCGAC
CCTGGCTGAAGAGATGGGCGTTCTGCAAGAACGTATCACTTCGACCAAGCAAGGCTCGATCACCTCGATCCAAGCGGTAT
ACGTACCTGCGGACGACTTGACCGACCCGTCGCCAGCGACCACCTTCGCCCACTTGGACGCCACCGTCGTTCTTTCCCGT
GACATCGCTTCCCTGGGTATCTACCCAGCGGTAGACCCACTGGACTCGACTTCCCGTCAGCTGGACCCGAACGTGATCGG
CAACGAGCACTATGAAACCGCTCGCGGCGTTCAGTACGTGCTGCAGCGCTACAAAGAGCTGAAGGACATCATTGCGATCC
TGGGTATGGACGAACTGTCCGAAGCCGACAAGCAACTGGTATCCCGCGCTCGTAAGATCCAGCGCTTCCTGTCGCAGCCG
TTCTTCGTGGCTGAAGTCTTCACTGGTTCTCCAGGCAAATACGTTTCCCTGAAAGACACCATCGCTGGCTTCAAAGGCAT
CCTCAACGGTGACTACGACCATCTGCCAGAACAAGCGTTCTACATGGTTGGTGGCATCGAAGAAGCGATCGAGAA

NCBI Reference Sequence: NZ_CP027746.1
Strain: DSM19603
Chained genes: 16S rRNA, recA, gyrB, rpoD, carA, atpD >NZ_CP027746_DSM19603
GAACTGAAGAGTTTGATCATGGCTCAGATTGAACGCTGGCGGCAGGCCTAACACATGCAAGTCGAGCGGTAGAGAGAAGC
TTGCTTCTCTTGAGAGCGGCGGACGGGTGAGTAATGCCTAGGAATCTGCCTGGTAGTGGGGGATAACGTCCGGAAACGGA
CGCTAATACCGCATACGTCCTACGGGAGAAAGCAGGGGACCTTCGGGCCTTGCGCTATCAGATGAGCCTAGGTCGGATTA
GCTAGTTGGTGAGGTAATGGCTCACCAAGGCGACGATCCGTAACTGGTCTGAGAGGATGATCAGTCACACTGGAACTGAG
ACACGGTCCAGACTCCTACGGGAGGCAGCAGTGGGGAATATTGGACAATGGGCGAAAGCCTGATCCAGCCATGCCGCGTG
TGTGAAGAAGGTCTTCGGATTGTAAAGCACTTTAAGTTGGGAGGAAGGGTACTTACCTAATACGTGAGTATTTTGACGTT
ACCGACAGAATAAGCACCGGCTAACTCTGTGCCAGCAGCCGCGGTAATACAGAGGGTGCAAGCGTTAATCGGAATTACTG
GGCGTAAAGCGCGCGTAGGTGGTTCGTTAAGTTGGATGTGAAATCCCCGGGCTCAACCTGGGAACTGCATCCAAAACTGG
CGAGCTAGAGTATGGTAGAGGGTGGTGGAATTTCCTGTGTAGCGGTGAAATGCGTAGATATAGGAAGGAACACCAGTGGC
GAAGGCGACCACCTGGACTGATACTGACACTGAGGTGCGAAAGCGTGGGGAGCAAACAGGATTAGATACCCTGGTAGTCC
ACGCCGTAAACGATGTCAACTAGCCGTTGGGAGCCTTGAGCTCTTAGTGGCGCAGCTAACGCATTAAGTTGACCGCCTGG
GGAGTACGGCCGCAAGGTTAAAACTCAAATGAATTGACGGGGGCCCGCACAAGCGGTGGAGCATGTGGTTTAATTCGAAG
CAACGCGAAGAACCTTACCAGGCCTTGACATCCAATGAACTTTCCAGAGATGGATTGGTGCCTTCGGGAACATTGAGACA
GGTGCTGCATGGCTGTCGTCAGCTCGTGTCGTGAGATGTTGGGTTAAGTCCCGTAACGAGCGCAACCCTTGTCCTTAGTT
ACCAGCACGTTATGGTGGGCACTCTAAGGAGACTGCCGGTGACAAACCGGAGGAAGGTGGGGATGACGTCAAGTCATCAT
GGCCCTTACGGCCTGGGCTACACACGTGCTACAATGGTCGGTACAGAGGGTTGCCAAGCCGCGAGGTGGAGCTAATCCCA
TAAAACCGATCGTAGTCCGGATCGCAGTCTGCAACTCGACTGCGTGAAGTCGGAATCGCTAGTAATCGCGAATCAGAATG
TCGCGGTGAATACGTTCCCGGGCCTTGTACACACCGCCCGTCACACCATGGGAGTGGGTTGCACCAGAAGTAGCTAGTCT
AACCTTCGGGAGGACGGTTACCACGGTGTGATTCATGACTGGGGTGAAGTCGTAACAAGGTAGCCGTAGGGGAACCTGCG
GCTGGATCACCTCCTTAAATGGACGACAACAAGAAGAAAGCCTTGGCTGCGGCCCTGGGTCAGATCGAACGTCAATTCGG
CAAGGGTGCCGTAATGCGTATGGGCGATCACGACCGCCAGGCGATCCCGGCCATTTCCACTGGCTCTCTGGGCCTGGACA
TCGCGCTCGGCATCGGCGGCCTGCCAAAAGGCCGTATTGTTGAAATCTACGGTCCGGAATCGTCCGGTAAAACCACCCTG
ACCCTGTCGGTGATTGCCCAGGCACAGAAGATGGGCGCCACCTGCGCCTTCGTCGACGCCGAGCACGCACTGGACCCGGA
ATACGCCGGCAAACTGGGGGTCAACGTTGACGACCTGCTGGTTTCCCAGCCGGACACCGGCGAACAGGCGCTGGAAATCA
CCGACATGCTGGTGCGTTCCAATGCCATCGACGTGATCGTGATCGACTCCGTGGCGGCACTGGTACCCAAGGCCGAGATC
GAAGGCGAGATGGGCGACATGCACGTGGGCCTGCAGGCCCGCCTGATGTCCCAGGCGCTGCGCAAGATCACCGGTAACAT
CAAGAACGCCAACTGCCTGGTGATCTTCATCAACCAGATCCGTATGAAAATCGGCGTGATGTTCGGCAGCCCGGAAACCA
CCACCGGCGGTAACGCGCTGAAGTTCTACGCCTCGGTTCGTCTGGACATCCGTCGTACTGGCGCGGTGAAGGAAGGCGAC
GAAGTCGTCGGTAGCGAAACCCGGGTCAAGATCGTCAAGAACAAGGTGGCTCCACCGTTCCGTCAGGCTGAATTCCAGAT
CCTGTACGGCAAGGGTATCTACCTGAACGGCGAGATCATCGATCTGGGCGTGCTGCACGGTTTCCTCGAGAAGTCCGGTG
CCTGGTACAGCTACCAGGGCAACAAGATCGGTCAGGGCAAGGCCAACTCGGCCAAGTTCCTGCAGGACAATCCGGAAATC
GGCAATGCCCTCGAGAAGCAGATTCGCGACAAGCTGCTGGCTCCAACCGCTGATGTCAAAGCTTCGCCGGTCAACGAGAC
CATCGATGACATGGCTGACGCGGATATCTGAATGAGCGAAGAAAACACGTACGACTCGAGCAGCATTAAAGTGCTGAAAG
GTTTGGATGCCGTACGCAAACGTCCCGGTATGTACATTGGTGACACCGACGATGGCAGCGGTCTGCACCATATGGTGTTC
GAGGTGGTCGATAACTCGATCGACGAAGCTCTGGCCGGCCACTGCGACGACATCAGCATCATCATCCACCCGGACGAATC
CATTACCGTGCGTGACAACGGTCGCGGCATCCCGGTAGACGTGCATAAAGAAGAAGGCGTTTCCGCGGCCGAGGTCATCA
TGACTGTGCTGCACGCCGGCGGTAAGTTCGACGACAACTCCTACAAAGTATCCGGCGGTCTGCACGGTGTGGGTGTGTCG
GTAGTGAACGCCCTGTCCGAAGAACTGGTCCTGACCGTTCGCCGCAGTGGCAAGATCTGGGAACAGACCTACGTTCACGG
TGTGCCTCAGGCGCCTATGGCGATCGTCGGTGACAGTGAAACCACCGGTACCCAGATTCACTTCAAGGCTTCCAGCGAGA
CCTTCAAGAACATCCATTTCAGCTGGGACATCCTGGCCAAGCGGATTCGTGAACTGTCCTTCCTCAACTCCGGTGTCGGT
ATCGTTCTGAAGGACGAGCGTAGCGGCAAGGAAGAACTGTTCAAGTACGAAGGCGGTCTGCGTGCGTTCGTTGAATACCT
GAACACCAACAAGACCGCGGTCAACCAGGTGTTCCATTTCAATGTGCAGCGTGAAGATGGCATCGGCGTGGAAATCGCCC
TGCAGTGGAATGACAGCTTCAACGAAAACCTGCAGTGCTTCACCAACAACATTCCGCAGCGCGATGGCGGCACCCACCTG
GTGGGCTTCCGTTCGGCACTGACACGTAACCTGAACAACTACATCGAACAGGAAGGTCTGGCGAAGAAGCACAAGGTCGC
CACCACCGGTGACGATGCCCGCGAAGGCCTGACCGCAATCATTTCGGTCAAGGTGCCGGATCCGAAGTTCAGCTCCCAGA
CCAAAGACAAGCTGGTGTCTTCCGAAGTGAAGACCGCGGTCGAACAGGAAATGGGCAAGTACTTCTCCGACTTCCTGCTG
GAAAACCCGAACGAAGCCAAGCTGGTGGTCGGCAAGATGCTCGACGCCGCCCGTGCCCGTGAAGCGGCGCGTAAGGCTCG
TGAGATGACCCGCCGTAAAGGTGCGCTGGATATCGCCGGCCTGCCGGGCAAACTGGCGGACTGCCAGGAAAAAGACCCTG
CCCTTTCCGAACTCTACCTGGTGGAAGGTGACTCTGCTGGCGGCTCCGCCAAGCAGGGACGCAACCGTAAGACCCAGGCG
ATTCTGCCGCTCAAGGGCAAGATCCTTAACGTCGAGAAAGCGCGTTTCGACAAGATGATTTCCTCGCAAGAGGTCGGCAC

CTTGATCACTGCACTCGGTTGCGGCATCGGCCGCGAAGAGTACAACATCGACAAGCTGCGTTATCACAACATCATCATCA
TGACTGACGCCGACGTCGATGGTTCGCACATCCGTACCCTGCTGCTGACCTTCTTCTTCCGTCAGCTGCCGGAGCTGATC
GAGCGTGGCTACATCTACATCGCTCAACCGCCGCTGTACAAGGTCAAGAAAGGCAAGCAAGAGCAATACATCAAAGACGA
CGACGCCATGGAAGAGTACATGACGCAGTCGGCCCTGGAAGATGCGAGCCTGCACCTGAACGAAGACGCACCGGGTATTT
CCGGCGAGGCGCTGGAGCGCCTGGTGAACGACTTCCGCATGGTCATGAAAACCCTCAAGCGTCTGTCGCGCCTGTACCCT
CAGGAGCTGACCGAGCACTTCATCTACCTGCCGGCCGTGAGCCTGGAGCAGCTCTCCGATCACGCGGCGATGCAGGATTG
GCTGGCCCAATATGAAGTCCGCCTGCGCACCGTCGAGAAGTCCGGCCTGGTCTACAAGGCCAGCCTGCGTGAAGACCGTG
AACGTAATGTCTGGCTGCCAGAGGTCGAACTGATCTCCCACGGCCTGTCGAACTACGTCACCTTCAACCGCGACTTCTTC
GGCAGCAATGACTACAAGACCGTCGTCACCCTCGGTGCTCAACTGAGCTCCCTGCTGGACGAAGGCGCTTATATTCAGCG
TGGCGAACGCAAGAAGGCAGTGACCGAGTTCAAGGAAGCCCTGGACTGGCTGATGACCGAAAGTACCAAGCGCCACACCA
TCCAGCGATACAAAGGTCTGGGCGAGATGAACCCGGATCAGCTGTGGGAAACCACCATGGACCCAAGCGTGCGCCGTATG
CTCAAGGTCACCATCGAAGACGCCATCGGCGCCGACCAGATCTTCAACACCCTGATGGGTGATGCGGTCGAGCCTCGTCG
CGACTTCATCGAAAGCAACGCCCTGGCGGTATCCAACTTGGACTTCTGAATGTCCGGAAAAGCGCAACAGCAGTCTCGCC
TCAAAGAGTTGATCAGCCGTGGTCGTGAGCAGGGTTACCTGACTTACGCGGAGGTCAACGACCACCTGCCGGAGGATATT
TCAGATCCGGAACAGGTGGAAGACATCATCCGCATGATCAACGACATGGGGATCAACGTATTCGAGAGTGCTCCGGATGC
GGATGCCCTTTTGTTGGCCGAAGCCGATACCGACGAAGCAGCAGCTGAAGAAGCCGCCGCAGCGTTGGCGGCTGTGGAAA
CCGACATTGGTCGCACTACCGACCCCGTGCGTATGTACATGCGCGAAATGGGTACGGTAGAGCTGCTCACACGTGAAGGC
GAAATCGAAATCGCCAAGCGTATCGAAGAGGGCATCCGTGAAGTGATGGGCGCGATCGCGCACTTCCCTGGCACGGTTGA
GCACATCCTCTCCGAATACACTCGCGTCACCACCGAAGGTGGCCGCCTGTCCGACGTCCTGAGCGGTTACATCGACCCGG
ACGACGGTATTGCGCCGCCTGCCGCCGAAGTACCACCGCCTGTCGATGCCAAGGCCGCAAAAGCGGACGACGACACCGAC
GACGATGACGCCGAAGCCAGTGACGACGAAGAAGAAGCCGAAAGCGGTCCGGATCCGGTCATCGCAGCCCAGCGCTTTGG
CGCCGTTGCCGACCAGATGGAAATTACCCGCAAGGCGCTGAAAAAGCACGGTCGCGAACACAAGCAAGCCCTGGCTGAAA
TGCTGGCCCTGGCTGAGCTGTTCATGCCGATCAAACTGGTTCCGAAGCAATTCGAAGGCCTGGTTGAACGTGTTCGTAGC
GCCCTGGATCGCCTGCGTCAGCAAGAGCGCGCGATCATGCAGCTCTGTGTTCGTGATGCCCGCATGCCACGCGCCGACTT
CCTGCGCCAGTTCCCTGGCAATGAAGTGGACGAAAGCTGGTCCGACGCATTGGCCAAAGGCAAGGCCAAGTACGCCGAAG
CCATCGGCCGCCTGCAGCCGGATATCATCCGTTGCCAGCAGAAGCTGACAGCGCTCGAGACCGAGACTGGCCTGACGATC
GCCGAGATCAAGGACATCAACCGTCGCATGTCGATCGGCGAGGCCAAGGCCCGTCGCGCGAAGAAAGAGATGGTCGAAGC
CAACTTGCGTCTGGTGATCTCCATCGCCAAGAAGTACACCAACCGTGGCCTGCAATTCCTCGACCTGATCCAGGAAGGCA
ACATCGGTTTGATGAAAGCGGTAGACAAGTTCGAATACCGTCGCGGCTACAAATTCTCGACTTATGCCACCTGGTGGATC
CGTCAGGCGATCACTCGTTCGATCGCCGACCAGGCCCGCACCATCCGTATTCCGGTGCACATGATCGAGACGATCAACAA
GCTCAACCGTATTTCCCGGCAGATGTTGCAGGAAATGGGTCGCGAACCGACTCCGGAAGAGCTGGGCGAACGCATGGAAA
TGCCTGAGGACAAGATCCGCAAGGTATTGAAGATCGCTAAAGAGCCGATCTCCATGGAAACCCCGATCGGTGATGACGAA
GACTCCCATCTGGGTGACTTCATCGAAGACTCGACCATGCAGTCGCCAATCGATGTCGCCACCGTTGAGAGCCTTAAAGA
AGCGACTCGCGAAGTACTCTCCGGCCTCACTGCCCGTGAAGCCAAGGTACTGCGCATGCGCTTCGGCATCGACATGAATA
CCGACCACACCCTCGAGGAAGTCGGTAAGCAGTTCGACGTTACCCGTGAGCGGATTCGTCAGATCGAAGCCAAGGCGCTG
CGCAAGCTGCGCCACCCGACGCGAAGCGAGCATCTGCGCTCCTTCCTCGACGAGTAATTGACTAAGCCAGCCATACTCGC
CCTTGCTGATGGCAGCATTTTTCGCGGCGAAGCCATTGGAGCCGACGGTCAAACCGTTGGTGAGGTGGTGTTTAACACCG
CAATGACCGGCTATCAGGAAATCCTTACCGATCCTTCCTACGCCCAACAGATCGTTACCCTGACTTACCCGCACATCGGC
AATACCGGCACCACGCCGGAAGACGCCGAGTCCGATCGTGTCTGGTCGGCCGGTCTGGTGATTCGCGACCTGCCACTGGT
TGCGAGCAACTGGCGTAACACCTTGTCCCTGTCCGACTACCTGAAAGCCAACAATGTTGTGGCGATCGCCGGTATCGACA
CCCGTCGTCTGACGCGCATCCTGCGCGAGAAAGGCGCGCAGAACGGCTGCATCATGGCCGGCGACAATATCTCCGACGAA
GCGGCGATTGCCGCTGCGCGCGGCTTCCCGGGCCTGAAAGGCATGGATCTGGCGAAGGTCGTCAGCACCAAGGAAAGCTA
CGAGTGGCGCTCCAGCGTCTGGAGCCTGAAGACCGACAGTCACCCGACCATCGAGGCTTCCGAGCTGCCTTACCACGTGG
TTGCCTACGACTACGGCGTCAAGCTGAACATCCTGCGCATGCTGGTCGAGCGCGGTTGCCGCGTGACCGTGGTACCTGCG
CAAACCCCGGCCAGCGACGTCCTGGCGCTCAAGCCTGACGGTGTGTTCCTGTCCAACGGTCCTGGCGACCCCGAGCCTTG
CGATTACGCGATCCAGGCGATCAAGGACGTGCTGGAGACCGAGATACCGGTCTTCGGGATCTGCCTGGGCCACCAACTGC
TGGCGCTGGCCGCCGGCGCCAAGACAGTGAAGATGGGCCACGGCCACCACGGTGCCAACCACCCGGTCCAGGACCTGGAC
AGCGGTGTAGTGATGATCACCAGTCAGAACCACGGTTTTGCGGTGGACGAAACCACCCTGCCGGGCAACGTGCGGGCGAT
CCACAAGTCGTTGTTCGACGGCACTCTGCAAGGCATCGAGCTGACCGACAAGAGCGCATTCAGCTTCCAGGGCCACCCTG
AAGCGAGCCCGGGCCCGAACGATGTGGCGCCGCTGTTCGATCGTTTCATCAACGAGATGGCCAAGCGACGCTGAATGAGT
AGCGGACGTATCGTTCAAATCATCGGCGCCGTTATCGACGTGGAATTTCCACGCGACAGCGTACCGAGCATCTACGACGC
CTTGAAGGTTCAAGGCGCCGAAACCACTCTGGAAGTTCAGCAGCAGCTGGGCGACGGCGTGGTACGTACCATTGCGATGG
GCTCCACCGAGGGCTTGAAGCGCGGTCTGGACGTCAACAACACTGGCGCAGCCATCTCCGTACCGGTCGGTAAAGCGACC
CTGGGCCGGATCATGGACGTACTGGGCAACCCGATCGACGAAGCTGGCCCGATCGGCGAAGAAGAGCGTTGGGGCATTCA
CCGTCCTGCGCCGACCTTCGCTGAACAAGCTGGTGGCAACGACCTGCTGGAAACCGGCATCAAGGTTATCGACCTGGTTT
GCCCGTTCGCCAAGGGCGGTAAAGTTGGTCTGTTCGGTGGTGCCGGTGTGGGCAAAACCGTAAACATGATGGAACTGATC
CGTAACATCGCCATCGAGCACAGCGGTTATTCCGTGTTCGCCGGTGTGGGCGAGCGTACTCGTGAGGGTAACGACTTCTA
CCACGAGATGAAGGATTCCAACGTTCTGGACAAAGTGGCACTGGTATACGGCCAGATGAACGAGCCGCCGGGAAACCGTC
TGCGCGTAGCTCTGACCGGCCTGACCATGGCCGAGAAGTTCCGTGACGAAGGTAACGACGTTCTGCTGTTCGTCGACAAC
ATCTATCGTTACACCCTGGCCGGTACCGAAGTATCCGCACTGCTGGGCCGTATGCCTTCGGCAGTAGGTTACCAGCCGAC
CCTGGCTGAAGAGATGGGCGTTCTGCAAGAACGTATCACTTCGACCAAGCAAGGCTCGATCACCTCGATCCAAGCGGTAT
ACGTACCTGCGGACGACTTGACCGACCCGTCGCCAGCGACCACCTTCGCCCACTTGGACGCCACCGTCGTTCTTTCCCGT
GACATCGCTTCCCTGGGTATCTACCCAGCGGTAGACCCACTGGACTCGACTTCCCGTCAGCTGGACCCGAACGTGATCGG
CAACGAGCACTATGAAACCGCTCGCGGCGTTCAGTACGTGCTGCAGCGCTACAAAGAGCTGAAGGACATCATTGCGATCC

TGGGTATGGACGAACTGTCCGAAGCCGACAAGCAACTGGTATCCCGCGCTCGTAAGATCCAGCGCTTCCTGTCGCAGCCG
TTCTTCGTGGCTGAAGTCTTCACTGGTTCTCCAGGCAAATACGTTTCCCTGAAAGACACCATCGCTGGCTTCAAAGGCAT
CCTCAACGGTGACTACGACCATCTGCCAGAACAAGCGTTCTACATGGTTGGTGGCATCGAAGAAGCGATCGAGAAAGCCA
AGAAACTGTAA

NCBI Reference Sequence: NZ_CP027709.1
Strain: ATCC17809
Chained genes: 16S rRNA, recA, gyrB, rpoD, carA, atpD >NZ_CP027709_ATCC17809
GAACTGAAGAGTTTGATCATGGCTCAGATTGAACGCTGGCGGCAGGCCTAACACATGCAAGTCGAGCGGTAGAGAGGTGC
TTGCACCTCTTGAGAGCGGCGGACGGGTGAGTAATGCCTAGGAATCTGCCTGGTAGTGGGGGATAACGTTCGGAAACGGA
CGCTAATACCGCATACGTCCTACGGGAGAAAGCAGGGGACCTTCGGGCCTTGCGCTATCAGATGAGCCTAGGTCGGATTA
GCTAGTTGGTGAGGTAATGGCTCACCAAGGCGACGATCCGTAACTGGTCTGAGAGGATGATCAGTCACACTGGAACTGAG
ACACGGTCCAGACTCCTACGGGAGGCAGCAGTGGGGAATATTGGACAATGGGCGAAAGCCTGATCCAGCCATGCCGCGTG
TGTGAAGAAGGTCTTCGGATTGTAAAGCACTTTAAGTTGGGAGGAAGGGTACTTACCTAATACGTGAGTATTTTGACGTT
ACCGACAGAATAAGCACCGGCTAACTCTGTGCCAGCAGCCGCGGTAATACAGAGGGTGCAAGCGTTAATCGGAATTACTG
GGCGTAAAGCGCGCGTAGGTGGTTCGTTAAGTTGAATGTGAAATCCCCGGGCTCAACCTGGGAACTGCATCCAAAACTGG
CGAGCTAGAGTATGGTAGAGGGTGGTGGAATTTCCTGTGTAGCGGTGAAATGCGTAGATATAGGAAGGAACACCAGTGGC
GAAGGCGACCACCTGGACTGATACTGACACTGAGGTGCGAAAGCGTGGGGAGCAAACAGGATTAGATACCCTGGTAGTCC
ACGCCGTAAACGATGTCAACTAGCCGTTGGGAGCCTTGAGCTCTTAGTGGCGCAGCTAACGCATTAAGTTGACCGCCTGG
GGAGTACGGCCGCAAGGTTAAAACTCAAATGAATTGACGGGGGCCCGCACAAGCGGTGGAGCATGTGGTTTAATTCGAAG
CAACGCGAAGAACCTTACCAGGCCTTGACATCCAATGAACTTTCCAGAGATGGATTGGTGCCTTCGGGAACATTGAGACA
GGTGCTGCATGGCTGTCGTCAGCTCGTGTCGTGAGATGTTGGGTTAAGTCCCGTAACGAGCGCAACCCTTGTCCTTAGTT
ACCAGCACGTAATGGTGGGCACTCTAAGGAGACTGCCGGTGACAAACCGGAGGAAGGTGGGGATGACGTCAAGTCATCAT
GGCCCTTACGGCCTGGGCTACACACGTGCTACAATGGTCGGTACAGAGGGTTGCCAAGCCGCGAGGTGGAGCTAATCCCA
TAAAACCGATCGTAGTCCGGATCGCAGTCTGCAACTCGACTGCGTGAAGTCGGAATCGCTAGTAATCGCGAATCAGAATG
TCGCGGTGAATACGTTCCCGGGCCTTGTACACACCGCCCGTCACACCATGGGAGTGGGTTGCACCAGAAGTAGCTAGTCT
AACCTTCGGGAGGACGGTTACCACGGTGTGATTCATGACTGGGGTGAAGTCGTAACAAGGTAGCCGTAGGGGAACCTGCG
GCTGGATCACCTCCTTAAATGGACGACAACAAGAAGAAAGCCTTGGCTGCGGCCCTGGGTCAGATCGAACGTCAATTCGG
CAAGGGTGCCGTAATGCGTATGGGCGATCACGATCGCCAGGCGATCCCGGCCATTTCCACTGGCTCTCTGGGTCTGGACA
TCGCACTCGGCATCGGCGGCCTGCCAAAAGGCCGTATTGTTGAAATCTACGGTCCGGAATCGTCCGGTAAAACCACCCTG
ACCCTGTCGGTGATCGCCCAGGCACAGAAGATGGGCGCCACCTGCGCCTTCGTCGACGCCGAGCACGCACTGGACCCGGA
ATACGCCGGCAAACTGGGGGTCAACGTTGACGACCTGCTGGTTTCCCAGCCGGACACCGGCGAACAGGCGCTGGAAATCA
CCGACATGCTGGTGCGCTCCAATGCCATCGACGTGATCGTGATCGACTCCGTGGCGGCGCTGGTACCCAAGGCCGAGATC
GAAGGCGAGATGGGCGACATGCACGTGGGCCTGCAGGCTCGCCTGATGTCCCAGGCGCTGCGCAAGATCACCGGTAACAT
CAAGAACGCCAACTGCCTGGTGATCTTCATCAACCAGATCCGTATGAAAATCGGCGTGATGTTCGGCAGCCCGGAAACCA
CCACCGGTGGTAACGCGCTGAAGTTCTACGCTTCGGTTCGTCTGGACATCCGTCGTACTGGCGCGGTGAAGGAAGGTGAC
GAAGTCGTCGGTAGCGAAACCCGGGTCAAGATCGTCAAGAACAAGGTGGCTCCACCTTTCCGTCAAGCTGAGTTCCAGAT
CCTGTACGGCAAGGGTATCTACCTGAACGGCGAGATCATCGATCTGGGCGTGCTGCACGGTTTCCTCGAGAAGTCCGGTG
CCTGGTACAGCTACCAGGGCAACAAGATCGGTCAGGGCAAGGCCAACTCGGCCAAGTTCCTGCAGGACAATCCGGAAATC
GGTAATGCCCTCGAGAAGCAGATTCGCGACAAGCTGCTGGCTCCGAGCGGAGATACCAAGGCTCTGCCCGTCAACGAGAC
CATCGATGACATGGCCGACGCGGATATCTGAATGAGCGAAGAAAACACGTACGACTCGAGCAGCATTAAAGTGCTGAAAG
GTTTGGATGCCGTACGCAAACGTCCCGGTATGTACATTGGTGACACCGACGATGGCAGCGGTCTGCACCATATGGTGTTC
GAGGTGGTCGACAACTCGATCGACGAAGCTCTGGCCGGCCACTGCGACGACATCAGCATCATCATCCACCCGGACGAATC
CATTACCGTGCGTGACAACGGTCGCGGCATCCCGGTAGACGTGCATAAAGAAGAAGGCGTTTCCGCAGCCGAGGTCATCA
TGACCGTGCTGCACGCCGGCGGTAAGTTCGACGACAACTCCTACAAGGTATCCGGCGGTCTGCACGGTGTGGGTGTGTCG
GTAGTGAACGCCCTGTCCGAAGAACTGGTGCTGACCGTTCGCCGCAGTGGCAAGATCTGGGAACAGACCTACGTTCACGG
TGTGCCTCAGGCGCCTATGGCAATTGTCGGTGACAGCGAGACCACCGGTACCCAGATTCACTTCAAGGCTTCCAGCGAGA
CCTTCAAGAACATCCATTTCAGCTGGGACATCCTGGCCAAGCGGATTCGTGAACTGTCCTTCCTCAACTCCGGTGTCGGT
ATCGTTCTGAAGGACGAGCGCAGCGGCAAGGAAGAGCTGTTCAAGTACGAAGGCGGCCTGCGCGCATTCGTTGAATACCT
GAACACCAACAAGACTGCGGTCAACCAGGTGTTCCACTTCAACGTGCAGCGTGAAGACGGCATCGGCGTGGAAATCGCCC
TGCAGTGGAACGACAGCTTCAACGAAAACCTGCAGTGCTTCACCAACAACATTCCGCAGCGCGACGGCGGCACCCACCTG
GTGGGCTTCCGCTCGGCACTGACGCGTAACCTGAACAACTACATCGAGCAGGAAGGTCTGGCGAAGAAGCACAAGGTGGC
CACCACCGGTGACGATGCCCGCGAAGGCCTGACCGCGATCATTTCGGTCAAGGTGCCGGATCCGAAGTTCAGCTCCCAGA
CCAAAGATAAGCTGGTGTCTTCCGAAGTGAAGACCGCAGTCGAACAGGAAATGGGCAAGTACTTCTCCGACTTCCTGCTG
GAAAACCCGAACGAAGCCAAGCTGGTGGTCGGCAAGATGCTCGACGCCGCCCGTGCCCGTGAAGCGGCGCGTAAGGCTCG
TGAGATGACCCGCCGTAAAGGTGCGCTGGATATCGCCGGCCTGCCGGGCAAACTGGCGGACTGCCAGGAAAAGGACCCTG
CCCTTTCCGAACTCTACCTGGTGGAAGGTGACTCTGCTGGCGGCTCCGCCAAGCAGGGACGCAACCGCAAGACCCAGGCG
ATTCTGCCGCTCAAGGGCAAGATTCTTAACGTCGAGAAAGCGCGCTTCGACAAGATGATTTCCTCGCAAGAGGTCGGCAC
CTTGATCACTGCACTCGGCTGCGGCATCGGCCGCGAAGAGTACAACATCGACAAGCTGCGTTATCACAACATCATCATCA
TGACCGACGCCGACGTCGACGGTTCGCACATCCGCACCCTGCTGCTGACTTTCTTCTTCCGTCAGCTGCCGGAGCTGATC
GAGCGTGGCTACATCTACATCGCTCAACCGCCGCTGTACAAGGTCAAGAAAGGCAAGCAAGAGCAATACATCAAAGACGA
CGACGCCATGGAAGAGTACATGACGCAGTCGGCCCTGGAAGATGCGAGCCTGCACCTGAACGAAGAAGCACCGGGTATTT
CCGGCGAGGCGCTGGAGCGCCTGGTGAACGACTTCCGCATGGTCATGAAAACCCTCAAGCGCCTGTCGCGCCTGTACCCT
CAGGAGCTGACCGAACACTTCATCTACCTGCCAGCCGTGAGCCTGGAGCAACTCTCCGATCACGCAGCGATGCAGGATTG

GTTGGCCCAATATGAAGTCCGCCTGCGCACCGTCGAGAAGTCCGGCCTGGTCTACAAGGCCAGCCTGCGTGAAGACCGTG
AACGTAATGTCTGGCTGCCAGAGGTCGAACTGATCTCCCACGGCCTGTCGAACTACGTCACCTTCAACCGCGACTTCTTC
GGCAGCAATGACTACAAGACCGTCGTCACCCTCGGCGCTCAACTGAGCTCCCTGCTGGACGAAGGCGCTTATATTCAGCG
TGGCGAACGCAAGAAGGCGGTGACCGAGTTCAAGGAAGCCCTGGACTGGCTGATGACCGAAAGCACCAAGCGCCACACCA
TCCAGCGATACAAAGGTCTGGGCGAGATGAACCCGGATCAGCTGTGGGAAACCACCATGGACCCAAGCGTGCGCCGCATG
CTCAAGGTCACCATCGAAGACGCCATCGGCGCCGACCAGATCTTCAACACCCTGATGGGTGATGCGGTCGAGCCTCGTCG
CGACTTCATCGAAAGCAACGCCCTGGCGGTATCCAACCTGGACTTCTGAATGTCCGGAAAAGCGCAACAGCAGTCTCGCC
TCAAAGAGTTGATCAGCCGTGGTCGTGAGCAGGGTTACCTGACTTACGCGGAGGTCAACGACCACCTGCCGGAGGATATT
TCAGATCCGGAACAGGTGGAAGACATCATCCGCATGATCAACGACATGGGGATCAACGTATTCGAGAGTGCTCCGGATGC
GGATGCCCTTTTGTTGGCCGAAGCCGATACCGACGAAGCAGCAGCTGAAGAAGCCGCCGCAGCGTTGGCGGCTGTGGAAA
CCGACATTGGTCGCACTACCGACCCAGTGCGTATGTACATGCGCGAAATGGGCACGGTAGAGCTGCTCACACGTGAAGGC
GAAATCGAAATCGCCAAGCGTATCGAAGAGGGCATCCGTGAAGTGATGGGCGCGATCGCGCACTTCCCTGGCACGGTTGA
GCACATCCTCTCCGAATACACTCGCGTCACCACCGAAGGTGGCCGCCTGTCCGACGTCCTGAGCGGTTACATCGACCCGG
ATGACGGCATTGCGCCGCCTGCCGCCGAAGTACCACCGCCTGTCGATGCCAAGGCCGCGAAAGCGGACGACGACACCGAC
GACGATGACGCCGAAGCCAGTGACGACGAAGAAGAAGCCGAAAGCGGTCCGGATCCGGTCATCGCAGCCCAGCGCTTTGG
CGCCGTTGCCGACCAGATGGAAATCACCCGCAAGGCGCTGAAAAAGCACGGTCGCGAACACAAGCAAGCCCTGGCTGAAA
TGCTGGCCCTGGCTGAACTGTTCATGCCGATCAAACTGGTTCCGAAGCAATTCGAAGGCCTGGTTGAACGTGTTCGTAGC
GCCCTGGATCGCCTGCGTCAGCAAGAGCGCGCGATCATGCAGCTCTGTGTTCGTGATGCCCGCATGCCACGCGCCGACTT
CCTGCGCCAGTTCCCTGGCAATGAAGTGGACGAAAGCTGGTCCGACGCGCTGGCCAAAGGCAAGGCCAAGTACGCCGAAG
CCATCGGCCGCCTGCAGCCGGACATCATCCGTTGCCAGCAGAAGCTGACCGCGCTCGAGACCGAGACCGGCCTGACGATT
GCCGAGATCAAGGACATCAACCGTCGCATGTCGATCGGCGAGGCCAAGGCCCGTCGCGCGAAGAAAGAGATGGTCGAAGC
CAACTTGCGCCTGGTGATCTCCATCGCCAAGAAGTACACCAACCGTGGCCTGCAGTTCCTCGACCTGATCCAGGAAGGCA
ACATCGGTTTGATGAAAGCGGTAGACAAGTTCGAATACCGTCGCGGCTACAAGTTCTCGACTTATGCCACCTGGTGGATC
CGTCAGGCGATCACTCGCTCGATCGCCGACCAGGCCCGCACCATCCGTATTCCGGTGCACATGATCGAGACGATCAACAA
GCTCAACCGTATTTCCCGGCAGATGTTGCAGGAAATGGGTCGCGAACCGACTCCGGAAGAGCTGGGCGAACGCATGGAAA
TGCCTGAGGACAAGATCCGCAAGGTATTGAAGATCGCTAAAGAGCCGATCTCCATGGAAACCCCGATCGGTGATGACGAA
GACTCCCATCTGGGTGACTTCATCGAAGACTCGACCATGCAGTCGCCAATCGATGTCGCCACCGTTGAGAGCCTCAAGGA
AGCGACTCGCGAAGTCCTCTCCGGCCTCACTGCCCGTGAAGCCAAGGTACTGCGCATGCGCTTCGGCATCGACATGAATA
CCGACCACACCCTTGAGGAAGTCGGTAAGCAGTTCGACGTTACCCGTGAGCGGATTCGTCAGATCGAAGCCAAGGCGCTG
CGCAAGCTGCGCCACCCGACGAGAAGCGAGCATCTGCGCTCCTTCCTCGACGAGTAATTGACTAAGCCAGCCATACTCGC
CCTTGCTGATGGCAGCATTTTTCGCGGCGAAGCCATTGGAGCCGACGGTCAGACCGTTGGTGAGGTGGTGTTTAACACCG
CAATGACCGGCTATCAGGAAATCCTTACCGATCCTTCCTACGCCCAACAGATCGTTACCCTGACTTACCCGCACATCGGC
AACACTGGCACCACGCCGGAAGACGCCGAGTCCGATCGTGTCTGGTCGGCCGGTCTGGTGATTCGCGACCTGCCACTGGT
TGCGAGCAACTGGCGTAACACCCTGTCCCTGTCCGATTACCTGAAAGCCAACAATGTCGTGGCGATCGCCGGTATCGACA
CCCGTCGCCTGACGCGCATCCTGCGCGAGAAAGGCGCACAGAACGGCTGCATCATGGCCGGCGACAACATCTCCGACGAA
GCGGCGATTGCTGCTGCACGCGGCTTCCCTGGCCTGAAAGGCATGGATCTGGCGAAGGTCGTCAGCACCAAGGAAAGCTA
CGAGTGGCGCTCCAGCGTCTGGAGCCTGAAGACCGACAGTCATCCGACTATCGAGGCTTCCGAGCTGCCTTACCACGTGG
TTGCCTACGACTACGGCGTCAAGCTGAACATCCTGCGCATGCTGGTCGAGCGCGGTTGCCGCGTGACCGTAGTGCCTGCG
CAAACCCCGGCCAGCGACGTCCTGGCACTCAAGCCTGACGGTGTGTTCCTGTCCAACGGTCCTGGCGACCCCGAGCCTTG
CGATTACGCCATCCAGGCGATCAAGGACGTGCTGGAAACCGAGATTCCGGTCTTCGGTATCTGCCTGGGCCACCAACTGC
TGGCACTGGCCTCCGGCGCCAAGACGGTGAAAATGGGCCACGGCCACCACGGCGCCAACCACCCGGTCCAGGACCTGGAC
AGCGGTGTGGTGATGATCACCAGCCAGAACCACGGTTTTGCGGTGGACGAAACCACCCTGCCGGGCAACGTGCGGGCGAT
CCACAAGTCGCTGTTCGATGGCACCCTGCAAGGCATCGAGCGTACCGACAAGAGCGCATTCAGCTTCCAGGGCCACCCTG
AAGCGAGCCCGGGCCCGAACGATGTGGCGCCGCTGTTCGATCGTTTCATCAACGAGATGGCCAAGCGACGCTGAATGAGT
AGCGGACGTATCGTTCAAATCATCGGCGCCGTTATCGACGTGGAATTTCCACGCGACAGCGTACCGAGCATCTACGACGC
CTTGAAGGTTCAAGGCGCCGAAACCACTCTGGAAGTTCAGCAGCAGCTGGGCGACGGCGTGGTACGTACCATTGCGATGG
GCTCCACCGAGGGCTTGAAGCGCGGTCTGGACGTCAACAACACTGGCGCCGCCATCTCCGTACCGGTCGGTAAAGCGACC
CTGGGCCGGATCATGGACGTGCTGGGCAACCCGATCGACGAAGCTGGCCCGATCGGCGAAGAAGAGCGTTGGGGCATTCA
CCGTCCTGCGCCGACCTTCGCTGAACAAGCTGGCGGCAACGACCTCCTGGAAACCGGCATCAAGGTTATCGACCTGGTTT
GCCCGTTCGCCAAGGGCGGTAAAGTCGGTCTGTTCGGTGGTGCCGGTGTGGGCAAAACCGTAAACATGATGGAACTGATC
CGTAACATCGCCATCGAGCACAGCGGTTATTCCGTGTTCGCTGGTGTGGGTGAGCGTACTCGTGAGGGTAACGACTTCTA
CCACGAGATGAAGGATTCCAACGTTCTGGACAAAGTGGCACTGGTATACGGCCAGATGAACGAGCCGCCGGGAAACCGTC
TGCGCGTAGCTCTGACCGGCCTGACCATGGCCGAGAAGTTCCGTGACGAAGGTAACGACGTTCTGCTGTTCGTCGACAAC
ATCTATCGTTACACCCTGGCCGGTACCGAAGTATCCGCACTGCTGGGCCGTATGCCTTCGGCAGTAGGTTACCAGCCGAC
CCTGGCTGAAGAGATGGGCGTTCTGCAAGAACGTATCACTTCGACCAAGCAAGGCTCGATCACCTCGATCCAAGCGGTAT
ACGTACCTGCGGACGACTTGACCGACCCGTCGCCAGCGACCACCTTCGCCCACTTGGACGCCACCGTCGTACTGTCCCGT
GACATCGCTTCCCTGGGTATCTACCCAGCGGTAGACCCACTGGATTCGACTTCCCGTCAGCTGGACCCGAACGTGATCGG
CAACGAGCACTACGAAACCGCTCGCGGCGTTCAGTACGTGCTGCAGCGCTACAAAGAGCTGAAGGACATCATTGCGATCC
TGGGTATGGACGAACTGTCCGAAGCCGACAAGCAACTGGTATCCCGCGCTCGTAAGATCCAGCGCTTCCTGTCGCAGCCG
TTCTTCGTGGCTGAAGTCTTCACTGGTTCTCCAGGCAAATACGTTTCCCTGAAAGACACCATCGCTGGCTTCAAAGGCAT
CCTCAACGGTGACTACGACCACCTGCCAGAACAAGCGTTCTACATGGTCGGCGGCATCGAAGAAGCGATCGAGAAAGCCA
AGAAACTGTAA

NCBI Reference Sequence: NZ_CP027719.1

Strain: P2
Chained genes: 16S rRNA, recA, gyrB, rpoD, carA, atpD
>NZ_CP027719_P2
GAACTGAAGAGTTTGATCATGGCTCAGATTGAACGCTGGCGGCAGGCCTAACACATGCAAGTCGAGCGGTAGAGAGAAGC
TTGCTTCTCTTGAGAGCGGCGGACGGGTGAGTAATGCCTAGGAATCTGCCTGGTAGTGGGGGATAACGTTCGGAAACGGA
CGCTAATACCGCATACGTCCTACGGGAGAAAGCAGGGGACCTTCGGGCCTTGCGCTATCAGATGAGCCTAGGTCGGATTA
GCTAGTTGGTGAGGTAATGGCTCACCAAGGCGACGATCCGTAACTGGTCTGAGAGGATGATCAGTCACACTGGAACTGAG
ACACGGTCCAGACTCCTACGGGAGGCAGCAGTGGGGAATATTGGACAATGGGCGAAAGCCTGATCCAGCCATGCCGCGTG
TGTGAAGAAGGTCTTCGGATTGTAAAGCACTTTAAGTTGGGAGGAAGGGTACTTACCTAATACGTGAGTATTTTGACGTT
ACCGACAGAATAAGCACCGGCTAACTCTGTGCCAGCAGCCGCGGTAATACAGAGGGTGCAAGCGTTAATCGGAATTACTG
GGCGTAAAGCGCGCGTAGGTGGTTCGTTAAGTTGGATGTGAAATCCCCGGGCTCAACCTGGGAACTGCATCCAAAACTGG
CGAGCTAGAGTATGGTAGAGGGTGGTGGAATTTCCTGTGTAGCGGTGAAATGCGTAGATATAGGAAGGAACACCAGTGGC
GAAGGCGACCACCTGGACTGATACTGACACTGAGGTGCGAAAGCGTGGGGAGCAAACAGGATTAGATACCCTGGTAGTCC
ACGCCGTAAACGATGTCAACTAGCCGTTGGGAGCCTTGAGCTCTTAGTGGCGCAGCTAACGCATTAAGTTGACCGCCTGG
GGAGTACGGCCGCAAGGTTAAAACTCAAATGAATTGACGGGGGCCCGCACAAGCGGTGGAGCATGTGGTTTAATTCGAAG
CAACGCGAAGAACCTTACCAGGCCTTGACATCCAATGAACTTTCCAGAGATGGATTGGTGCCTTCGGGAACATTGAGACA
GGTGCTGCATGGCTGTCGTCAGCTCGTGTCGTGAGATGTTGGGTTAAGTCCCGTAACGAGCGCAACCCTTGTCCTTAGTT
ACCAGCACGTCATGGTGGGCACTCTAAGGAGACTGCCGGTGACAAACCGGAGGAAGGTGGGGATGACGTCAAGTCATCAT
GGCCCTTACGGCCTGGGCTACACACGTGCTACAATGGTCGGTACAGAGGGTTGCCAAGCCGCGAGGTGGAGCTAATCCCA
TAAAACCGATCGTAGTCCGGATCGCAGTCTGCAACTCGACTGCGTGAAGTCGGAATCGCTAGTAATCGCGAATCAGAATG
TCGCGGTGAATACGTTCCCGGGCCTTGTACACACCGCCCGTCACACCATGGGAGTGGGTTGCACCAGAAGTAGCTAGTCT
AACCTTCGGGAGGACGGTTACCACGGTGTGATTCATGACTGGGGTGAAGTCGTAACAAGGTAGCCGTAGGGGAACCTGCG
GCTGGATCACCTCCTTAAATGGACGACAACAAGAAGAAAGCCTTGGCTGCGGCCCTGGGTCAGATCGAACGTCAATTCGG
CAAGGGTGCCGTAATGCGTATGGGCGATCACGACCGCCAGGCGATCCCGGCCATTTCCACTGGCTCTCTGGGTCTGGACA
TCGCACTCGGCATCGGCGGCCTGCCAAAAGGTCGTATTGTTGAAATCTACGGTCCGGAATCGTCCGGTAAAACCACCCTG
ACCCTGTCCGTGATTGCCCAGGCACAGAAGATGGGCGCCACCTGCGCCTTCGTCGACGCCGAGCACGCACTGGACCCGGA
ATACGCCGGCAAACTGGGGGTCAACGTTGACGACCTGCTGGTTTCCCAGCCGGACACCGGCGAACAGGCGCTGGAAATCA
CCGACATGCTGGTGCGCTCCAATGCCATCGACGTGATCGTGATCGACTCCGTGGCGGCACTGGTACCCAAGGCCGAGATC
GAAGGCGAGATGGGCGACATGCACGTGGGTCTGCAGGCCCGCCTGATGTCCCAGGCGCTGCGCAAGATCACCGGTAACAT
CAAGAACGCCAACTGCCTGGTGATCTTCATCAACCAGATCCGAATGAAAATCGGCGTGATGTTCGGCAGCCCGGAAACCA
CCACCGGCGGTAACGCGCTGAAGTTCTACGCTTCGGTTCGTCTGGACATCCGTCGTACTGGCGCGGTGAAGGAAGGCGAC
GAAGTCGTCGGTAGCGAAACCCGGGTCAAGATCGTCAAGAACAAGGTGGCTCCACCGTTCCGCCAGGCTGAATTCCAGAT
CCTGTACGGCAAGGGTATCTACCTGAACGGTGAGATCATCGATCTGGGCGTGCTGCACGGTTTCCTCGAGAAGTCCGGTG
CCTGGTACAGCTACCAGGGCAACAAGATCGGTCAGGGCAAGGCCAACTCGGCCAAGTTCCTGCAGGACAATCCGGAAATC
GGCAATGCCCTCGAGAAGCAGATTCGCGACAAGCTGCTGGCTCCAACCGCTGATGTCAAAGCTTCGCCGGTCAACGAGAC
CATCGATGACATGGCTGACGCGGATATCTGAATGAGCGAAGAAAACACGTACGACTCGAGCAGCATTAAAGTGCTGAAAG
GTTTGGATGCCGTACGCAAACGTCCCGGTATGTACATTGGTGACACCGACGATGGCAGCGGTCTGCACCATATGGTGTTC
GAGGTGGTCGATAACTCGATCGACGAAGCTCTGGCCGGCCATTGCGACGACATCAGCATCATCATCCACCCGGACGAATC
CATTACCGTGCGTGACAACGGTCGCGGCATCCCGGTAGACGTGCATAAAGAAGAAGGCGTTTCCGCGGCCGAGGTCATCA
TGACCGTACTGCACGCCGGCGGTAAGTTCGACGATAACTCCTACAAAGTATCCGGCGGTCTGCACGGTGTGGGTGTGTCG
GTAGTGAACGCCCTGTCCGAAGAACTGGTCCTGACCGTTCGCCGCAGCGGAAAGATCTGGGAACAGACCTACGTTCACGG
TGTGCCTCAGGCGCCTATGGCGATCGTCGGTGACAGCGAAACCACCGGTACCCAGATTCACTTCAAGGCGTCCAGCGAGA
CCTTCAAGAACATCCATTTCAGCTGGGACATCCTGGCCAAGCGGATTCGTGAACTGTCCTTCCTCAACTCCGGTGTCGGT
ATCGTTCTGAAGGACGAACGCAGTGGCAAGGAAGAGCTGTTCAAGTACGAAGGCGGCCTGCGTGCGTTCGTTGAATACCT
GAACACCAACAAGACCGCGGTCAACCAGGTGTTCCACTTCAATGTGCAGCGTGAAGATGGCATCGGCGTGGAAATCGCCC
TGCAGTGGAACGACAGCTTCAACGAAAACCTGCAGTGCTTCACCAACAACATTCCGCAGCGCGATGGCGGCACCCACTTG
GTGGGCTTCCGTTCGGCACTGACGCGTAACCTGAACAACTACATCGAACAGGAAGGTCTGGCGAAGAAGCACAAGGTCGC
CACCACCGGTGACGATGCCCGCGAAGGCCTGACCGCGATCATTTCGGTCAAGGTGCCGGATCCGAAGTTCAGCTCCCAGA
CCAAAGACAAGCTGGTGTCTTCCGAAGTGAAGACCGCGGTTGAACAGGAAATGGGCAAGTACTTCTCCGACTTCCTGCTG
GAAAACCCGAACGAAGCCAAGCTGGTGGTCGGCAAGATGCTCGACGCCGCCCGTGCCCGTGAAGCGGCGCGTAAGGCTCG
TGAGATGACCCGCCGTAAAGGCGCGCTGGATATCGCCGGCCTGCCGGGCAAACTGGCGGACTGCCAGGAAAAAGACCCTG
CCCTTTCCGAACTCTACCTGGTGGAAGGTGACTCTGCTGGCGGCTCCGCCAAGCAGGGACGCAACCGTAAGACCCAGGCG
ATTCTGCCGCTCAAGGGCAAGATCCTTAACGTCGAGAAAGCGCGCTTCGACAAGATGATTTCCTCGCAAGAGGTCGGCAC
CTTGATCACTGCACTCGGTTGCGGCATCGGCCGCGAAGAGTACAACATCGACAAGCTGCGTTATCACAACATCATCATCA
TGACCGACGCTGACGTCGACGGTTCGCACATCCGTACCCTGCTGCTGACCTTCTTCTTCCGTCAGCTGCCGGAGCTGATC
GAGCGTGGCTACATCTACATCGCTCAACCGCCGCTGTACAAGGTCAAGAAAGGCAAGCAAGAGCAATACATCAAAGACGA
CGACGCCATGGAAGAGTACATGACGCAGTCGGCCCTGGAAGATGCGAGCCTGCACCTGAACGAAGAAGCACCGGGTATTT
CCGGCGAGGCGCTGGAGCGCCTGGTGAACGACTTCCGCATGGTCATGAAAACCCTCAAGCGTCTGTCGCGCCTGTACCCT
CAGGAGCTGACCGAGCACTTCATCTACCTGCCGGCCGTGAGCCTGGAGCAACTCTCCGATCACGCGGCCATGCAGGATTG
GCTGGCCCAATATGAAGTCCGCCTGCGCACCGTCGAGAAGTCCGGCCTGGTCTACAAGGCCAGCCTGCGTGAAGACCGTG
AACGTAATGTCTGGCTGCCAGAGGTCGAACTGATCTCCCACGGCCTGTCGAACTACGTCACCTTCAACCGCGACTTCTTC
GGTAGCAATGACTACAAGACCGTCGTTACCCTCGGCGCTCAACTGAGCTCCCTGCTGGACGAAGGCGCTTATATTCAGCG
TGGCGAACGCAAGAAGGCGGTGACCGAGTTCAAGGAAGCCCTGGACTGGCTGATGACCGAAAGCACCAAGCGCCACACCA
TCCAGCGATACAAAGGTCTGGGCGAGATGAACCCGGATCAGCTGTGGGAAACCACCATGGACCCAAGCGTGCGCCGTATG

CTCAAGGTCACGATTGAAGATGCCATCGGCGCCGACCAGATCTTCAACACCCTGATGGGGATGCGGTCGAGCCTCGTCG
CGACTTCATCGAAAGCAACGCCCTGGCGGTATCCAATCTGGACTTCTGAATGTCCGGAAAAGCGCAACAGCAGTCTCGCC
TCAAAGAGTTGATCAGCCGTGGTCGTGAGCAGGGTTACCTGACTTACGCGGAGGTCAACGACCACCTGCCGGAGGATATT
TCAGATCCGGAACAGGTGGAAGACATCATCCGCATGATCAACGACATGGGGATCAACGTATTCGAGAGTGCTCCGGATGC
GGATGCCCTTTTGTTGGCCGAAGCCGATACCGACGAAGCAGCAGCTGAAGAAGCCGCCGCAGCGTTGGCGGCTGTGGAAA
CCGACATTGGTCGCACTACCGACCCCGTGCGTATGTACATGCGCGAAATGGGAACGGTAGAGCTGCTCACACGTGAAGGC
GAAATCGAAATCGCCAAGCGTATCGAAGAGGGCATCCGTGAAGTGATGGGCGCGATCGCGCACTTCCCTGGCACGGTTGA
GCACATCCTCTCCGAATACACTCGCGTCACCACCGAAGGTGGCCGCCTGTCCGACGTCCTGAGCGGTTACATCGACCCGG
ACGACGGCATTGCGCCGCCTGCCGCCGAAGTACCACCGCCTGTCGATGCCAAGGCTGCAAAAGCGGACGACGACACCGAC
GACGATGACGCCGAAGCCAGTGACGACGAAGAAGAAGCCGAAAGCGGTCCGGATCCGGTCATCGCAGCCCAGCGCTTTGG
CGCCGTTGCCGACCAGATGGAAATCACCCGCAAGGCGCTGAAGAAGCACGGTCGCGAACACAAGCAAGCCCTGGCCGAAA
TGCTGGCCCTGGCTGAACTGTTCATGCCGATCAAACTGGTTCCGAAGCAATTCGAAGGCCTGGTTGAACGTGTTCGTAGC
GCCCTGGATCGCCTGCGTCAGCAAGAGCGCGCGATCATGCAGCTCTGTGTTCGTGATGCCCGCATGCCACGCGCCGACTT
CCTGCGCCAGTTCCCTGGCAATGAAGTGGACGAAAGCTGGTCCGACGCGCTGGCCAAAGGCAAGGCCAAGTACGCCGAAG
CCATCGGCCGCCTGCAGCCGGACATCATCCGTTGCCAGCAGAAGCTGACCGCGCTCGAGACCGAGACCGGCCTGACGATC
GCCGAGATCAAGGACATCAACCGTCGCATGTCGATCGGCGAGGCCAAGGCCCGTCGCGCGAAGAAAGAGATGGTCGAAGC
CAACCTGCGTCTGGTGATCTCCATCGCCAAGAAGTACACCAACCGTGGCTTGCAATTCCTCGACCTGATCCAGGAAGGCA
ACATCGGTTTGATGAAAGCGGTAGACAAGTTCGAATACCGTCGCGGCTACAAATTCTCGACTTATGCCACCTGGTGGATC
CGTCAGGCGATCACTCGCTCGATCGCCGACCAGGCCCGCACCATCCGTATTCCGGTGCACATGATCGAGACGATCAACAA
GCTCAACCGTATTTCCCGGCAGATGTTGCAGGAAATGGGTCGCGAACCGACTCCGGAAGAGCTGGGCGAACGCATGGAAA
TGCCTGAGGACAAGATCCGCAAGGTATTGAAGATCGCTAAAGAGCCGATCTCCATGGAAACCCCGATCGGTGATGACGAA
GACTCCCATCTGGGTGACTTCATCGAAGACTCGACCATGCAGTCGCCAATCGATGTCGCCACCGTTGAGAGCCTTAAAGA
AGCGACTCGCGAAGTACTCTCCGGCCTCACTGCCCGTGAAGCCAAGGTACTGCGCATGCGCTTCGGCATCGACATGAATA
CCGACCACACCCTCGAGGAAGTCGGTAAGCAGTTCGACGTTACCCGTGAGCGGATTCGTCAGATCGAAGCCAAGGCGCTG
CGCAAGCTGCGCCACCCGACGCGAAGCGAGCACCTGCGCTCCTTCCTCGACGAGTAATTGACTAAGCCAGCCATACTCGC
CCTTGCTGATGGCAGCATTTTTCGCGGCGAAGCCATTGGAGCCGACGGTCAAACCGTTGGTGAGGTGGTGTTTAACACCG
CAATGACCGGCTATCAGGAAATCCTTACCGATCCTTCCTACGCCCAACAGATCGTTACCCTGACTTACCCGCATATCGGC
AATACCGGCACCACGCCGGAAGACGCCGAGTCCGATCGTGTCTGGTCGGCCGGTCTGGTGATTCGCGACCTGCCTCTGGT
TGCGAGCAACTGGCGTAACACCCTGTCCCTGTCCGACTACCTGAAAGCCAACAATGTCGTGGCGATCGCCGGTATCGACA
CCCGTCGCCTGACGCGCATCCTGCGCGAGAAAGGTGCGCAGAACGGCTGCATCATGGCCGGCGACAATATCTCCGACGAA
GCGGCGATTGCCGCTGCACGCGGCTTCCCGGGCCTGAAAGGCATGGATCTGGCGAAGGTCGTCAGTACCAAGGAAAGCTA
CGAGTGGCGCTCCAGTGTCTGGAACCTGAAGACCGACAGTCATCCGACCATCGAAGCTTCCGAGCTGCCTTACCACGTGG
TTGCCTACGACTACGGCGTCAAGCTGAACATCCTGCGCATGCTGGTCGAACGCGGTTGCCGCGTGACCGTGGTGCCTGCG
CAAACCCCGGCCAGCGAAGCTCTGGCGCTCAAGCCTGACGGTGTGTTCCTGTCCAACGGCCCTGGCGACCCCGAGCCTTG
CGATTACGCCATCCAGGCGATCAAGGACGTGCTGGAGACCGAGATTCCGGTCTTCGGTATCTGTCTGGGCCACCAACTGC
TGGCACTGGCCGCCGGCGCCAAGACAGTGAAGATGGGCCACGGCCACCACGGCGCCAACCACCCGGTCCAGGACCTGGAC
AGCGGTGTGGTGATGATCACCAGCCAGAACCACGGTTTTGCGGTGGACGAAGCCACCCTGCCGGGCAACGTGCGGGCGAT
CCACAAGTCGCTGTTCGACGGCACCCTGCAAGGCATCGAGCTGACCGACAAGAGCGCATTCAGCTTCCAGGGCCACCCTG
AAGCGAGCCCGGGCCCGAACGATGTGGCGCCGCTGTTCGATCGTTTCATCAACGAGATGGCCAAGCGACGCTGAATGAGT
AGCGGACGTATCGTTCAAATCATCGGCGCCGTTATCGACGTGGAATTTCCACGCGACAGCGTACCGAGCATCTACGACGC
CTTGAAGGTTCAAGGCGCCGAAACCACTCTGGAAGTTCAGCAGCAGCTGGGCGACGGCGTGGTACGTACCATTGCGATGG
GCTCCACCGAGGGCTTGAAGCGCGGTCTGGACGTCAACAACACTGGCGCAGCCATCTCCGTACCGGTCGGTAAAGCGACC
CTGGGCCGGATCATGGACGTACTGGGCAACCCGATCGACGAAGCTGGCCCGATCGGTGAAGAAGAGCGTTGGGGCATTCA
CCGTCCTGCGCCGACCTTCGCTGAACAAGCTGGCGGCAACGACCTGCTGGAAACCGGCATCAAGGTTATCGACCTGGTTT
GCCCGTTCGCCAAGGGCGGTAAAGTCGGTCTGTTCGGTGGTGCCGGTGTGGGCAAAACCGTAAACATGATGGAACTGATC
CGTAACATCGCCATCGAGCACAGCGGTTATTCCGTGTTCGCCGGTGTGGGTGAGCGTACTCGTGAGGGTAACGACTTCTA
CCACGAGATGAAGGATTCCAACGTTCTGGACAAAGTGGCACTGGTATACGGCCAGATGAACGAGCCGCCGGGAAACCGTC
TGCGCGTAGCTCTGACCGGCCTGACCATGGCCGAGAAGTTCCGTGACGAAGGTAACGACGTTCTGCTGTTCGTCGACAAC
ATCTATCGTTACACCCTGGCCGGTACCGAAGTATCCGCACTGCTGGGCCGTATGCCTTCGGCAGTAGGTTACCAGCCGAC
CCTGGCTGAAGAGATGGGCGTTCTGCAAGAACGTATCACTTCGACCAAGCAAGGCTCGATCACCTCGATCCAAGCGGTAT
ACGTACCTGCGGACGACTTGACCGACCCGTCGCCAGCGACCACCTTCGCCCACTTGGACGCCACCGTCGTTCTGTCCCGT
GACATCGCTTCCCTGGGTATCTACCCAGCGGTAGACCCACTGGACTCGACTTCCCGTCAGCTGGACCCGAACGTGATCGG
CACCGAGCACTACGAAACCGCTCGTGGCGTTCAGTACGTGCTGCAGCGCTACAAAGAGCTGAAGGACATCATTGCGATCC
TGGGTATGGACGAACTGTCCGAAGCCGACAAGCAACTGGTATCCCGCGCTCGTAAGATCCAGCGCTTCCTGTCGCAGCCG
TTCTTCGTGGCTGAAGTCTTCACTGGTTCTCCAGGCAAATACGTTTCCCTGAAAGACACCATCGCTGGCTTCAAAGGCAT
CCTCAACGGTGACTACGACCATCTGCCAGAACAAGCGTTCTACATGGTTGGTGGCATCGAAGAAGCGATCGAGAAAGCCA
AGAAACTGTAA

NCBI Reference Sequence: NZ_CP008696.1
Strain: PA23
Chained genes: 16S rRNA, recA, gyrB, rpoD, carA, atpD
>NZ_CP008696_PA23
AGAGTTTGATCATGGCTCAGATTGAACGCTGGCGGCAGGCCTAACACATGCAAGTCGAGCGGTAGAGAGAAGCTTGCTTC
TCTTGAGAGCGGCGGACGGGTGAGTAATGCCTAGGAATCTGCCTGGTAGTGGGGGATAACGTTCGGAAACGGACGCTAAT

ACCGCATACGTCCTACGGGAGAAAGCAGGGGACCTTCGGGCCTTGCGCTATCAGATGAGCCTAGGTCGGATTAGCTAGTT
GGTGAGGTAATGGCTCACCAAGGCGACGATCCGTAACTGGTCTGAGAGGATGATCAGTCACACTGGAACTGAGACACGGT
CCAGACTCCTACGGGAGGCAGCAGTGGGGAATATTGGACAATGGGCGAAAGCCTGATCCAGCCATGCCGCGTGTGTGAAG
AAGGTCTTCGGATTGTAAAGCACTTTAAGTTGGGAGGAAGGGTACTTACCTAATACGTGAGTATTTTGACGTTACCGACA
GAATAAGCACCGGCTAACTCTGTGCCAGCAGCCGCGGTAATACAGAGGGTGCAAGCGTTAATCGGAATTACTGGGCGTAA
AGCGCGCGTAGGTGGTTCGTTAAGTTGGATGTGAAATCCCCGGGCTCAACCTGGGAACTGCATCCAAAACTGGCGAGCTA
GAGTATGGTAGAGGGTGGTGGAATTTCCTGTGTAGCGGTGAAATGCGTAGATATAGGAAGGAACACCAGTGGCGAAGGCG
ACCACCTGGACTGATACTGACACTGAGGTGCGAAAGCGTGGGGAGCAAACAGGATTAGATACCCTGGTAGTCCACGCCGT
AAACGATGTCAACTAGCCGTTGGGAGCCTTGAGCTCTTAGTGGCGCAGCTAACGCATTAAGTTGACCGCCTGGGGAGTAC
GGCCGCAAGGTTAAAACTCAAATGAATTGACGGGGGCCCGCACAAGCGGTGGAGCATGTGGTTTAATTCGAAGCAACGCG
AAGAACCTTACCAGGCCTTGACATCCAATGAACTTTCCAGAGATGGATTGGTGCCTTCGGGAACATTGAGACAGGTGCTG
CATGGCTGTCGTCAGCTCGTGTCGTGAGATGTTGGGTTAAGTCCCGTAACGAGCGCAACCCTTGTCCTTAGTTACCAGCA
CGTAATGGTGGGCACTCTAAGGAGACTGCCGGTGACAAACCGGAGGAAGGTGGGGATGACGTCAAGTCATCATGGCCCTT
ACGGCCTGGGCTACACACGTGCTACAATGGTCGGTACAGAGGGTTGCCAAGCCGCGAGGTGGAGCTAATCCCATAAAACC
GATCGTAGTCCGGATCGCAGTCTGCAACTCGACTGCGTGAAGTCGGAATCGCTAGTAATCGCGAATCAGAATGTCGCGGT
GAATACGTTCCCGGGCCTTGTACACACCGCCCGTCACACCATGGGAGTGGGTTGCACCAGAAGTAGCTAGTCTAACCTTC
GGGAGGACGGTTACCACGGTGTGATTCATGACTGGGGTGAAGTCGTAACAAGGTAGCCGTAGGGGAACCTGCGGCTGGAT
CACCTCCTTAAATGGACGACAACAAGAAGAAAGCCTTGGCTGCGGCCCTGGGTCAGATCGAACGTCAATTCGGCAAGGGT
GCCGTAATGCGTATGGGCGATCACGACCGCCAGGCGATCCCGGCCATTTCCACTGGCTCTCTGGGTCTGGACATCGCACT
CGGCATCGGCGGCCTGCCAAAAGGTCGTATTGTTGAAATCTACGGTCCGGAATCGTCCGGTAAAACCACCCTGACCCTGT
CCGTGATTGCCCAGGCACAGAAGATGGGCGCCACCTGCGCCTTCGTCGACGCCGAGCACGCACTGGACCCGGAATACGCC
GGCAAACTGGGGGTCAACGTTGACGACCTGCTGGTTTCCCAGCCGGACACCGGCGAACAGGCGCTGGAAATCACCGACAT
GCTGGTGCGCTCCAATGCCATCGACGTGATCGTGATCGACTCCGTGGCGGCACTGGTACCCAAGGCCGAGATCGAAGGCG
AGATGGGCGACATGCACGTGGGCCTGCAGGCCCGCCTGATGTCCCAGGCGCTGCGCAAGATCACCGGTAACATCAAGAAC
GCCAACTGCCTGGTGATCTTCATCAACCAGATCCGTATGAAAATCGGCGTGATGTTCGGCAGCCCGGAAACCACCACCGG
CGGTAACGCGCTGAAGTTCTACGCTTCGGTTCGTCTGGACATCCGTCGTACTGGCGCGGTGAAGGAAGGCGACGAAGTCG
TCGGTAGCGAAACCCGGGTCAAGATCGTCAAGAACAAGGTGGCTCCACCGTTCCGCCAGGCTGAATTCCAGATCCTGTAC
GGCAAGGGTATCTACCTGAACGGTGAGATCATCGATCTGGGCGTGCTGCACGGTTTCCTCGAGAAGTCCGGTGCCTGGTA
CAGCTACCAGGGCAACAAGATCGGTCAGGGCAAGGCCAACTCGGCCAAGTTCCTGCAGGACAATCCGGAAATCGGCAATG
CCCTCGAGAAGCAGATTCGCGACAAGCTGCTGGCTCCAACCGCTGATGTCAAAGCTTCGCCGGTCAACGAGACCATCGAT
GACATGGCTGACGCGGATATCTGAATGAGCGAAGAAAACACGTACGACTCGAGCAGCATTAAAGTGCTGAAAGGTTTGGA
TGCCGTACGCAAACGTCCCGGTATGTACATTGGTGACACCGACGATGGCAGCGGTCTGCACCATATGGTGTTCGAGGTGG
TCGATAACTCGATCGACGAAGCTCTGGCCGGCCATTGCGACGACATCAGCATCATCATCCACCCGGACGAATCCATTACC
GTGCGTGACAACGGTCGCGGCATCCCGGTAGACGTGCATAAAGAAGAAGGCGTGTCCGCGGCCGAGGTCATCATGACCGT
GCTGCACGCTGGTGGTAAGTTCGACGACAACTCCTACAAAGTATCCGGCGGTCTGCACGGTGTGGGTGTGTCGGTAGTGA
ACGCCCTGTCCGAAGAACTGGTCCTGACCGTTCGCCGCAGCGGAAAGATCTGGGAACAGACCTACGTTCACGGTGTGCCT
CAGGCGCCTATGGCGATCGTCGGTGACAGCGAAACCACCGGTACCCAGATTCACTTCAAGGCGTCCAGCGAGACCTTCAA
GAACATCCATTTCAGCTGGGACATCCTGGCCAAGCGGATTCGTGAACTGTCCTTCCTCAACTCCGGTGTCGGTATCGTTC
TGAAGGACGAACGCAGCGGCAAGGAAGAGCTGTTCAAGTACGAAGGCGGCCTGCGTGCGTTCGTTGAATACCTGAACACC
AACAAGACCGCGGTCAACCAGGTGTTCCACTTCAATGTGCAGCGTGAAGATGGCATCGGCGTGGAAATCGCCCTGCAGTG
GAACGACAGCTTCAACGAAAACCTGCAGTGCTTCACCAACAACATTCCGCAGCGCGATGGCGGCACCCACTTGGTGGGCT
TCCGTTCGGCACTGACGCGTAACCTGAACAACTACATCGAACAGGAAGGTCTGGCGAAGAAGCACAAGGTCGCCACCACC
GGTGACGATGCCCGCGAAGGCCTGACCGCGATCATTTCGGTCAAGGTGCCGGATCCGAAGTTCAGCTCCCAGACCAAAGA
CAAGCTGGTGTCTTCCGAAGTGAAGACCGCGGTTGAACAGGAAATGGGCAAGTACTTCTCCGACTTCCTGCTGGAAAACC
CGAACGAAGCCAAGCTGGTGGTCGGCAAGATGCTCGACGCCGCCCGTGCCCGTGAAGCGGCGCGTAAGGCTCGTGAGATG
ACCCGCCGTAAAGGCGCGCTGGATATCGCCGGCCTGCCGGGCAAACTGGCGGACTGCCAGGAAAAAGACCCTGCCCTTTC
CGAACTCTACCTGGTGGAAGGTGACTCTGCTGGCGGCTCCGCCAAGCAGGGACGCAACCGTAAGACCCAGGCGATTCTGC
CGCTCAAGGGCAAGATCCTTAACGTCGAGAAAGCGCGCTTCGACAAGATGATTTCCTCGCAAGAGGTCGGCACCTTGATC
ACTGCACTCGGTTGCGGCATCGGCCGCGAAGAGTACAACATCGACAAGCTGCGTTATCACAACATCATCATCATGACCGA
CGCCGACGTCGACGGTTCGCACATCCGCACCCTGCTGCTGACCTTCTTCTTCCGTCAGCTGCCGGAGCTGATCGAGCGTG
GCTACATCTACATCGCTCAACCGCCGCTGTACAAGGTCAAGAAAGGCAAGCAAGAGCAATACATCAAAGACGACGACGCC
ATGGAAGAGTACATGACGCAGTCGGCCCTGGAAGATGCGAGCCTGCACCTGAACGAAGAAGCACCGGGTATTTCCGGCGA
GGCGCTGGAGCGCCTGGTGAACGACTTCCGCATGGTCATGAAAACCCTCAAGCGTCTGTCGCGCCTGTACCCTCAGGAGC
TGACCGAACACTTCATCTACCTGCCAGCCGTGAGCCTGGAGCAACTCTCCGATCACGCAGCGATGCAGGATTGGTTGGCC
CAATATGAAGTCCGCCTGCGCACCGTCGAGAAGTCCGGCCTGGTCTACAAGGCCAGCCTGCGTGAAGACCGTGAACGTAA
TGTCTGGCTGCCAGAGGTCGAACTGATCTCCCATGGCCTGTCGAACTACGTCACCTTCAACCGCGACTTCTTCGGCAGCA
ATGACTACAAGACCGTCGTCACCCTCGGCGCTCAACTGAGCTCCCTGCTGGACGAAGGCGCTTATATTCAGCGCGGCGAA
CGCAAGAAGGCGGTGACCGAGTTCAAGGAAGCCCTGGACTGGCTGATGACCGAAAGCACCAAGCGCCACACCATCCAGCG
ATACAAAGGTCTGGGCGAGATGAACCCGGACCAGCTGTGGGAAACCACCATGGATCCAAGCGTGCGCCGCATGCTCAAGG
TCACCATCGAAGACGCCATCGGCGCCGACCAGATCTTCAACACCCTGATGGGTGATGCGGTCGAGCCTCGTCGCGACTTC
ATCGAAAGCAACGCCCTGGCGGTATCCAACTTGGACTTCTGAATGTCCGGAAAAGCGCAACAGCAGTCTCGCCTCAAAGA
GTTGATCAGCCGTGGTCGTGAGCAGGGTTACCTGACTTACGCGGAGGTCAACGACCACCTGCCGGAGGATATTTCAGATC
CGGAACAGGTGGAAGACATCATCCGCATGATCAACGACATGGGGATCAACGTATTCGAGAGTGCTCCGGATGCGGATGCC
CTTTTGTTGGCCGAAGCCGATACCGACGAAGCAGCAGCTGAAGAAGCCGCCGCAGCGTTGGCGGCCGTGGAAACCGACAT

TGGTCGCACTACCGACCCCGTGCGTATGTACATGCGCGAAATGGGAACGGTAGAGCTGCTCACACGTGAAGGCGAAATCG
AAATCGCCAAGCGTATCGAAGAGGGCATCCGTGAAGTGATGGGCGCGATCGCGCACTTCCCTGGCACGGTTGAGCACATC
CTCTCCGAATACACTCGCGTCACCACCGAAGGTGGCCGCCTGTCCGACGTCCTGAGCGGTTACATCGACCCGGACGACGG
CATCGCGCCGCCTGCCGCCGAAGTACCACCGCCTGTCGATGCCAAGGCCGCAAAAGCGGACGACGACACCGACGACGATG
ACGCCGAAGCCAGTGACGACGAAGAAGAAGCCGAAAGCGGTCCGGATCCGGTCATCGCAGCCCAGCGCTTTGGCGCCGTT
GCCGACCAGATGGAAATCACCCGCAAGGCGCTGAAGAAGCACGGTCGCGAACACAAGCAAGCCCTGGCTGAAATGCTGGC
CCTGGCTGAACTGTTCATGCCGATCAAACTGGTTCCGAAGCAATTCGAAGGCCTGGTTGAACGTGTTCGTAGCGCCCTGG
ATCGCCTGCGTCAGCAAGAGCGCGCGATCATGCAGCTCTGTGTTCGTGATGCCCGCATGCCACGCGCCGACTTCCTGCGC
CAGTTCCCTGGCAATGAAGTGGACGAAAGCTGGTCCGACGCGCTGGCCAAAGGCAAGGCCAAGTACGCCGAAGCCATCGG
CCGCCTGCAGCCGGACATTATCCGTTGCCAGCAGAAGCTGACCGCGCTTGAGACCGAGACCGGCCTGACGATCGCCGAGA
TCAAGGACATCAACCGTCGCATGTCGATCGGCGAGGCCAAGGCCCGTCGCGCGAAGAAAGAGATGGTCGAAGCCAACCTG
CGTCTGGTGATCTCCATCGCCAAGAAGTACACCAACCGTGGCTTGCAATTCCTCGACCTGATCCAGGAAGGCAACATCGG
TTTGATGAAAGCGGTAGACAAGTTCGAATACCGTCGCGGCTACAAATTCTCGACTTATGCCACCTGGTGGATCCGTCAGG
CGATCACTCGCTCGATCGCCGACCAGGCCCGCACCATCCGTATTCCGGTGCACATGATCGAGACGATCAACAAGCTCAAC
CGTATTTCCCGGCAGATGTTGCAGGAAATGGGTCGCGAACCGACTCCGGAAGAGCTGGGCGAACGCATGGAAATGCCTGA
GGACAAGATCCGCAAGGTATTGAAGATCGCTAAAGAGCCGATCTCCATGGAAACCCCGATCGGTGATGACGAAGACTCCC
ATCTGGGTGACTTCATCGAAGACTCGACCATGCAGTCGCCAATCGATGTCGCCACCGTCGAGAGTCTTAAAGAAGCGACT
CGCGAAGTACTCTCCGGCCTCACTGCCCGTGAAGCCAAGGTACTGCGCATGCGCTTCGGCATCGACATGAATACCGACCA
CACCCTCGAGGAAGTCGGTAAGCAGTTTGACGTTACCCGTGAGCGGATTCGTCAGATCGAAGCCAAGGCGCTGCGCAAGC
TGCGCCACCCGACGCGAAGCGAGCACCTGCGCTCCTTCCTCGACGAGTAATTGACTAAGCCAGCCATACTCGCCCTTGCT
GATGGCAGCATTTTTCGCGGCGAAGCCATTGGAGCCGACGGTCAAACCGTTGGTGAGGTGGTGTTTAACACCGCAATGAC
CGGCTATCAGGAAATCCTTACCGATCCTTCCTACGCCCAACAGATCGTTACCCTGACTTACCCGCATATCGGCAATACCG
GCACCACGCCGGAAGACGCCGAGTCCGATCGTGTCTGGTCGGCCGGTCTGGTGATTCGCGACCTGCCTCTGGTTGCGAGC
AACTGGCGTAACACCCTGTCCCTGTCCGACTACCTGAAAGCCAACAATGTCGTGGCAATCGCCGGTATCGACACCCGTCG
CCTGACGCGCATCCTGCGCGAGAAAGGTGCGCAGAACGGCTGCATCATGGCCGGCGACAATATCTCCGACGAAGCGGCGA
TTGCCGCTGCACGCGGCTTCCCGGGCCTGAAAGGCATGGATCTGGCGAAGGTCGTCAGTACCAAGGAAAGCTACGAGTGG
CGCTCCAGTGTCTGGAACCTGAAGACCGACAGTCATCCGACCATCGAAGCTTCCGAGCTGCCTTACCACGTGGTTGCCTA
CGACTACGGCGTCAAGCTGAACATCCTGCGCATGCTGGTCGAACGCGGTTGCCGCGTGACCGTGGTGCCTGCGCAAACCC
CGGCCAGCGAAGCTCTGGCGCTCAAGCCTGACGGTGTGTTCCTGTCCAACGGCCCTGGCGACCCCGAGCCTTGCGATTAC
GCCATCCAGGCGATCAAGGACGTGCTGGAGACCGAGATTCCGGTCTTCGGTATCTGTCTGGGCCACCAACTGCTGGCACT
GGCCGCCGGCGCCAAGACAGTGAAGATGGGCCACGGCCACCACGGCGCCAACCACCCGGTCCAGGACCTGGACAGCGGTG
TGGTGATGATCACCAGCCAGAACCACGGTTTTGCGGTGGACGAAGCCACCCTGCCGGGCAACGTGCGGGCGATCCACAAG
TCGCTGTTCGACGGCACCCTGCAAGGCATCGAGCTGATCGACAAGAGCGCATTCAGCTTCCAGGGCCACCCTGAAGCGAG
CCCGGGCCCGAACGATGTGGCGCCGCTGTTCGATCGTTTCATCAACGAGATGGCCAAGCGACGCTGAATGAGTAGCGGAC
GTATCGTTCAAATCATCGGCGCCGTTATCGACGTGGAATTTCCACGCGACAGCGTACCGAGCATCTACGACGCCTTGAAG
GTTCAAGGCGCCGAAACCACTCTGGAAGTTCAGCAGCAGCTGGGCGACGGCGTGGTACGTACCATTGCGATGGGCTCCAC
CGAGGGCTTGAAGCGCGGTCTGGACGTCAACAACACTGGCGCAGCCATCTCCGTACCGGTCGGTAAAGCGACCCTGGGCC
GGATCATGGACGTACTGGGCAACCCGATCGACGAAGCTGGCCCGATCGGCGAAGAAGAGCGTTGGGGCATTCACCGTCCT
GCGCCGACCTTCGCTGAACAAGCTGGCGGCAACGACCTGCTGGAAACCGGCATCAAGGTTATCGACCTGGTTTGCCCGTT
CGCCAAGGGCGGTAAAGTCGGTCTGTTCGGTGGTGCCGGTGTGGGCAAAACCGTAAACATGATGGAACTGATCCGTAACA
TCGCCATCGAGCACAGCGGTTATTCCGTGTTCGCCGGTGTGGGTGAGCGTACTCGTGAGGGTAACGACTTCTACCACGAG
ATGAAGGATTCCAACGTTCTGGACAAAGTGGCACTGGTATACGGCCAGATGAACGAGCCGCCGGGAAACCGTCTGCGCGT
AGCTCTGACCGGCCTGACCATGGCCGAGAAGTTCCGTGACGAAGGTAACGACGTTCTGCTGTTCGTCGACAACATCTATC
GTTACACCCTGGCCGGTACCGAAGTATCCGCACTGCTGGGCCGTATGCCTTCGGCAGTAGGTTACCAGCCGACCCTGGCT
GAAGAGATGGGCGTTCTGCAAGAACGTATCACTTCGACCAAGCAAGGCTCGATCACCTCGATCCAAGCGGTATACGTACC
TGCGGACGACTTGACCGACCCGTCGCCAGCGACCACCTTCGCCCACTTGGACGCCACCGTCGTTCTGTCCCGTGACATCG
CTTCCCTGGGTATCTACCCAGCGGTAGACCCACTGGACTCGACTTCCCGCCAGCTGGACCCGAACGTGATCGGCACCGAG
CACTACGAAACCGCTCGTGGCGTTCAGTACGTGCTGCAGCGCTACAAAGAGCTGAAGGACATCATTGCGATCCTGGGTAT
GGACGAACTGTCCGAAGCCGACAAGCAACTGGTATCCCGCGCTCGTAAGATCCAGCGCTTCCTGTCGCAGCCGTTCTTCG
TGGCTGAAGTCTTCACTGGTTCTCCAGGCAAATACGTTTCCCTGAAAGACACCATCGCTGGCTTCAAAGGCATCCTCAAC
GGTGACTACGACCATCTGCCAGAACAAGCGTTCTACATGGTTGGTGGCATCGAAGAAGCGATCGAGAAAGCCAAGAAACT
GTAA

NCBI Reference Sequence: NC_007492.2
Strain: Pf01
Chained genes: 16S rRNA, recA, gyrB, rpoD, carA, atpD
>NC_007492_Pf01
GAACTGAAGAGTTTGATCATGGCTCAGATTGAACGCTGGCGGCAGGCCTAACACATGCAAGTCGAGCGGATGAAGGGAGC
TTGCTCCTGGATTCAGCGGCGGACGGGTGAGTAATGCCTAGGAATCTGCCTGGTAGTGGGGGACAACGTTTCGAAAGGAA
CGCTAATACCGCATACGTCCTACGGGAGAAAGCAGGGGACCTTCGGGCCTTGCGCTATCAGATGAGCCTAGGTCGGATTA
GCTAGTTGGTGAGGTAATGGCTCACCAAGGCGACGATCCGTAACTGGTCTGAGAGGATGATCAGTCACACTGGAACTGAG
ACACGGTCCAGACTCCTACGGGAGGCAGCAGTGGGGAATATTGGACAATGGGCGAAAGCCTGATCCAGCCATGCCGCGTG
TGTGAAGAAGGTCTTCGGATTGTAAAGCACTTTAAGTTGGGAGGAAGGGTTGTAGATTAATACTCTGCAATTTTGACGTT
ACCGACAGAATAAGCACCGGCTAACTCTGTGCCAGCAGCCGCGGTAATACAGAGGGTGCAAGCGTTAATCGGAATTACTG

GGCGTAAAGCGCGCGTAGGTGGTTCGTTAAGTTGGATGTGAAATCCCCGGGCTCAACCTGGGAACTGCATCCAAAACTGG
CGAGCTAGAGTATGGTAGAGGGTGGTGGAATTTCCTGTGTAGCGGTGAAATGCGTAGATATAGGAAGGAACACCAGTGGC
GAAGGCGACCACCTGGACTGATACTGACACTGAGGTGCGAAAGCGTGGGGAGCAAACAGGATTAGATACCCTGGTAGTCC
ACGCCGTAAACGATGTCAACTAGCCGTTGGGAGCCTTGAGCTCTTAGTGGCGCAGCTAACGCATTAAGTTGACCGCCTGG
GGAGTACGGCCGCAAGGTTAAAACTCAAATGAATTGACGGGGGCCCGCACAAGCGGTGGAGCATGTGGTTTAATTCGAAG
CAACGCGAAGAACCTTACCAGGCCTTGACATCCAATGAACTTTCCAGAGATGGATTGGTGCCTTCGGGAGCATTGAGACA
GGTGCTGCATGGCTGTCGTCAGCTCGTGTCGTGAGATGTTGGGTTAAGTCCCGTAACGAGCGCAACCCTTGTCCTTAGTT
ACCAGCACGTTATGGTGGGCACTCTAAGGAGACTGCCGGTGACAAACCGGAGGAAGGTGGGGATGACGTCAAGTCATCAT
GGCCCTTACGGCCTGGGCTACACACGTGCTACAATGGTCGGTACAAAGGGTTGCCAAGCCGCGAGGTGGAGCTAATCCCA
TAAAACCGATCGTAGTCCGGATCGCAGTCTGCAACTCGACTGCGTGAAGTCGGAATCGCTAGTAATCGCGAATCAGAATG
TCGCGGTGAATACGTTCCCGGGCCTTGTACACACCGCCCGTCACACCATGGGAGTGGGTTGCACCAGAAGTAGCTAGTCT
AACCTTCGGGAGGACGGTTACCACGGTGTGATTCATGACTGGGGTGAAGTCGTAACAAGGTAGCCGTAGGGGAACCTGCG
GCTGGATCACCTCCTTAAATGGACGACAACAAGAAGAAAGCCTTGGCTGCGGCCCTGGGTCAGATCGAACGTCAATTCGG
CAAGGGTGCCGTAATGCGTATGGGCGATCAGGACCGTCAGGCGATCCCGGCCATTTCCACCGGCTCTCTGGGTCTGGACA
TCGCACTCGGCATCGGCGGCCTGCCAAAAGGCCGTATCGTTGAAATCTACGGTCCTGAATCTTCCGGTAAAACCACACTG
ACGCTGTCCGTGATCGCCCAGGCTCAAAAAGCCGGTGCGACCTGCGCCTTCGTCGACGCCGAACACGCCCTCGACCCAGA
GTACGCCGGCAAGCTGGGCGTCAATGTCGACGACCTGCTGGTTTCCCAGCCGGACACCGGCGAGCAGGCCCTGGAAATCA
CCGACATGCTGGTGCGCTCCAACGCCGTTGACGTGATCATCGTCGACTCCGTGGCCGCTCTGGTACCGAAGGCAGAAATC
GAAGGCGAAATGGGTGACATGCACGTGGGCCTGCAAGCCCGTCTGATGTCCCAGGCGCTGCGTAAAATCACCGGTAACAT
CAAGAACGCCAACTGCCTGGTGATCTTCATCAACCAGATCCGCATGAAGATCGGCGTGATGTTCGGCAGCCCGGAAACCA
CCACCGGTGGTAACGCGCTGAAGTTCTACGCCTCGGTTCGTCTCGACATCCGCCGTACCGGCGCGGTGAAGGAAGGCGAC
GAAGTGGTCGGCAGCGAAACCCGCGTCAAGGTTGTGAAGAACAAGGTGGCTTCGCCGTTCCGTCAGGCCGAGTTCCAGAT
TCTCTACGGCAAGGGTATCTACCTGAACGGCGAGATGATCGACCTGGGCGTTCTGCACGGGTTCGTCGAGAAGTCCGGCG
CCTGGTATGCCTACGAAGGCACCAAGATCGGTCAGGGCAAGGCCAACTCGGCCAAGTTCCTGGCGGACAACCCGGAAGTC
GCGGCCAAGCTCGAGAAGCAACTGCGTGACAAGCTGCTGTCGCCAGCCGTGATCGCCGACTCCAAGGCTTCTGCGGTCAA
AGAGACCGAAGACGACCTGGCTGACGCTGACATCTGATTGAGCGAAGAAAATACGTACGACTCATCGAGCATTAAAGTGC
TGAAAGGCCTGGATGCCGTGCGCAAACGTCCCGGTATGTACATTGGTGACACCGACGATGGCAGCGGTCTGCACCACATG
GTGTTCGAGGTGGTCGACAACTCGATCGACGAAGCCCTCGCCGGCCACTGCGACGACATCAGCATCATCATCCACCCGGA
TGAGTCCATCACCGTTAAAGACAACGGCCGTGGCATCCCGGTAGACGTGCACAAAGAGGAAGGCGTTTCCGCCGCCGAGG
TCATCATGACCGTCCTCCACGCCGGCGGTAAGTTTGACGACAACTCCTACAAGGTATCCGGTGGTCTGCACGGTGTAGGT
GTTTCGGTCGTGAACGCGCTGTCTGAAGAACTGGTCCTGACCGTGCGCCGCAGCGGCAAGATCTGGGAACAGACCTACGT
CCACGGCGTGCCTCAGGCACCGATGGCGATCGTTGGCGACAGCGAAACCACTGGCACCCAGATTCACTTCAAGGCTTCCA
GCGAAACCTTCAAGAACATTCACTTCAGCTGGGATATCCTGGCCAAGCGCATTCGTGAACTGTCTTTCCTCAACTCCGGT
GTAGGCATCGTCCTCAAGGACGAGCGCAGCGGCAAGGAAGAACTGTTCAAGTACGAAGGCGGCCTGCGTGCGTTCGTTGA
ATACCTGAACACCAACAAGACTGCGGTCAACCAGGTGTTCCACTTCAACATCCAGCGTGAAGACGGCATCGGCGTGGAAA
TCGCCCTGCAGTGGAACGACAGCTTCAACGAGAACCTGTTGTGCTTCACCAACAACATTCCGCAGCGCGACGGTGGCACT
CACCTGGTGGGCTTCCGCTCGGCACTGACGCGTAACCTGAACAACTACATCGAGCAGGAAGGCCTGGCGAAGAAGCACAA
AGTCGCCACCACCGGTGACGATGCCCGTGAAGGCCTGACCGCGATCATCTCGGTGAAGGTGCCGGATCCGAAGTTCAGCT
CCCAGACCAAAGACAAGCTGGTTTCTTCCGAAGTGAAGACCGCGGTCGAACAGGAAATGGGCAAGTACTTCTCCGACTTC
CTGCTGGAAAACCCGAACGAAGCCAAACTGGTCGTCGGCAAGATGATCGACGCTGCCCGTGCCCGTGAAGCGGCGCGCAA
GGCCCGTGAGATGACCCGCCGCAAAGGCGCGCTGGACATCGCCGGCCTGCCGGGCAAGCTCGCTGACTGCCAGGAAAAAG
ACCCGGCGCTGTCCGAACTGTACCTCGTGGAAGGTGACTCCGCGGGCGGCTCTGCCAAGCAGGGACGTAACCGCAAGACC
CAGGCCATCCTGCCGCTGAAGGGCAAGATCCTCAACGTCGAGAAAGCCCGTTTCGACAAGATGATCTCGTCCCAGGAAGT
GGGCACCCTGATCACCGCGCTGGGCTGCGGTATCGGTCGCGAAGAGTACAACATCGACAAGCTGCGCTATCACAACATCA
TCATCATGACCGATGCTGACGTCGACGGTTCGCACATCCGTACCCTGCTGCTGACCTTCTTCTTCCGTCAGTTGCCTGAG
CTGATCGAGCGTGGCTACATCTACATCGCCCAGCCGCCGCTGTACAAGGTCAAGAAAGGCAAGCAAGAGCAATACATCAA
AGACGACGACGCCATGGAAGAGTACATGACGCAGTCGGCCCTGGAAGATGCGAGCCTGCACCTGAACGAAGACGCCCCGG
GCATTTCCGGCGAAGCGCTGGAACGTCTGGTGAACGACTTCCGCATGGTGATGAAGACCCTCAAGCGTCTGTCGCGCCTG
TACCCGCAGGAGCTGACCGAGCACTTCATCTACCTGCCGGCCGTCAGCCTGGAAATGCTCGGCGACCACGCGAAGATGCA
GGACTGGCTGGCCCAGTACGAAGTCCGTCTGCGCACCGTCGAGAAGTCGGGCCTGGTCTACAAGGCCAGCCTGCGTGAAG
ATCGTGAACGTGGCGTTTGGCTGCCAGAGGTCGAACTGATCTCCCACGGTCTGTCGAACTATGTCACCTTTAACCGCGAC
TTCTTCGGCAGCAACGACTACAAGACCGTCGTCACCCTCGGCGCTCAACTGAGCACCCTTCTTGATGAAGGCGCATACAT
CCAGCGTGGCGAGCGTAAAAAAGCGGTCACTGAGTTCAAGGAAGCCCTGGACTGGCTGATGGCCGAAAGCACCAAGCGCC
ACACCATTCAGCGATACAAAGGTCTGGGCGAAATGAACCCGGATCAGCTGTGGGAAACCACCATGGACCCAAGCGTGCGC
CGCATGCTGAAAGTCACCATCGAAGACGCCATTGGCGCAGACCAGATCTTCAACACCCTGATGGGTGATGCGGTCGAACC
TCGCCGTGACTTCATCGAGAGCAACGCTCTGGCGGTGTCCAACCTGGATTTCTGAATGTCCGGAAAAGCGCAACAGCAGT
CTCGTATTATTGAGTTGATCAAACTGGGTCGTGAGCAGAAGTATCTGACTTACGCCGAGGTCAACGACCACCTGCCCGAG
GATATTTCAGATCCGGAGCAGGTGGAAGACATCATCCGCATGATTAACGACATGGGGATCCCCGTACACGAGAGTGCTCC
GGATGCGGACGCCCTTATGCTGGCCGACGCCGATACCGACGAGGCCGCTGCGGAAGAAGCAGCCGCTGCGTTGGCGGCGG
TGGAGACCGATATCGGTCGCACCACTGACCCGGTGCGCATGTACATGCGTGAAATGGGTACGGTCGAGCTTCTGACTCGT
GAAGGCGAAATCGAAATCGCCAAGCGTATCGAAGAAGGCATCCGTGAAGTGATGAGCGCCATCGCGCACTTCCCTGGCAC
GGTTGACCATATTCTCTCCGAGTACACTCGCGTCACCACCGAAGGTGGTCGCCTGTCCGACGTTCTGAGCGGTTACATCG
ACCCGGACGACGGCATTACGCCGCCTGCCGCCGAAGTACCGCCACCGATCGACGCGAAAGCCGCGAAAGCGGATGACGAC
TCCGAGGACGATGACGCCGAAGCTTCCGATGACGAAGAAGAAGCCGAAAGCGGTCCGGATCCGGTCATTGCCGCACAGCG

CTTCGGTGCTGTCGCCGACCAGATGGAAATCACCCGCAAGGCCCTGAAAAAGCACGGTCGTCACAACAAGGCGGCAATTG
CCGAACTGTTGGCCCTGGCCGAGCTGTTCATGCCGATCAAGCTGGTGCCGAAGCAGTTCGAAGCCCTGGTCGAGCGTGTT
CGCAGCGCCCTGGATCGCCTGCGTCAGCAAGAGCGCGCGATCATGCAACTGTGCGTACGTGATGCACGTATGCCTCGTGC
CGACTTCCTGCGCCAGTTCCCGGGCAACGAAGTCGACGAAAGCTGGTCCGACGCCCTGGCCAAAGGCAAGAGCAAGTACG
CCGAAGCCATCGCCCGCGTGCAACCGGACATCATCCGTTGCCAGCAAAAGCTGACCGCGCTGGAAACCGAGACCGGTTTG
ACCATCGCCGAGATCAAGGACATCAACCGTCGCATGTCGATCGGTGAGGCGAAAGCCCGCCGCGCGAAGAAAGAGATGGT
TGAAGCGAACTTGCGTCTGGTGATCTCCATCGCCAAGAAGTACACCAACCGTGGCCTGCAATTCCTCGACCTGATCCAGG
AAGGCAACATCGGTCTGATGAAAGCGGTGGACAAGTTCGAATACCGTCGTGGTTACAAGTTCTCGACTTACGCCACCTGG
TGGATCCGTCAGGCGATCACTCGCTCGATCGCCGACCAGGCCCGCACCATCCGTATTCCGGTGCACATGATCGAGACGAT
CAACAAGCTCAACCGTATTTCCCGGCAGATGCTGCAGGAAATGGGTCGCGAACCGACCCCGGAAGAGCTGGGTGAACGCA
TGGAAATGCCTGAGGACAAGATCCGCAAGGTATTGAAGATCGCCAAAGAGCCGATCTCCATGGAAACCCCGATCGGTGAT
GACGAAGACTCTCATCTGGGTGACTTCATCGAAGACTCGACCATGCAGTCGCCAATCGATGTCGCCACTGTCGAGAGCCT
GAAAGAAGCGACCCGCGAAGTGCTGTCCGGCCTTACTGCCCGTGAAGCCAAGGTACTGCGCATGCGTTTCGGTATCGACA
TGAACACCGACCATACGCTTGAAGAAGTCGGCAAACAGTTTGACGTGACCCGCGAGCGGATCCGTCAGATCGAAGCCAAG
GCGCTGCGCAAGCTGCGCCACCCGACGCGAAGCGAGCATCTGCGCTCCTTCCTCGACGAGTGATTGACTAAGCCAGCCAT
ACTCGCCCTTGCTGATGGCAGCATTTTTCGCGGCGAAGCCATTGGAGCCGACGGTCAGACCGTTGGTGAGGTGGTGTTCA
ACACCGCAATGACCGGCTATCAGGAAATCCTTACCGATCCTTCCTACGCCCAACAGATCGTTACCCTGACTTACCCGCAC
ATCGGCAACACCGGCACCACGCCGGAAGACGCCGAGTCCGATCGCGTCTGGTCCGCTGGCCTGGTCATTCGTGACCTGCC
GCTGGTAGCGAGCAACTGGCGTAACACGATGTCCCTGTCCGATTACCTGAAAGCCAACAATGTTGTGGCGATCGCCGGTA
TCGACACCCGCCGCCTGACCCGCATCCTGCGTGAAAAAGGCGCACAGAACGGCTGCATCATGGCCGGCGACAACATCTCC
GAAGAGGCGGCCATCGCCGCGGCGCAAGGCTTCCCGGGCCTGAAGGGCATGGATCTGGCGAAAGTCGTCAGCACCAAGAC
CCAATACGAATGGCGCTCCACTGTCTGGGATCTGAAAACCGACAGCCACGCGACCATCGAAGCCTCCGAGCTGCCTTACC
ACGTGGTTGCCTACGACTACGGCGTCAAGGTCAACATCCTGCGCATGTTGGTCGAGCGCGGCTGCCGCGTCACTGTCGTT
CCGGCACAGACCCCGGCGGCCGACGTGCTGGCCTTGAAGCCGGACGGCGTGTTCCTGTCCAACGGTCCTGGTGATCCGGA
GCCTTGCGACTACGCGATCCAAGCGATCAAGGAAGTGCTGGAAACCGAAATTCCAGTCTTCGGCATCTGCCTCGGTCACC
AGCTGCTGGCTCTGGCCTCCGGCGCCAAGACCCTGAAAATGGGCCACGGCCACCACGGTGCCAACCACCCGGTGCAGGAT
CTGGACACTGGCGTCGTGATGATCACCAGCCAGAACCACGGTTTCGCGGTTGACGAAGAAACCCTGCCAGCCAACGTCCG
CGCGATCCATAAATCGCTGTTCGACGGCACCCTGCAAGGCATCGAGCGCACCGACAAGAGCGCGTTCAGCTTCCAGGGTC
ACCCTGAGGCGAGCCCGGGCCCGAACGATGTGGCCCCTCTGTTTGACCGCTTCATCAACGAGATGGCCAAGCGACGCTAA
ATGAGTAGCGGACGTATCGTTCAAATCATCGGCGCCGTTATCGACGTGGAATTTCCACGCGACAGCGTACCGAGCATCTA
CGACGCCTTGAAGGTTCAAGGCGCCGAAACCACTCTGGAAGTTCAGCAGCAGCTGGGCGACGGCGTGGTTCGTACCATTG
CGATGGGCTCCACCGAAGGCTTGAAGCGCGGTCTGGACGTCAACAACACTGGCGCAGCCATCTCCGTACCGGTCGGTAAA
GCGACTCTGGGCCGGATCATGGACGTACTGGGCAACCCGATCGACGAAGCTGGCCCGATCGGCGAAGAAGAGCGCTGGGG
TATCCACCGCGCCGCTCCTTCCTTCGCTGAACAAGCCGGTGGCAACGAGCTGCTGGAAACAGGCATCAAGGTTATCGACC
TGGTTTGCCCGTTCGCCAAGGGCGGTAAAGTCGGTCTGTTCGGTGGTGCCGGTGTAGGCAAGACCGTAAACATGATGGAA
CTGATCCGTAACATCGCCATCGAGCACAGCGGTTATTCCGTGTTCGCCGGTGTGGGTGAGCGTACTCGTGAGGGTAACGA
CTTCTACCACGAGATGAAGGACTCCAACGTTCTCGACAAGGTAGCCCTGGTCTACGGTCAGATGAACGAGCCACCGGGAA
ACCGTCTGCGCGTAGCGCTGACCGGTCTGACCATGGCCGAGAAGTTCCGTGACGAAGGTAACGACGTTCTGCTGTTCGTC
GACAACATCTATCGTTACACCCTGGCCGGTACCGAAGTATCCGCACTGCTGGGCCGTATGCCTTCGGCAGTAGGTTACCA
GCCGACCCTGGCTGAAGAGATGGGCGTGCTGCAAGAGCGCATCACTTCGACCAAGCAAGGTTCGATTACTTCGATCCAGG
CCGTATACGTACCAGCGGACGACTTGACTGACCCGTCGCCAGCGACCACGTTTGCTCACCTGGACGCCACCGTCGTTCTG
TCCCGTGACATCGCTTCTCTGGGTATCTACCCGGCGGTAGACCCACTGGACTCGACTTCGCGTCAGCTGGACCCGAACGT
GATCGGCAACGATCACTACGAGACCGCTCGTGGTGTTCAGTACGTGCTGCAGCGTTACAAAGAGCTGAAGGACATCATCG
CGATCCTGGGTATGGACGAGCTGTCGGAAGCCGACAAGCAGTTGGTAAACCGTGCTCGTAAGATCCAGCGCTTCTTGTCG
CAGCCGTTCTTCGTGGCTGAAGTCTTCACCGGTGCTTCGGGTAAATACGTTTCCCTGAAAGACACCATTGCTGGCTTCAA
AGGCATCCTCAACGGTGACTACGACCACCTGCCAGAACAAGCGTTCTACATGGTCGGCGGCATCGAAGAAGCGATCGAGA
AAGCCAAGAAACTGTAA

NCBI Reference Sequence: NZ_CP048051.1
Strain: zm-1
Chained genes: 16S rRNA, recA, gyrB, rpoD, carA, atpD
>NZ_CP048051_zm1
GAACTGAAGAGTTTGATCATGGCTCAGATTGAACGCTGGCGGCAGGCCTAACACATGCAAGTCGAGCGGTAGAGAGAAGC
TTGCTTCTCTTGAGAGCGGCGGACGGGTGAGTAATGCCTAGGAATCTGCCTGGTAGTGGGGGATAACGTCCGGAAACGGA
CGCTAATACCGCATACGTCCTACGGGAGAAAGCAGGGGACCTTCGGGCCTTGCGCTATCAGATGAGCCTAGGTCGGATTA
GCTAGTTGGTGAGGTAATGGCTCACCAAGGCGACGATCCGTAACTGGTCTGAGAGGATGATCAGTCACACTGGAACTGAG
ACACGGTCCAGACTCCTACGGGAGGCAGCAGTGGGGAATATTGGACAATGGGCGAAAGCCTGATCCAGCCATGCCGCGTG
TGTGAAGAAGGTCTTCGGATTGTAAAGCACTTTAAGTTGGGAGGAAGGGTACTTACCTAATACGTGAGTATTTTGACGTT
ACCGACAGAATAAGCACCGGCTAACTCTGTGCCAGCAGCCGCGGTAATACAGAGGGTGCAAGCGTTAATCGGAATTACTG
GGCGTAAAGCGCGCGTAGGTGGTTCGTTAAGTTGGATGTGAAATCCCCGGGCTCAACCTGGGAACTGCATCCAAAACTGG
CGAGCTAGAGTATGGTAGAGGGTGGTGGAATTTCCTGTGTAGCGGTGAAATGCGTAGATATAGGAAGGAACACCAGTGGC
GAAGGCGACCACCTGGACTGATACTGACACTGAGGTGCGAAAGCGTGGGGAGCAAACAGGATTAGATACCCTGGTAGTCC
ACGCCGTAAACGATGTCAACTAGCCGTTGGGAGCCTTGAGCTCTTAGTGGCGCAGCTAACGCATTAAGTTGACCGCCTGG
GGAGTACGGCCGCAAGGTTAAAACTCAAATGAATTGACGGGGGCCCGCACAAGCGGTGGAGCATGTGGTTTAATTCGAAG

CAACGCGAAGAACCTTACCAGGCCTTGACATCCAATGAACTTTCCAGAGATGGATTGGTGCCTTCGGGAACATTGAGACA
GGTGCTGCATGGCTGTCGTCAGCTCGTGTCGTGAGATGTTGGGTTAAGTCCCGTAACGAGCGCAACCCTTGTCCTTAGTT
ACCAGCACGTTATGGTGGGCACTCTAAGGAGACTGCCGGTGACAAACCGGAGGAAGGTGGGGATGACGTCAAGTCATCAT
GGCCCTTACGGCCTGGGCTACACACGTGCTACAATGGTCGGTACAGAGGGTTGCCAAGCCGCGAGGTGGAGCTAATCCCA
TAAAACCGATCGTAGTCCGGATCGCAGTCTGCAACTCGACTGCGTGAAGTCGGAATCGCTAGTAATCGCGAATCAGAATG
TCGCGGTGAATACGTTCCCGGGCCTTGTACACACCGCCCGTCACACCATGGGAGTGGGTTGCACCAGAAGTAGCTAGTCT
AACCTTCGGGAGGACGGTTACCACGGTGTGATTCATGACTGGGGTGAAGTCGTAACAAGGTAGCCGTAGGGGAACCTGCG
GCTGGATCACCTCCTTAAATGGACGACAACAAGAAGAAAGCCTTGGCTGCGGCCCTGGGTCAGATCGAACGTCAATTCGG
CAAGGGTGCCGTAATGCGTATGGGCGATCACGACCGCCAGGCGATCCCGGCCATTTCCACTGGCTCTCTGGGTCTGGACA
TCGCACTCGGCATCGGCGGCCTGCCAAAAGGCCGTATTGTTGAAATCTACGGTCCGGAATCGTCCGGTAAAACCACCCTG
ACCCTGTCGGTGATTGCCCAGGCACAGAAGATGGGCGCCACCTGCGCCTTCGTCGACGCCGAGCACGCACTGGACCCGGA
ATACGCAGGCAAACTGGGGGTCAACGTTGACGACCTGCTGGTTTCCCAGCCGGACACCGGCGAACAGGCGCTGGAAATCA
CCGACATGCTGGTGCGCTCCAATGCCATCGACGTGATCGTGATCGACTCCGTGGCGGCACTGGTACCCAAGGCCGAGATC
GAAGGCGAGATGGGCGACATGCACGTGGGCCTGCAGGCCCGCCTGATGTCCCAGGCGCTGCGCAAGATCACCGGTAACAT
CAAGAACGCCAACTGCCTGGTGATCTTCATCAACCAGATCCGTATGAAAATCGGCGTGATGTTCGGCAGCCCGGAAACCA
CCACCGGTGGTAACGCGCTGAAGTTCTACGCTTCGGTTCGTCTGGACATCCGTCGTACTGGCGCGGTGAAGGAAGGCGAC
GAAGTCGTCGGTAGCGAAACCCGGGTCAAGATCGTCAAGAACAAGGTGGCTCCACCGTTCCGTCAGGCTGAATTCCAGAT
CCTGTACGGCAAGGGTATCTACCTGAACGGCGAGATCATCGATCTGGGCGTGCTGCACGGTTTCCTCGAGAAGTCCGGTG
CCTGGTACAGCTACCAGGGCAACAAGATCGGTCAGGGCAAGGCCAACTCGGCCAAGTTCCTGCAGGACAATCCGGAAATC
GGCAATGCCCTCGAGAAGCAGATTCGCGACAAGCTGCTGGCTCCAACCGCTGATGTCAAAGCTTCGCCGGTCAACGAGAC
CATCGATGACATGGCTGACGCGGATATCTGAATGAGCGAAGAAAACACGTACGACTCGAGCAGCATTAAAGTGCTGAAAG
GTTTGGATGCCGTACGCAAACGTCCCGGTATGTACATTGGTGACACCGACGATGGCAGCGGTCTGCACCATATGGTGTTC
GAGGTGGTCGATAACTCGATCGACGAAGCTCTGGCCGGCCACTGCGACGACATCAGCATCATCATCCACCCGGACGAATC
CATTACCGTGCGTGACAACGGTCGCGGCATCCCGGTAGACGTGCATAAAGAAGAAGGCGTTTCCGCGGCCGAGGTCATCA
TGACTGTGCTGCACGCCGGCGGTAAGTTCGACGACAACTCCTACAAAGTATCCGGCGGTCTGCACGGTGTGGGTGTGTCG
GTAGTGAACGCCCTGTCCGAAGAACTGGTCCTGACCGTTCGCCGCAGTGGCAAGATCTGGGAACAGACCTACGTTCACGG
TGTGCCTCAGGCGCCTATGGCGATCGTCGGTGACAGTGAAACCACCGGTACCCAGATTCACTTCAAGGCTTCCAGCGAGA
CCTTCAAGAACATCCATTTCAGCTGGGACATCCTGGCCAAGCGGATTCGTGAACTGTCCTTCCTCAACTCCGGTGTCGGT
ATCGTTCTGAAGGACGAGCGTAGCGGCAAGGAAGAACTGTTCAAGTACGAAGGCGGTCTGCGTGCGTTCGTTGAATACCT
GAACACCAACAAGACCGCGGTCAACCAGGTGTTCCATTTCAATGTGCAGCGTGAAGATGGCATCGGCGTGGAAATCGCCC
TGCAGTGGAATGACAGCTTCAACGAAAACCTGCAGTGCTTCACCAACAACATTCCGCAGCGCGATGGCGGCACCCACCTG
GTGGGCTTCCGTTCGGCACTGACACGTAACCTGAACAACTACATCGAACAGGAAGGTCTGGCGAAGAAGCACAAGGTCGC
CACCACCGGTGACGATGCCCGCGAAGGCCTGACCGCAATCATTTCGGTCAAGGTGCCGGATCCGAAGTTCAGCTCCCAGA
CCAAAGACAAGCTGGTGTCTTCCGAAGTGAAGACCGCGGTCGAACAGGAAATGGGCAAGTACTTCTCCGACTTCCTGCTG
GAAAACCCGAACGAAGCCAAGCTGGTGGTCGGCAAGATGCTCGACGCCGCCCGTGCCCGTGAAGCGGCGCGTAAGGCTCG
TGAGATGACCCGCCGTAAAGGTGCGCTGGATATCGCCGGCCTGCCGGGCAAACTGGCGGACTGCCAGGAAAAAGACCCTG
CCCTTTCCGAACTCTACCTGGTGGAAGGTGACTCTGCTGGCGGCTCCGCCAAGCAGGGACGCAACCGTAAGACCCAGGCG
ATTCTGCCGCTCAAGGGCAAGATCCTTAACGTCGAGAAAGCGCGTTTCGACAAGATGATTTCCTCGCAAGAGGTCGGCAC
CTTGATCACTGCACTCGGTTGCGGCATCGGCCGCGAAGAGTACAACATCGACAAGCTGCGTTATCACAACATCATCATCA
TGACCGACGCCGACGTCGACGGTTCGCACATCCGTACCCTGCTGCTGACCTTCTTCTTCCGTCAGCTGCCGGAGCTGATC
GAGCGTGGCTACATCTACATCGCTCAACCGCCGCTGTACAAGGTCAAGAAAGGCAAGCAAGAGCAATACATCAAAGACGA
CGACGCCATGGAAGAGTACATGACGCAGTCGGCCCTGGAAGATGCGAGCCTGCACCTGAACGAAGACGCACCGGGTATTT
CCGGCGAGGCGCTGGAGCGCCTGGTGAACGACTTCCGCATGGTCATGAAAACCCTCAAGCGTCTGTCGCGCCTGTACCCT
CAGGAGCTGACCGAGCACTTCATCTACCTGCCGGCCGTGAGCCTGGAGCAGCTCTCCGATCACGCGGCGATGCAGGATTG
GCTGGCCCAATATGAAGTCCGCCTGCGCACCGTCGAGAAGTCCGGCCTGGTCTACAAGGCCAGCCTGCGTGAAGACCGTG
AACGTAATGTCTGGCTGCCAGAGGTCGAACTGATCTCCCACGGCCTGTCGAACTACGTCACCTTCAACCGTGACTTCTTC
GGCAGCAATGACTACAAGACCGTCGTCACCCTCGGTGCTCAACTGAGCTCCCTGCTGGACGAAGGCGCTTATATTCAGCG
TGGCGAACGCAAGAAGGCAGTGACCGAGTTCAAGGAAGCCCTGGACTGGCTGATGACCGAAAGTACCAAGCGCCACACCA
TCCAGCGATACAAAGGTCTGGGCGAGATGAACCCGGATCAGCTGTGGGAAACCACCATGGACCCAAGCGTGCGCCGTATG
CTCAAGGTCACCATCGAAGACGCCATCGGCGCCGACCAGATCTTCAACACCCTGATGGGTGATGCGGTCGAGCCTCGTCG
CGACTTCATCGAAAGCAACGCCCTGGCGGTATCCAACCTGGACTTCTGAATGTCCGGAAAAGCGCAACAGCAGTCTCGCC
TCAAAGAGTTGATCAGCCGTGGTCGTGAGCAGGGTTACCTGACTTACGCGGAGGTCAACGACCACCTGCCGGAGGATATT
TCAGATCCGGAACAGGTGGAAGACATCATCCGCATGATCAACGACATGGGGATCAACGTATTCGAGAGTGCTCCGGATGC
GGATGCCCTTTTGTTGGCCGAAGCCGATACCGACGAAGCAGCAGCTGAAGAAGCCGCCGCAGCGTTGGCGGCTGTGGAAA
CCGACATTGGTCGCACTACCGACCCCGTGCGTATGTACATGCGCGAAATGGGTACGGTAGAGCTGCTCACACGTGAAGGC
GAAATCGAAATCGCCAAGCGTATCGAAGAGGGCATCCGTGAAGTGATGGGCGCGATCGCGCACTTCCCTGGCACGGTTGA
GCACATCCTCTCCGAATACACTCGCGTCACCACCGAAGGTGGCCGCCTGTCCGACGTCCTGAGCGGTTACATCGACCCGG
ACGACGGTATTGCGCCGCCTGCCGCCGAAGTACCACCGCCTGTCGATGCCAAGGCCGCAAAAGCGGACGACGACACCGAC
GACGATGACGCCGAAGCCAGTGACGACGAAGAAGAAGCCGAAAGCGGTCCGGATCCGGTCATCGCAGCCCAGCGCTTTGG
CGCCGTTGCCGACCAGATGGAAATTACCCGCAAGGCGCTGAAAAAGCACGGTCGCGAACACAAGCAAGCCCTGGCTGAAA
TGCTGGCCCTGGCTGAGCTGTTCATGCCGATCAAACTGGTTCCGAAGCAATTCGAAGGCCTGGTTGAACGTGTTCGTAGT
GCCCTGGATCGCCTGCGTCAGCAAGAGCGCGCGATCATGCAGCTCTGTGTTCGTGATGCCCGCATGCCACGCGCCGACTT
CCTGCGCCAGTTCCCTGGCAATGAAGTGGACGAAAGCTGGTCCGACGCACTGGCCAAAGGCAAGGCCAAGTACGCCGAAG
CCATTGGCCGCCTGCAGCCGGACATCATCCGTTGCCAGCAGAAGCTGACCGCACTCGAGACCGAGACCGGCCTGACGATC

GCCGAGATCAAGGACATCAACCGTCGCATGTCGATCGGCGAGGCCAAGGCCCGTCGCGCGAAGAAAGAGATGGTCGAAGC
CAACTTGCGTCTGGTGATCTCCATCGCCAAGAAGTACACCAACCGTGGCCTGCAATTCCTCGACCTGATCCAGGAAGGCA
ACATCGGTTTGATGAAAGCAGTAGACAAGTTCGAATACCGTCGCGGCTACAAGTTCTCGACTTATGCCACCTGGTGGATC
CGTCAGGCGATCACTCGCTCGATCGCCGACCAGGCCCGCACCATCCGTATTCCGGTGCACATGATCGAGACGATCAACAA
GCTCAACCGTATTTCCCGGCAGATGTTGCAGGAAATGGGTCGCGAACCGACTCCGGAAGAGCTGGGCGAACGCATGGAAA
TGCCTGAGGACAAGATCCGCAAGGTATTGAAGATCGCTAAAGAGCCGATCTCCATGGAAACCCCGATCGGTGATGACGAA
GACTCCCATCTGGGTGACTTCATCGAAGACTCGACCATGCAGTCGCCAATCGATGTCGCCACCGTTGAGAGCCTTAAAGA
AGCGACTCGCGAAGTACTCTCCGGCCTCACTGCCCGTGAAGCCAAGGTACTGCGCATGCGCTTCGGCATCGACATGAATA
CCGACCACACCCTTGAGGAAGTCGGTAAGCAGTTCGATGTTACCCGTGAGCGGATTCGTCAGATCGAAGCCAAGGCGCTG
CGCAAGCTGCGCCACCCGACGCGAAGCGAGCATCTGCGCTCCTTCCTCGACGAGTAATTGACTAAGCCAGCCATACTCGC
CCTTGCTGATGGCAGCATTTTTCGCGGCGAAGCCATTGGAGCCGACGGTCAAACCGTTGGTGAGGTGGTGTTTAACACCG
CAATGACCGGCTATCAGGAAATCCTTACCGATCCTTCCTACGCCCAACAGATCGTTACCCTGACTTACCCACATATCGGC
AATACCGGCACCACGCCGGAAGACGCCGAGTCCGATCGTGTCTGGTCGGCCGGTCTGGTGATTCGCGACCTGCCACTGGT
TGCGAGCAACTGGCGTAACACCTTGTCCCTGTCCGACTACCTGAAAGCCAACAATGTTGTGGCGATCGCCGGTATCGACA
CCCGTCGTCTGACGCGCATCCTGCGCGAGAAAGGCGCGCAGAACGGCTGCATCATGGCCGGCGACAATATCTCCGACGAA
GCGGCGATTGCCGCTGCGCGCGGCTTCCCGGGCCTGAAAGGCATGGATCTGGCGAAGGTCGTCAGCACCAAGGAAAGCTA
CGAGTGGCGCTCCAGCGTCTGGAGCCTGAAGACCGACAGTCACCCGACCATCGAGGCTTCCGAGCTGCCTTACCACGTGG
TTGCCTACGACTACGGCGTCAAGCTGAACATCCTGCGCATGCTGGTCGAGCGCGGTTGCCGCGTGACCGTGGTACCTGCG
CAAACCCCGGCCAGCGACGTCCTGGCGCTCAAGCCTGACGGTGTGTTCCTGTCCAACGGTCCTGGCGACCCCGAGCCTTG
CGATTACGCCATCCAGGCGATCAAGGACGTGCTGGAAACCGAGATTCCGGTCTTCGGTATCTGCCTGGGCCACCAACTGC
TGGCGCTGGCCGCCGGCGCCAAGACAGTGAAGATGGGCCACGGCCACCACGGTGCCAACCACCCGGTCCAGGACCTGGAC
AGCGGTGTAGTGATGATCACCAGCCAGAACCACGGTTTTGCGGTGGACGAAACCACCCTGCCGGGCAACGTGCGGGCGAT
CCACAAGTCGCTGTTCGACGGCACCCTGCAAGGCATCGAGTTGACCGACAAGAGCGCATTCAGCTTCCAGGGCCACCCTG
AAGCGAGCCCGGGCCCGAACGATGTGGCGCCGCTGTTCGATCGTTTCATCAACGAGATGGCCAAGCGACGCTGAATGAGT
AGCGGACGTATCGTTCAAATCATCGGCGCCGTTATCGACGTGGAATTTCCACGCGACAGCGTACCGAGCATCTACGACGC
CTTGAAGGTTCAAGGCGCCGAAACCACTCTGGAAGTTCAGCAGCAGCTGGGCGACGGCGTGGTACGTACCATTGCGATGG
GCTCCACCGAGGGCTTGAAGCGCGGTCTGGACGTCAACAACACTGGCGCAGCCATCTCCGTACCGGTCGGTAAAGCGACC
CTGGGCCGGATCATGGACGTACTGGGCAACCCGATCGACGAAGCTGGCCCGATCGGCGAAGAAGAGCGTTGGGGCATTCA
CCGTCCTGCGCCGACCTTCGCTGAACAAGCTGGCGGCAACGACCTGCTGGAAACCGGCATCAAGGTTATCGACCTGGTTT
GCCCGTTCGCCAAGGGCGGTAAAGTCGGTCTGTTCGGTGGTGCCGGTGTGGGCAAAACCGTAAACATGATGGAACTGATC
CGTAACATCGCCATCGAGCACAGCGGTTATTCCGTGTTCGCCGGTGTGGGTGAGCGTACTCGTGAGGGTAACGACTTCTA
CCACGAGATGAAGGATTCCAACGTTCTGGACAAAGTGGCACTGGTATACGGCCAGATGAACGAGCCGCCGGGAAACCGTC
TGCGCGTAGCTCTGACCGGCCTGACCATGGCCGAGAAGTTCCGTGACGAAGGTAACGACGTTCTGCTGTTCGTCGACAAC
ATCTATCGTTACACCCTGGCCGGTACCGAAGTATCCGCACTGCTGGGCCGTATGCCTTCGGCAGTAGGTTACCAGCCGAC
CCTGGCTGAAGAGATGGGCGTTCTGCAAGAACGTATCACTTCGACCAAGCAAGGCTCGATCACCTCGATCCAAGCGGTAT
ACGTGCCTGCGGACGACTTGACCGACCCGTCGCCAGCGACCACCTTCGCCCACTTGGACGCCACCGTCGTTCTGTCCCGT
GACATCGCTTCCCTGGGTATCTACCCAGCGGTAGACCCACTGGACTCGACTTCCCGTCAGCTGGACCCGAACGTGATCGG
CAACGAGCACTACGAAACCGCTCGCGGCGTTCAGTACGTGCTGCAGCGCTACAAAGAGCTGAAGGACATCATTGCGATCC
TGGGTATGGACGAACTGTCCGAAGCCGACAAGCAACTGGTATCCCGCGCTCGTAAGATCCAGCGCTTCCTGTCGCAGCCG
TTCTTCGTGGCTGAAGTCTTCACTGGTTCTCCAGGCAAATACGTTTCCCTGAAAGACACCATCGCTGGCTTCAAAGGCAT
CCTCAACGGTGACTACGACCATCTGCCAGAACAAGCGTTCTACATGGTTGGTGGCATCGAAGAAGCGATCGAGAAAGCCA
AGAAACTGTAA

NCBI Reference Sequence: NZ_LT799039.1
Strain: KT2440
Chained genes: 16S rRNA, recA, gyrB, rpoD, carA, atpD
>NZ_LT799039_KT2440
GAACTGAAGAGTTTGATCATGGCTCAGATTGAACGCTGGCGGCAGGCCTAACACATGCAAGTCGAGCGGATGACGGGAGC
TTGCTCCTTGATTCAGCGGCGGACGGGTGAGTAATGCCTAGGAATCTGCCTGGTAGTGGGGGACAACGTTTCGAAAGGAA
CGCTAATACCGCATACGTCCTACGGGAGAAAGCAGGGGACCTTCGGGCCTTGCGCTATCAGATGAGCCTAGGTCGGATTA
GCTAGTTGGTGGGGTAATGGCTCACCAAGGCGACGATCCGTAACTGGTCTGAGAGGATGATCAGTCACACTGGAACTGAG
ACACGGTCCAGACTCCTACGGGAGGCAGCAGTGGGGAATATTGGACAATGGGCGAAAGCCTGATCCAGCCATGCCGCGTG
TGTGAAGAAGGTCTTCGGATTGTAAAGCACTTTAAGTTGGGAGGAAGGGCAGTAAGTTAATACCTTGCTGTTTTGACGTT
ACCGACAGAATAAGCACCGGCTAACTCTGTGCCAGCAGCCGCGGTAATACAGAGGGTGCAAGCGTTAATCGGAATTACTG
GGCGTAAAGCGCGCGTAGGTGGTTTGTTAAGTTGGATGTGAAAGCCCCGGGCTCAACCTGGGAACTGCATCCAAAACTGG
CAAGCTAGAGTACGGTAGAGGGTGGTGGAATTTCCTGTGTAGCGGTGAAATGCGTAGATATAGGAAGGAACACCAGTGGC
GAAGGCGACCACCTGGACTGATACTGACACTGAGGTGCGAAAGCGTGGGGAGCAAACAGGATTAGATACCCTGGTAGTCC
ACGCCGTAAACGATGTCAACTAGCCGTTGGAATCCTTGAGATTTAGTGGCGCAGCTAACGCATTAAGTTGACCGCCTGG
GGAGTACGGCCGCAAGGTTAAAACTCAAATGAATTGACGGGGGCCCGCACAAGCGGTGGAGCATGTGGTTTAATTCGAAG
CAACGCGAAGAACCTTACCAGGCCTTGACATGCAGAGAACTTTCCAGAGATGGATTGGTGCCTTCGGGAACTCTGACACA
GGTGCTGCATGGCTGTCGTCAGCTCGTGTCGTGAGATGTTGGGTTAAGTCCCGTAACGAGCGCAACCCTTGTCCTTAGTT
ACCAGCACGTAATGGTGGGCACTCTAAGGAGACTGCCGGTGACAAACCGGAGGAAGGTGGGGATGACGTCAAGTCATCAT
GGCCCTTACGGCCTGGGCTACACACGTGCTACAATGGTCGGTACAGAGGGTTGCCAAGCCGCGAGGTGGAGCTAATCTCA
CAAAACCGATCGTAGTCCGGATCGCAGTCTGCAACTCGACTGCGTGAAGTCGGAATCGCTAGTAATCGCGAATCAGAATG

TCGCGGTGAATACGTTCCCGGGCCTTGTACACACCGCCCGTCACACCATGGGAGTGGGTTGCACCAGAAGTAGCTAGTCT
AACCTTCGGGGGGACGGTTACCACGGTGTGATTCATGACTGGGGTGAAGTCGTAACAAGGTAGCCGTAGGGGAACCTGCG
GCTGGATCACCTCCTTAAATGGACGACAACAAGAAGCGCGCCTTGGCTGCGGCCCTGGGTCAGATCGAACGCCAATTCGG
TAAAGGCGCGGTCATGCGCATGGGTGACCATGAGCGTCAAGGCATTCCGGCCATCTCCACCGGCTCGCTGGGGCTGGATA
TCGCCCTGGGCATCGGCGGTCTGCCAAAAGGCCGTATCGTCGAGATCTACGGCCCGGAATCGTCGGGTAAGACCACGCTG
ACCCTGTCTGTCATCGCTGAAGCGCAAAAGAACGGTGCTACCTGCGCCTTCGTCGACGCCGAACATGCCCTTGACCCTGA
GTACGCCGGCAAGCTGGGCGTCAACGTCGATGACCTGCTGGTTTCGCAGCCGGACACCGGTGAGCAGGCCCTTGAAATCA
CCGACATGCTGGTGCGTTCCAACGCGGTCGACGTGATCATTGTCGACTCCGTTGCCGCGCTGGTACCGAAGGCCGAGATC
GAAGGCGAGATGGGTGACATGCATGTGGGCCTGCAGGCCCGCCTGATGTCCCAGGCACTGCGTAAGATCACCGGTAACAT
CAAGAACGCGAACTGCCTGGTCATCTTCATCAACCAGATCCGTATGAAGATCGGTGTGATGTTCGGCAGCCCGGAAACCA
CTACCGGTGGTAACGCCCTGAAGTTCTATGCTTCGGTCCGTCTGGACATCCGCCGTACCGGCGCGGTCAAGGAAGGCGAC
GAAGTGGTCGGCAGCGAAACCCGCGTCAAGATCGTCAAGAACAAGGTCTCGCCTCCGTTCCGTCAGGCTGAGTTCCAGAT
TCTTTACGGGAAAGGTATCTACCGTAACGGCGAGATCATTGATCTGGGGGTATCTCAGGGCCTGGTCGAAAAGTCCGGCG
CCTGGTACGCCTACCAAGGCAACAAGATCGGTCAAGGCAAAGCCAACGCTGCCAAGTACCTGGCTGAGAACCCGGCTATT
GGTGCCGAGATCGAGAAGCAGATTCGTGAGAAGTTGCTGAAAGCGGGTGCTGCTGCTGAAGCCGGCAAAGCTGCTGCTGC
TGAAGCTGATGCCGATGACATGGCCGACGCTGACGCCGGTTATTGAATGAGCGAAAATCAAACGTACGACTCCTCCAGCA
TCAAGGTGCTGAAAGGTTTGGATGCCGTACGCAAGCGTCCCGGCATGTACATTGGCGACACCGATGATGGTAGTGGCCTG
CACCACATGGTCTTCGAGGTGGTCGACAACTCGATCGACGAAGCCCTCGCCGGTCACTGCGATGACATTACCGTCATCAT
CCACCCGGACGAATCTATCAGTGTGCGCGACAACGGTCGCGGCATTCCGGTCGATGTGCATAAGGAAGAAGGCGTTTCCG
CAGCCGAGGTCATCATGACTGTGCTGCACGCCGGCGGTAAGTTTGACGACAACTCCTACAAAGTATCCGGCGGTCTGCAC
GGTGTAGGTGTGTCGGTTGTGAACGCCCTGTCCGAGAAGCTGGTTTTGACTGTTCGCCGTAGCGGCAAGATCTGGGAACA
GACTTACGTTCACGGTGTTCCACAAGCGCCCATGGCGGTTGTCGGTGACAGTGAAACCACGGGTACCCACATCCACTTCA
AGCCATCGGCTGAAACCTTCAAGAACATTCACTTCAGCTGGGACATCCTGGCCAAGCGCATCCGCGAGCTGTCGTTCCTC
AACTCGGGCGTTGGCATTCTGCTGAAGGATGAGCGCAGCGGTAAGGAAGAGTTCTTCAAGTACGAAGGCGGTCTGCGTGC
GTTCGTCGAGTACTTGAACACCAACAAGACGCCGGTCAACTCCCAGGTGTTCCACTTCAACGTTCAGCGTGACGATGGCG
TGGGTGTTGAAGTCGCCCTGCAATGGAACGACAGCTTCAACGAAAACCTGCTGTGCTTTACCAACAATATTCCGCAGCGT
GATGGCGGTACCCACCTGGTGGGTTTCCGTTCCTCGCTGACCCGTAGCCTTAACAGCTACATCGAGCAGGAAGGCCTGGC
CAAGAAGAACAAGGTGGCAACCACTGGCGACGACGCCCGTGAAGGCCTGACCGCGATCATCTCGGTGAAGGTACCGGACC
CGAAGTTCAGCTCGCAGACCAAGGACAAGCTGGTCTCCTCGGAGGTGAAAACCGCCGTGGAACAGGAGATGAACAAGTAC
TTCGCCGATTTCCTCCTGGAAAACCCGAACGAGGCGAAGGCCGTCGTTGGCAAGATGATCGACGCCGCTCGCGCCCGTGA
AGCCGCCCGTAAAGCCCGTGAGATGACCCGCCGTAAAGGTGCGCTGGATATCGCGGGTCTGCCGGGCAAGCTGGCCGACT
GCCAAGAGAAGGATCCTGCTCTCTCCGAACTGTACCTGGTGGAGGGTGACTCCGCGGGTGGCTCGGCCAAGCAAGGCCGC
AACCGTCGTACCCAGGCGATCTTGCCGCTGAAGGGTAAAATCCTCAACGTCGAGAAAGCGCGCTTCGACAAGATGATTTC
GTCCCAGGAAGTGGGCACGCTGATCACTGCGCTGGGCTGTGGCATCGGCCGCGAAGAGTACAACATCGACAAACTGCGTT
ATCACAACATCATCATCATGACCGATGCTGACGTTGACGGTTCGCACATCCGTACGCTGCTGCTGACCTTCTTCTTCCGT
CAGCTGCCGGAGCTGGTCGAGCGTGGCTACATCTATATTGCCCAGCCGCCGCTGTACAAGGTGAAACGAGGCAAGCAGGA
GCAGTACATCAAGGACGATGAGGCCATGGAAGAGTACATGACCCAGTCGGCTCTGGAAGATGCCAGCCTGCACCTGGACG
AATCGGCGCCAGCAGTTTCCGGCGTGCAGCTGGAAGCGCTGGTGAATGAGTTCCGTAGTGTCATGAAGACTCTCAAGCGC
CTGTCGCGCTTGTACCCGGAAGAGCTGACCGAGCACTTCGTCTACCTGCCTGAGGTGACCCTGGAGCAGTTGGGTGACCA
CGCAGTGATGCAGGCCTGGCTGGCCCAGTTCCAGGCGCGTCTGAACTCCAGCCAGAAGTCTGGCCTGGCTTACAACGCCA
GCCTGCGTGAAGACAAAGAGCGCAACGTATGGCTGCCTGAAGTGGAAATTACCTCTCACGGTCTGGCCAGCTACATCACC
TTCAACCGCGATTTCTTCGGCAGCAATGACTACCGTACCGTAGTCAACATTGGTGCCAAGCTTTCGAGCCTGTTGGGTGA
AGGTGCGTACGTGCAGCGCGGTGAACGCCGCAAGGCAATCGTCGAGTTCAAAGAGGGCCTGGATTGGCTGATGAACGAGA
CCACCAAGCGCCATACGATTCAGCGATACAAAGGGCTGGGTGAGATGAACCCGGATCAACTGTGGGAAACCACCATGGAC
CCGACCGTTCGCCGTATGCTCAAGGTCACGATCGAAGATGCTATCGCCGCTGACCAGATCTTCAACACCCTGATGGGTGA
TGCGGTCGAGCCGCGTCGTGACTTCATCGAAAGTAACGCGCTGTCGGTGTCGAACCTGGACTTCTGAATGTCCGGAAAAG
CGCAACAGCAGTCTCGTATCAAAGAGTTGATCACCCGCGGTCGTGAGCAGGGCTACCTGACTTACGCGGAGGTCAACGAC
CACCTGCCTGAGGATATTTCAGATCCGGAACAGGTGGAAGACATCATCCGCATGATCAACGACATGGGGATCAACGTATT
CGAGAGTGCTCCGGATGCGGATGCCCTTCTGTTGGCGGAAGCCGACACCGATGAAGCCGCGGCCGAAGAAGCCGCTGCTG
CATTGGCGGCAGTTGAAACCGATATCGGCCGCACGACCGACCCGGTGCGCATGTATATGCGCGAAATGGGTACTGTCGAG
CTGCTGACCCGCGAAGGCGAGATCGAAATCGCCAAGCGTATCGAGGAAGGCATTCGTGAAGTCATGGGCGCCATCGCCCA
CTTCCCGGGCACTGTCGACTACATTCTCGGCGAATATGACCGCGTCACCACCGAGGGTGGCCGCCTGTCGGACGTTCTCA
GCGGTTACATCGACCCTGACGACAACATTGCCGCGCCAACCGAAGAAGTGCCGATCCCAGGTGCCAAGGCCGCTGCGGCG
AAGGAAGAGTCCGACGACGACGAAGAAGAATCCGAAAGCGGTGACGACGAGGAAGAGGCCGAGAGCGGCCCGGATCCGGT
CGTCGCAGCCCAACGCTTCGGTGCGGTATCCGATCAGCTTCAGGCAACCTCCAAGGTCCTGAAGAAAAATGGTCGCAACC
ACAAGGAAAGCATCGAGGCCCTGCAGGCCCTGGCTGACCTGTTCATGCCGATCAAGCTGGTACCGAAGCAGTTCGAGGTA
CTGGTCGAGCGTGTCCGTGACGCCCTGAACCGTCTGCGACAGCAAGAACGCGCCATCATGCAGCTGTGCGTACGTGACGC
CCGCATGCCGCGAGCCGACTTCCTGCGCATGTTCCCAAGCAACGAAACCGACCAGACCTGGAGCGGTGACCTGGCCAAGC
GCAACACCAAGTGGGCTGCCGCCCTGGGTGAAAAGAACGCTGCCATCGTCGCTTGCCAACAGAAGCTGATCGACCTTGAG
ACCGAAACCGGCCTGACCGTTGCCGAGATCAAGGAAATCAACCGTCGCATGTCGATCGGTGAAGCCAAGGCCCGCCGCGC
CAAGAAAGAAATGGTCGAGGCGAACCTGCGTCTGGTGATTTCCATCGCCAAGAAGTACACCAACCGTGGCCTGCAGTTCC
TCGACCTGATCCAGGAAGGCAACATCGGTCTGATGAAAGCGGTGGACAAGTTCGAATACCGTCGCGGCTACAAGTTCTCG
ACCTATGCCACCTGGTGGATCCGTCAGGCGATCACCCGTTCGATCGCCGACCAGGCACGCACCATCCGCATTCCGGTGCA
CATGATCGAGACGATCAACAAGCTCAACCGTATTTCCCGCCAGATGCTGCAGGAAATGGGTCGCGAACCGACTCCGGAAG

AGCTGGGCGAGCGCATGGAAATGCCTGAGGACAAAATCCGCAAGGTATTGAAGATCGCCAAAGAGCCGATCTCCATGGAA
ACCCCGATCGGTGACGACGAAGATTCGCACCTGGGCGACTTCATCGAGGACTCGACCATGCAGTCCCCGATCGACGTGGC
CACGGTCGAAAGCCTCAAGGAAGCCACCCGTGACGTGCTCTCGGGCCTGACCGCACGTGAAGCCAAGGTGCTGCGCATGC
GTTTCGGTATCGACATGAACACCGACCACACCCTCGAAGAGGTGGGCAAGCAGTTTGACGTAACGCGTGAGCGGATCCGT
CAGATCGAAGCGAAGGCGTTGCGCAAGCTGCGCCACCCGACTCGCAGCGAGCACCTGCGCTCCTTCCTCGACGAGTGATT
GACAAAGCCAGCCATACTCGCCCTTGCCGACGGCAGTATTTTCCGCGGTGAAGCCATCGGTGCCGACGGTCAGACCGTTG
GTGAGGTGGTATTCAACACCGCTATGACCGGCTACCAGGAAATCCTTACAGACCCTTCCTACGCGCAGCAAATCGTTACC
CTGACCTACCCGCACATCGGCAACACCGGTACTACCCCGGAAGACGCCGAGTCGAGCCGCGTCTGGTCCGCTGGCCTGGT
CATCCGTGACCTGCCGCTGCTGGCCAGCAACTGGCGTAACACCCAGTCGCTGCCTGAGTACCTCAAGGCCAACAACGTCG
TCGCCATCGCCGGCATCGACACCCGTCGCCTGACCCGTATCCTGCGTGAAAAGGGCGCCCAGAACGGCTGCATTCTGGCG
GGTGACAACATCAGCGAAGAAGCTGCCATCGCTGCTGCCCGCGGCTTCCCGGGCCTGAAGGGCATGGACCTGGCCAAGGT
CGTCTCCACCAAGGAACGTTACGAGTGGCGCTCCAGCGTGTGGGAGCTGAAAACCGACAGCCACCCGACCATCGACGCTG
CCGACCTGCCGTACCACGTGGTTGCCTTCGACTATGGCGTCAAGCTGAACATCCTGCGCATGCTGGTGGCCCGCGGCTGC
CGCGTGACCGTGGTACCAGCCCAGACCCCGGCCAGCGAAGTACTGGCACTCAACCCGGACGGCGTGTTCCTGTCCAACGG
CCCTGGTGACCCTGAGCCGTGCGACTACGCGATCCAGGCGATCAAGGAAATCCTCGAAACCGAGATCCCGGTATTCGGCA
TCTGCCTCGGCCACCAGCTGCTGGCCCTGGCGTCCGGCGCCAAGACCGTGAAAATGGGCCACGGCCACCACGGTGCCAAC
CACCCGGTCCAGGACCTGGATACTGGTGTGGTCATGATCACCAGCCAGAACCACGGTTTCGCCGTTGACGAGGCGACCCT
GCCGGGCAACGTTCGCGCCATTCACAAGTCGCTGTTCGACGGCACCCTGCAGGGTATCGAGCGTACCGACAAGAGCGCGT
TCAGCTTCCAGGGCCACCCTGAAGCGAGCCCGGGCCCGACCGACGTCGCGCCTCTGTTCGATCGTTTCACCGATGCCATG
GCCAAGCGCCGCTGAATGAGTAGCGGACGTATCGTTCAAATCATCGGCGCCGTCATCGACGTGGAATTCCCACGTGACGT
CGTGCCGAGTGTATACAACGCGCTTAAAGTACAAGGCGCGGAAACCACCCTGGAAGTTCAGCAGCAGCTGGGCGACGGCG
TGGTTCGTACCATTGCGATGGGCTCGACCGAAGGCCTGAAGCGCGGTCTGGATGTCGTCGACACCGGCGCTGCCATTTCC
GTTCCAGTTGGTAAGGCCACCCTGGGCCGTATCATGGACGTACTGGGCAACCCGATCGACGAAGCCGGCCCGATCGGCGA
AGAAGAGCGTCGCGGTATCCACCAGCCAGCGCCTTCGTTCGCTGACCAGGCAGGCGGCAACGACCTGCTGGAAACCGGCA
TCAAGGTTATCGACCTGGTTTGCCCGTTCGCCAAGGGTGGTAAGGTTGGTCTGTTCGGTGGTGCCGGTGTCGGCAAGACC
GTAAACATGATGGAACTGATCCGTAACATCGCCATGGAACACAGCGGTTACTCCGTGTTCGCTGGTGTGGGTGAGCGTAC
TCGTGAGGGTAACGACTTCTACCACGAGATGAAGGACTCCAACGTTCTCGACAAAGTAGCGCTGGTCTACGGTCAGATGA
ACGAGCCACCAGGAAACCGTCTGCGTGTAGCGCTGACCGGCCTGACCATGGCCGAGAAGTTCCGTGACGAAGGTAACGAC
GTTCTGCTGTTCGTCGACAACATCTATCGTTACACCCTGGCCGGTACCGAAGTATCCGCACTGCTGGGCCGTATGCCTTC
GGCAGTAGGTTACCAGCCGACCCTGGCTGAAGAGATGGGCGTTCTGCAAGAGCGCATCACCTCCACCAAGGAAGGTTCGA
TCACCTCCGTACAGGCCGTATACGTACCTGCGGACGACTTGACCGACCCGTCGCCAGCGACCACCTTCGCCCACTTGGAC
GCCACCGTCGTTCTGTCCCGTGACATCGCCTCCCTGGGTATCTACCCAGCGGTCGACCCACTGGACTCGACTTCGCGCCA
GCTGGACCCGAACGTGATCGGCAACGAGCACTACGAGACCGCTCGTGGCGTTCAGTATGTTCTGCAGCGCTATAAAGAGC
TGAAGGACATCATTGCGATCCTGGGTATGGACGAACTGTCCGAAGCCGACAAGCAACTGGTAGCCCGCGCTCGTAAGATC
CAGCGCTTCCTGTCGCAGCCGTTCTTCGTGGCAGAAGTCTTCACCGGTTCGCCAGGCAAGTACGTTTCCCTGAAAGACAC
CATCGCTGGCTTCAGCGGCATCCTCAAAGGTGACTACGACCACCTGCCAGAACAAGCGTTCTACATGGTCGGCAGCATCG
ACGAAGCGATCGAGAAAGCCAAGAAACTGTAA
